# The valine-arginine dipeptide repeat protein encoded by mammalian telomeric RNA appears highly expressed in mitosis and may repress global translation

**DOI:** 10.1101/2024.07.24.604971

**Authors:** Taghreed M. Al-Turki, Venkata Mantri, Smaranda Willcox, C. Allie Mills, Laura E. Herring, Su-Ji Cho, Hannah Lee, Cailyn Meyer, E. S. Anton, Jack D. Griffith

## Abstract

Translation of mammalian telomeric G-rich RNA via the Repeat Associated non-AUG translation mechanism can produce two dipeptide repeat proteins: repeating valine-arginine (VR) and repeating glycine-leucine (GL). Their potentially toxic nature suggests that one or both must play a needed role in the cell. Using light microscopy combined with antibody staining we discovered that cultured human cells stain brightly for VR during mitosis with VR staining co-localizing with ribosomes. *In vitro*, VR protein represses translation in a firefly luciferase assay. Affinity purification combined with mass spectrometry identified ribosomal proteins as the major class of VR interacting proteins. Extension to mouse embryonic cerebral cortical development showed strong staining in the ventricular zone where high mitotic index neural progenitor cells proliferate and in the cortical plate where new neurons settle. These observations point to VR playing a key role in mitosis very possibly depressing global translation, a role mediated by the telomere.

**Teaser:** The telomeric valine-arginine dipeptide repeat protein is highly expressed in mitotic
cells in culture and in mouse embryonic neural tissue.

## INTRODUCTION

For decades, telomeres have been assumed to be largely passive elements at chromosome ends, serving only to block end-to-end fusions and undergo slow replication-dependent attrition until senescence is triggered. The 6 nucleotide repeat in telomeric DNA is transcribed into a G-rich RNA (UUAGGG)n termed TERRA. TERRA plays important structural roles at the telomere (*1, 2*), but the simple repeat, and lack of canonical start signals argued against its translation into protein(s). However, in studies of expanded nucleotide repeat diseases in which the RNA transcripts are known to form stable secondary structures such as hairpins or G quartets, Zu et al (*3*) described a novel translation mechanism termed Repeat Associated non-AUG translation (RAN). RAN allows ribosomes to load onto these secondary structures and initiate translation at any nucleotide in the repeat, independent of the start codon ATG. This results in multiple amino acid homopolymers (triplet repeats), as observed in diseases such as SCA8 and Myotonic Dystrophy type 1 (*4*) or dipeptide repeat proteins (6 nt repeats) as seen in the inherited form of Amyotrophic Lateral Sclerosis/Frontotemporal Dementia (ALS/FTD) (*5*–*7*). We recently reported that RAN translation of mammalian TERRA will produce two dipeptide repeat proteins, repeating valine-arginine (VR) and repeating glycine-leucine (GL) (*8*).

An expanded 6 nt repeat (CCGGGG)n at the orf72 locus on chromosome 9 is the underlying cause of inherited ALS/FTD. RAN translation of the RNA transcript produces five dipeptide repeat proteins, GA, GP, and PA, which are hydrophobic and PR, and GR which are highly charged (*5*– *7*). GA forms long amyloid-like protein filaments *in vitro*, (*8, 9*) and *in vivo* they are associated with amyloid deposits in the brain of ALS/FTD patients (*9*). GR and PR which are highly charged are toxic when added directly to cells or expressed from plasmids in cultured cells (*10*–*15*). Zhang et al (*14*) expressed a plasmid encoding a GR dipeptide with 100 repeats in mice resulting in memory deficiency, brain atrophy, and age-dependent neurodegeneration. These changes paralleled an accumulation of diffuse GR in the cytoplasm which over time changed into large aggregates and inclusions. They also noted that the diffuse and aggregated forms of GR colocalize with ribosomes in mice. Other studies showed that when expressed from plasmids, GR will compromise mitochondrial function and increase oxidative stress in motor neurons (*16, 17*). PR expressed from plasmids forms toxic nuclear aggregates that result in neuronal death (*13*) and PR is seen localized to nucleoli where it impedes RNA biogenesis (*18*). A recent cryo-EM study from Loveland et al (*19*) demonstrated that PR binds to the exit tunnel in the large ribosomal subunit, repressing translation.

In our recent study we generated a polyclonal antibody to repeating VR dipeptide protein and verified its specificity by demonstrating that it reacts with VR but not GL or other repeating dipeptides. Further, upon expression of VR_60_ in cells from a plasmid containing conventional start and stop codons with a Flag tag, examination by light microscopy showed nuclear inclusions in which 85% of the Flag and VR signals co-localized (*8*). Employing the VR antibody and a labelled secondary antibody together with laser scanning confocal light microscopy, we observed that VR is more abundant in an osteosarcoma cell line (U2OS) and cells from a patient with a telomere biology disease known as Immunodeficiency-Centromeric instability-Facial anomalies syndrome (ICF). Both cell lines have elevated TERRA levels as contrasted to primary human foreskin cells. The staining was diffuse in the three cell lines and predominantly nuclear. Telomere disruption employing an oligonucleotide to reduce TERRA levels and a lentivirus targeting the telomere factor TRF2, led to higher levels of staining and large nuclear inclusions that stained brightly with the VR antibody.

Recent work has revealed how TERRA can move from the nucleus to the cytoplasm where it could undergo RAN translation. With the nuclear membrane intact, TERRA is sequestered in the nucleus in three forms: 1) in telomeric DNA:RNA hybrids, 2) associated with heterochromatin, and 3) free in the nucleoplasm (*1*). During cell division, TERRA hybrids tether the two sister telomeres together and overexpression of RnaseH results in loss of telomere cohesion during mitosis (*20*). In cycling cells, the generation of dysfunctional telomeres via knockdown of TRF2 results in TERRA being released into the cytoplasm resulting in mitochondrial dysfunction and activation of innate immune pathways (*21*). While experiments designed to examine the effect of eliminating VR by knocking out TERRA would be valuable, experimentally, the greatest reduction in TERRA achieved by us or others is at most 40% (*8, 22*) since TERRA is transcribed from most all telomeres.

It is relevant to ask why VR and GL proteins were previously missed. Our studies (*8*) and those from the ALS/FTD field (*6*) found that this family of proteins aggregates severely. They do not enter standard SDS gels remaining at the top with other aggregated proteins. Moreover, the nature of the RAN process is expected to generate a spectrum of polypeptide sizes as contrasted to a single species. This and the high content of arginine in PR, GR, and VR lead to their being missed in mass spectrometry studies using traditional bottom-up proteomics methods. Finally, the lack of canonical start and stop codons in the RNA prior to the discovery of the RAN mechanism, argued against their production. Thus, VR and GL may be members of the “dark proteome” encompassing many previously overlooked micro-proteins (*23*).

## RESULTS

### The VR dipeptide repeat protein shows strong diffuse staining in mitotic cells

The toxic nature of PR and GR and their similarity to telomeric VR suggested that for VR to have been retained in mammalian cells it must play some important functional role in the cell. During a routine staining of cycling U2OS cells with the VR antibody, we observed a field (Fig. 1A) in which most of the cells showed only DAPI staining (blue) for nuclear DNA. However, two cells in the field undergoing mitosis, stained brightly (green) for VR. Imaging of cycling primary human foreskin cells (Fig. 1B) revealed similar patterns of brightly staining cells engaged in cell division.

**Fig. 1.**
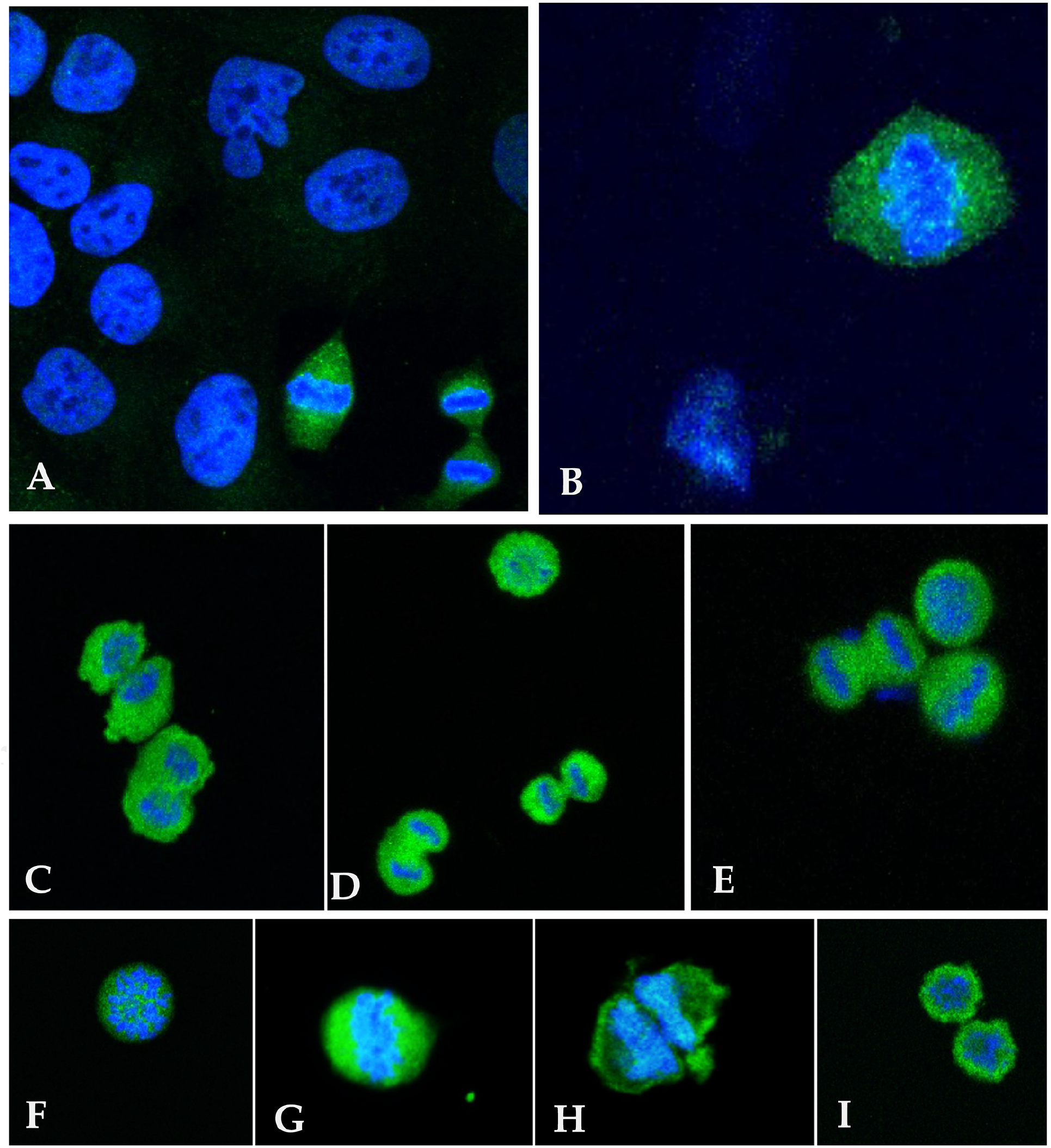
Visualization of VR dipeptide repeat protein in cultured mitotic cells. Examination of cycling U2OS (A) and primary foreskin cells (B) in culture occasionally show cells undergoing cell division. Staining with an antibody to the VR dipeptide repeat protein (green) and DAPI (blue) for DNA revealed strong staining for VR in mitotic cells but not in adjacent non-dividing cells (A). (C-D) U2OS cells were blocked in G2 with the drug RO-3306 and released into mitosis, followed by shake-off of mitotic cells and preparation for light microscopy (Methods). Scoring fields of cells, 41% (n=177), 43% (n=186), 11% (n=49), and 6% (n=25) were in prophase, metaphase, anaphase, and telophase respectively. Staining as in A, B. Examples of stained mitotic U2OS cells in: (F) prophase, (G) metaphase, (H) anaphase and (I) telophase.

To pursue this observation further, we treated 50% confluent U2OS cells for 16 hours with 9 μM CDK-1 kinase inhibitor (RO-3306) to block cells in late G2. The drug was removed, and cells were incubated for 45 min to allow them to progress into mitosis. Mitotic cells were isolated by mechanical shake-off, collected and replated on microscopy slides followed by fixation, incubation with the primary VR antibody, and finally incubation with an AlexaFluor 488 conjugated secondary antibody (Methods). The slides were then examined by laser confocal scanning microscopy. Examples of cells in prophase, metaphase, anaphase, and telophase are shown in Fig. 1 C-G. One hundred percent of the mitotic cells stained very brightly with the VR antibody. These experiments verified the observation of brightly staining cycling cells caught in mitosis. Unlike the varied size dots and punctate forms of VR we observed in nuclei of cycling cells, VR was diffusely distributed throughout the mitotic cell in which both nuclear and cytoplasmic constituents are present together.

These observations parallel findings in the ALS/FTD studies showing that PR and GR accumulate in cytoplasmic and cellular organelles (*11, 13*–*18, 24*). In addition, PR and GR colocalize with ribosomes in the cytoplasm of cells from patients’ brains(*25*). Paralleling the studies of PR and GR, if VR is bound to ribosomes this would yield a diffuse staining pattern and may point to its function in mitosis. To examine this, we tested the activity of VR in an *in vitro* translation system.

### VR dipeptides inhibit protein translation *in vitro*

Loveland et al (*19*) employed an *in vitro* translation system consisting of a rabbit reticulocyte extract supplemented with mRNA for firefly luciferase to ask if the addition of PR or GR would inhibit translation. They observed a strong signal of newly synthesized luciferase in the absence of PR or GR but found that translation was quenched upon adding PR or GR in increasing amounts. We carried out this same translation assay using a VR_10_ dipeptide (Fig. 2) and observed a strong luciferase signal with the luciferase mRNA alone. However, upon inclusion of 25, 50, and 100 μM VR_10_ dipeptide with the luciferase mRNA, expression of luciferase protein was reduced 6.8, and 19.7-fold for 50 and 100 μM respectively (Fig. 2). We observed little signal when the mRNA was absent, or Harringtonine, a known inhibitor of translation initiation, was present. These results together with the light microscopy point to the hypothesis that VR binds ribosomes and will depress translation.

**Fig. 2.**
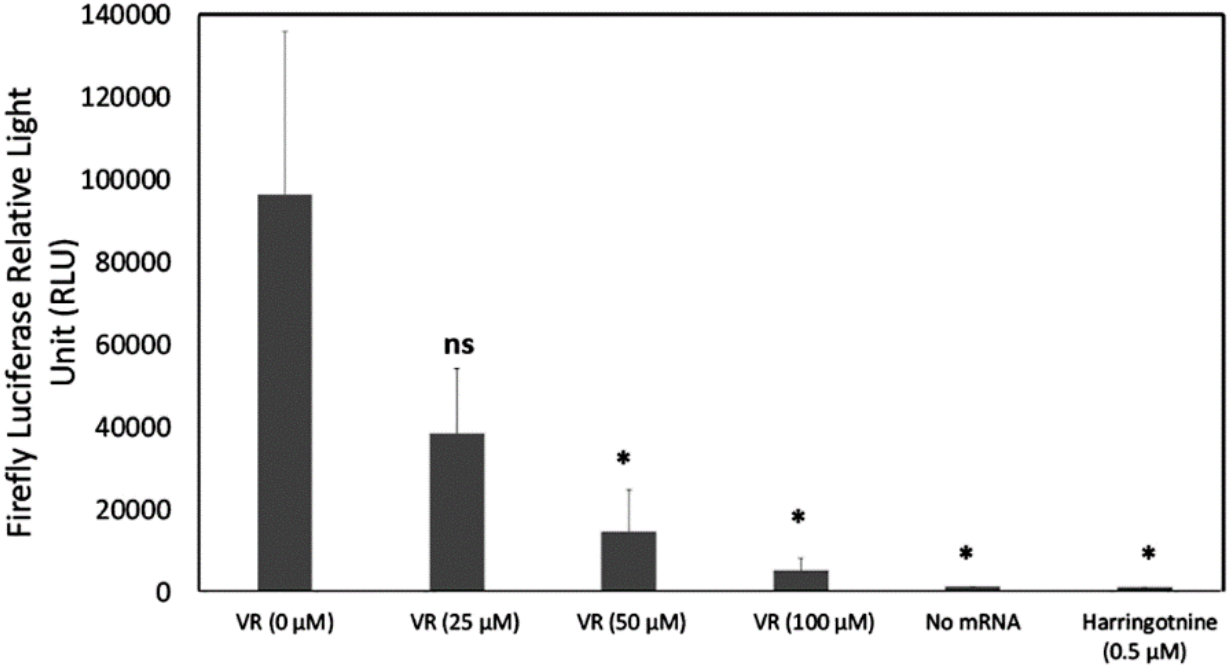
VR dipeptide repeat protein inhibits protein translation in a firefly luciferase assay. A commercial firefly luciferase assay kit was used to examine the ability of added VR_10_ to quench protein translation as measured by relative fluorescence units (RLU) of firefly luciferase (Methods). Amounts of 25, 50, and 100 μM VR_10_ were added to each reaction. Controls included no firefly luciferase mRNA, and Harringtonine, a known inhibitor of translation. Data presented are the ±SEM of three independent experiments. Unpaired two-tailed t test; P= 0.12 for 25 μM, *P = 0.05 for 50 μM VR versus untreated, and *P = 0.04 for 100 μM VR versus untreated. *P = 0.03 for minus mRNA versus untreated. *P = 0.03 for Harringtonine versus untreated.

### VR dipeptides colocalize with L4 ribosomal protein during mitosis

The cryo-EM data reported by Loveland et al (*19*) revealed that *in vitro*, PR and GR dipeptides bind in the exit tunnel in the large subunit of the ribosome. The exit tunnel of the ribosome is lined with ribosomal protein L4 for which an antibody is available. Thus, we asked if L4 and VR might co-localize in the cell. To probe this, we enriched cells in mitosis and co-stained with the rabbit antibody to VR and a mouse antibody to L4. As seen by laser scanning confocal microscopy, the VR peptides (green diffuse signals with some puncta) strongly colocalized with L4 protein (red diffuse signals) during mitotic progression (Fig. 3). These combined results suggest that like GR and PR, VR associates with ribosomes during mitosis.

**Fig. 3.**
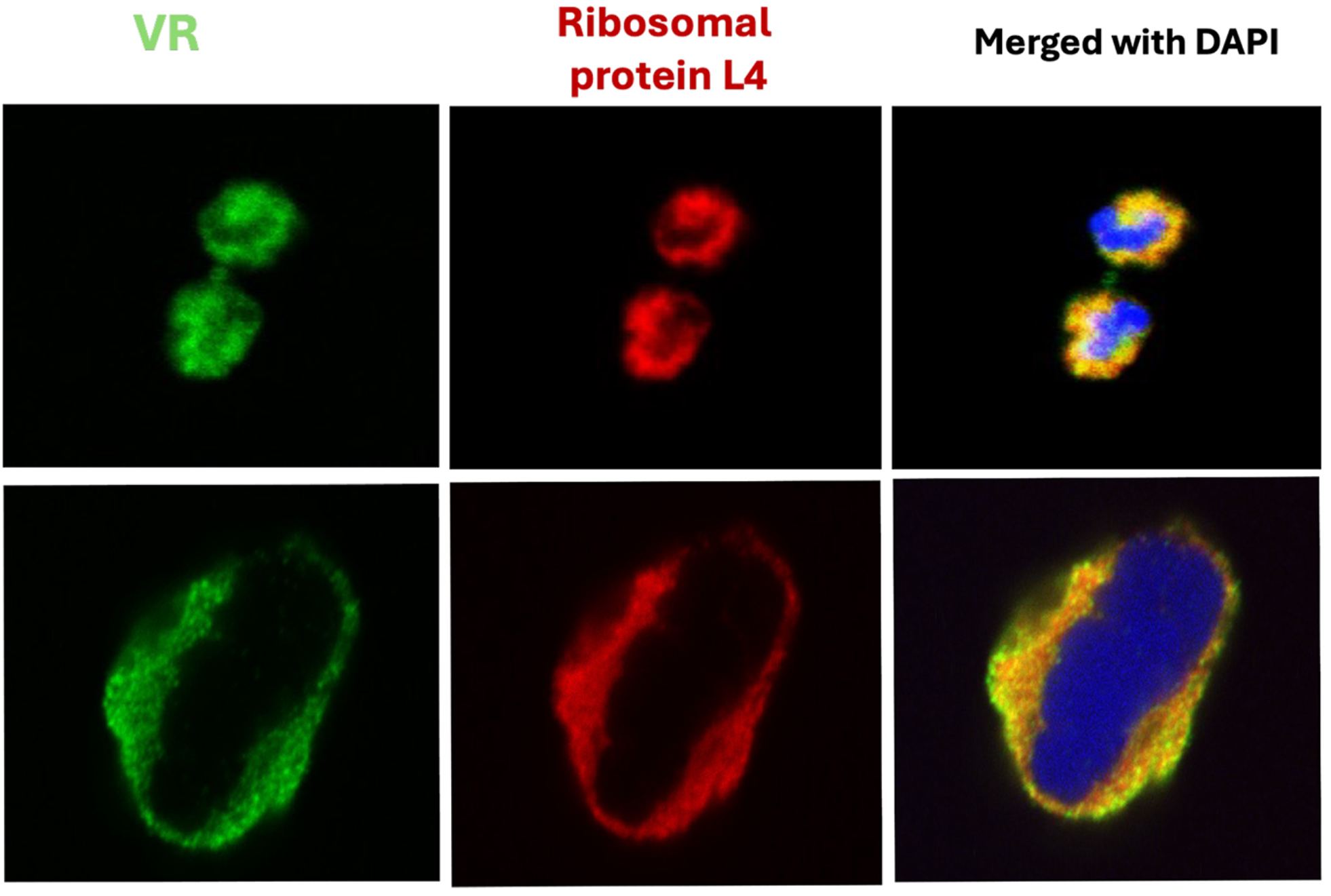
Co-localization of the VR dipeptide repeat protein and ribosomal protein L4. U2OS cells were grown in culture, plated on microscope slides, fixed and prepared for laser scanning microscopy (Methods). Microscopy images show two mitotic cells labeled with VR (green) and ribosomal protein L4 (red). The merged signals (yellow) indicate the colocalization of VR and L4 in the cytoplasm. VR and L4 were used at 1:50 dilution. Top row: cell in anaphase, bottom row: cell in metaphase.

### Immunoprecipitation of proteins from U2OS cells combined with mass spectrometry identifies ribosomes as a major class of VR interacting proteins

Affinity purification-mass spectrometry analysis of VR immunoprecipitation (IP) samples allowed unbiased identification of VR interacting proteins. Briefly, either VR antibody or IgG control antibody was bound to magnetic beads, incubated with U2OS cell extracts, and subjected to on-bead tryptic digestion and LC-MS/MS analysis. Over 1,600 proteins were identified as significantly enriched (p-value < 0.05, log2 fold change > 1) in the VR IP compared to the negative IgG IP control (Fig.4, supplemental Table 1). Of these enriched proteins, 209 are known ribosome-related proteins (Fig. 4 blue datapoints). Gene Ontology (GO) enrichment analysis revealed that among the VR associated proteins, ribosomal, proteasomal, translational and mitotic GO terms (Supplemental Table 2) were highly enriched in the VR IP (Fig. 4). These data further support an interaction between VR and ribosomes.

**Fig. 4.**
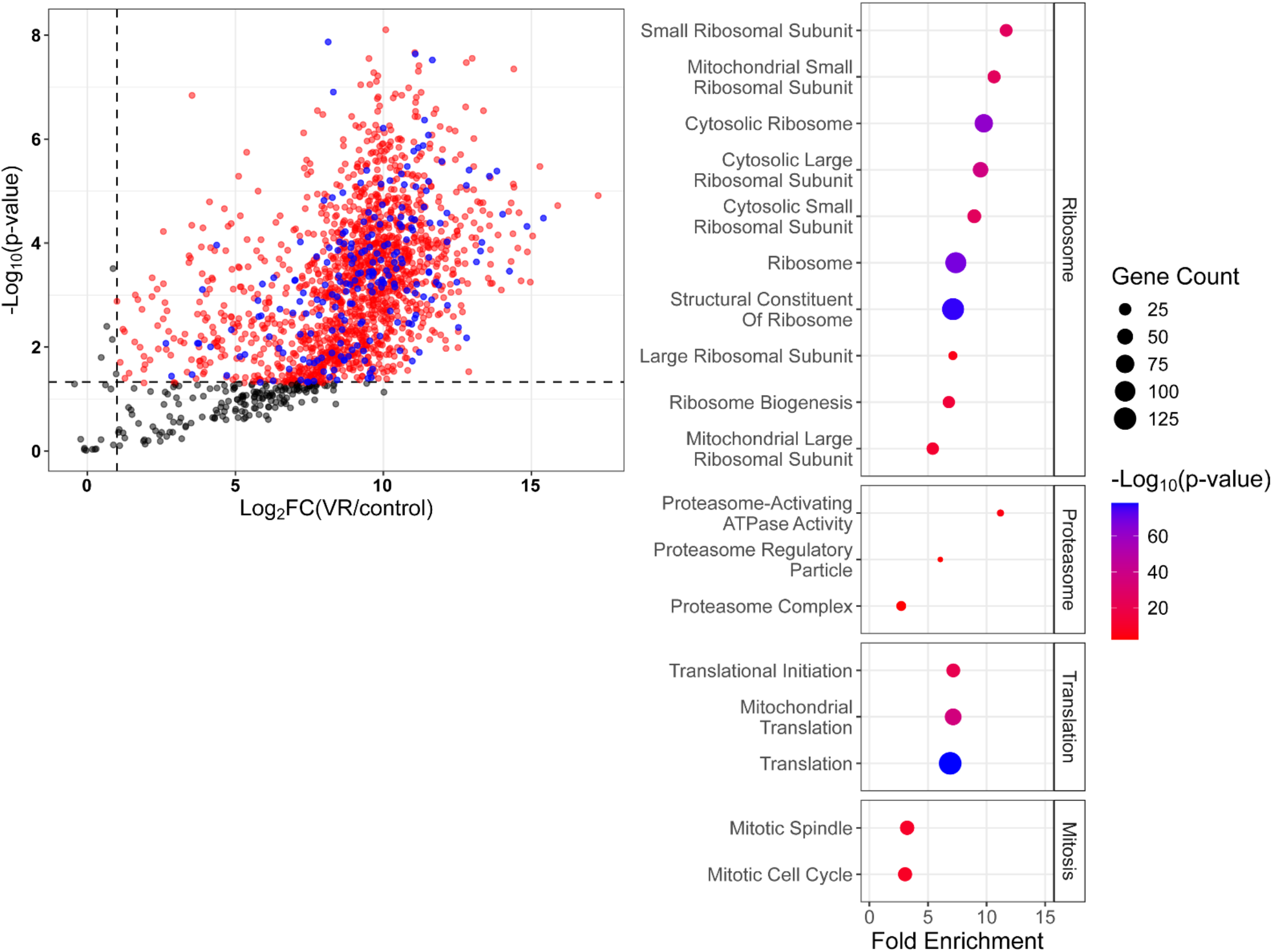
VR affinity purification-mass spectrometry analysis reveals significant enrichment of ribosomal proteins among VR interactors. (A) Volcano plot of VR AP-MS results. Significant interactors, defined as proteins with p-value < 0.05 and log2 fold change > 1, are highlighted in red and blue (1,665 total proteins). Blue datapoints are proteins that have ribosome-associated GO terms (209 of the 1,665 total). (B) GO enrichment analysis of the significant proteins identified in A revealed enrichment of ribosome, proteasome, translation and mitotic terms.

### VR dipeptide protein is highly expressed in a developing mouse embryonic cerebral cortex

Our observations using human cultured cells suggest that mammalian tissue with a high mitotic index might show elevated VR staining. To explore this, we turned to a developing mouse embryonic cerebral cortex. In the ventricular zone (VZ) of the embryonic cortex, progenitors divide symmetrically and asymmetrically to generate the appropriate number and types of precursors and neurons necessary to construct a functional cerebral cortex. Progenitor proliferation is at its peak in the embryonic day 16 mouse cortex, thus providing an ideal *in vivo* model to explore the expression of VR in rapidly dividing cells during organogenesis. We immunolabeled embryonic cortical slices with anti-VR and cortical progenitor specific antibodies (i.e., RC2). We found selective high expression of VR in the mitotically active progenitors in the VZ, consistent with the notion that tissue domains of high mitotic index display elevated VR expression (Fig. 5). VZ expression of VR is also evident at other stages of cortical development (e.g., E14 and P0). Intriguingly, VR expression was also noticed in the cortical plate (CP) where newly generated neurons settle, grow, and differentiate, after migration from the VZ, indicating potential role VR in post-mitotic neuronal differentiation independent of proliferation.

**Fig. 5.**
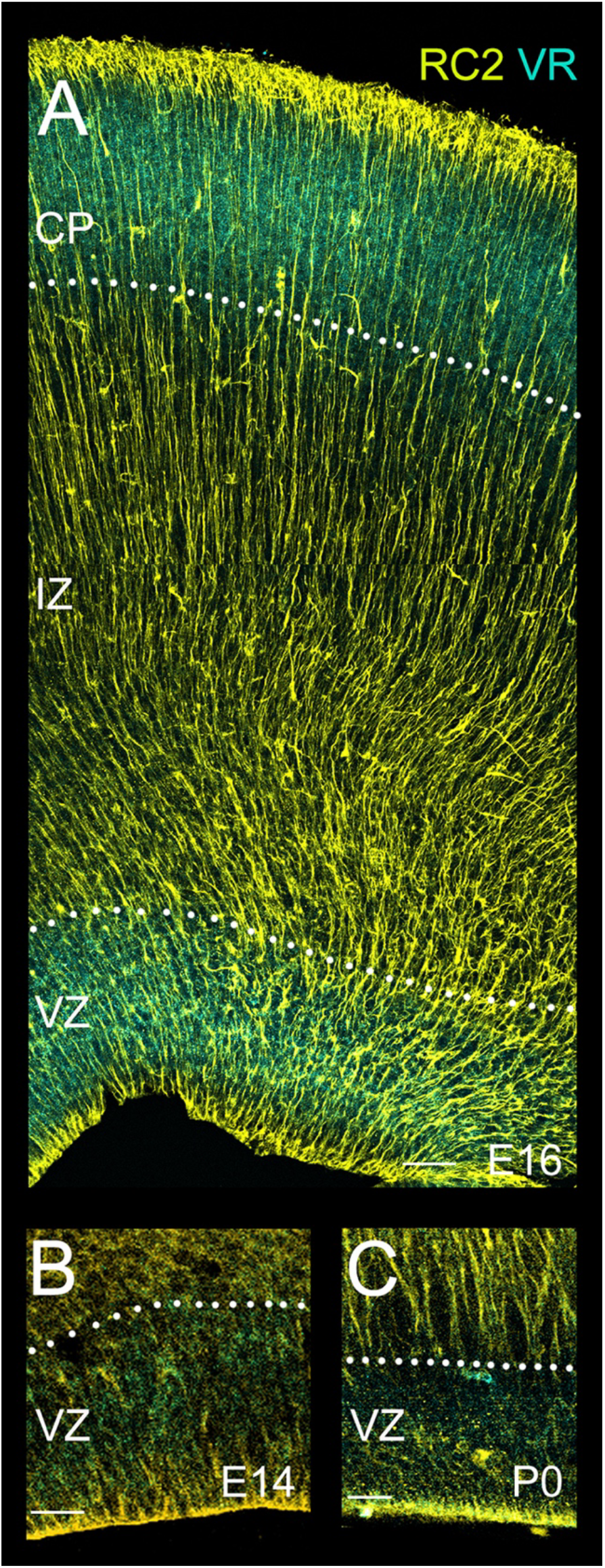
Expression of VR dipeptide protein in dividing progenitors of mouse embryonic cerebral cortex. (A) E16 mouse embryonic cortical section was labeled with anti-VR (teal) and cortical progenitor-specific RC2 (Yellow) antibodies. VR labels the ventricular zone (VZ) where progenitors proliferate as well as the cortical plate (CP) where new neurons settle. (B-C) VR expression is also evident in the early (E14) and late-stage (P0) cortical ventricular zone. Anti-VR and RC2 antibodies were used at 1:200 dilution. IZ, intermediate zone. Scale bar: 50µm (A), 25µm (B-C).

## DISCUSSION

We observed that when cultured human cells are stained with an antibody to the VR telomeric dipeptide repeat protein, cells in mitosis show very bright staining throughout the cell whereas non-mitotic cells show only low level, mostly nuclear staining. VR staining co-localized with ribosomes and *in vitro*, VR repressed translation in a firefly luciferase assay. Affinity purification-mass spectrometry analysis identified ribosomal proteins as a major class of VR interacting proteins. Extension of these findings to mouse embryonic cerebral cortical development showed strong staining in the ventricular zone where high mitotic index neural progenitor cells proliferate and in the cortical plate where new neurons settle. These observations point to VR playing a key role in mitosis, possibly suppressing global translation. If so, this suggests that the telomere has a more direct role in the cell cycle than had previously been considered.

The affinity purification-mass spectrometry studies comprised a pilot run (not shown) with one control and 2 VR affinity pull-downs, and the experiment shown which included 3 controls and 3 VR affinity pull-downs. Very similar results were obtained in both, with ample material recovered in the VR pull-downs arguing that significant amounts of VR were present in the U2OS cell extracts. This observation supports our previous work (8), and the results and conclusions presented here.

The most direct model explaining these results is that as cells enter mitosis, the nuclear envelope breaks down, releasing the fraction of free TERRA into the milieu of the cytoplasm replete with ribosomes. RAN translation then generates a burst of VR, and as the VR level builds up it binds the ribosomes, quenching global translation. As both translation and mitosis are highly energy-dependent, downregulating global translation may ensure enough energy for cell division. The bright mitotic staining by VR might also reflect a change in the oligomerization state of VR, which in the nucleus may be oligomerized into amyloid filaments or present in liquid droplets. Upon the breakdown of the nuclear membrane, VR may disperse into smaller oligomers/species more easily detected by the antibody. In either model the downstream events of repressing global translation would be the same. This argues for the presence of a significant amount of VR present in the mitotic cell. Due to the presumably random initiation of RAN translation on TERRA, if VR is produced, we must assume GL is also. However, since we have not been able to raise specific GL antibodies, we have no information about its cellular expression.

It has been known since the 1970’s that protein synthesis in mammalian cells is depressed during mitosis (*26*) with the overall degree having been estimated to be in the order of 70%. In earlier studies, cells were treated with nocodazole to inhibit microtubule polymerization and thus arresting cells before metaphase. Because nocodazole was later found to affect translation (*27*), Tanenbaum et al enriched cells in mitosis with the CDK1 inhibitor RO-3306 as used here and reported that the rate of protein synthesis in mitotic mammalian cells is repressed by about 35%, although the overall reduction may have been higher (*28*).

Mass spectrometry analysis of proteins immunoprecipitated with the VR antibody provided direct evidence that VR binds ribosomes. Ribosomal proteins were significantly enriched among the VR interacting proteins (p-value < 0.05 and log2 fold change > 1). This included ribosomal protein L4, a protein that lines the large subunit exit tunnel, and which we observed to co-localize with VR by light microscopy. Numerous other proteins related to ribosome assembly were present including nucleophosphim and nucleolin. Mitochondrial ribosomes and translational factors were highly enriched in the analysis paralleling the finding that GR will compromise mitochondrial function (*16,17*). In addition, this analysis provided insights into other cellular pathways in which VR may play a role including mRNA processing, and DNA repair (supplemental table 1). Cytoskeletal proteins and cell division proteins were also detected in abundance, suggesting other potential functions for VR outside of ribosomal regulation.

Cellular metabolic gene expression is tightly regulated in neural precursor cells to facilitate the appropriate generation of neurons and glia during cortical development (*29*). Translational repression via the expression of VR from TERRA may provide a hitherto unexplored axis of metabolic pathway regulation in these cells. It will be critical to establish if VR is differentially sequestered in symmetrically or asymmetrically dividing neural precursors and whether VR expression changes in precursors as cortical development unfolds and mitotic drive produces different types of neurons and glia. Further, the refinement of transcriptional programs by translation is a feature of corticogenesis (*30*) and VR might participate in this process by facilitating appropriate gene expression needed to promote progenitor proliferation and emerging neuronal identities. Intriguingly, VR is also expressed in the cortical plate where new neurons settle to make their circuit connections. VR binds to ribosomes, possibly quenching ribosome biogenesis and thus translation. Whether VR plays a role in decreased ribosomal numbers and translational downregulation seen in nonmitotic new neurons (*30*) during this stage of cortical development, via such a mechanism, remains an intriguing possibility.

## METHODS

### Cell growth and blocking cells in metaphase

Cycling U2OS cells were grown in Dulbecco’s modified Eagle medium (DMEM, Gibco), supplemented with 10% FBS and 3X Glutamax. For blocking cells in metaphase, cells were seeded at 50-60% confluency in a T75 flask and treated with 9 μM CDK-1 inhibitor (RO-3306) and incubated at 37° C with 5% CO_2_ for 16 h to block cells at late G2 phase. Cells were allowed to enter mitosis by washing three times with warm 1x PBS, adding drug-free media, and incubating them for 45 min. The isolation of mitotic cells was performed by tapping the flask (mitotic shake-off). Mitotic cells were collected in a 50 ml conical tube, centrifuged at 1000 rpm for 5 min, and replated in microscopy slides. Two hours later, the attached cells were fixed with 4% paraformaldehyde for 10 min at 4° C and permeabilized with 0.2% Triton X-100 in 1x PBS for 12 min.

### Immunostaining

Fixed cells were incubated with a blocking solution containing 10% Normal Goat Serum in 1X PBS for 1 h at room temperature. The cells on the slide were then incubated with a primary polyclonal VR antibody raised in rabbits (Biosynth inc.) (1:50 dilution) and a 1:100 dilution of a mouse monoclonal antibody to the ribosomal protein L4 (Sc-100838 Santa Cruz Biotechnology) in a blocking solution at 4° C overnight. The slides were washed 3X (10 min each) in 1X PBS and cells stained with AlexaFluor 488 and 647 conjugated secondary antibodies (Invitrogen A11034, and A21236 at a dilution of 1:750) for 40 min at room temperature. Following incubation with the secondary antibody, cells were washed 3X (10 min each) in 1X PBS and stained and mounted with prolong Gold Antifade mountant containing DAPI (Invitrogen P36931).

### Translation assay

The Promega Rabbit Reticulocyte Lysate system (L4960, Promega) was reconstituted in 50 μL aliquots consisting of 70% Rabbit Reticulocyte Lysate, 0.01 mM amino acid mixture minus methionine, 0.01 mM amino acid mixture minus leucine, 0.8 U Rnase inhibitor, and 50 μg of luciferase substrate as provided in the kit. VR dipeptides (25, 50 and 100 μM) were added to separate reactions. As a negative control, luciferase RNA was omitted or 0.5 μM Harringotnine was added to a separate reaction. Each reaction was incubated at 30° C for 20 min. Aliquots (2.5 μL) from each reaction were added to a 50 μL mixture containing the luciferase assay reagent (E1483 Promega) in a luminometer microplate. The illuminated light was read using Cytation 5 Imaging Reader.

### Immunohistochemistry

Cerebral cortices were removed from embryonic day 14,16, or P0 mice, fixed in 4% paraformaldehyde/PBS, embedded in 3% low melting point agarose in complete Hanks Balanced Salt Solution, and coronally sectioned (150µm) in a vibratome (Leica VT 1000S). Immunohistochemical labeling of embryonic brain sections or isolated neural precursor cells was performed as described earlier (*31*–*33*). The following primary antibodies were used: anti-VR (*8*) and RC2 (1:200 dilution; mouse IgM, Developmental Studies Hybridoma Bank, University of Iowa). Appropriate Cy3 and Alexa 488-conjugated secondary antibodies (Jackson ImmunoResearch, Molecular Probes) were used to detect primary antibody binding. DAPI (Sigma-Aldrich, D9542) was used as a nuclear counterstain.

### Light microscopy

Visualization of cells in tissue culture following our previously described imaging strategy (*8*). An Olympus FV-1000 confocal microscope equipped with a 60X oil immersion objective was used to acquire the Z-stacks images. The acquisition employed both sequential and simultaneous multi-channel imaging with optimized laser and detector settings. Alexa 488 excitation utilized an argon-ion laser, while Alexa 647 signals were detected using a far-red detector. Acquisition parameters were optimized for each fluorescence channel using cells with the highest Signal-to-Noise Ratio (SNR). PMT voltage, amplifier, and offset levels were adjusted to further enhance SNR. Laser power was maintained below 7% and PMT HV set below 700 to prevent photobleaching and signal saturation. Z-slices were acquired with a step size of 0.40 μm. These optimized settings were applied consistently across independent experiments. Stacked images were processed into maximum and average intensity projections using Fiji (*34*)

Mouse cerebral cortical tissue was visualized using a similar imaging strategy (*33, 35, 36*). Images were obtained using a Zeiss LSM780 confocal microscope.

### Immunoprecipitation

U2OS cells were grown to 90% confluence on 150 mm plates, washed twice with ice-cold PBS, harvested by scraping, and pelleted. PBS was removed and cells were resuspended in 1 mL lysis buffer (0.1% NP-40, 50 mM Hepes-NaOH pH 8, 150 mM NaCl) containing protease and phosphatase inhibitor cocktail (PPC1010, Sigma). Whole cell lysates were incubated on ice for 30 min with one snap freeze in liquid nitrogen at 15 min into the incubation and centrifuged for 15 min at maximum speed in a refrigerated microfuge to pellet DNA. Supernatant was recovered and total protein concentration determined using a Nanodrop at 280 nm. Equal amounts of lysate were aliquoted and pre-cleared with 20 μl of Dynabeads protein A (10001D, Invitrogen) per ml of lysate for 1 hour at 4°C, and then again with another 20 μl of protein A beads per ml, overnight, on a rocker at 4°C. After pre-clearing, beads were removed (using magnet) and lysate was transferred to a fresh tube. A magnetic stand was used to isolate beads, and the cleared lysate was transferred to fresh tubes. Concomitantly, 10 μg of rabbit anti-VR antibody was incubated with 25 μl of protein A beads for 1 hr on a rocker at 4°C. For control, we used 10ug of rabbit IgG Isotype control antibody (08-6199, Invitrogen). The beads were washed 2x with lysis buffer to eliminate unbound antibody and then added to the pre-cleared lysate and the mix was incubated overnight at 4°C with gentle agitation. The beads were washed 3x with 1 ml ice cold lysis buffer (without protease/phosphatase inhibitors) and then washed 3x with 1 ml ice cold wash buffer (50 mM Ammonium Bicarbonate, pH 7.8). Triplicate samples for both IgG control and VR conditions were submitted to the UNC Metabolomics and Proteomics Core Facility for mass spectrometry analysis in 50 μl wash buffer.

### Mass Spectrometry

#### Sample preparation

Immunoprecipitated protein samples were subjected to on-bead trypsin digestion as previously described (*37*). Briefly, on-bead digestion was performed by adding 1µg trypsin and incubating, with shaking, overnight at 37ºC. The next day, 1µg trypsin was added and samples were incubated at 37ºC for an additional 3h. Beads were pelleted, and supernatants transferred to fresh tubes. The beads were washed twice with 100µl LC-MS grade water, and washes were added to the original supernatants. Samples were acidified by adding trifluoracetic acid for a final concentration of 2%. Peptides were dried and desalted using peptide desalting spin columns (Pierce), lyophilized and stored at -80ºC until further analysis.

#### LC-MS/MS

The peptide samples were analyzed by liquid chromatography-tandem mass spectrometry (LC-MS/MS) using an Ultimate3000 coupled to an Exploris480 mass spectrometer (Thermo Scientific). Samples were injected onto an IonOpticks Aurora series 2 C18 column (75 μm id × 15 cm, 1.6 μm particle size; IonOpticks) and separated over a 90-minute method. The gradient for separation consisted of 2–40% mobile phase B at a 250 nl/min flow rate, where mobile phase A was 0.1% formic acid in water and mobile phase B consisted of 0.1% formic acid in ACN. The Exploris480 was operated in data-dependent mode with a cycle time of 2s. Resolution for the precursor scan (m/z 375–1500) was set to 120,000, while MS/MS scans resolution was set to 15,000. The normalized collision energy was set to 30% for HCD. Peptide match was set to preferred, and precursors with unknown charge or a charge state of 1 and ≥ 7 were excluded.

### Data Analysis

Raw data files were processed using MaxQuant version 1.6.15.0 and searched against the Uniprot reviewed human database (containing 20,396 entries, downloaded March 2021) appended with a common contaminants database (245 sequences), using Andromeda within MaxQuant (*38, 39*) Enzyme specificity was set to trypsin, up to two missed cleavage sites were allowed, methionine oxidation and N-terminus acetylation were set as variable modifications. A 1% FDR was used to filter all data. Match between runs was enabled (5 min match time window, 20 min alignment window), and a minimum of two unique peptides was required for label-free quantitation using the LFQ intensities. The mass spectrometry proteomics data have been deposited to the ProteomeXchange Consortium (*40*) via the PRIDE *(43)* partner repository with the dataset identifier PXD054040.

Perseus was used for further processing (*41*). Reverse hits, and proteins with only 1 unique+razor peptide, were removed from the dataset. Proteins with >50% missing values were removed, and missing values were imputed from normal distribution within Perseus. Log2 fold change (FC) ratios were calculated using the averaged Log2 LFQ intensities of and students t-test performed for each pairwise comparison, with p-values calculated. Proteins with significant p-values (<0.05, Student’s T-test) and Log2 FC >1 were considered biological interactors.

Gene Ontology and enrichment analyses were conducted using DAVID (*42, 43*). Genes were searched against the default databases. GO terms used are described in supplemental Table 1.

Fig. 4 was made using tidyverse, ggplot2 (*44*), and janitor (*44*) packages in R version 4.3.1 (45).

## Supporting information

Al Turki et al

## Author contributions

TMA and VM carried out the light microscopy studies of cultured cells, S-JC, HL, CM, and EA carried out the studies of the mouse embryonic tissue. TMA and VM conducted the *in vitro* translation studies. SW carried out the preparation for the affinity purification and mass spectrometry. JDG, TMA, SW, VM, EA, CAM and LEH contributed to the design of the work and writing the manuscript and data analysis.

## Data availability

All data needed to evaluate the conclusions in the paper are present in the paper.

