## Supplementary material for "The valine-arginine dipeptide repeat protein encoded by mammalian telomeric RNA appears highly expressed in mitosis and may repress global translation": Al Turki et al

Supplemental Table 1
Mass spectrometry Results

| Majority protein IDs | Protein names | Gene names | Fasta headers | Log2 Fold Change VR v IgG | Student's T-test p-value VR v IgG | Log2 LFQ intensity VR R1 | Log2 LFQ intensity VR R2 | Log2 LFQ intensity VR R3 |
|---|---|---|---|---|---|---|---|---|
| P35579 | Myosin-9 | MYH9 | Myosin-9 O | 4.608827 | 0.003083 | 29.32622 | 29.44729 | 29.96747 |
| P63261;P6 | Actin, cytop | ACTG1;ACT | Actin, cytop | 4.519056 | 0.002213 | 31.08036 | 31.10106 | 31.45484 |
| P08670 | Vimentin | VIM | Vimentin OS | 10.2434 | 0.000528 | 29.37478 | 30.89077 | 29.06283 |
| P62906 | 60S ribosor | RPL10A | Large ribosc | 9.95155 | 0.000506 | 32.4845 | 33.09475 | 33.2292 |
| P62805 | Histone H4 | HIST1H4A | Histone H4 | 6.822599 | 0.000323 | 30.65881 | 30.59023 | 30.65233 |
| Q6NZI2 | Polymerase | PTRF | Caveolae-as | 10.3406 | 0.000617 | 30.20222 | 30.09207 | 30.2889 |
| Q70J99 | Protein unc | UNC13D | Protein unc | 17.25345 | 1.22E-05 | 29.47046 | 28.91684 | 29.58484 |
| P16403 | Histone H1. | HIST1H1C | Histone H1. | 6.399636 | 0.000413 | 30.49781 | 30.19625 | 30.5401 |
| Q6P2Q9 | Pre-mRNA-p | PRPF8 | Pre-mRNA-p | 13.58161 | 4.63E-05 | 27.50393 | 27.35035 | 27.76812 |
| Q9NQW7 | Xaa-Pro am | XPNPEP1 | Xaa-Pro am | 14.96216 | 0.000564 | 29.48719 | 29.35466 | 29.50042 |
| P07437 | Tubulin bet | TUBB | Tubulin bet | 7.528848 | 0.000262 | 29.37148 | 29.27824 | 29.40046 |
| P22626 | Heterogene | HNRNPA2B | Heterogene | 13.54131 | 0.000473 | 29.56129 | 29.38354 | 29.55541 |
| O75643 | U5 small nu | SNRNP200 | U5 small nu | 15.28595 | 3.35E-06 | 28.58692 | 28.53267 | 28.7907 |
| P06748 | Nucleopho | NPM1 | Nucleopho | 9.626893 | 0.000877 | 29.83005 | 29.81507 | 29.94377 |
| Q96RE7 | Nucleus acc | NACC1 | Nucleus acc | 14.64955 | 0.000532 | 29.06361 | 30.32195 | 29.01518 |
| P68363;Q9 | Tubulin alp | TUBA1B;TU | Tubulin alp | 7.264825 | 0.000475 | 29.43098 | 29.25784 | 29.49373 |
| Q9UBU9 | Nuclear RNA | NXF1 | Nuclear RNA | 13.61909 | 0.000256 | 28.6188 | 28.1795 | 28.54768 |
| P62701;Q8 | 40S ribosor | RPS4X;RPS | Small ribos | 7.177037 | 0.001328 | 29.59001 | 29.29307 | 29.54624 |
| P36578 | 60S ribosor | RPL4 | Large ribosc | 9.64334 | 0.000235 | 28.89009 | 28.8064 | 28.95316 |
| Q5JTV8 | Torsin-1A-ir | TOR1AIP1 | Torsin-1A-ir | 15.04104 | 7.30E-05 | 28.71607 | 28.65308 | 28.78419 |
| Q16778;P3 | Histone H2I | HIST2H2BE | Histone H2I | 8.27676 | 0.000879 | 30.43002 | 30.07979 | 30.65574 |
| P19338 | Nucleolin | NCL | Nucleolin C | 8.758216 | 0.000146 | 28.43249 | 28.57384 | 28.52977 |
| Q92614 | Unconventi | MYO18A | Unconventi | 13.35988 | 2.37E-05 | 28.00426 | 27.0427 | 27.42614 |
| P62249 | 40S ribosor | RPS16 | Small ribos | 8.943441 | 0.001468 | 29.57376 | 29.44983 | 29.90038 |
| P26373 | 60S ribosor | RPL13 | Large ribosc | 8.376099 | 0.001701 | 29.17067 | 28.82596 | 29.10522 |
| Q02878 | 60S ribosor | RPL6 | Large ribosc | 10.01755 | 0.000592 | 29.04466 | 29.0234 | 29.17515 |
| P46781 | 40S ribosor | RPS9 | Small ribos | 8.87212 | 0.000629 | 29.08446 | 28.85979 | 29.09419 |
| Q15029 | 116 kDa U5 | EFTUD2 | 116 kDa U5 | 12.52971 | 0.005032 | 27.59135 | 27.87079 | 28.24921 |
| P61247 | 40S ribosor | RPS3A | Small ribos | 8.690842 | 0.003059 | 28.75525 | 28.90314 | 28.89067 |
| P18124 | 60S ribosor | RPL7 | Large ribosc | 9.266002 | 0.000152 | 29.04396 | 28.95144 | 29.13912 |
| P62424 | 60S ribosor | RPL7A | Large ribosc | 9.567764 | 0.000666 | 28.99337 | 28.50624 | 28.75178 |
| P61353 | 60S ribosor | RPL27 | Large ribosc | 8.634131 | 6.53E-05 | 29.23932 | 29.06116 | 29.53057 |
| P62241 | 40S ribosor | RPS8 | Small ribos | 8.169163 | 0.000123 | 29.10282 | 28.97443 | 29.08509 |
| Q99878;Q9 | Histone H2A | HIST1H2AJ; | Histone H2A | 7.742718 | 0.001856 | 30.58388 | 30.26629 | 30.42402 |
| Q00839 | Heterogene | HNRNPU | Heterogene | 9.316052 | 0.002472 | 28.92841 | 28.82935 | 28.97356 |
| P07355;A6 | Annexin A2; | ANXA2;ANX | Annexin A2 | 4.856369 | 0.00259 | 28.55644 | 28.29885 | 28.54189 |
| P60660 | Myosin ligh | MYL6 | Myosin ligh | 4.869793 | 0.002674 | 29.16531 | 29.18641 | 29.66385 |
| P39023 | 60S ribosor | RPL3 | Large ribosc | 10.79563 | 0.000817 | 28.33756 | 28.35203 | 28.33782 |
| Q07020 | 60S ribosor | RPL18 | Large ribosc | 10.38633 | 0.000701 | 29.80158 | 29.39344 | 29.57051 |
| P15880 | 40S ribosor | RPS2 | Small ribos | 9.628021 | 0.000424 | 29.31164 | 29.09705 | 29.21294 |
| P52272 | Heterogene | HNRNPM | Heterogene | 10.77956 | 0.000349 | 27.67976 | 27.3901 | 27.65267 |
| Q5VTE0;P6 | Putative elo | EEF1A1P5; | Putative elo | 4.974856 | 0.004206 | 29.97049 | 29.80327 | 29.91157 |
| P05388;Q8 | 60S acidic r | RPLP0;RPLI | Large ribosc | 9.965565 | 1.83E-05 | 28.78457 | 28.42858 | 28.4554 |
| O94906 | Pre-mRNA-p | PRPF6 | Pre-mRNA-p | 14.58086 | 2.60E-05 | 27.13291 | 26.95892 | 27.07408 |

| | | | | | | | | |
|---|---|---|---|---|---|---|---|---|
| P61978 | Heterogene | HNRNPK | Heterogene | 8.868413 | 0.00211 | 28.30125 | 28.11222 | 28.26137 |
| P62277 | 40S ribosor | RPS13 | Small ribos | 8.096869 | 0.000235 | 28.79238 | 28.54915 | 28.67522 |
| P62269 | 40S ribosor | RPS18 | Small ribos | 3.763571 | 0.008493 | 28.74256 | 28.55877 | 28.83708 |
| P62244 | 40S ribosor | RPS15A | Small ribos | 8.542702 | 0.000553 | 29.49016 | 29.37824 | 29.62871 |
| P61313 | 60S ribosor | RPL15 | Large ribosc | 10.10858 | 0.000337 | 28.85991 | 28.86997 | 28.91268 |
| Q15149 | Plectin | PLEC | Plectin OS= | 3.010346 | 0.094254 | 29.38856 | 31.16471 | 31.22596 |
| P11940;Q9 | Polyadenyl | PABPC1;PA | Polyadenyl | 11.0268 | 0.000309 | 28.06564 | 27.74817 | 28.07368 |
| P25705 | ATP syntha: | ATP5A1 | ATP syntha: | 8.714914 | 0.004682 | 27.70293 | 27.57788 | 27.6569 |
| P05141 | ADP/ATP tra | SLC25A5 | ADP/ATP tra | 8.065065 | 0.001147 | 28.47197 | 28.37069 | 28.54617 |
| Q9BUQ8 | Probable AT | DDX23 | Probable AT | 14.41089 | 8.43E-06 | 27.32629 | 27.22343 | 27.37758 |
| Q71DI3;Q1 | Histone H3. | HIST2H3A;I | Histone H3. | 9.191062 | 0.005906 | 29.59137 | 29.45683 | 30.26282 |
| P09651;A0 | Heterogene | HNRNPA1;I | Heterogene | 8.20788 | 0.003226 | 28.40892 | 28.30213 | 28.49739 |
| P07305 | Histone H1. | H1F0 | Histone H1. | 5.99012 | 0.000221 | 29.11354 | 29.04682 | 29.17138 |
| Q08211 | ATP-depenc | DHX9 | ATP-depenc | 6.731261 | 0.000719 | 28.62422 | 27.01255 | 27.14773 |
| Q8TDD1 | ATP-depenc | DDX54 | ATP-depenc | 6.250025 | 0.082103 | 30.58441 | 30.59094 | 30.79157 |
| P62753 | 40S ribosor | RPS6 | Small ribos | 11.62894 | 0.001229 | 28.72433 | 28.65884 | 28.34079 |
| P23396 | 40S ribosor | RPS3 | Small ribos | 9.540844 | 0.007583 | 28.24364 | 28.11515 | 28.26078 |
| P62917 | 60S ribosor | RPL8 | Large ribosc | 8.460595 | 0.000138 | 29.04257 | 29.03808 | 29.03149 |
| P19105;O1 | Myosin regu | MYL12A;MY | Myosin regu | 5.332739 | 0.005582 | 28.47232 | 28.45325 | 29.1118 |
| P17844 | Probable AT | DDX5 | Probable AT | 7.564681 | 0.001442 | 27.60187 | 27.40186 | 27.60738 |
| Q8TDM6 | Disks large I | DLG5 | Disks large I | 12.99935 | 2.79E-08 | 26.86561 | 26.73815 | 26.82408 |
| Q02809 | Procollager | PLOD1 | Procollager | 14.86226 | 0.000148 | 27.90236 | 27.71299 | 27.82972 |
| P27635 | 60S ribosor | RPL10 | Large ribosc | 9.626429 | 0.000349 | 29.59155 | 29.1541 | 29.93733 |
| P05387 | 60S acidic r | RPLP2 | Large ribosc | 8.934376 | 0.000246 | 28.41925 | 28.42174 | 28.59399 |
| P18621 | 60S ribosor | RPL17 | Large ribosc | 7.992825 | 1.51E-05 | 28.83569 | 28.53103 | 28.75863 |
| O43290 | U4/U6.U5 t | SART1 | U4/U6.U5 t | 8.681949 | 0.00018 | 26.49433 | 26.46466 | 26.80649 |
| P62750 | 60S ribosor | RPL23A | Large ribosc | 7.294573 | 0.000907 | 28.92369 | 28.64341 | 28.80002 |
| P84098 | 60S ribosor | RPL19 | Large ribosc | 7.194441 | 9.50E-05 | 28.82223 | 28.85261 | 28.91886 |
| P62851 | 40S ribosor | RPS25 | Small ribos | 5.766127 | 0.001276 | 28.9687 | 28.97016 | 29.02923 |
| P46779 | 60S ribosor | RPL28 | Large ribosc | 10.17844 | 0.000421 | 28.62408 | 28.65816 | 28.65779 |
| P51991 | Heterogene | HNRNPA3 | Heterogene | 12.36932 | 0.000389 | 27.35313 | 27.39705 | 27.56384 |
| P62081 | 40S ribosor | RPS7 | Small ribos | 7.293025 | 0.017322 | 27.75098 | 27.68392 | 27.8408 |
| P35251 | Replication | RFC1 | Replication | 5.16538 | 0.11475 | 28.31514 | 30.4343 | 30.54324 |
| P11142 | Heat shock | HSPA8 | Heat shock | 5.104549 | 0.000286 | 27.29035 | 27.13555 | 27.2226 |
| P14866 | Heterogene | HNRNPL | Heterogene | 13.61729 | 0.000134 | 28.26638 | 28.22522 | 26.94592 |
| P46778 | 60S ribosor | RPL21 | Large ribosc | 9.229631 | 0.000374 | 29.09343 | 28.94517 | 29.11823 |
| P11586 | C-1-tetrahy | MTHFD1 | C-1-tetrahy | 11.28565 | 7.89E-05 | 26.51474 | 26.44905 | 26.77354 |
| P49454 | Centromere | CENPF | Centromere | 11.57343 | 1.24E-06 | 25.14717 | 25.30009 | 25.09027 |
| P62829 | 60S ribosor | RPL23 | Large ribosc | 10.78001 | 8.58E-05 | 29.15381 | 28.74477 | 28.85712 |
| P25054 | Adenomatc | APC | Adenomatc | 10.15225 | 0.000375 | 26.08912 | 26.11428 | 26.83882 |
| P42766 | 60S ribosor | RPL35 | Large ribosc | 6.297078 | 0.000464 | 28.90805 | 28.99113 | 28.89908 |
| P83731 | 60S ribosor | RPL24 | Large ribosc | 9.13168 | 0.000756 | 29.05512 | 28.86809 | 28.72533 |
| P07858 | Cathepsin E | CTSB | Cathepsin E | 14.40347 | 4.47E-08 | 28.85085 | 28.78759 | 28.74612 |
| P08708 | 40S ribosor | RPS17 | Small ribos | 5.789654 | 0.002596 | 28.46494 | 28.3789 | 28.5575 |
| Q9NR30 | Nucleolar R | DDX21 | Nucleolar R | 12.98849 | 7.82E-05 | 26.84482 | 26.69445 | 26.78831 |
| Q8TF01 | Arginine/ser | PNISR | Arginine/ser | 11.35039 | 1.91E-05 | 27.72683 | 27.65867 | 28.07214 |
| Q15418 | Ribosomal | RPS6KA1 | Ribosomal | 14.84996 | 4.77E-05 | 26.99701 | 26.44608 | 27.02454 |
| P61254 | 60S ribosor | RPL26 | Large ribosc | 11.78479 | 0.000562 | 27.88239 | 27.75557 | 28.14705 |

| | | | | | | | | |
|---|---|---|---|---|---|---|---|---|
| P40429 | 60S riboso | RPL13A | Large ribos | 8.897094 | 0.000173 | 28.40483 | 28.48324 | 28.60184 |
| Q53GS9 | U4/U6.U5 t | USP39 | Ubiquitin ca | 14.31337 | 0.000226 | 27.22903 | 27.13095 | 27.27844 |
| P14618 | Pyruvate kir | PKM | Pyruvate kir | 5.705334 | 0.006802 | 27.06034 | 27.08477 | 27.07521 |
| P18077 | 60S riboso | RPL35A | Large ribos | 14.02831 | 2.79E-05 | 27.8929 | 27.66194 | 27.81672 |
| Q8IVL6 | Prolyl 3-hy | LEPREL2 | Prolyl 3-hy | 13.3793 | 2.84E-07 | 28.44608 | 28.37306 | 28.40373 |
| P62280 | 40S riboso | RPS11 | Small ribos | 8.436052 | 0.001076 | 28.44514 | 28.21803 | 28.26973 |
| P67809 | Nuclease-s | YBX1 | Y-box-bindi | 13.36423 | 1.67E-05 | 27.91102 | 27.76477 | 28.03039 |
| Q07955 | Serine/argin | SRSF1 | Serine/argin | 10.52448 | 0.001013 | 28.05734 | 27.86106 | 27.94145 |
| P78527 | DNA-depen | PRKDC | DNA-depen | 12.14547 | 0.001934 | 24.90141 | 25.54822 | 25.78843 |
| P62263 | 40S riboso | RPS14 | Small ribos | 7.951312 | 0.001365 | 27.79532 | 27.56296 | 27.81818 |
| P62899 | 60S riboso | RPL31 | Large ribos | 8.031346 | 0.000638 | 28.7293 | 28.65465 | 28.84119 |
| P49207 | 60S riboso | RPL34 | Large ribos | 9.668446 | 0.000376 | 28.56362 | 28.3593 | 28.82796 |
| Q9Y2W1 | Thyroid hor | THRAP3 | Thyroid hor | 9.192651 | 0.000175 | 26.79724 | 26.62282 | 26.90724 |
| P50914 | 60S riboso | RPL14 | Large ribos | 8.522248 | 9.48E-05 | 29.3952 | 29.09617 | 29.28094 |
| P30050 | 60S riboso | RPL12 | Large ribos | 8.923412 | 0.000102 | 28.88945 | 28.68392 | 28.76066 |
| Q12906 | Interleukin | ILF3 | Interleukin | 13.27127 | 1.69E-05 | 26.83111 | 26.14307 | 26.90081 |
| Q9UQ35 | Serine/argin | SRRM2 | Serine/argin | 8.184154 | 0.000391 | 26.35187 | 26.08458 | 26.24098 |
| Q9UMS4 | Pre-mRNA- | PRPF19 | Pre-mRNA- | 12.36264 | 0.002583 | 27.45065 | 27.32697 | 27.51721 |
| P35580 | Myosin-10 | MYH10 | Myosin-10 | 5.074209 | 0.003518 | 25.44199 | 25.64067 | 26.25364 |
| Q96DI7 | U5 small nu | SNRNP40 | U5 small nu | 13.98301 | 0.000798 | 27.77725 | 27.61154 | 27.87531 |
| P62266 | 40S riboso | RPS23 | Small ribos | 7.74306 | 0.000181 | 28.55578 | 28.65096 | 28.83032 |
| P23246 | Splicing fac | SFPQ | Splicing fac | 3.995047 | 0.022194 | 26.97705 | 26.40813 | 26.98772 |
| P39019 | 40S riboso | RPS19 | Small ribos | 2.973832 | 0.053143 | 27.63547 | 27.03328 | 27.51616 |
| P51114 | Fragile X me | FXR1 | RNA-bindin | 7.175042 | 7.28E-05 | 27.04677 | 26.97465 | 27.06948 |
| Q13247 | Serine/argin | SRSF6 | Serine/argin | 13.61114 | 5.53E-06 | 28.41058 | 28.43289 | 28.64509 |
| P11021 | 78 kDa gluc | HSPA5 | Endoplasm | 4.368656 | 0.000109 | 26.53991 | 26.28194 | 26.48278 |
| Q02543 | 60S riboso | RPL18A | Large ribos | 10.15144 | 0.000104 | 28.41969 | 28.38895 | 28.71313 |
| P50454 | Serpin H1 | SERPINH1 | Serpin H1 C | 6.354118 | 0.008697 | 27.1054 | 26.96873 | 27.2141 |
| P62847 | 40S riboso | RPS24 | Small ribos | 8.877694 | 0.001451 | 28.58628 | 28.34266 | 28.51247 |
| P52597 | Heterogene | HNRNPF | Heterogene | 13.61883 | 0.000601 | 27.69465 | 27.59996 | 27.11852 |
| O43390 | Heterogene | HNRNPR | Heterogene | 12.75799 | 3.09E-05 | 27.13212 | 26.52709 | 27.11575 |
| O76021 | Ribosomal | RSL1D1 | Ribosomal | 11.60929 | 8.01E-06 | 26.34221 | 26.05631 | 26.1778 |
| Q8N1F8 | Serine/thre | STK11IP | Serine/thre | 11.9107 | 0.002275 | 26.78307 | 24.49619 | 27.58241 |
| Q9Y3U8 | 60S riboso | RPL36 | Large ribos | 10.59215 | 3.15E-05 | 27.84212 | 28.17269 | 28.23786 |
| Q92522 | Histone H1 | H1FX | Histone H1 | 8.686282 | 0.000825 | 27.61246 | 27.4243 | 27.48208 |
| P08238 | Heat shock | HSP90AB1 | Heat shock | 5.635888 | 0.001153 | 26.66871 | 26.28169 | 26.29642 |
| Q14980 | Nuclear mit | NUMA1 | Nuclear mit | 10.33301 | 3.22E-05 | 24.94597 | 24.96746 | 24.92021 |
| Q02539 | Histone H1 | HIST1H1A | Histone H1 | 7.872903 | 0.013104 | 28.17555 | 27.97049 | 28.25016 |
| P46776 | 60S riboso | RPL27A | Large ribos | 8.667514 | 0.000103 | 28.72981 | 28.57438 | 28.61547 |
| P53621 | Coatomer s | COPA | Coatomer s | 9.049112 | 8.23E-06 | 25.71366 | 25.504 | 25.7297 |
| O60506 | Heterogene | SYNCRIP | Heterogene | 10.71928 | 0.000175 | 26.57888 | 26.31254 | 26.58849 |
| P12956 | X-ray repair | XRCC6 | X-ray repair | 8.461068 | 0.000779 | 26.3674 | 26.29511 | 26.31244 |
| Q7Z406 | Myosin-14 | MYH14 | Myosin-14 | 4.402656 | 0.009728 | 28.07796 | 28.05874 | 28.65676 |
| P68371 | Tubulin bet | TUBB4B | Tubulin bet | 12.88375 | 0.029814 | 27.71764 | 27.69226 | 27.72423 |
| Q99459 | Cell divisio | CDC5L | Cell divisio | 10.51257 | 7.67E-05 | 25.71398 | 25.62701 | 25.74052 |
| Q12797 | Aspartyl/as | ASPH | Aspartyl/as | 12.22579 | 6.43E-07 | 26.3227 | 26.19551 | 26.2981 |
| Q99986 | Serine/thre | VRK1 | Serine/thre | 15.8884 | 1.90E-05 | 27.04239 | 27.17864 | 27.25368 |
| Q14674 | Separin | ESPL1 | Separin OS= | 12.65481 | 0.000526 | 26.13283 | 26.00443 | 26.11795 |

| | | | | | | | | |
|---|---|---|---|---|---|---|---|---|
| Q8NC51 | Plasminoge | SERBP1 | SERPINE1 r | 8.143191 | 0.010022 | 26.82226 | 26.9788 | 26.77718 |
| P26599 | Polypyrimic | PTBP1 | Polypyrimic | 10.73262 | 0.000697 | 26.92751 | 26.84302 | 26.86773 |
| Q9UHF7 | Zinc finger t | TRPS1 | Zinc finger t | 12.43442 | 3.54E-05 | 26.34591 | 26.18675 | 26.40746 |
| Q9H3F6 | BTB/POZ dc | KCTD10 | BTB/POZ dc | 10.81996 | 0.028136 | 26.45808 | 27.48738 | 27.63858 |
| Q07666 | KH domain- | KHDRBS1 | KH domain- | 11.74908 | 0.000628 | 26.73467 | 26.83087 | 26.89134 |
| P62913 | 60S ribosor | RPL11 | Large ribosc | 8.746763 | 0.000846 | 28.36903 | 28.77809 | 27.70286 |
| P62854;Q5 | 40S ribosor | RPS26;RPS | Small ribos | 15.40607 | 3.31E-05 | 28.92844 | 28.67101 | 28.90113 |
| P62910 | 60S ribosor | RPL32 | Large ribosc | 14.25801 | 0.000347 | 27.58706 | 27.27604 | 27.49518 |
| Q12905 | Interleukin | ILF2 | Interleukin | 13.67681 | 0.00059 | 26.74548 | 26.43025 | 26.58449 |
| P47914 | 60S ribosor | RPL29 | Large ribosc | 9.937965 | 0.000707 | 28.45046 | 28.93253 | 28.59989 |
| P62888 | 60S ribosor | RPL30 | Large ribosc | 11.21459 | 0.012637 | 28.29758 | 26.48743 | 28.52332 |
| Q15233 | Non-POU d | NONO | Non-POU d | 3.948858 | 0.030182 | 26.13796 | 25.71707 | 26.28175 |
| Q9Y3I0 | tRNA-splicir | RTCB | RNA-splicin | 12.46576 | 1.04E-05 | 27.26051 | 27.01116 | 27.17293 |
| P60866 | 40S ribosor | RPS20 | Small ribos | 6.47485 | 0.010566 | 28.2109 | 27.90782 | 28.0776 |
| O60841 | Eukaryotic t | EIF5B | Eukaryotic t | 10.76089 | 0.001305 | 26.86432 | 26.99453 | 27.09265 |
| P46777 | 60S ribosor | RPL5 | Large ribosc | 12.81108 | 0.006656 | 27.41369 | 26.8007 | 26.91205 |
| O95793 | Double-stra | STAU1 | Double-stra | 13.17917 | 1.59E-05 | 25.78272 | 25.48312 | 25.64419 |
| O00571;O1 | ATP-depenc | DDX3X;DD | ATP-depenc | 9.882279 | 0.006135 | 25.72002 | 25.57664 | 25.74891 |
| Q96SI1 | BTB/POZ dc | KCTD15 | BTB/POZ dc | 13.12336 | 7.91E-05 | 25.95868 | 26.21638 | 25.99723 |
| Q9NYF8 | Bcl-2-assoc | BCLAF1 | Bcl-2-assoc | 10.0993 | 0.001928 | 26.168 | 26.05815 | 26.09485 |
| Q8IVF2 | Protein AHN | AHNAK2 | Protein AHN | 9.25832 | 1.01E-05 | 25.59744 | 25.2775 | 24.48025 |
| O60716 | Catenin del | CTNND1 | Catenin del | 10.72982 | 0.000139 | 26.97826 | 25.79521 | 27.12445 |
| P43243 | Matrin-3 | MATR3 | Matrin-3 OS | 12.28417 | 0.000891 | 26.38187 | 26.38334 | 26.59178 |
| P31943 | Heterogene | HNRNPH1 | Heterogene | 11.09208 | 0.011103 | 27.84266 | 27.54149 | 27.5969 |
| P55081 | Microfibrilla | MFAP1 | Microfibrilla | 4.209181 | 0.000468 | 26.06878 | 25.85956 | 26.59705 |
| P84103 | Serine/argin | SRSF3 | Serine/argin | 11.93617 | 0.005849 | 26.36453 | 28.5241 | 28.69169 |
| P60903 | Protein S10 | S100A10 | Protein S10 | 8.375788 | 3.08E-05 | 28.85377 | 28.72293 | 28.34875 |
| Q9Y697 | Cysteine de | NFS1 | Cysteine de | 14.23317 | 2.46E-05 | 26.30859 | 26.11428 | 26.29258 |
| P09493 | Tropomyos | TPM1 | Tropomyos | 4.262768 | 0.005308 | 26.75392 | 26.81433 | 27.47816 |
| P63173 | 60S ribosor | RPL38 | Large ribosc | 9.115784 | 0.009367 | 28.58273 | 28.46972 | 28.50309 |
| Q96HS1 | Serine/threc | PGAM5 | Serine/threc | 12.52053 | 0.000248 | 26.06201 | 27.12859 | 26.18391 |
| Q9BVG8 | Kinesin-like | KIFC3 | Kinesin-like | 11.28215 | 8.51E-05 | 25.02682 | 24.9049 | 25.12751 |
| Q13310 | Polyadenyl | PABPC4 | Polyadenyl | 8.372602 | 0.00071 | 26.51644 | 28.60424 | 26.53508 |
| Q9UGU0 | Transcriptio | TCF20 | Transcriptio | 12.6004 | 4.63E-05 | 25.80784 | 25.01448 | 25.82723 |
| P84090 | Enhancer o | ERH | Enhancer o | 10.98044 | 0.000243 | 27.78113 | 27.55552 | 27.72793 |
| P13010 | X-ray repair | XRCC5 | X-ray repair | 9.126753 | 0.003985 | 26.07333 | 25.98233 | 26.02115 |
| P35658 | Nuclear por | NUP214 | Nuclear por | 11.9563 | 5.94E-05 | 26.0394 | 25.79422 | 25.74597 |
| P49411 | Elongation | TUFM | Elongation | 9.660715 | 0.002753 | 25.8903 | 27.24245 | 27.39067 |
| P32969 | 60S ribosor | RPL9 | Large ribosc | 5.151283 | 0.015264 | 26.9516 | 26.62826 | 26.81702 |
| O43143 | Pre-mRNA-s | DHX15 | ATP-depenc | 9.426997 | 6.88E-05 | 25.69548 | 25.51346 | 25.75241 |
| Q9BYW2 | Histone-lys | SETD2 | Histone-lys | 10.8976 | 0.00141 | 24.83755 | 24.70277 | 24.94427 |
| Q01130 | Serine/argin | SRSF2 | Serine/argin | 5.864452 | 0.00019 | 27.35136 | 27.17617 | 27.40219 |
| Q9UDY2 | Tight junctic | TJP2 | Tight junctic | 12.10599 | 0.000304 | 23.83103 | 25.13387 | 25.21223 |
| O43809 | Cleavage ar | NUDT21 | Cleavage ar | 8.877431 | 0.001701 | 26.62617 | 26.49043 | 26.86891 |
| Q5SW79 | Centrosom | CEP170 | Centrosom | 8.834185 | 0.020894 | 27.28295 | 22.87425 | 22.99704 |
| P33992 | DNA replica | MCM5 | DNA replica | 11.78782 | 0.000282 | 25.4982 | 25.10398 | 25.47853 |
| Q92841 | Probable A | DDX17 | Probable A | 10.4557 | 0.002013 | 26.13811 | 25.60925 | 25.57868 |
| P38919 | Eukaryotic i | EIF4A3 | Eukaryotic i | 11.74845 | 0.000604 | 27.21002 | 26.35252 | 26.67828 |

| | | | | | | | | |
|---|---|---|---|---|---|---|---|---|
| Q86V81 | THO compl | ALYREF | THO compl | 3.671442 | 0.040486 | 26.80501 | 26.79017 | 27.29588 |
| Q13573 | SNW doma | SNW1 | SNW doma | 11.72863 | 0.005263 | 26.36791 | 24.2604 | 26.32631 |
| P62841 | 40S ribosor | RPS15 | Small ribos | 9.284569 | 0.003268 | 27.25115 | 27.61835 | 27.74015 |
| Q7L2E3 | Putative ATI | DHX30 | ATP-depend | 11.15437 | 3.69E-07 | 25.41671 | 25.32261 | 25.33919 |
| Q99880;Q9 | Histone H2I | HIST1H2BL | Histone H2I | 8.468016 | 6.29E-05 | 28.54134 | 28.16581 | 28.40088 |
| P07910;P0 | Heterogene | HNRNPC;H | Heterogene | 12.54226 | 0.000864 | 26.73635 | 26.45309 | 26.78619 |
| Q92791 | Synaptonei | LEPREL4 | Endoplasm | 5.118354 | 0.00394 | 27.21225 | 26.84254 | 24.53386 |
| P46783 | 40S ribosor | RPS10 | Small ribos | 3.537247 | 0.033857 | 27.21651 | 26.59078 | 26.83978 |
| O75533 | Splicing fac | SF3B1 | Splicing fac | 5.429484 | 0.000286 | 24.78581 | 24.70192 | 24.95385 |
| P01031 | Compleme | C5 | Compleme | 11.99802 | 1.58E-06 | 26.58219 | 26.7128 | 26.70965 |
| P35222 | Catenin bet | CTNNB1 | Catenin bet | 11.99786 | 0.000676 | 25.92073 | 25.69498 | 25.92291 |
| Q8IUG5 | Unconventi | MYO18B | Unconventi | 8.051589 | 0.001033 | 23.81646 | 23.66644 | 26.03521 |
| Q8IYB3 | Serine/argin | SRRM1 | Serine/argin | 8.464254 | 0.000461 | 26.44081 | 26.25877 | 26.33173 |
| Q9H6W3 | Bifunctiona | NO66 | Ribosomal | 11.86289 | 0.0002 | 26.22806 | 25.85472 | 25.90355 |
| P38159;Q9 | RNA-bindin | RBMX;RBM | RNA-bindin | 11.27994 | 0.002147 | 25.86971 | 25.6142 | 25.76684 |
| Q5SSJ5 | Heterochroi | HP1BP3 | Heterochroi | 12.09154 | 0.000201 | 26.77077 | 25.3025 | 24.91228 |
| P10412;P1 | Histone H1. | HIST1H1E;I | Histone H1. | 7.258095 | 0.000431 | 27.91016 | 28.19348 | 28.0425 |
| Q92576 | PHD finger p | PHF3 | PHD finger p | 11.95222 | 0.000788 | 25.84661 | 26.06739 | 25.71072 |
| P27169 | Serum para | PON1 | Serum para | 6.329823 | 0.003535 | 28.61161 | 28.34299 | 28.13501 |
| Q13435 | Splicing fac | SF3B2 | Splicing fac | 4.674305 | 0.00014 | 25.7465 | 25.42332 | 25.80312 |
| Q9HCD6 | Protein TAN | TANC2 | Protein TAN | 12.52192 | 0.010267 | 26.55961 | 26.69724 | 30.18376 |
| Q92499 | ATP-depend | DDX1 | ATP-depend | 9.906712 | 0.001885 | 26.53401 | 26.29102 | 25.84859 |
| Q14118 | Dystroglyca | DAG1 | Dystroglyca | 12.50485 | 0.00022 | 25.46242 | 25.45967 | 25.53561 |
| P21333 | Filamin-A | FLNA | Filamin-A O | 5.554386 | 0.003215 | 24.64807 | 24.2827 | 24.68153 |
| O75165 | DnaJ homo | DNAJC13 | DnaJ homo | 10.40303 | 0.000335 | 25.22586 | 23.27142 | 23.50069 |
| Q9BUJ2 | Heterogene | HNRNPUL1 | Heterogene | 10.9567 | 0.00062 | 25.62274 | 25.16588 | 25.6709 |
| Q9NVP1 | ATP-depend | DDX18 | ATP-depend | 13.59276 | 2.82E-05 | 26.02682 | 24.54486 | 25.93256 |
| Q13829 | BTB/POZ dc | TNFAIP1 | BTB/POZ dc | 14.54103 | 0.000125 | 26.6983 | 26.49998 | 26.71973 |
| Q9BUF5 | Tubulin bet | TUBB6 | Tubulin bet | 11.65239 | 0.00478 | 26.42472 | 26.3892 | 26.44859 |
| P21980 | Protein-glut | TGM2 | Protein-glut | 10.95165 | 0.000362 | 24.93481 | 26.54078 | 26.42339 |
| Q13151 | Heterogene | HNRNPA0 | Heterogene | 8.602475 | 0.007525 | 26.91251 | 25.98548 | 26.99723 |
| Q16629 | Serine/argin | SRSF7 | Serine/argin | 12.67161 | 0.002102 | 26.73867 | 26.55871 | 26.68793 |
| O95810 | Serum depr | SDPR | Caveolae-as | 10.10861 | 0.002217 | 27.5369 | 25.6916 | 25.78589 |
| Q14444 | Caprin-1 | CAPRIN1 | Caprin-1 OS | 6.295597 | 0.006688 | 26.68378 | 26.4325 | 26.36721 |
| P17987 | T-complex p | TCP1 | T-complex p | 9.299987 | 0.000523 | 28.52951 | 27.48838 | 25.78322 |
| O00159 | Unconventi | MYO1C | Unconventi | 7.788104 | 0.003936 | 25.1268 | 25.2314 | 25.93661 |
| P11387 | DNA topois | TOP1 | DNA topois | 3.904925 | 5.09E-05 | 25.45986 | 25.27331 | 25.39558 |
| P49792 | E3 SUMO-p | RANBP2 | E3 SUMO-p | 10.3319 | 0.000107 | 24.65818 | 24.35003 | 24.6485 |
| O75400 | Pre-mRNA-p | PRPF40A | Pre-mRNA-p | 2.56992 | 0.000606 | 25.9198 | 25.651 | 26.15457 |
| Q9UKV3 | Apoptotic c | ACIN1 | Apoptotic c | 10.85635 | 0.000178 | 26.10803 | 25.1795 | 23.84929 |
| Q9NPA8 | Transcriptio | ENY2 | Transcriptio | 5.21572 | 0.016934 | 29.13092 | 25.22229 | 25.29537 |
| O95400 | CD2 antiger | CD2BP2 | CD2 antiger | 12.11313 | 0.010677 | 24.74468 | 27.76989 | 24.84441 |
| P27708 | CAD proteir | CAD | CAD proteir | 11.9442 | 2.10E-05 | 24.92764 | 24.86179 | 25.04302 |
| Q16643 | Drebrin | DBN1 | Drebrin OS= | 5.536641 | 0.003884 | 25.55766 | 25.38578 | 26.1403 |
| P46013 | Antigen KI-6 | MKI67 | Proliferatior | 5.626944 | 0.002968 | 24.90362 | 25.20419 | 24.74175 |
| Q14690 | Protein RRP | PDCD11 | Protein RRP | 9.233534 | 0.003529 | 25.162 | 24.81831 | 24.2553 |
| Q9NWB6 | Arginine anc | ARGLU1 | Arginine anc | 0.843501 | 0.007148 | 25.96926 | 25.75407 | 26.2759 |
| Q96QC0 | Serine/threc | PPP1R10 | Serine/threc | 9.839097 | 0.00121 | 26.45605 | 25.01125 | 25.17268 |

| | | | | | | | | |
|---|---|---|---|---|---|---|---|---|
| O14683 | Tumor prot | TP53I11 | Tumor prot | 12.25922 | 0.000621 | 25.70617 | 26.73557 | 26.87631 |
| Q6PCB7 | Long-chain | SLC27A1 | Long-chain | 11.7233 | 0.000124 | 25.15217 | 25.1411 | 25.16955 |
| Q7L014 | Probable AT | DDX46 | Probable AT | 1.779706 | 0.010084 | 24.57173 | 24.27056 | 25.16699 |
| P23528;Q9 | Cofilin-1;Co | CFL1;CFL2 | Cofilin-1 OS | 4.928462 | 0.000769 | 26.3434 | 26.31752 | 26.5349 |
| P22087 | rRNA 2-O-m | FBL | rRNA 2-O-m | 12.53503 | 0.004196 | 26.29827 | 25.69906 | 25.57311 |
| P49756 | RNA-bindin | RBM25 | RNA-bindin | 2.183235 | 0.001069 | 25.52352 | 25.28613 | 25.72637 |
| Q9NVI7 | ATPase fam | ATAD3A | ATPase fam | 7.968064 | 0.000145 | 25.7984 | 24.27491 | 25.79855 |
| P02545 | Prelamin-A/ | LMNA | Prelamin-A/ | 1.47699 | 0.162043 | 27.70675 | 27.08801 | 26.93089 |
| P62861 | 40S ribosor | FAU | Ubiquitin-lil | 3.72006 | 0.008463 | 27.56761 | 27.5343 | 27.6633 |
| Q96PK6 | RNA-bindin | RBM14 | RNA-bindin | 5.044727 | 0.016563 | 25.56616 | 25.49796 | 25.9844 |
| P82650 | 28S ribosor | MRPS22 | Small ribos | 8.134302 | 1.35E-08 | 25.71427 | 25.58368 | 25.68509 |
| Q14498 | RNA-bindin | RBM39 | RNA-bindin | 2.358063 | 0.000769 | 25.75913 | 25.4249 | 25.75251 |
| Q09666 | Neuroblast | AHNAK | Neuroblast | 7.127731 | 0.006245 | 25.40419 | 24.87891 | 25.25544 |
| P25205 | DNA replica | MCM3 | DNA replica | 11.6463 | 0.000225 | 24.89837 | 24.61403 | 24.69057 |
| Q07065 | Cytoskeleto | CKAP4 | Cytoskeleto | 8.197094 | 0.020315 | 24.8242 | 24.45425 | 24.76263 |
| Q9Y224 | UPF0568 p | C14orf166 | RNA transcr | 11.81848 | 0.00025 | 25.49677 | 25.30065 | 25.53124 |
| P0DI83 | Ras-related | RAB34 | Ras-related | 11.44806 | 0.001937 | 27.06487 | 27.05248 | 27.23187 |
| P11388 | DNA topois | TOP2A | DNA topois | 8.251927 | 0.000736 | 24.63362 | 23.92642 | 25.4038 |
| P46782 | 40S ribosor | RPS5 | Small ribos | 6.147282 | 0.006295 | 26.924 | 26.52752 | 26.84661 |
| Q9P258 | Protein RCC | RCC2 | Protein RCC | 11.18745 | 0.00621 | 26.0425 | 25.89751 | 26.07953 |
| Q14103 | Heterogene | HNRNPD | Heterogene | 8.576066 | 0.001411 | 27.22177 | 26.0831 | 27.23306 |
| Q9P2E9 | Ribosome-l | RRBP1 | Ribosome-l | 11.37062 | 2.03E-05 | 24.13993 | 24.15731 | 24.65376 |
| Q96T58 | Msx2-intera | SPEN | Msx2-intera | 10.49739 | 0.000458 | 25.52636 | 24.81753 | 25.29544 |
| Q9Y383 | Putative RN | LUC7L2 | Putative RN | 3.435976 | 0.000175 | 25.98333 | 25.84563 | 26.24313 |
| Q15678 | Tyrosine-pr | PTPN14 | Tyrosine-pr | 2.587894 | 0.126798 | 27.84128 | 26.09376 | 27.80815 |
| Q15393 | Splicing fac | SF3B3 | Splicing fac | 5.527947 | 0.000733 | 24.44607 | 24.6698 | 24.46812 |
| P10809 | 60 kDa heat | HSPD1 | 60 kDa heat | 5.451792 | 0.002292 | 25.00933 | 25.06846 | 25.06595 |
| P42677 | 40S ribosor | RPS27 | Small ribos | 11.07228 | 0.000919 | 26.65772 | 26.45345 | 26.67331 |
| O95239 | Chromosor | KIF4A | Chromosor | 9.178984 | 5.40E-05 | 24.33926 | 24.21519 | 24.31686 |
| P35637 | RNA-bindin | FUS | RNA-bindin | 2.967424 | 0.049618 | 26.20034 | 25.82065 | 26.93382 |
| Q00610 | Clathrin hea | CLTC | Clathrin hea | 5.86926 | 0.020478 | 24.91953 | 24.73536 | 25.33207 |
| Q14839 | Chromodor | CHD4 | Chromodor | 11.27224 | 2.29E-05 | 24.29978 | 24.31347 | 24.16791 |
| Q14676 | Mediator of | MDC1 | Mediator of | 11.43189 | 0.000553 | 25.61082 | 25.50561 | 25.68223 |
| Q9BQG0 | Myb-bindin | MYBBP1A | Myb-bindin | 10.48527 | 0.002547 | 22.73256 | 24.97277 | 25.07737 |
| P12268 | Inosine-5-n | IMPDH2 | Inosine-5-n | 7.665135 | 0.000249 | 24.55625 | 25.62224 | 25.53234 |
| P35268 | 60S ribosor | RPL22 | Large ribosc | 5.86959 | 0.002503 | 27.42966 | 27.29325 | 27.43849 |
| P33993 | DNA replica | MCM7 | DNA replica | 10.68584 | 0.002964 | 24.93881 | 25.35202 | 25.01253 |
| P0DMV9;P( | Heat shock | HSPA1B;HS | Heat shock | 4.966745 | 0.005112 | 24.98485 | 24.76593 | 24.84057 |
| Q8NE71 | ATP-binding | ABCF1 | ATP-binding | 7.500042 | 0.019859 | 24.49924 | 24.4535 | 24.56017 |
| Q9HDC9 | Adipocyte p | APMAP | Adipocyte p | 3.085224 | 0.124884 | 27.52701 | 27.32646 | 27.7051 |
| O00567 | Nucleolar p | NOP56 | Nucleolar p | 11.52398 | 6.47E-06 | 25.23714 | 24.9805 | 24.78736 |
| Q96EY7 | Pentatricop | PTCD3 | Small ribos | 13.8342 | 4.12E-06 | 25.8157 | 25.60281 | 25.78962 |
| P82933 | 28S ribosor | MRPS9 | Small ribos | 8.750508 | 0.000601 | 25.85505 | 25.5251 | 24.30911 |
| Q9H0A0 | N-acetyltrar | NAT10 | RNA cytidin | 11.3775 | 5.05E-06 | 24.94873 | 24.72416 | 24.9423 |
| P62316 | Small nucle | SNRPD2 | Small nucle | 9.435141 | 0.050126 | 27.14861 | 27.02549 | 27.57029 |
| Q9HCE1 | Putative hel | MOV10 | Helicase MC | 10.3307 | 1.61E-07 | 24.28835 | 23.9581 | 24.27306 |
| P15924 | Desmoplak | DSP | Desmoplak | 4.346822 | 0.16337 | 25.9035 | 24.28729 | 24.22597 |
| O14979 | Heterogene | HNRNPDL | Heterogene | 12.09678 | 0.007393 | 26.94659 | 26.54821 | 26.60922 |

| | | | | | | | | |
|---|---|---|---|---|---|---|---|---|
| Q99729 | Heterogene | HNRNPAB | Heterogene | 11.61557 | 0.001933 | 26.16663 | 26.11388 | 26.23635 |
| Q7Z460 | CLIP-assoc | CLASP1 | CLIP-assoc | 11.12203 | 1.85E-05 | 24.86032 | 24.96759 | 24.9697 |
| Q92552 | 28S ribosoı | MRPS27 | Small ribos | 10.95879 | 0.003639 | 26.97509 | 26.7442 | 26.8236 |
| P05976;P0 | Myosin ligh | MYL1;MYL3 | Myosin ligh | 6.239691 | 0.002476 | 27.60915 | 27.7078 | 28.33219 |
| P63244 | Guanine nu | GNB2L1 | Small ribos | 7.193154 | 0.005662 | 25.92266 | 25.81553 | 26.05393 |
| Q8IY81 | pre-rRNA pr | FTSJ3 | pre-rRNA 2- | 12.89496 | 3.94E-06 | 25.06628 | 25.03376 | 25.18518 |
| P62995 | Transforme | TRA2B | Transforme | 13.87298 | 6.98E-05 | 26.12613 | 26.10057 | 26.1362 |
| Q8TDN6 | Ribosome b | BRIX1 | Ribosome b | 13.14299 | 0.000231 | 25.91246 | 25.92958 | 26.02335 |
| Q9Y2R9 | 28S ribosoı | MRPS7 | Small ribos | 7.761148 | 0.001229 | 26.08714 | 25.15607 | 25.3931 |
| Q96N67 | Dedicator o | DOCK7 | Dedicator o | 9.761524 | 1.25E-05 | 24.27689 | 24.87291 | 25.28694 |
| Q00325 | Phosphate | SLC25A3 | Solute carri | 9.198836 | 0.009789 | 26.39926 | 26.19479 | 26.62659 |
| Q13243 | Serine/argin | SRSF5 | Serine/argin | 14.3882 | 7.32E-06 | 26.31734 | 26.14211 | 26.29213 |
| Q6UB35 | Monofuncti | MTHFD1L | Monofuncti | 7.799019 | 0.006941 | 24.30062 | 24.15831 | 25.15584 |
| P13639 | Elongation | EEF2 | Elongation | 8.286987 | 0.001537 | 25.38828 | 25.07533 | 25.0496 |
| Q16630 | Cleavage ar | CPSF6 | Cleavage ar | 7.195659 | 0.000682 | 25.07631 | 25.63536 | 25.15108 |
| Q9Y2A7 | Nck-associa | NCKAP1 | Nck-associa | 10.15328 | 0.023925 | 24.44088 | 24.3199 | 26.44188 |
| Q6PKG0 | La-related p | LARP1 | La-related p | 10.42829 | 0.001396 | 24.60462 | 24.35299 | 24.58927 |
| P06576 | ATP syntha | ATP5B | ATP syntha | 5.996464 | 0.045157 | 25.36833 | 25.13061 | 25.10597 |
| Q9GZT3 | SRA stem-lo | SLIRP | SRA stem-lo | 6.922012 | 0.003832 | 26.37255 | 26.10883 | 26.09093 |
| P49327 | Fatty acid sy | FASN | Fatty acid sy | 7.736866 | 0.000188 | 25.90192 | 25.5322 | 24.08132 |
| Q96CW1 | AP-2 compl | AP2M1 | AP-2 compl | 10.70573 | 1.31E-05 | 25.59087 | 24.96504 | 25.43793 |
| Q9Y223 | Bifunctiona | GNE | Bifunctiona | 11.13613 | 0.000989 | 24.43307 | 24.26707 | 24.57202 |
| Q01780 | Exosome co | EXOSC10 | Exosome co | 11.23877 | 0.000133 | 24.51971 | 24.30556 | 24.49057 |
| Q9UBB9 | Tuftelin-inte | TFIP11 | Tuftelin-inte | 10.61461 | 1.04E-06 | 24.28954 | 24.22251 | 24.8615 |
| A6ND36 | Protein FAM | FAM83G | Protein FAM | 10.04758 | 0.0027 | 24.26456 | 25.33685 | 24.39885 |
| Q59GN2;P6 | Putative 60S | RPL39P5;R | Putative ribo | 3.887218 | 0.000782 | 27.69525 | 27.33083 | 27.57355 |
| Q92769 | Histone dea | HDAC2 | Histone dea | 11.84738 | 9.92E-05 | 25.38258 | 25.11957 | 25.38897 |
| P26368 | Splicing fac | U2AF2 | Splicing fac | 5.160448 | 0.000695 | 25.58006 | 25.77668 | 25.9104 |
| Q07157 | Tight junctic | TJP1 | Tight junctic | 9.469402 | 0.001127 | 24.2203 | 24.19768 | 24.83113 |
| Q13283 | Ras GTPase | G3BP1 | Ras GTPase | 9.491087 | 0.015611 | 25.58294 | 25.52987 | 25.58995 |
| O43242 | 26S proteas | PSMD3 | 26S proteas | 1.728188 | 0.29479 | 26.55985 | 26.88064 | 27.67378 |
| Q9NZ01 | Very-long-c | TECR | Very-long-c | 10.98262 | 0.001348 | 26.13195 | 24.66899 | 25.99339 |
| P60842 | Eukaryotic i | EIF4A1 | Eukaryotic i | 7.506214 | 0.015731 | 25.24876 | 25.08946 | 25.19126 |
| Q9H307 | Pinin | PNN | Pinin OS=H | 13.12006 | 0.00011 | 26.10129 | 25.8319 | 25.99185 |
| O15042 | U2 snRNP-a | U2SURP | U2 snRNP-a | 6.765359 | 0.000137 | 26.97246 | 25.60575 | 26.27936 |
| Q86Y91 | Kinesin-like | KIF18B | Kinesin-like | 7.679937 | 0.033257 | 26.04183 | 25.04439 | 23.41764 |
| O75534 | Cold shock | CSDE1 | Cold shock | 10.67366 | 0.001138 | 24.90573 | 24.68859 | 23.75399 |
| Q13268 | Dehydroger | DHRS2 | Dehydroger | 3.987349 | 0.003039 | 25.73474 | 25.62399 | 25.72632 |
| P00734 | Prothrombi | F2 | Prothrombi | 8.938124 | 0.000578 | 26.77153 | 26.69392 | 26.6698 |
| Q9Y230 | RuvB-like 2 | RUVBL2 | RuvB-like 2 | 9.290932 | 0.007931 | 23.82163 | 25.25108 | 23.73727 |
| P04792 | Heat shock | HSPB1 | Heat shock | 3.371098 | 0.030396 | 25.79357 | 25.45745 | 25.19731 |
| P14923 | Junction pla | JUP | Junction pla | 5.91835 | 0.068726 | 25.20038 | 24.50513 | 24.75552 |
| P27816 | Microtubule | MAP4 | Microtubule | 5.633565 | 0.018342 | 25.26896 | 25.08837 | 25.1164 |
| Q15287 | RNA-bindin | RNPS1 | RNA-bindin | 12.84712 | 0.000601 | 25.95762 | 25.89352 | 25.78778 |
| Q8NF37 | Lysophosp | LPCAT1 | Lysophosp | 12.42234 | 6.96E-07 | 25.28355 | 25.15734 | 25.76155 |
| Q8TDB6 | E3 ubiquitir | DTX3L | E3 ubiquitir | 11.71914 | 0.000576 | 24.86311 | 24.72941 | 24.84929 |
| Q96AG4 | Leucine-rich | LRRC59 | Leucine-rich | 4.382474 | 0.009648 | 26.16877 | 25.31195 | 25.63838 |
| P51398 | 28S ribosoı | DAP3 | Small ribos | 13.58042 | 5.15E-06 | 25.74109 | 25.53668 | 25.79858 |

| | | | | | | | | |
|---|---|---|---|---|---|---|---|---|
| Q8N684 | Cleavage ar | CPSF7 | Cleavage ar | 6.136028 | 0.001411 | 25.28394 | 24.88702 | 25.26821 |
| P48047 | ATP syntha: | ATP5O | ATP syntha: | 4.696931 | 0.037818 | 26.17619 | 24.85538 | 25.2352 |
| Q14807 | Kinesin-like | KIF22 | Kinesin-like | 10.86379 | 0.000519 | 25.18786 | 25.18722 | 25.25035 |
| Q7L576 | Cytoplasmi | CYFIP1 | Cytoplasmi | 11.27158 | 0.004154 | 24.34415 | 24.1848 | 24.34116 |
| Q9NTI5 | Sister chron | PDS5B | Sister chron | 2.90937 | 0.058441 | 26.39913 | 24.13461 | 24.31036 |
| Q14764 | Major vault | MVP | Major vault | 4.644842 | 0.014407 | 24.1817 | 24.12849 | 24.56128 |
| Q9Y3F4 | Serine-threc | STRAP | Serine-threc | 10.8396 | 0.000137 | 25.57473 | 25.40015 | 24.77886 |
| P62273 | 40S ribosor | RPS29 | Small ribos | 4.160185 | 0.009915 | 27.62554 | 27.17617 | 27.51796 |
| Q06587 | E3 ubiquitir | RING1 | E3 ubiquitir | 12.75967 | 0.000248 | 25.62976 | 25.60947 | 25.60919 |
| P36542 | ATP syntha: | ATP5C1 | ATP syntha: | 7.361031 | 0.008612 | 25.42291 | 25.11394 | 25.27323 |
| P07602 | Prosaposin | PSAP | Prosaposin | 14.37686 | 6.63E-05 | 26.70108 | 26.49072 | 26.74124 |
| Q14966 | Zinc finger p | ZNF638 | Zinc finger p | 11.05712 | 0.000373 | 25.30344 | 24.11124 | 23.62567 |
| O75122 | CLIP-assoc | CLASP2 | CLIP-assoc | 10.60438 | 5.48E-05 | 23.59446 | 23.69349 | 24.95247 |
| P78371 | T-complex p | CCT2 | T-complex p | 10.71528 | 0.000129 | 24.94311 | 25.41764 | 25.32721 |
| Q15717 | ELAV-like pı | ELAVL1 | ELAV-like pı | 10.38985 | 0.000603 | 25.2078 | 24.9522 | 25.56392 |
| P63104 | 14-3-3 prot | YWHAZ | 14-3-3 prot | 6.697039 | 0.06145 | 26.76509 | 25.97891 | 26.02726 |
| O95782 | AP-2 compl | AP2A1 | AP-2 compl | 10.76928 | 0.000816 | 24.49381 | 24.19535 | 24.37789 |
| Q99848 | Probable rR | EBNA1BP2 | Probable rR | 10.23266 | 0.00203 | 26.27482 | 26.03781 | 23.82755 |
| P40937 | Replication | RFC5 | Replication | 12.60693 | 1.98E-06 | 25.14067 | 25.09063 | 25.24822 |
| P38646 | Stress-70 p | HSPA9 | Stress-70 p | 4.162488 | 0.000122 | 24.514 | 24.53659 | 24.46526 |
| Q6UXN9 | WD repeat-c | WDR82 | WD repeat-c | 11.30353 | 0.000355 | 25.41351 | 25.15526 | 25.36175 |
| P52292 | Importin su | KPNA2 | Importin su | 10.29961 | 0.000457 | 24.95731 | 24.81191 | 25.04101 |
| P23258;Q9 | Tubulin gan | TUBG1;TUE | Tubulin gan | 8.417503 | 2.93E-05 | 24.63904 | 24.49564 | 24.97893 |
| P25398 | 40S ribosor | RPS12 | Small ribos | 8.574065 | 0.0025 | 26.20226 | 26.04293 | 26.20094 |
| P41212 | Transcriptio | ETV6 | Transcriptio | 10.47455 | 0.010159 | 23.81431 | 26.36489 | 23.81948 |
| P52701 | DNA misma | MSH6 | DNA misma | 10.43699 | 0.000106 | 24.31652 | 24.21104 | 24.18813 |
| Q8NAV1 | Pre-mRNA-s | PRPF38A | Pre-mRNA-s | 4.232405 | 0.000502 | 25.1686 | 24.86391 | 25.83273 |
| Q03701 | CCAAT/enh | CEBPZ | CCAAT/enh | 8.722643 | 9.54E-05 | 23.65371 | 24.807 | 25.32186 |
| Q8NCA5 | Protein FAM | FAM98A | Protein FAM | 12.62468 | 0.001585 | 25.31948 | 25.17364 | 25.29165 |
| Q96CW5 | Gamma-tub | TUBGCP3 | Gamma-tub | 7.564051 | 0.041814 | 25.32965 | 21.79042 | 22.03584 |
| Q9BWF3 | RNA-bindin | RBM4 | RNA-bindin | 11.50791 | 0.00012 | 24.44569 | 24.26428 | 24.49582 |
| Q92900 | Regulator o | UPF1 | Regulator o | 10.32117 | 0.001695 | 24.78492 | 24.53973 | 24.66563 |
| Q14257 | Reticulocall | RCN2 | Reticulocall | 10.21637 | 0.000407 | 25.21897 | 25.38073 | 25.38486 |
| Q9Y265 | RuvB-like 1 | RUVBL1 | RuvB-like 1 | 10.13244 | 0.005297 | 25.5811 | 23.41158 | 25.64927 |
| Q9Y399 | 28S ribosor | MRPS2 | Small ribos | 11.342 | 1.33E-06 | 24.99183 | 24.73299 | 24.97972 |
| Q9BY77 | Polymerase | POLDIP3 | Polymerase | 5.810755 | 0.042156 | 25.04352 | 24.99998 | 23.78556 |
| Q8IXQ6 | Poly [ADP-r | PARP9 | Protein mor | 9.519972 | 0.001567 | 24.7251 | 24.66503 | 24.79213 |
| Q96BF6 | Nucleus ac | NACC2 | Nucleus ac | 11.35264 | 0.002744 | 25.44313 | 25.19565 | 25.56011 |
| Q9Y6M1 | Insulin-like | IGF2BP2 | Insulin-like | 10.25996 | 2.31E-05 | 24.58509 | 24.43498 | 24.51599 |
| Q9UHI6 | Probable AT | DDX20 | Probable AT | 12.11556 | 0.00014 | 24.72291 | 24.55415 | 24.77415 |
| Q9H6R4 | Nucleolar p | NOL6 | Nucleolar p | 10.95086 | 3.38E-05 | 24.20292 | 24.08018 | 24.21519 |
| Q9HC52 | Chromobo: | CBX8 | Chromobo: | 9.669274 | 0.003229 | 22.92919 | 25.06378 | 25.16665 |
| Q9Y3D9 | 28S ribosor | MRPS23 | Small ribos | 12.4797 | 0.000566 | 25.30942 | 25.00928 | 25.28465 |
| P82930 | 28S ribosor | MRPS34 | Small ribos | 10.97717 | 0.018578 | 24.96552 | 24.86684 | 24.90357 |
| Q9NZB2 | Constitutive | FAM120A | Constitutive | 8.374669 | 0.002929 | 25.04519 | 23.95828 | 24.88413 |
| O94964 | Protein SOC | SOGA1 | Protein SOC | 8.014596 | 0.005007 | 25.50406 | 24.5852 | 21.13094 |
| Q5JTH9 | RRP12-like | RRP12 | RRP12-like | 12.34786 | 0.000323 | 24.36078 | 24.15607 | 24.37716 |
| P61927 | 60S ribosor | RPL37 | Large ribosc | 8.276544 | 1.33E-05 | 26.99873 | 26.68405 | 26.64238 |

| | | | | | | | | |
|---|---|---|---|---|---|---|---|---|
| Q969Q0 | 60S riboso | RPL36AL | Ribosomal | 13.28108 | 0.000143 | 25.77691 | 25.79744 | 25.84024 |
| P0DOX5;P( | Ig gamma-1 | IGHG1;IGH | Immunoglo | 4.63145 | 0.054685 | 25.92789 | 23.86032 | 27.44971 |
| Q9BSJ2 | Gamma-tuk | TUBGCP2 | Gamma-tuk | 5.956358 | 0.004421 | 23.83913 | 23.85614 | 25.22258 |
| Q9Y5B9 | FACT comp | SUPT16H | FACT comp | 11.24425 | 0.001455 | 24.52103 | 24.4263 | 24.37716 |
| O15479 | Melanoma- | MAGEB2 | Melanoma- | 10.79618 | 4.95E-05 | 25.96693 | 24.63589 | 25.80326 |
| P06396 | Gelsolin | GSN | Gelsolin OS | 4.288263 | 0.008361 | 24.89551 | 24.77077 | 24.97758 |
| Q8WVK2 | U4/U6.U5 s | SNRNP27 | U4/U6.U5 s | 1.407763 | 0.031816 | 25.75279 | 25.50655 | 26.41703 |
| P46087 | Probable 28 | NOP2 | Probable 28 | 12.50554 | 6.94E-05 | 24.60823 | 24.50894 | 24.55069 |
| Q9BVP2 | Guanine nu | GNL3 | Guanine nu | 9.521494 | 0.000357 | 25.01014 | 24.94114 | 23.79243 |
| Q96GQ7 | Probable A | DDX27 | Probable A | 9.945359 | 0.000521 | 21.7233 | 24.55157 | 24.33987 |
| Q9BW19 | Kinesin-like | KIFC1 | Kinesin-like | 11.88381 | 1.29E-05 | 24.57744 | 24.75801 | 23.78756 |
| Q08J23 | tRNA (cytos | NSUN2 | RNA cytosir | 9.459096 | 3.36E-05 | 23.9115 | 24.62962 | 24.88874 |
| Q9H0D6 | 5-3 exoribo | XRN2 | 5-3 exoribo | 8.540315 | 0.004987 | 24.28079 | 24.08578 | 24.30647 |
| P04843 | Dolichyl-dip | RPN1 | Dolichyl-dip | 8.394532 | 0.000184 | 25.35901 | 25.41538 | 23.52277 |
| Q96ME1 | F-box/LRR-r | FBXL18 | F-box/LRR-r | 12.12528 | 9.59E-05 | 24.43161 | 24.40204 | 24.47673 |
| Q92974 | Rho guanin | ARHGEF2 | Rho guanin | 9.513181 | 0.020194 | 25.01005 | 24.84427 | 24.90747 |
| P62140 | Serine/threo | PPP1CB | Serine/threo | 11.09032 | 0.000545 | 25.649 | 25.51087 | 25.59215 |
| O75934 | Pre-mRNA-s | BCAS2 | Pre-mRNA-s | 10.97036 | 0.000147 | 25.79486 | 25.56436 | 24.12163 |
| Q13242 | Serine/argin | SRSF9 | Serine/argin | 11.33447 | 0.003465 | 25.6216 | 22.81359 | 25.5378 |
| Q9H7N4 | Splicing fac | SCAF1 | Splicing fac | 4.383733 | 0.007624 | 26.58075 | 25.1885 | 28.21304 |
| Q15397 | Pumilio dor | KIAA0020 | Pumilio hor | 10.73859 | 0.016219 | 23.53386 | 23.32622 | 25.94679 |
| Q99567 | Nuclear por | NUP88 | Nuclear por | 11.02925 | 1.13E-06 | 24.64136 | 24.43759 | 24.63317 |
| P82673 | 28S riboso | MRPS35 | Small ribos | 12.15402 | 9.47E-05 | 24.6856 | 24.57202 | 24.7638 |
| Q9H0S4 | Probable A | DDX47 | Probable A | 12.81372 | 9.27E-06 | 24.27845 | 24.40295 | 24.60218 |
| Q12789 | General tran | GTF3C1 | General tran | 10.46286 | 0.000706 | 23.77254 | 22.74125 | 24.35373 |
| P14868 | Aspartate--t | DARS | Aspartate--t | 7.065467 | 0.06604 | 24.19144 | 24.65256 | 24.14569 |
| P04406 | Glyceraldeh | GAPDH | Glyceraldeh | 3.918896 | 0.088198 | 24.70725 | 25.00933 | 24.11323 |
| P41223 | Protein BUD | BUD31 | Protein BUD | 9.437287 | 0.009156 | 25.72231 | 25.65875 | 24.18026 |
| P28340 | DNA polym | POLD1 | DNA polym | 10.23577 | 0.000303 | 23.97797 | 23.88204 | 24.06538 |
| Q92917 | G patch dor | GPKOW | G-patch dor | 12.39807 | 7.55E-05 | 24.90224 | 24.78202 | 24.93279 |
| Q14692 | Ribosome b | BMS1 | Ribosome b | 9.329202 | 0.011937 | 22.4818 | 24.78771 | 22.61693 |
| P41091;Q2 | Eukaryotic t | EIF2S3;EIF: | Eukaryotic t | 6.512702 | 0.008035 | 24.73133 | 24.58789 | 24.79347 |
| P56545 | C-terminal-l | CTBP2 | C-terminal-l | 8.816113 | 5.48E-06 | 25.46364 | 24.85191 | 25.37537 |
| O60832 | H/ACA ribor | DKC1 | H/ACA ribor | 10.14308 | 0.006961 | 23.44316 | 26.12558 | 23.50288 |
| P06702 | Protein S10 | S100A9 | Protein S10 | 3.186079 | 0.5793 | 26.67882 | 20.80136 | 18.77192 |
| Q71UM5 | 40S riboso | RPS27L | Ribosomal | 8.902903 | 0.000122 | 26.22993 | 25.77693 | 26.25712 |
| Q10570 | Cleavage ar | CPSF1 | Cleavage ar | 8.169274 | 0.001049 | 24.02859 | 21.49858 | 24.06964 |
| Q9BZJ0 | Crooked ne | CRNKL1 | Crooked ne | 11.14133 | 0.001908 | 24.48984 | 24.1823 | 24.51455 |
| Q9Y2X3 | Nucleolar p | NOP58 | Nucleolar p | 10.85145 | 0.003477 | 24.87098 | 24.4872 | 24.68549 |
| Q9BXP5 | Serrate RNA | SRRT | Serrate RNA | 8.512765 | 0.020877 | 24.2712 | 24.58629 | 24.34367 |
| P82675 | 28S riboso | MRPS5 | Small ribos | 11.87035 | 0.000641 | 25.90667 | 25.57603 | 25.82343 |
| Q9ULU4 | Protein kina | ZMYND8 | MYND-type | 10.85263 | 1.03E-07 | 24.50628 | 24.22361 | 24.35454 |
| Q86U86 | Protein poly | PBRM1 | Protein poly | 11.40928 | 2.00E-05 | 24.28778 | 23.93724 | 23.12978 |
| P57678 | Gem-assoc | GEMIN4 | Gem-assoc | 11.47422 | 0.000252 | 24.12345 | 23.95633 | 24.0181 |
| P42285 | Superkiller | SKIV2L2 | Exosome R | 9.281468 | 0.005191 | 24.1885 | 23.96622 | 24.00607 |
| P40938 | Replication | RFC3 | Replication | 11.83097 | 0.000423 | 25.00299 | 24.76466 | 25.03741 |
| P52948 | Nuclear por | NUP98 | Nuclear por | 8.568663 | 0.003218 | 23.58898 | 23.47754 | 25.40659 |
| P17858 | ATP-depen | PFKL | ATP-depen | 10.36372 | 9.30E-06 | 24.48806 | 24.64971 | 24.36366 |

| | | | | | | | | |
|---|---|---|---|---|---|---|---|---|
| O00422 | Histone dea | SAP18 | Histone dea | 13.06129 | 4.08E-05 | 25.4992 | 25.14301 | 25.27149 |
| O95819 | Mitogen-act | MAP4K4 | Mitogen-act | 10.27451 | 0.000641 | 24.2815 | 23.51461 | 24.08554 |
| P37108 | Signal recog | SRP14 | Signal recog | 12.04432 | 0.000162 | 25.41254 | 25.04168 | 25.20684 |
| Q7L4I2 | Arginine/ser | RSRC2 | Arginine/ser | 1.600357 | 0.007987 | 24.68678 | 24.413 | 25.16488 |
| Q13459 | Unconventi | MYO9B | Unconventi | 7.244436 | 0.002729 | 25.01359 | 22.73167 | 24.69275 |
| P08621 | U1 small nu | SNRNP70 | U1 small nu | 8.662959 | 0.000429 | 25.52292 | 25.66907 | 25.67366 |
| P78406 | mRNA expo | RAE1 | mRNA expo | 9.89675 | 0.013999 | 26.43028 | 23.57577 | 23.81235 |
| O14578 | Citron Rho- | CIT | Citron Rho- | 6.117874 | 0.041969 | 23.49814 | 23.23923 | 24.73236 |
| Q9H2U1 | ATP-depend | DHX36 | ATP-depend | 11.19591 | 1.48E-05 | 24.40568 | 24.30159 | 24.42066 |
| P19525 | Interferon-i | EIF2AK2 | Interferon-i | 11.09849 | 0.002407 | 24.14663 | 24.32643 | 24.59685 |
| Q92665 | 28S riboso | MRPS31 | Small ribos | 12.16625 | 0.001577 | 25.15831 | 25.1689 | 25.26999 |
| Q14684 | Ribosomal | RRP1B | Ribosomal | 9.544895 | 0.031831 | 23.72536 | 23.50773 | 25.36369 |
| P55265 | Double-stra | ADAR | Double-stra | 7.406585 | 0.000625 | 23.62243 | 24.03069 | 24.23717 |
| Q9UKD2 | mRNA turno | MRTO4 | mRNA turno | 11.0601 | 0.002803 | 24.25523 | 24.08974 | 24.22986 |
| O60306 | Intron-bind | AQR | RNA helicas | 10.15914 | 0.000843 | 24.61801 | 24.47797 | 22.14401 |
| Q9UPT8 | Zinc finger C | ZC3H4 | Zinc finger C | 10.59216 | 0.000259 | 24.58262 | 24.7053 | 24.48561 |
| Q9P206 | Uncharacte | KIAA1522 | Uncharacte | 8.490037 | 0.01939 | 26.02102 | 26.10763 | 23.21488 |
| P50990 | T-complex p | CCT8 | T-complex p | 7.922047 | 0.000801 | 25.85489 | 25.03913 | 23.87558 |
| P38117 | Electron tra | ETFB | Electron tra | 11.5674 | 0.007196 | 23.10068 | 24.90564 | 25.24169 |
| O00411 | DNA-directe | POLRMT | DNA-directe | 11.13285 | 9.20E-05 | 25.09889 | 24.93751 | 25.12802 |
| Q13123 | Protein Red | IK | Protein Red | 8.686181 | 0.00056 | 24.58388 | 23.92615 | 24.37166 |
| Q04637 | Eukaryotic t | EIF4G1 | Eukaryotic t | 9.093708 | 0.00024 | 23.73138 | 24.1411 | 23.64317 |
| Q9HD34 | LYR motif-c | LYRM4 | LYR motif-c | 10.22542 | 0.010565 | 26.77228 | 24.01018 | 24.11466 |
| Q71UI9;P0 | Histone H2 | H2AFV;H2A | Histone H2 | 3.79058 | 0.001357 | 25.81039 | 25.68864 | 25.80324 |
| P63010 | AP-2 compl | AP2B1 | AP-2 compl | 10.27084 | 0.002854 | 24.64834 | 24.40529 | 24.46899 |
| Q8TF72 | Protein Shr | SHROOM3 | Protein Shr | 0.468241 | 0.015883 | 26.14165 | 26.09073 | 25.90692 |
| P47929 | Galectin-7 | LGALS7 | Galectin-7 ( | 1.951351 | 0.736934 | 26.8087 | 19.03094 | 16.66678 |
| Q96SB4 | SRSF protei | SRPK1 | SRSF protei | 7.095304 | 0.065306 | 23.92171 | 26.63284 | 23.98564 |
| Q2M1P5 | Kinesin-like | KIF7 | Kinesin-like | 9.055078 | 0.001144 | 23.47902 | 23.729 | 24.0242 |
| P68400;Q8 | Casein kina | CSNK2A1;C | Casein kina | 6.511126 | 2.98E-05 | 24.64493 | 25.35901 | 24.50288 |
| Q92804 | TATA-bindir | TAF15 | TATA-bindir | 8.304609 | 0.000192 | 26.2191 | 26.09493 | 26.39221 |
| P31942 | Heterogene | HNRNPH3 | Heterogene | 9.021837 | 0.002327 | 26.06322 | 23.69943 | 23.8399 |
| P51116 | Fragile X me | FXR2 | RNA-bindin | 7.544974 | 0.005494 | 26.32201 | 27.48185 | 23.40367 |
| Q5BKY9 | Protein FAM | FAM133B | Protein FAM | 0.656974 | 0.004003 | 24.66909 | 24.46407 | 24.69152 |
| P42166 | Lamina-ass | TMPO | Lamina-ass | 4.857599 | 4.21E-05 | 24.1776 | 23.85842 | 24.21993 |
| Q8N9M1 | Uncharacte | C19orf47 | Uncharacte | 10.93839 | 0.007005 | 24.18873 | 24.43549 | 24.51761 |
| P49368 | T-complex p | CCT3 | T-complex p | 8.083886 | 3.68E-05 | 24.39441 | 24.3377 | 24.3377 |
| P35249 | Replication | RFC4 | Replication | 8.47125 | 0.000241 | 24.85072 | 24.71356 | 22.76729 |
| P35250 | Replication | RFC2 | Replication | 10.14485 | 0.002902 | 25.58216 | 25.35333 | 25.32378 |
| P23588 | Eukaryotic t | EIF4B | Eukaryotic t | 8.383088 | 0.001148 | 25.39323 | 25.44133 | 25.67093 |
| Q86X51 | Uncharacte | CXorf67 | EZH inhibito | 10.65643 | 0.000332 | 24.70762 | 25.18317 | 24.44127 |
| P63167 | Dynein ligh | DYNLL1 | Dynein ligh | 10.71873 | 0.012201 | 26.0185 | 25.87898 | 26.07339 |
| P35221 | Catenin alp | CTNNA1 | Catenin alp | 7.330605 | 3.61E-06 | 24.55631 | 24.43905 | 24.3995 |
| P61513 | 60S riboso | RPL37A | Large ribos | 5.32144 | 0.013934 | 27.04312 | 23.3672 | 26.7661 |
| Q05519 | Serine/argin | SRSF11 | Serine/argin | 2.568506 | 5.98E-05 | 25.53611 | 25.29918 | 25.65149 |
| P19474 | E3 ubiquitir | TRIM21 | E3 ubiquitir | 8.969701 | 6.97E-05 | 24.96036 | 25.06829 | 25.36986 |
| Q9NYV4 | Cyclin-depe | CDK12 | Cyclin-depe | 11.85057 | 0.002409 | 25.03204 | 24.76714 | 24.96437 |
| O95470 | Sphingosin | SGPL1 | Sphingosin | 9.966524 | 0.009128 | 24.15012 | 23.85091 | 24.24293 |

| | | | | | | | | |
|---|---|---|---|---|---|---|---|---|
| P07814 | Bifunctiona | EPRS | Bifunctiona | 9.618167 | 3.53E-05 | 23.91698 | 22.59115 | 23.82571 |
| O14579 | Coatomer s | COPE | Coatomer s | 10.69644 | 0.003599 | 25.44244 | 23.01929 | 25.44758 |
| Q93009 | Ubiquitin ca | USP7 | Ubiquitin ca | 11.43626 | 9.05E-05 | 24.59844 | 24.56069 | 24.7016 |
| O60264 | SWI/SNF-re | SMARCA5 | SWI/SNF-re | 11.11275 | 1.57E-07 | 23.55473 | 23.47407 | 23.60761 |
| Q9NUQ6 | SPATS2-like | SPATS2L | SPATS2-like | 11.11983 | 0.000354 | 23.91278 | 23.78835 | 23.88129 |
| Q01081;Q8 | Splicing fac | U2AF1;U2A | Splicing fac | 4.629379 | 2.18E-05 | 25.91536 | 25.6525 | 25.67206 |
| Q99496 | E3 ubiquitir | RNF2 | E3 ubiquitir | 9.654783 | 0.024136 | 26.83485 | 23.03144 | 25.65565 |
| P54886 | Delta-1-pyr | ALDH18A1 | Delta-1-pyr | 9.955987 | 0.000911 | 23.58405 | 23.55625 | 23.59252 |
| O60762 | Dolichol-ph | DPM1 | Dolichol-ph | 10.24113 | 0.001596 | 24.23096 | 24.03406 | 25.50918 |
| O75306 | NADH dehy | NDUFS2 | NADH dehy | 11.74442 | 0.00106 | 25.16814 | 25.46086 | 25.3869 |
| O95573 | Long-chain- | ACSL3 | Fatty acid C | 9.62446 | 0.002711 | 24.22575 | 24.0685 | 24.29537 |
| Q9Y5B6 | PAX3- and I | PAXBP1 | PAX3- and I | 9.344043 | 0.00525 | 23.85453 | 24.12345 | 24.99213 |
| P67936 | Tropomyos | TPM4 | Tropomyos | 4.339197 | 0.008162 | 24.32629 | 24.31022 | 24.85005 |
| P62937 | Peptidyl-pr | PPIA | Peptidyl-pr | 3.149277 | 0.007696 | 26.10011 | 25.08946 | 25.1612 |
| Q9BYN8 | 28S ribosor | MRPS26 | Small ribos | 11.38955 | 4.30E-07 | 24.56122 | 24.48228 | 24.71581 |
| Q96RF0 | Sorting nexi | SNX18 | Sorting nexi | 14.31501 | 1.69E-05 | 26.03318 | 26.01504 | 26.2676 |
| Q12849 | G-rich sequ | GRSF1 | G-rich sequ | 11.53993 | 0.002417 | 25.0874 | 24.86524 | 22.3374 |
| Q13263 | Transcriptio | TRIM28 | Transcriptio | 10.43525 | 0.001635 | 24.05961 | 23.7543 | 23.95527 |
| O60231 | Putative pre | DHX16 | Pre-mRNA-s | 6.973022 | 5.62E-05 | 23.94369 | 23.16829 | 23.07364 |
| Q9UHX1 | Poly(U)-bin | PUF60 | Poly(U)-bin | 1.845448 | 0.00188 | 23.97622 | 23.77595 | 24.07863 |
| P06753 | Tropomyos | TPM3 | Tropomyos | 7.903063 | 0.036417 | 24.71654 | 24.78895 | 25.19656 |
| O43684 | Mitotic chec | BUB3 | Mitotic chec | 6.028721 | 0.001823 | 24.62148 | 24.13571 | 24.41848 |
| Q2TAY7 | WD40 repea | SMU1 | WD40 repea | 8.691575 | 0.003548 | 25.67987 | 22.66544 | 25.53931 |
| O95816 | BAG family | BAG2 | BAG family | 8.541317 | 0.000132 | 25.52683 | 25.34391 | 25.47246 |
| P02511 | Alpha-cryst | CRYAB | Alpha-cryst | 5.768549 | 1.00E-05 | 24.90674 | 24.8691 | 24.79991 |
| Q9Y5S9 | RNA-bindin | RBM8A | RNA-bindin | 13.26922 | 5.40E-05 | 25.16135 | 24.91077 | 25.10873 |
| Q96KR1 | Zinc finger F | ZFR | Zinc finger F | 8.699187 | 0.005137 | 21.29322 | 24.2422 | 24.44821 |
| Q9BPW8 | Protein NipS | NIPSNAP1 | Protein NipS | 10.36694 | 0.015046 | 25.35548 | 25.09591 | 25.59126 |
| Q96T37 | Putative RN | RBM15 | RNA-bindin | 10.86718 | 0.002003 | 23.79966 | 24.31714 | 24.37212 |
| Q6P5R6 | 60S ribosor | RPL22L1 | Ribosomal | 4.865447 | 0.004643 | 25.67933 | 25.79536 | 25.37322 |
| Q7L0Y3 | Mitochondr | TRMT10C | tRNA methy | 11.09945 | 0.001873 | 24.03783 | 23.50628 | 24.14359 |
| Q9BQ39 | ATP-depenc | DDX50 | ATP-depenc | 11.00632 | 7.82E-06 | 24.13806 | 23.86533 | 24.05854 |
| Q9UKK6 | NTF2-relate | NXT1 | NTF2-relate | 13.05069 | 3.36E-06 | 25.7476 | 25.48062 | 25.85928 |
| Q9NQT8 | Kinesin-like | KIF13B | Kinesin-like | 10.01849 | 0.073753 | 26.60767 | 26.37564 | 26.06396 |
| Q9Y676 | 28S ribosor | MRPS18B | Small ribos | 9.368429 | 0.018319 | 24.46998 | 24.50664 | 24.55976 |
| Q13428 | Treacle prot | TCOF1 | Treacle prot | 4.634265 | 0.002516 | 24.80985 | 24.99304 | 24.82114 |
| O75489 | NADH dehy | NDUFS3 | NADH dehy | 9.389198 | 0.009106 | 24.39878 | 24.21505 | 24.41086 |
| Q08945 | FACT comp | SSRP1 | FACT comp | 9.147404 | 0.013602 | 25.03477 | 20.9356 | 23.63677 |
| O94842 | TOX high m | TOX4 | TOX high m | 7.359081 | 2.17E-05 | 23.93418 | 24.70472 | 24.74919 |
| Q9UKM9 | RNA-bindin | RALY | RNA-bindin | 9.674999 | 0.00012 | 24.36379 | 24.31927 | 24.57537 |
| P49750 | YLP motif-c | YLPM1 | YLP motif-c | 11.03251 | 7.79E-05 | 23.25753 | 23.0329 | 23.10162 |
| P61619 | Protein tran | SEC61A1 | Protein tran | 10.56032 | 0.011751 | 24.60077 | 25.31907 | 24.55467 |
| P30566 | Adenylosuc | ADSL | Adenylosuc | 9.074756 | 0.019353 | 24.68416 | 24.63034 | 25.84587 |
| A6NHR9 | Structural n | SMCHD1 | Structural n | 8.801606 | 0.000424 | 22.85812 | 23.84384 | 22.9798 |
| Q9HCS7 | Pre-mRNA-s | XAB2 | Pre-mRNA-s | 8.943747 | 0.004947 | 23.90637 | 22.02052 | 23.96983 |
| P30876 | DNA-directe | POLR2B | DNA-directe | 10.21112 | 0.003606 | 24.05333 | 24.7112 | 22.27395 |
| Q9NQW6 | Actin-bindir | ANLN | Anillin OS=I | 10.03501 | 7.56E-05 | 23.95287 | 23.59923 | 23.97998 |
| P16615 | Sarcoplasm | ATP2A2 | Sarcoplasm | 10.26135 | 0.004681 | 24.30459 | 22.03996 | 24.39297 |

| | | | | | | | | |
|---|---|---|---|---|---|---|---|---|
| Q9NY93 | Probable AT | DDX56 | Probable AT | 10.79071 | 0.016256 | 23.99343 | 24.02378 | 23.73779 |
| Q06830 | Peroxiredox | PRDX1 | Peroxiredox | 5.020504 | 0.037598 | 25.03065 | 23.90674 | 24.20814 |
| Q13523 | Serine/threc | PRPF4B | Serine/threc | 3.53604 | 1.44E-07 | 24.75394 | 24.66389 | 24.76005 |
| P51532 | Transcriptio | SMARCA4 | Transcriptio | 11.78517 | 8.81E-05 | 23.54851 | 23.2369 | 23.75399 |
| P11717 | Cation-inde | IGF2R | Cation-inde | 8.039404 | 0.012962 | 23.55859 | 24.75797 | 21.73178 |
| Q96A72 | Protein mag | MAGOHB | Protein mag | 10.11731 | 0.001649 | 25.19381 | 24.89998 | 24.90587 |
| Q9UNQ2 | Probable di | DIMT1 | Probable di | 9.99772 | 0.003168 | 24.32265 | 24.29488 | 23.98059 |
| Q9UI10 | Translation | EIF2B4 | Translation | 1.203726 | 0.00454 | 24.91798 | 25.5349 | 25.35971 |
| Q14974 | Importin su | KPNB1 | Importin su | 7.827278 | 0.002434 | 23.28644 | 23.82997 | 23.45638 |
| Q9NW13 | RNA-bindin | RBM28 | RNA-bindin | 6.10331 | 0.002877 | 23.87436 | 25.10769 | 23.94092 |
| P08865;A0 | 40S ribosor | RPSA | Small ribos | 7.433194 | 0.063241 | 24.72302 | 24.32327 | 24.59895 |
| Q86UP2 | Kinectin | KTN1 | Kinectin OS | 9.216894 | 0.006655 | 24.20411 | 23.36386 | 23.3835 |
| P12236 | ADP/ATP tra | SLC25A6 | ADP/ATP tra | 9.834144 | 0.001648 | 25.24786 | 25.12888 | 25.18941 |
| P16989 | Y-box-bindi | YBX3 | Y-box-bindi | 11.47227 | 0.001507 | 25.57071 | 25.32701 | 25.48062 |
| Q8TA86 | Retinitis pig | RP9 | Retinitis pig | 1.000376 | 0.001317 | 23.9534 | 24.01044 | 24.18782 |
| Q9UNF1 | Melanoma- | MAGED2 | Melanoma- | 8.603641 | 6.97E-05 | 23.70356 | 23.62366 | 23.85975 |
| Q6PJG2 | ELM2 and S | ELMSAN1 | Mitotic deac | 3.851983 | 0.000118 | 23.96482 | 24.15931 | 24.41261 |
| Q5BKZ1 | DBIRD com | ZNF326 | DBIRD com | 9.981958 | 0.001296 | 24.44531 | 24.16177 | 24.32663 |
| Q01844 | RNA-bindin | EWSR1 | RNA-bindin | 2.714064 | 0.080716 | 24.78252 | 24.22883 | 25.77884 |
| P42704 | Leucine-rich | LRPPRC | Leucine-rich | 6.4798 | 0.000202 | 23.68014 | 22.15357 | 23.57311 |
| P09382 | Galectin-1 | LGALS1 | Galectin-1 ( | 10.37432 | 0.005143 | 25.3571 | 25.29978 | 25.30831 |
| Q8WXX5 | DnaJ homo | DNAJC9 | DnaJ homo | 11.70684 | 3.23E-07 | 24.4729 | 24.38837 | 24.49265 |
| Q9BRK5 | 45 kDa calc | SDF4 | 45 kDa calc | 6.080615 | 0.003516 | 24.79615 | 24.73231 | 24.77107 |
| Q99873 | Protein argi | PRMT1 | Protein argi | 9.253094 | 9.18E-05 | 24.75674 | 23.99473 | 24.14965 |
| O60524 | Nuclear exp | NEMF | Ribosome c | 11.20257 | 3.85E-08 | 23.24512 | 23.1504 | 23.23332 |
| Q6PK04 | Coiled-coil | CCDC137 | Coiled-coil | 1.912307 | 0.015069 | 25.10949 | 25.02968 | 25.33368 |
| O00483 | Cytochrom | NDUFA4 | Cytochrom | 10.62225 | 0.010451 | 25.5491 | 25.1057 | 25.3662 |
| P33991 | DNA replica | MCM4 | DNA replica | 8.335948 | 0.010059 | 24.819 | 22.58953 | 22.69695 |
| P30530 | Tyrosine-pr | AXL | Tyrosine-pr | 9.852856 | 0.021328 | 24.55777 | 23.11816 | 24.53469 |
| Q9Y2Q9 | 28S ribosor | MRPS28 | Small ribos | 13.3275 | 9.71E-05 | 24.8853 | 24.76875 | 24.91036 |
| Q92575 | UBX domai | UBXN4 | UBX domai | 12.72743 | 0.000138 | 25.21767 | 25.28465 | 25.27803 |
| P31689 | DnaJ homo | DNAJA1 | DnaJ homo | 11.23488 | 4.52E-05 | 25.25883 | 24.79 | 24.92678 |
| Q9NYL9 | Tropomodu | TMOD3 | Tropomodu | 6.237191 | 0.00292 | 23.88939 | 23.94387 | 24.51905 |
| Q66PJ3 | ADP-ribosyl | ARL6IP4 | ADP-ribosyl | 1.70568 | 0.001224 | 25.36132 | 25.31368 | 25.25223 |
| Q7Z2W4 | Zinc finger C | ZC3HAV1 | Zinc finger C | 10.05904 | 0.001409 | 23.74345 | 23.72078 | 23.57877 |
| O75330 | Hyaluronan | HMMR | Hyaluronan | 9.661244 | 0.035673 | 23.45939 | 24.993 | 23.56929 |
| P05109 | Protein S10 | S100A8 | Protein S10 | 2.461071 | 0.649114 | 26.79216 | 18.47979 | 17.37152 |
| P09874 | Poly [ADP-r | PARP1 | Poly [ADP-r | 6.039686 | 0.006987 | 23.76436 | 23.40574 | 23.99231 |
| Q9BY12 | S phase cyc | SCAPER | S phase cyc | 10.33127 | 0.000836 | 23.76233 | 23.38179 | 23.7082 |
| Q13868 | Exosome co | EXOSC2 | Exosome co | 10.81775 | 0.001782 | 24.84886 | 24.63633 | 24.87718 |
| O43660 | Pleiotropic | PLRG1 | Pleiotropic | 8.38397 | 0.011795 | 23.16598 | 25.27987 | 23.19819 |
| P19367 | Hexokinase | HK1 | Hexokinase | 8.724781 | 0.000601 | 23.56616 | 23.22136 | 23.40093 |
| P20042 | Eukaryotic t | EIF2S2 | Eukaryotic t | 8.206213 | 0.04269 | 24.50857 | 24.41132 | 24.10757 |
| O00139 | Kinesin-like | KIF2A | Kinesin-like | 7.842752 | 0.005358 | 23.20654 | 24.00161 | 23.47988 |
| P55209 | Nucleosom | NAP1L1 | Nucleosom | 10.06464 | 0.000341 | 23.78387 | 24.49473 | 24.75389 |
| Q13724 | Mannosyl-c | MOGS | Mannosyl-c | 5.395753 | 8.55E-05 | 23.39126 | 24.33088 | 24.44278 |
| Q9UN86 | Ras GTPase | G3BP2 | Ras GTPase | 12.43037 | 2.48E-06 | 24.88795 | 24.52564 | 24.77299 |
| Q9Y520 | Protein PRR | PRRC2C | Protein PRR | 8.828921 | 0.004985 | 23.82755 | 23.60761 | 23.97342 |

| | | | | | | | | |
|---|---|---|---|---|---|---|---|---|
| Q9UJS0 | Calcium-bi | SLC25A13 | Electrogeni | 11.61957 | 8.17E-06 | 23.97106 | 23.89652 | 24.03061 |
| P05198 | Eukaryotic t | EIF2S1 | Eukaryotic t | 8.27117 | 0.004794 | 23.31133 | 24.62567 | 25.03107 |
| P48729 | Casein kina | CSNK1A1 | Casein kina | 8.870131 | 0.014674 | 23.99231 | 23.86684 | 23.97351 |
| Q9NWH9 | SAFB-like tr | SLTM | SAFB-like tr | 6.128384 | 0.008649 | 24.14965 | 23.73727 | 24.17242 |
| Q92598 | Heat shock | HSPH1 | Heat shock | 7.509601 | 0.056544 | 22.19877 | 25.1264 | 22.22247 |
| Q6R327 | Rapamycin- | RICTOR | Rapamycin- | 10.61552 | 4.86E-05 | 23.5386 | 23.328 | 23.44455 |
| O15234 | Protein CAS | CASC3 | Protein CAS | 10.66051 | 9.60E-06 | 23.86524 | 23.63755 | 23.75369 |
| Q16576 | Histone-bin | RBBP7 | Histone-bin | 9.699529 | 0.006501 | 24.50634 | 22.38588 | 24.50149 |
| O75569 | Interferon-i | PRKRA | Interferon-i | 11.08552 | 2.92E-07 | 24.69408 | 24.6226 | 24.7025 |
| Q9GZR7 | ATP-depend | DDX24 | ATP-depend | 3.262609 | 0.008336 | 24.60258 | 22.47187 | 23.66069 |
| Q05682 | Caldesmon | CALD1 | Caldesmon | 5.1946 | 0.000692 | 23.10839 | 23.07385 | 24.02673 |
| P98175 | RNA-bindin | RBM10 | RNA-bindin | 7.591273 | 0.012197 | 23.75338 | 24.40243 | 23.87296 |
| Q8NC60 | Nitric oxide- | NOA1 | Nitric oxide- | 10.7066 | 7.50E-06 | 23.56964 | 23.5559 | 23.66796 |
| Q9UKN8 | General tran | GTF3C4 | General tran | 9.498964 | 7.22E-07 | 23.92579 | 23.58038 | 23.37915 |
| Q5T9A4 | ATPase fam | ATAD3B | ATPase fam | 1.269619 | 0.0398 | 24.03246 | 24.7381 | 24.97487 |
| Q96IZ7 | Serine/Argin | RSRC1 | Serine/Argin | 2.621415 | 0.001397 | 24.96358 | 24.99809 | 25.40048 |
| P30153 | Serine/threo | PPP2R1A | Serine/threo | 10.64052 | 0.004322 | 24.41171 | 24.15236 | 23.89837 |
| P0C0L4;P0 | Compleme | C4A;C4B | Compleme | 5.134156 | 0.000298 | 25.07602 | 24.72265 | 24.68528 |
| Q9Y3Y2 | Chromatin t | CHTOP | Chromatin t | 9.005244 | 0.014803 | 24.55116 | 24.52355 | 24.56935 |
| P53618 | Coatomer s | COPB1 | Coatomer s | 9.573834 | 0.000183 | 23.83171 | 23.52121 | 23.52707 |
| Q96EL2 | 28S riboso | MRPS24 | Small ribos | 11.6538 | 3.01E-08 | 25.24623 | 25.16408 | 25.25061 |
| Q9NS87 | Kinesin-like | KIF15 | Kinesin-like | 7.744914 | 0.005605 | 24.2292 | 21.77278 | 21.97365 |
| Q93077;Q7 | Histone H2 | HIST1H2AC | Histone H2 | 10.35343 | 0.029368 | 26.24358 | 25.80292 | 25.67139 |
| Q13148 | TAR DNA-bi | TARDBP | TAR DNA-bi | 10.49638 | 0.000522 | 24.23885 | 23.884 | 24.45111 |
| Q13509 | Tubulin bet | TUBB3 | Tubulin bet | 10.76159 | 0.000128 | 24.42573 | 24.48751 | 24.70092 |
| Q9Y5V3 | Melanoma- | MAGED1 | Melanoma- | 10.27761 | 0.002612 | 23.99886 | 21.60137 | 24.0385 |
| Q6SJ93 | Protein FAM | FAM111B | Serine prote | 5.79625 | 0.19851 | 26.07867 | 24.80671 | 24.79838 |
| Q5JTC6 | APC memb | AMER1 | APC memb | 9.768677 | 0.005877 | 24.07594 | 22.01028 | 24.19596 |
| Q6UN15 | Pre-mRNA 3 | FIP1L1 | Pre-mRNA 3 | 6.696484 | 0.000936 | 23.83016 | 23.83413 | 24.2094 |
| Q7Z478 | ATP-depend | DHX29 | ATP-depend | 9.783149 | 0.000159 | 23.24417 | 23.11339 | 23.37875 |
| Q8TER5 | Rho guanin | ARHGEF40 | Rho guanin | 9.610962 | 0.001096 | 21.20902 | 23.39976 | 23.61493 |
| Q6P1L8 | 39S riboso | MRPL14 | Large riboso | 2.844257 | 0.0363 | 24.53089 | 24.15846 | 24.58411 |
| Q02241 | Kinesin-like | KIF23 | Kinesin-like | 9.977134 | 1.79E-05 | 24.19438 | 24.12447 | 24.3505 |
| P62318 | Small nucle | SNRPD3 | Small nucle | 10.00762 | 0.030749 | 25.62803 | 25.36098 | 25.53106 |
| P04844 | Dolichyl-di | RPN2 | Dolichyl-di | 8.668143 | 0.018041 | 23.89282 | 24.74709 | 24.71816 |
| Q8WUA4 | General tran | GTF3C2 | General tran | 4.126479 | 1.61E-05 | 24.2071 | 24.17668 | 24.12361 |
| Q6PCB5 | Round sper | RSBN1L | Lysine-spec | 8.656334 | 0.011196 | 22.23119 | 22.13317 | 24.61689 |
| P41250 | Glycine--tR | GARS | Glycine--tR | 10.19089 | 0.011147 | 22.1598 | 23.79243 | 23.79134 |
| P51812 | Ribosomal | RPS6KA3 | Ribosomal | 10.6075 | 4.67E-06 | 23.60309 | 23.43237 | 23.67282 |
| P26196 | Probable A | DDX6 | Probable A | 10.0987 | 0.006548 | 24.12384 | 23.88437 | 23.78107 |
| P37198 | Nuclear po | NUP62 | Nuclear po | 12.03649 | 0.001118 | 24.85329 | 24.32065 | 24.95522 |
| Q9NUL3 | Double-stra | STAU2 | Double-stra | 9.474129 | 0.01559 | 22.8714 | 24.83253 | 22.8645 |
| Q8WVV9 | Heterogene | HNRNPLL | Heterogene | 9.743464 | 1.15E-05 | 23.64603 | 23.82561 | 24.12589 |
| P18085 | ADP-ribosyl | ARF4 | ADP-ribosyl | 7.454924 | 6.44E-05 | 24.22971 | 24.16392 | 24.87886 |
| P83881 | 60S riboso | RPL36A | Large riboso | 7.334666 | 0.067429 | 24.00547 | 23.21056 | 26.94101 |
| Q15365 | Poly(rC)-bir | PCBP1 | Poly(rC)-bir | 9.896303 | 0.009014 | 24.3377 | 24.32971 | 24.36959 |
| E9PRG8 | Uncharacte | C11orf98 | Uncharacte | 8.84679 | 0.025747 | 25.53133 | 24.41809 | 25.61709 |
| Q13557 | Calcium/ca | CAMK2D | Calcium/ca | 9.870729 | 0.01595 | 23.84642 | 25.07014 | 25.12786 |

| | | | | | | | | |
|---|---|---|---|---|---|---|---|---|
| Q2NL82 | Pre-rRNA-pı | TSR1 | Pre-rRNA-pı | 11.07641 | 2.30E-08 | 23.33101 | 23.39728 | 23.46376 |
| Q9Y2R4 | Probable AT | DDX52 | Probable AT | 10.04763 | 0.001423 | 24.28778 | 24.30208 | 24.49893 |
| Q9NPD3 | Exosome co | EXOSC4 | Exosome co | 10.9201 | 0.005483 | 24.62014 | 24.54032 | 23.98146 |
| Q5VTL8 | Pre-mRNA-s | PRPF38B | Pre-mRNA-s | 2.189095 | 0.000465 | 23.72432 | 23.4405 | 23.87961 |
| Q9Y5Q8 | General trar | GTF3C5 | General trar | 10.96561 | 3.45E-05 | 24.04935 | 23.72557 | 24.02293 |
| Q86V48 | Leucine zip | LUZP1 | Leucine zip | 8.571283 | 0.007122 | 23.08858 | 23.06528 | 23.1069 |
| Q9BWJ5 | Splicing fac | SF3B5 | Splicing fac | 7.608018 | 0.000966 | 24.37915 | 24.32217 | 24.53279 |
| Q9BZH6 | WD repeat-c | WDR11 | WD repeat-c | 6.135737 | 0.001132 | 24.16046 | 24.28277 | 24.35151 |
| Q5JPH6 | Probable gl | EARS2 | Probable gl | 3.720777 | 0.053817 | 25.37139 | 22.53196 | 22.6805 |
| Q9Y3T9 | Nucleolar c | NOC2L | Nucleolar c | 8.868053 | 0.003236 | 23.89597 | 23.57796 | 23.61392 |
| O95232 | Luc7-like pı | LUC7L3 | Luc7-like pı | 3.430501 | 0.000237 | 23.85386 | 23.56442 | 24.12605 |
| O95347 | Structural n | SMC2 | Structural n | 6.794629 | 0.033809 | 22.51994 | 22.42997 | 24.0181 |
| O95248 | Myotubular | SBF1 | Myotubular | 9.645396 | 0.001461 | 23.29698 | 23.36399 | 25.00359 |
| Q8WTT2 | Nucleolar c | NOC3L | Nucleolar c | 10.18856 | 0.004381 | 24.24851 | 24.12652 | 24.24198 |
| P02751 | Fibronectin | FN1 | Fibronectin | 9.153691 | 0.00013 | 24.24822 | 22.88138 | 24.46737 |
| Q9HCD5 | Nuclear rec | NCOA5 | Nuclear rec | 10.45389 | 4.06E-05 | 24.07798 | 23.90205 | 23.94762 |
| P19784 | Casein kina | CSNK2A2 | Casein kina | 9.184038 | 0.02158 | 25.42111 | 23.23077 | 23.6753 |
| O94776 | Metastasis-a | MTA2 | Metastasis-a | 8.995196 | 0.024645 | 22.92419 | 24.41397 | 22.72819 |
| P62857 | 40S riboson | RPS28 | Small ribos | 7.004255 | 0.000526 | 25.43094 | 25.19276 | 25.08058 |
| Q9Y4W6 | AFG3-like p | AFG3L2 | AFG3-like p | 10.86838 | 0.000161 | 23.64878 | 23.81627 | 23.77625 |
| Q9Y3C6 | Peptidyl-pro | PPIL1 | Peptidyl-pro | 10.62309 | 0.007126 | 25.0228 | 24.70055 | 24.72067 |
| O96019 | Actin-like pı | ACTL6A | Actin-like pı | 12.80011 | 3.37E-08 | 24.87666 | 24.81577 | 24.92352 |
| P41252 | Isoleucine-- | IARS | Isoleucine-- | 9.673073 | 0.001236 | 23.22273 | 22.95536 | 23.75052 |
| Q8TAQ2 | SWI/SNF co | SMARCC2 | SWI/SNF co | 9.976213 | 0.002043 | 23.75328 | 23.42073 | 23.50458 |
| Q9Y3B7 | 39S riboson | MRPL11 | Large riboso | 6.505765 | 0.001861 | 24.8551 | 24.87145 | 24.04193 |
| P46940 | Ras GTPase | IQGAP1 | Ras GTPase | 7.830725 | 0.016632 | 22.98216 | 23.46638 | 23.55988 |
| O75367 | Core histon | H2AFY | Core histon | 7.909432 | 0.033955 | 25.45908 | 25.08022 | 24.06538 |
| Q9Y285 | Phenylalan | FARSA | Phenylalan | 9.569276 | 0.002393 | 23.99438 | 21.82544 | 24.14032 |
| P48643 | T-complex p | CCT5 | T-complex p | 10.73212 | 0.000768 | 23.75379 | 23.56546 | 23.65491 |
| P55795 | Heterogene | HNRNPH2 | Heterogene | 10.28949 | 0.026885 | 23.6725 | 23.70641 | 25.6871 |
| Q9UNX4 | WD repeat-c | WDR3 | WD repeat-c | 6.941692 | 0.002208 | 22.99245 | 23.67616 | 23.71272 |
| Q719H9 | BTB/POZ do | KCTD1 | BTB/POZ do | 5.839342 | 0.133097 | 25.65987 | 24.45864 | 25.69633 |
| Q9NQT5 | Exosome co | EXOSC3 | Exosome co | 9.367858 | 0.004537 | 24.01112 | 24.01742 | 24.24837 |
| Q8N163 | Cell cycle a | CCAR2 | Cell cycle a | 7.755222 | 0.024468 | 21.56593 | 21.19226 | 25.21063 |
| P0DP25;P0DP24;P0DP23 | | | Calmodulin | 6.114026 | 0.141159 | 25.15503 | 24.71371 | 25.07716 |
| Q96A26 | Protein FAM | FAM162A | Protein FAM | 10.07939 | 0.003459 | 24.73345 | 24.24605 | 24.5956 |
| P46060 | Ran GTPase | RANGAP1 | Ran GTPase | 7.980324 | 6.65E-05 | 23.49143 | 23.31091 | 23.57693 |
| Q96PU8 | Protein qua | QKI | KH domain- | 8.248273 | 0.004202 | 25.11847 | 24.59611 | 24.95172 |
| P27348 | 14-3-3 prot | YWHAQ | 14-3-3 prot | 6.683088 | 0.003331 | 25.46768 | 25.19693 | 25.34313 |
| Q5T200 | Zinc finger C | ZC3H13 | Zinc finger C | 9.615918 | 0.001911 | 23.11801 | 22.67474 | 23.11445 |
| Q658P3 | Metalloredu | STEAP3 | Metalloredu | 5.07903 | 0.01076 | 24.43702 | 22.16708 | 25.55078 |
| O15504 | Nucleoporin | NUPL2 | Nucleoporin | 11.9522 | 0.000122 | 24.3761 | 24.21475 | 24.33402 |
| Q00341 | Vigilin | HDLBP | Vigilin OS=H | 6.233327 | 0.000648 | 22.8536 | 22.24954 | 24.37908 |
| P40227 | T-complex p | CCT6A | T-complex p | 10.45798 | 0.000909 | 23.6371 | 23.57738 | 23.56279 |
| Q658Y4 | Protein FAM | FAM91A1 | Protein FAM | 10.1247 | 0.00211 | 23.47618 | 23.24339 | 23.5099 |
| P82663 | 28S riboson | MRPS25 | Small ribos | 10.47901 | 0.003356 | 24.19603 | 23.92814 | 24.17912 |
| P82914 | 28S riboson | MRPS15 | Small ribos | 11.54008 | 0.00037 | 24.17341 | 24.0077 | 24.15977 |
| Q8NI36 | WD repeat-c | WDR36 | WD repeat-c | 7.65342 | 0.045188 | 23.8956 | 22.65801 | 22.96019 |

| | | | | | | | | |
|---|---|---|---|---|---|---|---|---|
| P62826 | GTP-bindin | RAN | GTP-bindin | 7.389516 | 0.002271 | 25.33817 | 23.96146 | 23.56999 |
| Q9NX58 | Cell growth | LYAR | Cell growth | 9.238298 | 0.000594 | 23.99084 | 23.7043 | 23.82279 |
| P05023 | Sodium/po | ATP1A1 | Sodium/po | 10.82924 | 0.000216 | 23.22491 | 22.99376 | 23.02174 |
| Q96CB9 | 5-methylcy | NSUN4 | 5-methylcy | 8.392184 | 0.124834 | 24.76137 | 24.62845 | 24.49448 |
| Q01813 | ATP-depen | PFKP | ATP-depen | 8.204659 | 0.008359 | 23.53029 | 23.72067 | 23.53813 |
| Q8IZP0 | Abl interact | ABI1 | Abl interact | 7.547178 | 4.22E-06 | 24.54062 | 24.40185 | 24.60958 |
| Q08170 | Serine/argin | SRSF4 | Serine/argin | 11.5425 | 6.60E-06 | 24.13477 | 23.79867 | 24.02411 |
| Q96QV6;P1 | Histone H2 | HIST1H2AA | Histone H2 | 12.33009 | 0.000403 | 25.28623 | 25.02559 | 25.25075 |
| P35232 | Prohibitin | PHB | Prohibitin 1 | 7.725075 | 0.027582 | 23.56151 | 23.40756 | 23.42291 |
| Q96EE3 | Nucleopori | SEH1L | Nucleopori | 9.12475 | 0.005359 | 24.29418 | 24.09112 | 24.25104 |
| Q9BYJ9 | YTH domair | YTHDF1 | YTH domair | 8.795407 | 0.016637 | 24.18896 | 23.92034 | 24.00958 |
| Q9BRJ6 | Uncharacte | C7orf50 | Uncharacte | 13.05449 | 9.73E-07 | 24.87042 | 24.6746 | 24.76764 |
| P18031 | Tyrosine-pr | PTPN1 | Tyrosine-pr | 10.88721 | 5.20E-05 | 23.58026 | 23.48419 | 23.65513 |
| P63208 | S-phase kin | SKP1 | S-phase kin | 11.09812 | 0.001545 | 24.25954 | 24.03145 | 24.35636 |
| Q9NZM5 | Glioma tum | GLTSCR2 | Ribosome b | 10.78871 | 0.000236 | 24.56145 | 24.45268 | 24.64581 |
| O94966 | Ubiquitin ca | USP19 | Ubiquitin ca | 11.27021 | 1.16E-05 | 24.23264 | 24.04444 | 24.25839 |
| P61964 | WD repeat- | WDR5 | WD repeat- | 4.148806 | 0.019853 | 25.67627 | 22.33653 | 22.41426 |
| Q8IX01 | SURP and C | SUGP2 | SURP and C | 8.650893 | 0.015853 | 23.42278 | 23.33811 | 23.57519 |
| Q13200 | 26S proteas | PSMD2 | 26S proteas | 10.08749 | 0.000748 | 24.2793 | 23.84632 | 24.16008 |
| Q9BQ70 | Transcriptio | TCF25 | Ribosome c | 8.809597 | 0.010005 | 23.88148 | 22.3633 | 23.97255 |
| Q86Y79 | Probable pe | PTRH1 | Peptidyl-tRI | 13.04941 | 9.43E-06 | 24.68929 | 24.95496 | 25.22122 |
| O95900 | Probable tR | TRUB2 | Pseudourid | 7.561799 | 0.004827 | 24.34204 | 22.66075 | 22.64172 |
| Q9H6N6 | Putative un | MYH16 | Putative un | 6.369462 | 4.00E-05 | 22.84073 | 22.738 | 22.94831 |
| O00148;Q1 | ATP-depen | DDX39A;DI | ATP-depen | 7.722573 | 0.006848 | 24.04051 | 23.63256 | 23.8812 |
| Q69YN4 | Protein virili | KIAA1429 | Protein virili | 8.569437 | 0.017534 | 22.67401 | 24.56587 | 22.67653 |
| P52907 | F-actin-cap | CAPZA1 | F-actin-cap | 6.216681 | 0.035179 | 24.46918 | 24.62729 | 24.40237 |
| P50991 | T-complex | CCT4 | T-complex | 7.309812 | 0.005582 | 23.32663 | 23.3347 | 23.57496 |
| Q9UBV2 | Protein sel- | SEL1L | Protein sel- | 9.403461 | 0.008848 | 24.25176 | 23.93733 | 22.46993 |
| Q8WZ19 | BTB/POZ dc | KCTD13 | BTB/POZ dc | 10.62617 | 0.000145 | 23.99723 | 23.60286 | 23.98433 |
| P26641 | Elongation | EEF1G | Elongation | 4.923383 | 0.003589 | 24.37617 | 24.12605 | 24.23652 |
| P24752 | Acetyl-CoA | ACAT1 | Acetyl-CoA | 11.20746 | 4.05E-05 | 23.70599 | 23.57808 | 23.56906 |
| Q9Y2R5 | 28S riboso | MRPS17 | Small ribos | 9.593095 | 0.038579 | 24.66395 | 25.888 | 24.46258 |
| O75494 | Serine/argin | SRSF10 | Serine/argin | 10.26221 | 0.001225 | 24.37756 | 24.00325 | 24.20284 |
| Q14315 | Filamin-C | FLNC | Filamin-C O | 9.284804 | 0.000114 | 23.1124 | 22.7383 | 23.14571 |
| Q9NTJ3 | Structural n | SMC4 | Structural n | 9.077415 | 0.004954 | 21.63887 | 23.43314 | 23.47531 |
| Q13438 | Protein OS- | OS9 | Protein OS- | 12.60467 | 8.49E-06 | 24.70092 | 24.3946 | 24.54404 |
| Q8IWX8 | Calcium ho | CHERP | Calcium ho | 11.27238 | 5.78E-05 | 23.79887 | 23.37862 | 23.37941 |
| Q9H0H5 | Rac GTPase | RACGAP1 | Rac GTPase | 9.446646 | 0.003269 | 21.94135 | 23.73892 | 24.07325 |
| O60783 | 28S riboso | MRPS14 | Small ribos | 11.35783 | 0.001078 | 24.52815 | 24.81157 | 24.77062 |
| Q9BRT6 | Protein LLP | LLPH | Protein LLP | 12.15069 | 0.000112 | 25.10817 | 25.09969 | 25.09595 |
| Q14669 | E3 ubiquitir | TRIP12 | E3 ubiquitir | 6.788412 | 0.044004 | 22.5556 | 23.52217 | 21.29058 |
| Q13885 | Tubulin bet | TUBB2A | Tubulin bet | 10.1974 | 0.013939 | 25.37401 | 24.84891 | 23.36346 |
| P62195 | 26S proteas | PSMC5 | 26S proteas | 9.244399 | 0.016531 | 23.61313 | 23.63887 | 23.6058 |
| Q8NI27 | THO compl | THOC2 | THO compl | 11.31851 | 0.000599 | 23.76729 | 23.36346 | 23.63998 |
| Q96I24 | Far upstrear | FUBP3 | Far upstrear | 7.514179 | 0.049409 | 24.4247 | 24.79328 | 24.52958 |
| O95864 | Fatty acid d | FADS2 | Acyl-CoA 6- | 8.18616 | 0.000872 | 24.45168 | 22.03513 | 24.50415 |
| Q14697 | Neutral alpl | GANAB | Neutral alpl | 8.968376 | 0.014229 | 23.84355 | 23.52862 | 23.58405 |
| Q53GS7 | Nucleopori | GLE1 | mRNA expo | 10.62714 | 2.06E-06 | 23.46164 | 23.37769 | 23.44897 |

| | | | | | | | | |
|---|---|---|---|---|---|---|---|---|
| Q6P158 | Putative ATI | DHX57 | Putative ATI | 7.543293 | 0.010216 | 23.78746 | 23.30452 | 23.76699 |
| P47897 | Glutamine-- | QARS | Glutamine-- | 8.386383 | 0.009994 | 23.16797 | 23.1225 | 23.10468 |
| P28799 | Granulins;A | GRN | Progranulin | 12.93545 | 2.36E-05 | 25.67694 | 25.33906 | 25.73149 |
| Q09028 | Histone-bin | RBBP4 | Histone-bin | 7.284952 | 0.005055 | 21.84659 | 25.35525 | 25.67573 |
| Q8N1F7 | Nuclear por | NUP93 | Nuclear por | 5.955805 | 0.010809 | 22.63657 | 23.71366 | 23.79897 |
| Q49A26 | Putative oxi | GLYR1 | Cytokine-lik | 10.25455 | 0.00497 | 23.4073 | 23.17502 | 23.43632 |
| Q09161 | Nuclear cap | NCBP1 | Nuclear cap | 9.921042 | 0.002897 | 24.16262 | 23.82026 | 24.01759 |
| Q86UE4 | Protein LYRI | MTDH | Protein LYRI | 8.116399 | 0.000915 | 23.58348 | 23.0641 | 23.4179 |
| P08237 | ATP-depend | PFKM | ATP-depend | 9.798073 | 0.000348 | 23.23493 | 23.28743 | 23.50312 |
| Q3ZCQ8 | Mitochondı | TIMM50 | Mitochondı | 9.394094 | 0.017995 | 23.09528 | 22.8632 | 25.0611 |
| P09543 | 2,3-cyclic-n | CNP | 2,3-cyclic-n | 5.609162 | 0.005752 | 21.61027 | 22.96907 | 24.97941 |
| Q9BXS6 | Nucleolar a | NUSAP1 | Nucleolar a | 5.653872 | 0.01565 | 22.73849 | 22.91683 | 23.45161 |
| P51148 | Ras-related | RAB5C | Ras-related | 9.655534 | 0.001601 | 23.95056 | 23.85405 | 23.78676 |
| Q8WWY3 | U4/U6 sma | PRPF31 | U4/U6 sma | 9.641208 | 0.001246 | 23.81812 | 23.43122 | 24.08894 |
| Q15084 | Protein disu | PDIA6 | Protein disu | 10.15805 | 0.000251 | 23.31257 | 23.84345 | 23.70588 |
| P25685 | DnaJ homo | DNAJB1 | DnaJ homo | 10.33095 | 0.023287 | 24.37292 | 24.30159 | 24.3579 |
| Q9Y3D3 | 28S ribosor | MRPS16 | Small ribos | 10.80729 | 3.35E-06 | 24.61801 | 24.42554 | 24.6392 |
| P83111 | Serine beta- | LACTB | Serine beta- | 11.05438 | 7.85E-05 | 23.38416 | 23.0418 | 23.30703 |
| P24844 | Myosin regu | MYL9 | Myosin regu | 6.49809 | 0.003683 | 24.87619 | 24.65633 | 25.14406 |
| Q9NNW5 | WD repeat-c | WDR6 | tRNA (34-2- | 10.72713 | 0.000186 | 23.37358 | 23.16682 | 23.1559 |
| Q86VI3 | Ras GTPase | IQGAP3 | Ras GTPase | 9.302052 | 0.003013 | 22.76099 | 22.78225 | 22.95828 |
| P49591 | Serine--tRN | SARS | Serine--tRN | 8.644161 | 0.007997 | 23.57346 | 23.27242 | 23.67874 |
| Q9NP66 | High mobili | HMG20A | High mobili | 8.625382 | 0.027983 | 22.65207 | 22.77794 | 24.36112 |
| P01024 | Compleme | C3 | Compleme | 5.842711 | 0.013909 | 23.0911 | 25.8892 | 25.71406 |
| Q9BQ67 | Glutamate-r | GRWD1 | Glutamate-r | 10.6446 | 0.000552 | 24.06645 | 23.70746 | 24.80666 |
| P31947 | 14-3-3 prot | SFN | 14-3-3 prot | 4.68224 | 0.250652 | 24.95362 | 21.61297 | 22.176 |
| Q9Y291 | 28S ribosor | MRPS33 | Small ribos | 11.97435 | 2.68E-06 | 24.33477 | 24.21497 | 24.3505 |
| Q9UH73;Q9 | Transcriptio | EBF1;EBF3 | Transcriptio | 10.53283 | 0.000119 | 23.16909 | 23.36546 | 23.38337 |
| Q8N302 | Angiogenic | AGGF1 | Angiogenic | 7.434838 | 0.006204 | 24.62193 | 19.82265 | 21.80377 |
| Q99623 | Prohibitin-2 | PHB2 | Prohibitin-2 | 8.449574 | 0.026038 | 23.87502 | 23.88446 | 23.55321 |
| P43246 | DNA misma | MSH2 | DNA misma | 7.678647 | 0.021731 | 23.78497 | 23.57554 | 24.04827 |
| O75962 | Triple functi | TRIO | Triple functi | 10.08769 | 0.000171 | 23.04332 | 22.8507 | 22.82318 |
| Q9BYD3 | 39S ribosor | MRPL4 | Large ribosc | 11.5384 | 0.000188 | 24.24909 | 24.05953 | 24.17919 |
| Q06210 | Glutamine-- | GFPT1 | Glutamine-- | 10.4985 | 0.000788 | 23.25379 | 23.03098 | 23.27938 |
| O43663 | Protein regu | PRC1 | Protein regu | 3.782284 | 0.018627 | 24.25039 | 24.2781 | 22.3904 |
| Q9UBX3 | Mitochondı | SLC25A10 | Mitochondı | 7.243025 | 0.033481 | 23.57335 | 23.72411 | 23.85633 |
| Q99575 | Ribonuclea | POP1 | Ribonuclea | 11.18055 | 4.95E-08 | 22.98718 | 22.82229 | 22.87316 |
| Q9BRX2 | Protein pelc | PELO | Protein pelc | 11.49673 | 0.000195 | 23.34841 | 23.41054 | 23.53576 |
| Q9Y4P3 | Transducin | TBL2 | Transducin | 10.11212 | 0.040889 | 23.93031 | 25.22582 | 24.03725 |
| P24539 | ATP syntha | ATP5F1 | ATP syntha | 9.834692 | 7.50E-05 | 23.89264 | 23.87699 | 24.05374 |
| Q86XZ4 | Spermatoge | SPATS2 | Spermatoge | 8.913795 | 0.014494 | 23.70852 | 23.8188 | 23.27696 |
| P31151 | Protein S10 | S100A7 | Protein S10 | 0.297216 | 0.91179 | 23.18492 | 20.618 | 20.35767 |
| P06703 | Protein S10 | S100A6 | Protein S10 | 6.111384 | 0.247082 | 24.38561 | 25.73988 | 24.75979 |
| P78364 | Polyhomec | PHC1 | Polyhomec | 11.36874 | 0.000288 | 24.11593 | 23.94262 | 23.94432 |
| Q9Y3E5 | Peptidyl-tRI | PTRH2 | Peptidyl-tRI | 10.56485 | 0.021694 | 24.94833 | 25.04185 | 24.70762 |
| Q5BJF6 | Outer dense | ODF2 | Outer dense | 11.11665 | 5.66E-06 | 23.46339 | 23.52241 | 23.6781 |
| P46063 | ATP-depend | RECQL | ATP-depend | 10.40451 | 1.35E-07 | 23.35797 | 23.13751 | 23.45801 |
| Q15007 | Pre-mRNA-s | WTAP | Pre-mRNA-s | 9.30459 | 0.010926 | 23.49741 | 23.28673 | 23.46837 |

| | | | | | | | | |
|---|---|---|---|---|---|---|---|---|
| Q969S3 | Zinc finger p | ZNF622 | Cytoplasmi | 6.422797 | 0.020919 | 21.65071 | 21.49287 | 24.77992 |
| O60313 | Dynamin-lil | OPA1 | Dynamin-lil | 8.98011 | 0.000865 | 22.4784 | 23.88948 | 22.68964 |
| P19388 | DNA-directe | POLR2E | DNA-directe | 9.939596 | 0.007632 | 23.67906 | 23.53825 | 23.72588 |
| Q9NY61 | Protein AAT | AATF | Protein AAT | 10.64057 | 0.000241 | 23.16711 | 23.13691 | 23.30883 |
| Q9H089 | Large subur | LSG1 | Large subur | 5.63439 | 0.093008 | 23.43479 | 23.20129 | 23.50045 |
| Q9UKS6 | Protein kina | PACSIN3 | Protein kina | 8.333481 | 0.01169 | 24.34645 | 23.90325 | 23.60513 |
| P17812 | CTP syntha | CTPS1 | CTP syntha | 11.17481 | 5.04E-06 | 23.33756 | 23.32924 | 23.50506 |
| Q14562 | ATP-depend | DHX8 | ATP-depend | 10.84437 | 0.000201 | 23.09291 | 22.9735 | 23.15709 |
| Q12788 | Transducin | TBL3 | Transducin | 8.60507 | 0.004622 | 23.15089 | 22.90985 | 23.05641 |
| P53999 | Activated RI | SUB1 | Activated RI | 1.392047 | 0.145001 | 23.22742 | 23.17713 | 23.55438 |
| Q13144 | Translation | EIF2B5 | Translation | 12.30177 | 3.17E-05 | 24.93369 | 25.01167 | 25.15217 |
| Q86VY4 | Testis-speci | TSPYL5 | Testis-speci | 9.827106 | 0.013682 | 25.24242 | 24.68977 | 25.14215 |
| Q8WX93 | Palladin | PALLD | Palladin OS | 10.05616 | 4.13E-05 | 23.23301 | 22.93272 | 23.07253 |
| Q9BSC4 | Nucleolar p | NOL10 | Nucleolar p | 10.82426 | 2.04E-05 | 23.21379 | 23.12819 | 23.30131 |
| P60468 | Protein tran | SEC61B | Protein tran | 9.74115 | 0.011468 | 24.42291 | 24.34394 | 24.52719 |
| O43172 | U4/U6 sma | PRPF4 | U4/U6 sma | 9.699447 | 0.004119 | 24.05705 | 23.03157 | 23.26098 |
| Q9NZT1 | Calmodulir | CALML5 | Calmodulir | 1.091165 | 0.789136 | 19.35171 | 17.23781 | 20.20393 |
| P40939 | Trifunctiona | HADHA | Trifunctiona | 7.805161 | 0.000382 | 23.70314 | 22.86735 | 23.96358 |
| P12004 | Proliferating | PCNA | Proliferating | 8.736897 | 0.005419 | 24.38765 | 23.27597 | 24.3581 |
| Q16718 | NADH dehy | NDUFA5 | NADH dehy | 12.09255 | 4.29E-05 | 24.56034 | 24.61611 | 24.81929 |
| Q9C0J8 | pre-mRNA 3 | WDR33 | pre-mRNA 3 | 9.15129 | 0.000163 | 22.23599 | 23.06314 | 23.38759 |
| P33176 | Kinesin-1 h | KIF5B | Kinesin-1 h | 6.7217 | 0.002081 | 22.66297 | 23.88772 | 22.71452 |
| P02452 | Collagen al | COL1A1 | Collagen al | 9.406274 | 0.003808 | 23.7684 | 24.46389 | 24.48603 |
| Q9H6H4 | Receptor ex | REEP4 | Receptor ex | 12.7781 | 2.40E-07 | 24.86141 | 24.57848 | 24.73268 |
| Q9NSI2 | Protein FAM | FAM207A | Ribosome b | 7.500683 | 0.00738 | 24.11696 | 22.39239 | 22.80033 |
| Q8IX12 | Cell divisior | CCAR1 | Cell divisior | 11.49921 | 0.000113 | 23.76709 | 23.42329 | 23.80686 |
| Q86VM9 | Zinc finger C | ZC3H18 | Zinc finger C | 9.485063 | 0.002335 | 23.32937 | 23.13806 | 22.85144 |
| O75396 | Vesicle-traff | SEC22B | Vesicle-traff | 8.031575 | 0.019426 | 23.79956 | 23.46874 | 23.48247 |
| P48634 | Protein PRR | PRRC2A | Protein PRR | 9.75471 | 9.83E-05 | 23.30605 | 23.13573 | 23.41867 |
| O94766 | Galactosylg | B3GAT3 | Galactosylg | 8.941032 | 0.000516 | 23.86259 | 24.08805 | 24.08278 |
| P35568 | Insulin rece | IRS1 | Insulin rece | 10.17213 | 4.47E-05 | 22.90347 | 22.79921 | 23.05391 |
| P11177 | Pyruvate de | PDHB | Pyruvate de | 10.44429 | 4.02E-06 | 23.59286 | 23.50337 | 23.57242 |
| Q8WUD4 | Coiled-coil | CCDC12 | Coiled-coil | 10.12532 | 0.000104 | 24.1303 | 23.57369 | 24.10526 |
| P53007 | Tricarboxyla | SLC25A1 | Tricarboxyla | 11.23004 | 2.15E-06 | 23.71429 | 23.54416 | 23.78706 |
| Q14152 | Eukaryotic t | EIF3A | Eukaryotic t | 11.96083 | 0.000183 | 23.22366 | 23.39139 | 23.41326 |
| Q02880 | DNA topois | TOP2B | DNA topois | 9.201582 | 0.042834 | 22.62333 | 22.49194 | 24.053 |
| P61962 | DDB1- and | DCAF7 | DDB1- and | 7.84098 | 0.00172 | 23.829 | 23.61336 | 22.62983 |
| Q12873 | Chromodor | CHD3 | Chromodor | 9.278301 | 0.002516 | 23.3048 | 21.06842 | 23.41015 |
| P49815 | Tuberin | TSC2 | Tuberin OS= | 6.733426 | 0.095318 | 22.2055 | 23.72338 | 23.64933 |
| Q9BX40 | Protein LSM | LSM14B | Protein LSM | 11.25087 | 9.76E-06 | 23.68581 | 23.48579 | 23.77686 |
| O43164 | E3 ubiquitir | PJA2 | E3 ubiquitir | 10.40231 | 1.51E-05 | 23.85405 | 23.70324 | 23.73376 |
| Q13595 | Transforme | TRA2A | Transforme | 9.568227 | 0.027896 | 24.93805 | 23.00522 | 23.12958 |
| O95714 | E3 ubiquitir | HERC2 | E3 ubiquitir | 6.94083 | 0.002402 | 23.32937 | 22.5603 | 21.84824 |
| Q8WVB6 | Chromosor | CHTF18 | Chromosor | 5.812448 | 0.041078 | 21.75034 | 22.78173 | 23.51689 |
| P63241 | Eukaryotic t | EIF5A | Eukaryotic t | 7.461783 | 0.047052 | 24.09136 | 24.15885 | 24.11648 |
| Q8N3C0 | Activating si | ASCC3 | Activating si | 7.862555 | 0.04581 | 22.45761 | 23.07009 | 23.11429 |
| Q8NBJ5 | Procollager | COLGALT1 | Procollager | 8.869309 | 0.009969 | 23.58176 | 21.99341 | 23.94851 |
| P39748 | Flap endon | FEN1 | Flap endon | 3.915484 | 0.032821 | 22.24307 | 24.52648 | 22.14833 |

| | | | | | | | | |
|---|---|---|---|---|---|---|---|---|
| Q9UM54 | Unconventi | MYO6 | Unconventi | 5.208214 | 0.007032 | 22.93817 | 21.70291 | 23.41995 |
| P47756 | F-actin-cap | CAPZB | F-actin-cap | 7.06488 | 0.05311 | 24.04277 | 23.12992 | 24.02538 |
| Q8NHU0;Q | Cancer/test | CT45A3;CT | Cancer/test | 9.072809 | 0.001889 | 24.19189 | 23.57565 | 24.14849 |
| O00566 | U3 small nu | MPHOSPH1 | U3 small nu | 12.80211 | 7.79E-06 | 24.26127 | 24.24075 | 24.3064 |
| P00338 | L-lactate de | LDHA | L-lactate de | 2.213456 | 0.340964 | 24.20135 | 21.11674 | 21.26161 |
| O75683 | Surfeit locu | SURF6 | Surfeit locu | 10.55654 | 4.33E-05 | 23.57738 | 23.381 | 23.36105 |
| Q15050 | Ribosome b | RRS1 | Ribosome b | 8.627771 | 1.07E-05 | 22.957 | 23.14936 | 23.19895 |
| Q9Y3C1 | Nucleolar p | NOP16 | Nucleolar p | 12.14564 | 2.63E-05 | 23.50069 | 23.42163 | 23.55789 |
| O43175 | D-3-phosph | PHGDH | D-3-phosph | 7.471389 | 0.007294 | 23.38588 | 23.12799 | 23.005 |
| P06400 | Retinoblast | RB1 | Retinoblast | 8.63647 | 0.000623 | 22.95862 | 23.16308 | 23.8225 |
| P82932 | 28S riboso | MRPS6 | Small ribos | 11.57182 | 4.09E-05 | 24.26757 | 24.15298 | 24.38423 |
| Q9P0L0 | Vesicle-ass | VAPA | Vesicle-ass | 8.149836 | 0.01258 | 23.63101 | 20.84896 | 23.97806 |
| Q5T5U3 | Rho GTPase | ARHGAP21 | Rho GTPase | 10.49567 | 1.00E-06 | 22.72715 | 22.64068 | 22.78337 |
| O94880 | PHD finger | PHF14 | PHD finger | 9.900209 | 0.002171 | 21.5566 | 23.65305 | 23.79858 |
| Q9UG63 | ATP-binding | ABCF2 | ATP-binding | 9.25596 | 0.001787 | 23.34989 | 23.04395 | 23.08541 |
| Q9UQB8 | Brain-specif | BAIAP2 | Brain-specif | 10.07936 | 0.006154 | 23.4957 | 23.37425 | 23.82736 |
| Q9H4H8 | Protein FAM | FAM83D | Protein FAM | 8.748205 | 0.005633 | 24.84316 | 22.88455 | 23.92416 |
| O15231 | Zinc finger p | ZNF185 | Zinc finger p | 7.880119 | 0.010182 | 24.22486 | 22.17621 | 22.35103 |
| Q15024 | Exosome co | EXOSC7 | Exosome co | 10.04034 | 0.001864 | 24.54593 | 24.03187 | 24.10358 |
| Q96ME7 | Zinc finger p | ZNF512 | Zinc finger p | 8.328785 | 0.051015 | 22.99836 | 23.98729 | 22.90296 |
| O43159 | Ribosomal | RRP8 | Ribosomal | 11.79152 | 0.000481 | 24.60772 | 24.60688 | 24.28023 |
| P82912 | 28S riboso | MRPS11 | Small ribos | 11.12595 | 0.000621 | 24.28835 | 24.14973 | 24.37106 |
| Q96QR8 | Transcriptio | PURB | Transcriptio | 7.564685 | 0.037383 | 26.02851 | 22.63103 | 22.74759 |
| Q92620 | Pre-mRNA-s | DHX38 | Pre-mRNA-s | 9.700736 | 0.000854 | 23.07096 | 23.00587 | 23.13865 |
| O75525 | KH domain- | KHDRBS3 | KH domain- | 5.109092 | 5.17E-06 | 24.3057 | 24.05796 | 24.53884 |
| Q9BYG3 | MKI67 FHA | NIFK | MKI67 FHA | 8.591617 | 0.004223 | 23.82357 | 23.24069 | 23.73293 |
| Q9Y4C8 | Probable RI | RBM19 | Probable RI | 8.501668 | 0.016623 | 23.16987 | 23.07256 | 23.2856 |
| Q99661 | Kinesin-like | KIF2C | Kinesin-like | 9.050515 | 0.012661 | 22.35587 | 23.91871 | 24.05408 |
| Q8NFW8 | N-acylneura | CMAS | N-acylneura | 9.312336 | 0.007551 | 24.1876 | 22.32169 | 24.10709 |
| Q92922 | SWI/SNF co | SMARCC1 | SWI/SNF co | 9.038756 | 0.001325 | 23.37039 | 23.34935 | 23.43619 |
| P98179 | RNA-bindin | RBM3 | RNA-bindin | 9.47896 | 0.040028 | 25.21937 | 24.75119 | 24.98481 |
| Q7KZI7 | Serine/thre | MARK2 | Serine/thre | 9.810803 | 0.008741 | 23.89208 | 23.18349 | 23.3806 |
| P29692 | Elongation | EEF1D | Elongation | 6.130554 | 0.001358 | 23.07099 | 23.50616 | 23.0945 |
| Q9H227 | Cytosolic b | GBA3 | Cytosolic b | 12.48804 | 5.88E-05 | 24.60275 | 25.90681 | 25.36333 |
| P82921 | 28S riboso | MRPS21 | Small ribos | 11.92036 | 4.13E-05 | 24.15236 | 24.32526 | 24.24779 |
| O75616 | GTPase Era, | ERAL1 | GTPase Era, | 10.9505 | 5.05E-05 | 23.37928 | 22.99177 | 23.24761 |
| Q8WVM0 | Dimethylad | TFB1M | Dimethylad | 9.762632 | 0.001224 | 23.60241 | 23.168 | 23.32787 |
| Q14258 | E3 ubiquitir | TRIM25 | E3 ubiquitir | 8.341145 | 0.006115 | 23.2439 | 22.47155 | 22.79309 |
| Q13322 | Growth fact | GRB10 | Growth fact | 11.43403 | 4.20E-05 | 23.50264 | 23.59582 | 23.64878 |
| Q9P013 | Spliceosom | CWC15 | Spliceosom | 7.854965 | 0.001018 | 23.34408 | 22.96134 | 24.68817 |
| P78362 | SRSF protei | SRPK2 | SRSF protei | 9.71962 | 0.004802 | 23.20177 | 21.14327 | 23.23681 |
| Q9GZS1 | DNA-directe | POLR1E | DNA-directe | 10.37995 | 0.000591 | 23.41351 | 23.19492 | 23.4665 |
| Q5RKV6 | Exosome co | EXOSC6 | Exosome co | 10.09842 | 0.013956 | 24.2438 | 23.96869 | 23.93211 |
| P49458 | Signal recog | SRP9 | Signal recog | 8.109642 | 0.006612 | 25.20527 | 24.55022 | 24.41125 |
| Q13751 | Laminin sul | LAMB3 | Laminin sul | 10.12126 | 7.98E-05 | 22.51696 | 22.36568 | 22.59099 |
| Q9UGN5 | Poly [ADP-r | PARP2 | Poly [ADP-r | 9.209489 | 1.28E-05 | 23.47877 | 23.6953 | 23.40133 |
| P82664 | 28S riboso | MRPS10 | Small ribos | 11.5391 | 0.000355 | 24.2764 | 24.30946 | 24.29985 |
| Q9NUL7 | Probable AT | DDX28 | Probable AT | 9.249054 | 0.00022 | 22.98402 | 22.73132 | 22.73444 |

| | | | | | | | | |
|---|---|---|---|---|---|---|---|---|
| Q13418 | Integrin-link | ILK | Integrin-link | 10.28723 | 0.000218 | 22.99517 | 23.06019 | 23.02553 |
| O94851 | Protein-met | MICAL2 | [F-actin]-mo | 10.33552 | 6.36E-06 | 22.92883 | 22.54593 | 22.85641 |
| P62136 | Serine/threo | PPP1CA | Serine/threo | 9.589416 | 0.030747 | 23.89143 | 25.31752 | 23.66861 |
| Q9H0U4;Q9 | Ras-related | RAB1B;RAB | Ras-related | 9.081979 | 0.01949 | 24.9198 | 24.81431 | 25.0056 |
| Q9H6T3 | RNA polym | RPAP3 | RNA polym | 1.946873 | 0.021778 | 22.71477 | 22.59259 | 24.19865 |
| O15371 | Eukaryotic t | EIF3D | Eukaryotic t | 9.46472 | 0.020471 | 22.66566 | 24.66438 | 22.77286 |
| Q6DKI1 | 60S riboson | RPL7L1 | Ribosomal | 11.5212 | 8.36E-07 | 23.59628 | 23.7213 | 23.72578 |
| Q99523 | Sortilin | SORT1 | Sortilin OS= | 11.27492 | 1.26E-07 | 23.72099 | 23.52779 | 23.98807 |
| P51571 | Translocon | SSR4 | Translocon | 9.681181 | 0.013036 | 24.31997 | 24.15661 | 22.53141 |
| Q01469 | Fatty acid-b | FABP5 | Fatty acid-b | 0.880146 | 0.764461 | 24.25803 | 21.23818 | 20.2764 |
| Q8WYA6 | Beta-catenin | CTNNBL1 | Beta-catenin | 10.34295 | 0.000303 | 23.61358 | 23.52719 | 23.56023 |
| Q9Y221 | 60S riboson | NIP7 | 60S riboson | 11.44709 | 9.84E-06 | 24.37019 | 23.76739 | 24.01402 |
| O60568 | Procollagen | PLOD3 | Multifunctio | 5.644885 | 0.089506 | 22.85439 | 22.56141 | 23.58451 |
| O43896 | Kinesin-like | KIF1C | Kinesin-like | 7.480975 | 0.00034 | 22.90997 | 22.88813 | 22.58518 |
| Q13547 | Histone dea | HDAC1 | Histone dea | 10.01024 | 9.39E-05 | 23.85158 | 23.71701 | 23.80164 |
| Q8WXF1 | Paraspeckle | PSPC1 | Paraspeckle | 5.919027 | 0.099594 | 24.89643 | 22.48478 | 22.71314 |
| Q16637 | Survival mo | SMN1 | Survival mo | 11.06531 | 3.67E-07 | 23.37769 | 23.33497 | 23.35662 |
| Q8TEQ6 | Gem-assoc | GEMIN5 | Gem-assoc | 9.282248 | 0.001177 | 23.41506 | 23.37305 | 23.3769 |
| Q9UGJ1 | Gamma-tub | TUBGCP4 | Gamma-tub | 7.933688 | 0.023983 | 22.50232 | 23.58692 | 23.75511 |
| O43670 | BUB3-intera | ZNF207 | BUB3-intera | 6.544586 | 0.004733 | 21.43517 | 24.22942 | 24.82731 |
| Q9UBS4 | DnaJ homo | DNAJB11 | DnaJ homo | 8.564466 | 0.00163 | 23.10716 | 23.14911 | 23.48996 |
| Q99547 | M-phase ph | MPHOSPH6 | M-phase ph | 9.604301 | 0.00011 | 23.80104 | 23.42547 | 23.71156 |
| O15226 | NF-kappa-B | NKRF | NF-kappa-B | 8.050986 | 0.003007 | 23.03475 | 22.41398 | 22.62156 |
| Q9NP72 | Ras-related | RAB18 | Ras-related | 9.050338 | 0.00257 | 23.76051 | 23.74468 | 23.43505 |
| Q9Y5Q9 | General tran | GTF3C3 | General tran | 8.433688 | 0.003468 | 22.83644 | 22.53974 | 22.77398 |
| Q9BV38 | WD repeat-c | WDR18 | WD repeat-c | 8.39224 | 0.013958 | 24.00513 | 21.78804 | 21.90551 |
| O94763 | Unconventi | URI1 | Unconventi | 11.77556 | 2.65E-07 | 23.22297 | 22.93659 | 23.22258 |
| P42696 | RNA-bindin | RBM34 | RNA-bindin | 7.587286 | 0.010796 | 23.49069 | 23.20332 | 20.81995 |
| Q8IY37 | Probable AT | DHX37 | Probable AT | 8.188301 | 0.003088 | 22.77864 | 22.72594 | 22.87252 |
| P04181 | Ornithine ar | OAT | Ornithine ar | 7.306731 | 7.58E-07 | 23.40665 | 22.96118 | 23.24329 |
| P43487 | Ran-specific | RANBP1 | Ran-specific | 8.608916 | 0.002182 | 23.25969 | 23.66308 | 23.78706 |
| Q9Y3B4 | Splicing fac | SF3B6 | Splicing fac | 9.941984 | 0.025967 | 24.7304 | 24.40925 | 24.84695 |
| P39656 | Dolichyl-dip | DDOST | Dolichyl-dip | 9.226241 | 0.025336 | 23.5956 | 23.93076 | 23.97937 |
| P07237 | Protein disu | P4HB | Protein disu | 5.911473 | 0.003797 | 23.30619 | 22.54184 | 22.76935 |
| Q6NUJ5 | PWWP dom | PWWP2B | PWWP dom | 8.35285 | 0.022283 | 23.42201 | 21.8428 | 22.94244 |
| P17980 | 26S proteas | PSMC3 | 26S proteas | 8.776052 | 0.011583 | 23.19795 | 22.95015 | 23.32869 |
| O75531 | Barrier-to-au | BANF1 | Barrier-to-au | 5.759344 | 0.075102 | 24.45563 | 23.50191 | 24.21941 |
| P54136 | Arginine--tR | RARS | Arginine--tR | 8.701652 | 0.000867 | 23.21855 | 22.82009 | 23.07821 |
| Q14139 | Ubiquitin co | UBE4A | Ubiquitin co | 9.782076 | 0.000237 | 22.98936 | 22.86756 | 23.08508 |
| Q9NVV4 | Poly(A) RNA | MTPAP | Poly(A) RNA | 9.70048 | 0.001962 | 23.69774 | 23.38759 | 23.77997 |
| Q9H6F5 | Coiled-coil | CCDC86 | Coiled-coil | 9.50943 | 0.004519 | 23.73138 | 22.69304 | 23.07946 |
| O75369 | Filamin-B | FLNB | Filamin-B O | 3.563021 | 0.095083 | 23.4946 | 21.68192 | 21.58394 |
| P30837 | Aldehyde d | ALDH1B1 | Aldehyde d | 7.857864 | 0.0085 | 21.7737 | 23.78676 | 21.73372 |
| Q86U42 | Polyadenyl | PABPN1 | Polyadenyl | 10.75725 | 1.99E-05 | 24.0818 | 23.9411 | 23.92886 |
| P63151 | Serine/threo | PPP2R2A | Serine/threo | 8.474394 | 0.014161 | 23.2967 | 22.94951 | 23.07279 |
| Q9H4L5 | Oxysterol-b | OSBPL3 | Oxysterol-b | 10.45314 | 8.07E-05 | 22.59423 | 22.38182 | 22.70744 |
| Q9Y5X2 | Sorting nexi | SNX8 | Sorting nexi | 7.422711 | 0.025346 | 23.61639 | 22.07473 | 22.08132 |
| Q9NQ55 | Suppressor | PPAN | Suppressor | 9.805251 | 2.92E-07 | 23.04475 | 22.90705 | 23.00108 |

| | | | | | | | | |
|---|---|---|---|---|---|---|---|---|
| O15235 | 28S riboso | MRPS12 | Small ribos | 8.570186 | 0.015134 | 23.81431 | 23.63345 | 23.78856 |
| Q8IUD2 | ELKS/Rab6- | ERC1 | ELKS/Rab6- | 3.924685 | 0.002264 | 23.3434 | 21.53203 | 22.0424 |
| P41743 | Protein kina | PRKCI | Protein kina | 11.30206 | 0.000163 | 23.74981 | 23.71722 | 23.87296 |
| P06493 | Cyclin-depe | CDK1 | Cyclin-depe | 8.968635 | 0.020393 | 24.03221 | 22.11082 | 22.32177 |
| P11413 | Glucose-6-| | G6PD | Glucose-6-| | 10.45042 | 0.000126 | 22.99091 | 22.92091 | 22.85673 |
| Q96MX6 | WD repeat-c | WDR92 | Dynein axon | 5.539362 | 0.008272 | 22.01308 | 20.89815 | 24.70804 |
| Q15366 | Poly(rC)-bir | PCBP2 | Poly(rC)-bir | 9.4213 | 0.0063 | 24.21215 | 23.94566 | 24.38706 |
| P35240 | Merlin | NF2 | Merlin OS=I | 4.05002 | 0.002717 | 23.15179 | 23.96093 | 23.1102 |
| O75947 | ATP synthas | ATP5H | ATP synthas | 11.81377 | 0.000106 | 23.77746 | 23.74366 | 23.91351 |
| P56134 | ATP synthas | ATP5J2 | ATP synthas | 12.19084 | 8.67E-05 | 24.80391 | 24.60043 | 24.9371 |
| P78559 | Microtubule | MAP1A | Microtubule | 8.760647 | 0.007942 | 23.45111 | 21.82958 | 23.57865 |
| Q5VWQ0 | Round sper | RSBN1 | Lysine-spec | 9.90357 | 1.59E-05 | 23.1588 | 22.89828 | 23.18259 |
| O15260 | Surfeit locu | SURF4 | Surfeit locu | 10.13492 | 0.029204 | 25.46529 | 23.65535 | 23.7417 |
| Q15424 | Scaffold att | SAFB | Scaffold att | 10.74079 | 0.00042 | 23.23216 | 22.9225 | 23.12373 |
| Q9Y3Z3 | Deoxynucle | SAMHD1 | Deoxynucle | 8.583958 | 0.020842 | 22.48207 | 24.02158 | 22.44622 |
| Q9P2I0 | Cleavage ar | CPSF2 | Cleavage ar | 10.86571 | 2.58E-05 | 22.83761 | 22.74446 | 22.92649 |
| Q9UKB1;Q9 | F-box/WD re | FBXW11;BT | F-box/WD re | 2.396522 | 0.010866 | 23.96825 | 22.41516 | 22.43041 |
| Q969G3 | SWI/SNF-re | SMARCE1 | SWI/SNF-re | 11.28854 | 3.30E-05 | 23.18723 | 23.04352 | 23.34177 |
| Q12904 | Aminoacyl t | AIMP1 | Aminoacyl t | 11.53822 | 0.000123 | 23.99654 | 23.97727 | 23.64251 |
| Q9UKL3 | CASP8-asso | CASP8AP2 | CASP8-asso | 8.702913 | 0.014111 | 22.78955 | 21.46981 | 23.16131 |
| P15170 | Eukaryotic p | GSPT1 | Eukaryotic p | 1.048546 | 0.437382 | 24.40067 | 23.15791 | 22.62854 |
| O00541 | Pescadillo h | PES1 | Pescadillo h | 6.612845 | 0.012919 | 23.19022 | 22.94976 | 23.14465 |
| Q9ULW0 | Targeting pr | TPX2 | Targeting pr | 8.411944 | 0.006864 | 22.71364 | 21.00402 | 23.21435 |
| Q9BSJ8 | Extended sy | ESYT1 | Extended sy | 8.630154 | 4.35E-05 | 22.55063 | 22.42726 | 22.55506 |
| Q8N9Q2 | Protein SRE | SREK1IP1 | Protein SRE | 0.796392 | 0.064028 | 23.85918 | 23.32457 | 23.28065 |
| P62258 | 14-3-3 prot | YWHAE | 14-3-3 prot | 5.254179 | 0.084859 | 23.60388 | 22.7058 | 23.1453 |
| O43395 | U4/U6 sma | PRPF3 | U4/U6 sma | 9.0659 | 0.00469 | 22.97165 | 22.92991 | 23.72046 |
| Q14194 | Dihydropyri | CRMP1 | Dihydropyri | 9.274884 | 0.008588 | 22.76551 | 22.69797 | 22.89013 |
| P14373 | Zinc finger p | TRIM27 | Zinc finger p | 10.31589 | 0.012156 | 24.95651 | 23.06787 | 24.97136 |
| Q969R5 | Lethal(3)ma | L3MBTL2 | Lethal(3)ma | 7.918356 | 0.025928 | 22.46286 | 22.19087 | 23.95544 |
| Q9BPZ7 | Target of rap | MAPKAP1 | Target of rap | 11.00497 | 0.000816 | 23.10327 | 23.29418 | 23.16608 |
| P12814 | Alpha-actin | ACTN1 | Alpha-actin | 2.782215 | 0.21649 | 23.60354 | 20.17231 | 20.50019 |
| Q8NEY8 | Periphilin-1 | PPHLN1 | Periphilin-1 | 11.27121 | 4.50E-05 | 23.47605 | 23.44607 | 23.54581 |
| Q9NW64 | Pre-mRNA-s | RBM22 | Pre-mRNA-s | 10.11245 | 0.0011 | 23.02572 | 22.76943 | 22.99988 |
| P53985 | Monocarbo | SLC16A1 | Monocarbo | 10.39081 | 0.000619 | 23.30048 | 23.20973 | 23.41584 |
| Q14566 | DNA replica | MCM6 | DNA replica | 9.735214 | 2.60E-06 | 23.15023 | 23.01842 | 23.10823 |
| Q9P2K5 | Myelin expr | MYEF2 | Myelin expr | 10.20645 | 0.000842 | 23.50543 | 23.72359 | 23.35703 |
| Q15427 | Splicing fac | SF3B4 | Splicing fac | 4.300529 | 0.00014 | 23.79303 | 23.77826 | 24.03154 |
| O00178 | GTP-binding | GTPBP1 | GTP-binding | 8.841247 | 0.007055 | 23.30675 | 22.61374 | 23.42586 |
| P63220 | 40S riboso | RPS21 | Small ribos | 9.636065 | 0.026483 | 23.00758 | 24.02073 | 23.99886 |
| Q8NDX5 | Polyhomec | PHC3 | Polyhomec | 10.24474 | 1.12E-06 | 23.50579 | 23.4443 | 23.59764 |
| Q9NWT1 | p21-activat | PAK1IP1 | p21-activat | 11.12809 | 2.50E-06 | 24.10789 | 23.70725 | 23.96358 |
| Q9UKF6 | Cleavage ar | CPSF3 | Cleavage ar | 10.61356 | 0.000285 | 23.55742 | 23.55532 | 23.65916 |
| P36406 | E3 ubiquitir | TRIM23 | E3 ubiquitir | 9.291173 | 0.036322 | 25.03808 | 24.30006 | 25.09184 |
| Q14694 | Ubiquitin ca | USP10 | Ubiquitin ca | 9.399557 | 3.22E-06 | 22.82184 | 22.5904 | 22.78955 |
| P18583 | Protein SON | SON | Protein SON | 10.49305 | 1.34E-05 | 23.60479 | 23.37888 | 23.47717 |
| Q9NX20 | 39S riboso | MRPL16 | Large ribos | 6.048985 | 0.056605 | 23.44847 | 22.88582 | 23.39139 |
| Q13555 | Calcium/ca | CAMK2G | Calcium/ca | 10.18446 | 5.13E-05 | 23.28093 | 23.26528 | 23.25969 |

| | | | | | | | | |
|---|---|---|---|---|---|---|---|---|
| Q92610 | Zinc finger p | ZNF592 | Zinc finger p | 4.671144 | 0.071322 | 22.27555 | 22.17045 | 24.14118 |
| Q9H7B2 | Ribosome p | RPF2 | Ribosome p | 9.981249 | 0.000556 | 23.34394 | 23.19864 | 23.4675 |
| Q96A33 | Coiled-coil | CCDC47 | PAT comple | 9.709073 | 0.000879 | 22.92283 | 22.91568 | 22.99421 |
| O14980 | Exportin-1 | XPO1 | Exportin-1 ( | 8.941249 | 0.011316 | 22.98541 | 22.79487 | 22.90058 |
| Q13045 | Protein fligh | FLII | Protein fligh | 7.753334 | 0.000196 | 22.61544 | 22.22497 | 22.8203 |
| Q02413 | Desmoglein | DSG1 | Desmoglein | 3.16159 | 0.231963 | 23.86155 | 21.07862 | 22.33906 |
| Q9UHV9 | Prefoldin su | PFDN2 | Prefoldin su | 11.33012 | 9.92E-05 | 23.56848 | 23.3644 | 23.48481 |
| Q14683 | Structural m | SMC1A | Structural m | 8.480953 | 0.001104 | 22.85398 | 22.77211 | 20.90426 |
| Q8NHQ9 | ATP-depend | DDX55 | ATP-depend | 10.03814 | 4.52E-05 | 23.60286 | 23.28292 | 23.45801 |
| Q9Y4B5 | Microtubule | MTCL1 | Microtubule | 8.441087 | 0.011534 | 23.22255 | 21.75193 | 23.17942 |
| P17655 | Calpain-2 c | CAPN2 | Calpain-2 c | 8.313328 | 0.014102 | 22.82139 | 21.60779 | 23.43721 |
| Q96I25 | Splicing fac | RBM17 | Splicing fac | 9.609277 | 0.000863 | 23.50227 | 23.36906 | 23.42175 |
| O60884 | DnaJ homo | DNAJA2 | DnaJ homo | 9.706572 | 0.000232 | 23.13513 | 22.88902 | 23.2548 |
| Q14204 | Cytoplasmi | DYNC1H1 | Cytoplasmi | 9.777198 | 0.00064 | 22.29182 | 22.11642 | 22.29036 |
| Q13043 | Serine/threo | STK4 | Serine/threo | 10.51805 | 0.000311 | 23.23959 | 23.36533 | 23.51989 |
| Q9Y3A4 | Ribosomal | RRP7A | Ribosomal | 9.919116 | 7.34E-06 | 23.45726 | 23.34272 | 23.74016 |
| P05546 | Heparin cof | SERPIND1 | Heparin cof | 7.405867 | 2.52E-06 | 24.05068 | 23.56023 | 23.56616 |
| Q9Y3B2 | Exosome co | EXOSC1 | Exosome co | 10.2072 | 5.52E-07 | 23.88558 | 23.82221 | 23.90766 |
| Q12769 | Nuclear por | NUP160 | Nuclear por | 8.514005 | 0.00124 | 22.8253 | 22.70626 | 23.17225 |
| Q9Y4W2 | Ribosomal | LAS1L | Ribosomal | 10.04176 | 1.72E-06 | 23.25153 | 22.86486 | 23.06408 |
| Q9H857 | 5-nucleotid | NT5DC2 | 5-nucleotid | 9.778452 | 0.007634 | 23.53552 | 23.40574 | 23.30201 |
| Q92925 | SWI/SNF-re | SMARCD2 | SWI/SNF-re | 10.65863 | 4.90E-05 | 23.27099 | 23.31437 | 23.34286 |
| P24928 | DNA-directe | POLR2A | DNA-directe | 3.295647 | 0.005618 | 23.66047 | 22.03187 | 23.93463 |
| Q12792 | Twinfilin-1 | TWF1 | Twinfilin-1 ( | 7.240829 | 0.115359 | 23.30103 | 23.17353 | 23.29922 |
| Q3KQU3 | MAP7 doma | MAP7D1 | MAP7 doma | 10.0966 | 0.000115 | 23.01511 | 22.93335 | 23.10831 |
| P21127;Q9 | Cyclin-depe | CDK11B;CI | Cyclin-depe | 4.130133 | 0.00077 | 24.09764 | 23.4206 | 23.06062 |
| P27482 | Calmodulin | CALML3 | Calmodulin | 0.251168 | 0.927496 | 24.22979 | 18.41481 | 18.26188 |
| P17480 | Nucleolar tr | UBTF | Nucleolar tr | 8.178509 | 0.013796 | 21.82714 | 23.50531 | 21.80515 |
| P50402 | Emerin | EMD | Emerin OS= | 9.301038 | 0.006096 | 23.92425 | 23.44884 | 23.96226 |
| P62333 | 26S proteas | PSMC6 | 26S proteas | 9.562313 | 0.000267 | 22.78642 | 22.97926 | 22.98508 |
| P52815 | 39S riboson | MRPL12 | Large ribosc | 11.17777 | 1.46E-06 | 23.96869 | 23.5798 | 24.01784 |
| Q7Z417 | Nuclear frag | NUFIP2 | FMR1-intera | 8.554372 | 0.008455 | 23.21849 | 22.46824 | 23.17597 |
| Q16698 | 2,4-dienoyl | DECR1 | 2,4-dienoyl | 7.883569 | 0.023446 | 22.97235 | 21.6574 | 23.30952 |
| Q7RTV0 | PHD finger-l | PHF5A | PHD finger-l | 5.716668 | 0.00014 | 24.28933 | 23.97692 | 24.59252 |
| Q9NXC5 | WD repeat-( | MIOS | GATOR2 co | 8.412267 | 0.002554 | 21.28647 | 22.95562 | 23.15814 |
| P53680 | AP-2 compl | AP2S1 | AP-2 compl | 9.585607 | 0.01071 | 23.80971 | 23.51099 | 23.69721 |
| P46939 | Utrophin | UTRN | Utrophin OS | 8.042137 | 0.000893 | 22.17014 | 22.13195 | 22.27105 |
| Q8WWM7 | Ataxin-2-like | ATXN2L | Ataxin-2-like | 10.9031 | 0.000678 | 22.69852 | 22.5736 | 22.65823 |
| Q9UJF2 | Ras GTPase | RASAL2 | Ras GTPase | 10.28004 | 2.54E-06 | 23.21583 | 23.02901 | 23.09462 |
| P35226;P3 | Polycomb ( | BMI1;PCGF | Polycomb ( | 12.03175 | 9.33E-05 | 23.66514 | 23.87923 | 24.0411 |
| Q96ST3 | Paired ampl | SIN3A | Paired ampl | 10.40118 | 2.31E-06 | 22.796 | 22.66757 | 22.7043 |
| Q9BTC0 | Death-indu | DIDO1 | Death-indu | 5.290005 | 0.000852 | 22.16186 | 23.13325 | 21.10374 |
| Q12874 | Splicing fac | SF3A3 | Splicing fac | 3.527869 | 4.54E-05 | 23.08809 | 22.71944 | 22.85278 |
| P28331 | NADH-ubiq | NDUFS1 | NADH-ubiq | 6.912384 | 0.008626 | 21.90437 | 21.13715 | 23.35299 |
| P61981 | 14-3-3 prot | YWHAG | 14-3-3 prot | 8.091122 | 0.018921 | 23.27498 | 22.61077 | 23.00199 |
| Q9Y5A9 | YTH domain | YTHDF2 | YTH domain | 9.112696 | 0.038291 | 22.99948 | 22.76624 | 24.21682 |
| Q9UH99 | SUN domai | SUN2 | SUN domai | 8.538739 | 0.003559 | 23.19448 | 23.20033 | 23.30591 |
| P42167 | Lamina-ass | TMPO | Lamina-ass | 8.540972 | 0.001357 | 24.0436 | 23.68475 | 23.74273 |

| | | | | | | | | |
|---|---|---|---|---|---|---|---|---|
| O60287 | Nucleolar p | URB1 | Nucleolar p | 10.96189 | 5.02E-06 | 23.00417 | 22.96564 | 22.98083 |
| Q15269 | Periodic tryp | PWP2 | Periodic tryp | 7.770968 | 0.018125 | 22.42846 | 22.33634 | 24.22052 |
| P78346 | Ribonuclea | RPP30 | Ribonuclea | 9.831866 | 0.04012 | 23.74643 | 23.43466 | 23.09636 |
| Q15417 | Calponin-3 | CNN3 | Calponin-3 | 9.435 | 0.020753 | 22.80654 | 22.97456 | 23.17962 |
| Q9BWM7 | Sideroflexin | SFXN3 | Sideroflexin | 9.635303 | 0.00033 | 23.00421 | 22.7269 | 22.93434 |
| Q06787 | Fragile X me | FMR1 | Fragile X me | 11.50734 | 9.49E-05 | 23.72088 | 23.46214 | 23.78077 |
| O60825 | 6-phospho | PFKFB2 | 6-phospho | 7.035021 | 5.32E-05 | 22.34519 | 22.26224 | 22.50307 |
| Q96R06 | Sperm-asso | SPAG5 | Sperm-asso | 11.43206 | 7.26E-05 | 23.53517 | 23.39585 | 23.48358 |
| Q96HP0 | Dedicator o | DOCK6 | Dedicator o | 4.413581 | 0.006779 | 22.03053 | 23.16487 | 20.56272 |
| P62314 | Small nucle | SNRPD1 | Small nucle | 5.63669 | 0.150011 | 24.54881 | 24.50839 | 24.86382 |
| O75694 | Nuclear por | NUP155 | Nuclear por | 5.29782 | 0.049101 | 23.86835 | 21.33478 | 22.04973 |
| O94761 | ATP-depend | RECQL4 | ATP-depend | 8.294777 | 0.036212 | 22.89999 | 22.64665 | 22.93873 |
| P17706 | Tyrosine-pro | PTPN2 | Tyrosine-pro | 5.220072 | 0.172715 | 23.42201 | 23.30591 | 22.99231 |
| P17026 | Zinc finger p | ZNF22 | Zinc finger p | 9.659725 | 0.010026 | 22.85085 | 24.29929 | 24.40113 |
| Q8IUF8 | Bifunctiona | MINA | Ribosomal | 8.262968 | 0.012991 | 21.90437 | 21.6759 | 23.87774 |
| Q14978 | Nucleolar a | NOLC1 | Nucleolar a | 2.927012 | 0.048091 | 22.61365 | 22.92575 | 23.16156 |
| Q68CP9 | AT-rich inter | ARID2 | AT-rich inter | 9.648527 | 0.001585 | 22.81047 | 22.37883 | 22.7071 |
| O60508 | Pre-mRNA-p | CDC40 | Pre-mRNA-p | 6.649261 | 0.058261 | 23.27881 | 22.96686 | 24.14476 |
| Q9P2N5 | RNA-bindin | RBM27 | RNA-bindin | 7.192719 | 0.006459 | 22.80158 | 22.6428 | 23.38811 |
| Q9Y3B9 | RRP15-like | RRP15 | RRP15-like | 11.3459 | 8.41E-06 | 24.04368 | 23.87033 | 23.9453 |
| Q06265 | Exosome co | EXOSC9 | Exosome co | 11.99279 | 2.83E-06 | 24.06932 | 23.90848 | 23.99792 |
| P41208 | Centrin-2 | CETN2 | Centrin-2 O | 11.1144 | 0.000568 | 23.18677 | 23.15533 | 23.36773 |
| P23921 | Ribonucleo | RRM1 | Ribonucleo | 7.922894 | 0.031066 | 22.49802 | 23.61964 | 22.30536 |
| P06730 | Eukaryotic t | EIF4E | Eukaryotic t | 9.845131 | 0.008434 | 23.70388 | 23.51617 | 23.61583 |
| Q5HYW2 | NHS-like pro | NHSL2 | NHS-like pro | 7.528241 | 0.049008 | 22.89717 | 22.30772 | 23.84699 |
| Q92878 | DNA repair p | RAD50 | DNA repair p | 7.900885 | 0.001958 | 22.79231 | 22.49441 | 22.71053 |
| Q96RT7 | Gamma-tub | TUBGCP6 | Gamma-tub | 10.59772 | 0.000489 | 23.32169 | 23.03745 | 23.25129 |
| Q9BRR8 | G patch dor | GPATCH1 | G patch dor | 11.00709 | 3.73E-05 | 23.12177 | 23.15007 | 23.32251 |
| Q13185 | Chromobox | CBX3 | Chromobox | 6.635757 | 0.114615 | 23.17708 | 23.04652 | 24.28299 |
| Q96PZ2 | Protein FAM | FAM111A | Serine prote | 5.139396 | 0.0507 | 23.33538 | 22.71381 | 23.05786 |
| O95299 | NADH dehy | NDUFA10 | NADH dehy | 10.45327 | 5.83E-06 | 24.02149 | 24.03179 | 24.39034 |
| P61106 | Ras-related | RAB14 | Ras-related | 10.69546 | 0.000392 | 23.56151 | 23.34908 | 23.52528 |
| O00116 | Alkyldihydr | AGPS | Alkyldihydr | 11.91713 | 5.70E-07 | 23.80282 | 23.56883 | 23.70345 |
| P06733 | Alpha-enola | ENO1 | Alpha-enola | 6.858191 | 0.14767 | 23.51244 | 22.34819 | 21.41044 |
| O43852 | Calumenin | CALU | Calumenin | 10.147 | 0.000246 | 23.05217 | 23.16734 | 23.34204 |
| Q9NXF1 | Testis-expre | TEX10 | Testis-expre | 9.926868 | 0.000546 | 23.36813 | 23.37862 | 23.44101 |
| Q9NWS0 | PIH1 doma | PIH1D1 | PIH1 doma | 7.484543 | 0.023147 | 22.16423 | 23.72057 | 22.24048 |
| Q16527 | Cysteine an | CSRP2 | Cysteine an | 6.36539 | 0.062289 | 23.55742 | 23.2498 | 23.50154 |
| O15160 | DNA-directe | POLR1C | DNA-directe | 9.176162 | 5.92E-07 | 22.74505 | 22.61923 | 22.78195 |
| O60732 | Melanoma- | MAGEC1 | Melanoma- | 7.31193 | 0.036412 | 23.39034 | 23.28729 | 23.39963 |
| O75116 | Rho-associa | ROCK2 | Rho-associa | 9.872602 | 1.37E-06 | 22.90413 | 22.56456 | 22.7996 |
| Q9BWT7 | Caspase rec | CARD10 | Caspase rec | 4.972027 | 0.060571 | 21.6219 | 21.17414 | 23.57658 |
| P04075 | Fructose-bis | ALDOA | Fructose-bis | 2.881216 | 0.436177 | 23.45576 | 18.1834 | 19.21634 |
| P52298 | Nuclear cap | NCBP2 | Nuclear cap | 11.07943 | 6.76E-05 | 23.88008 | 23.77122 | 23.61538 |
| Q99832 | T-complex p | CCT7 | T-complex p | 7.395329 | 0.004106 | 22.41491 | 22.07179 | 23.21328 |
| Q52LJ0 | Protein FAM | FAM98B | Protein FAM | 8.280696 | 0.003517 | 22.94061 | 22.72061 | 23.00779 |
| P32119 | Peroxiredox | PRDX2 | Peroxiredox | 1.205282 | 0.447268 | 22.2894 | 21.6718 | 21.80896 |
| Q96Q07 | BTB/POZ do | BTBD9 | BTB/POZ do | 9.310657 | 0.011257 | 23.18091 | 21.48168 | 23.85633 |

| | | | | | | | | |
|---|---|---|---|---|---|---|---|---|
| Q15637 | Splicing fac | SF1 | Splicing fac | 9.367243 | 0.00014 | 23.29978 | 23.26098 | 23.13287 |
| Q9BZG1 | Ras-related | RAB34 | Ras-related | 11.86082 | 1.59E-05 | 23.44594 | 23.35931 | 23.3931 |
| O75127 | Pentatricop | PTCD1 | Pentatricop | 9.746563 | 0.000107 | 22.73715 | 22.45879 | 22.5695 |
| O94915 | Protein fury | FRYL | Protein fury | 8.147867 | 0.001093 | 22.27977 | 21.99421 | 22.27002 |
| Q8N1G0 | Zinc finger p | ZNF687 | Zinc finger p | 8.249212 | 0.015456 | 23.51917 | 21.38137 | 21.44076 |
| Q9BQ75 | Protein CMS | CMSS1 | Protein CMS | 9.281089 | 0.036611 | 24.69864 | 22.93224 | 22.99905 |
| P09496 | Clathrin ligh | CLTA | Clathrin ligh | 5.866007 | 8.60E-05 | 22.57549 | 23.13028 | 23.54274 |
| Q8ND30 | Liprin-beta-: | PPFIBP2 | Liprin-beta-: | 3.501684 | 0.290072 | 22.36461 | 22.51022 | 22.4066 |
| Q04323 | UBX domai | UBXN1 | UBX domai | 10.59554 | 2.96E-05 | 23.25796 | 23.3864 | 23.34272 |
| O15212 | Prefoldin su | PFDN6 | Prefoldin su | 10.63758 | 0.000609 | 22.79747 | 22.71055 | 22.92635 |
| Q96CS3 | FAS-associ | FAF2 | FAS-associ | 7.392591 | 0.034344 | 22.01662 | 21.83013 | 23.30814 |
| Q9NP81 | Serine--tRN | SARS2 | Serine--tRN | 5.716708 | 0.072619 | 22.94815 | 21.82291 | 23.53848 |
| Q93008 | Probable ul | USP9X | Probable ul | 9.281088 | 0.004733 | 22.20771 | 22.21718 | 22.11133 |
| Q92574 | Hamartin | TSC1 | Hamartin O: | 10.21239 | 7.16E-05 | 22.63159 | 22.55967 | 22.76004 |
| Q16795 | NADH dehy | NDUFA9 | NADH dehy | 6.432792 | 0.010693 | 23.17054 | 22.82085 | 22.09799 |
| Q9H5Z1 | Probable A | DHX35 | Probable A | 9.610716 | 1.03E-06 | 22.42096 | 22.49336 | 22.59327 |
| O75909 | Cyclin-K | CCNK | Cyclin-K OS | 10.26396 | 9.22E-06 | 23.0104 | 22.6051 | 22.80428 |
| Q9Y4Z0 | U6 snRNA-a | LSM4 | U6 snRNA-a | 9.029846 | 0.009264 | 23.83413 | 24.08383 | 24.37285 |
| Q8IYB1 | Protein MB2 | MB21D2 | Nucleotidyl | 0.867406 | 0.000309 | 22.9274 | 22.70468 | 22.82388 |
| O00469 | Procollager | PLOD2 | Procollager | 8.526831 | 0.012953 | 22.85555 | 22.71104 | 21.35192 |
| P27448 | MAP/microt | MARK3 | MAP/microt | 9.4337 | 0.000177 | 22.61178 | 22.55272 | 22.53796 |
| Q8WXI9 | Transcriptio | GATAD2B | Transcriptio | 10.79462 | 1.39E-05 | 23.22688 | 23.28292 | 23.31437 |
| P43686 | 26S proteas | PSMC4 | 26S proteas | 10.07734 | 1.66E-05 | 22.44139 | 22.3641 | 22.49668 |
| Q9Y4B6 | Protein VPR | VPRBP | DDB1- and | 10.50008 | 1.82E-05 | 23.02953 | 22.80235 | 22.75792 |
| Q16531 | DNA damag | DDB1 | DNA damag | 9.45521 | 2.95E-06 | 22.88878 | 22.69394 | 22.73254 |
| Q9Y305 | Acyl-coenzy | ACOT9 | Acyl-coenzy | 10.40191 | 1.39E-05 | 22.72151 | 22.45899 | 22.56586 |
| Q86UD1 | Out at first p | OAF | Out at first p | 10.81989 | 1.34E-05 | 23.63898 | 23.4188 | 23.57427 |
| Q9NR50 | Translation | EIF2B3 | Translation | 9.413227 | 0.01094 | 21.5882 | 23.31451 | 23.22844 |
| Q13895 | Bystin | BYSL | Bystin OS=I | 9.593843 | 1.27E-05 | 22.51138 | 22.2826 | 22.50727 |
| O95373 | Importin-7 | IPO7 | Importin-7 ( | 3.719896 | 0.002327 | 22.14485 | 23.66709 | 22.01475 |
| Q12899 | Tripartite mo | TRIM26 | Tripartite mo | 8.031729 | 0.000355 | 22.19608 | 22.33797 | 22.65864 |
| Q7Z2T5 | TRMT1-like | TRMT1L | TRMT1-like | 10.30918 | 0.000123 | 23.09088 | 22.94569 | 23.02146 |
| O95926 | Pre-mRNA-s | SYF2 | Pre-mRNA-s | 10.18003 | 0.000207 | 22.8101 | 22.73047 | 22.8535 |
| Q9NUG6 | p53 and DN | PDRG1 | p53 and DN | 9.973376 | 0.000939 | 22.99755 | 22.89164 | 23.09215 |
| Q9BRZ2 | E3 ubiquitir | TRIM56 | E3 ubiquitir | 10.54124 | 4.29E-05 | 22.62547 | 22.71249 | 22.89794 |
| Q5T3I0 | G patch dor | GPATCH4 | G patch dor | 9.827771 | 1.32E-05 | 22.64588 | 22.6368 | 22.61113 |
| Q96P11 | Probable 28 | NSUN5 | 28S rRNA (c | 9.607172 | 0.000571 | 22.33049 | 22.19997 | 22.28328 |
| Q99700 | Ataxin-2 | ATXN2 | Ataxin-2 OS | 6.950965 | 0.020214 | 23.02242 | 22.77147 | 20.61009 |
| Q96B26 | Exosome co | EXOSC8 | Exosome co | 5.38118 | 1.79E-06 | 22.90672 | 23.05963 | 23.23285 |
| P28288 | ATP-binding | ABCD3 | ATP-binding | 10.52428 | 1.44E-06 | 22.70018 | 22.59871 | 22.90998 |
| P08240 | Signal recog | SRPR | Signal recog | 9.276341 | 0.001076 | 22.75729 | 22.47025 | 22.73242 |
| Q15554 | Telomeric re | TERF2 | Telomeric re | 9.749523 | 1.80E-06 | 22.9316 | 22.5456 | 22.89019 |
| P28482 | Mitogen-act | MAPK1 | Mitogen-act | 5.168008 | 2.98E-05 | 22.70197 | 23.0142 | 22.74591 |
| Q9P0V3 | SH3 domai | SH3BP4 | SH3 domai | 10.39468 | 3.20E-06 | 23.60173 | 22.74041 | 23.41313 |
| Q1KMD3 | Heterogene | HNRNPUL2 | Heterogene | 10.62094 | 4.83E-05 | 23.52361 | 23.25753 | 23.42714 |
| Q9BWF2 | E3 ubiquitir | TRAIP | E3 ubiquitir | 9.471263 | 2.54E-05 | 22.89341 | 22.9925 | 23.2037 |
| P55884 | Eukaryotic t | EIF3B | Eukaryotic t | 7.944439 | 0.01381 | 22.84299 | 22.3041 | 21.50523 |
| Q9UQE7 | Structural n | SMC3 | Structural n | 4.515252 | 0.015421 | 20.31694 | 23.64867 | 20.33331 |

| | | | | | | | | |
|---|---|---|---|---|---|---|---|---|
| P35998 | 26S proteas | PSMC2 | 26S proteas | 7.134192 | 0.024658 | 21.89992 | 21.75898 | 23.00982 |
| P08559 | Pyruvate de | PDHA1 | Pyruvate de | 4.753713 | 4.75E-05 | 22.96853 | 22.34237 | 22.33928 |
| P36873 | Serine/threo | PPP1CC | Serine/threo | 10.12973 | 2.86E-05 | 23.97684 | 24.08464 | 24.43968 |
| P49757 | Protein num | NUMB | Protein num | 7.281459 | 0.008009 | 20.91225 | 23.01274 | 23.83846 |
| P62330 | ADP-ribosyl | ARF6 | ADP-ribosyl | 8.206407 | 0.007093 | 23.43619 | 23.15278 | 23.22682 |
| P62304 | Small nucle | SNRPE | Small nucle | 11.64491 | 8.35E-07 | 24.26757 | 24.39585 | 24.13038 |
| P56192 | Methionine- | MARS | Methionine- | 11.23947 | 0.000228 | 23.53148 | 23.44088 | 23.33129 |
| Q5VZL5 | Zinc finger M | ZMYM4 | Zinc finger M | 10.55566 | 4.88E-05 | 22.70181 | 22.73849 | 22.68419 |
| P60228 | Eukaryotic t | EIF3E | Eukaryotic t | 8.345635 | 4.29E-05 | 22.33748 | 21.62462 | 22.67691 |
| Q13769 | THO compl | THOC5 | THO compl | 9.797696 | 4.58E-05 | 23.20868 | 22.71597 | 23.13207 |
| P46977 | Dolichyl-di | STT3A | Dolichyl-di | 8.913338 | 4.40E-05 | 22.83076 | 22.77404 | 23.22948 |
| Q9HAU5 | Regulator o | UPF2 | Regulator o | 7.942996 | 0.025915 | 22.10172 | 21.61621 | 23.50688 |
| O76031 | ATP-depend | CLPX | ATP-depend | 9.387301 | 0.000387 | 23.00953 | 22.76953 | 22.91914 |
| P27824 | Calnexin | CANX | Calnexin OS | 8.181529 | 0.037899 | 22.92678 | 22.84571 | 23.5093 |
| P63162;P1 | Small nucle | SNRPN;SNI | Small nucle | 10.88992 | 0.000186 | 22.99303 | 22.97181 | 23.08361 |
| Q86WX3 | Active regul | RPS19BP1 | Active regul | 9.72053 | 0.029014 | 22.46752 | 23.57034 | 23.80961 |
| P49916 | DNA ligase : | LIG3 | DNA ligase : | 6.578239 | 0.035572 | 20.91918 | 22.76939 | 23.02756 |
| Q9NW82 | WD repeat-c | WDR70 | WD repeat-c | 8.868975 | 1.59E-05 | 22.5013 | 22.49126 | 22.14805 |
| Q9NQZ2 | Something | UTP3 | Something | 8.332065 | 0.02768 | 23.51003 | 22.20428 | 22.30833 |
| O43148 | mRNA cap g | RNMT | mRNA cap g | 7.082736 | 0.049721 | 23.16999 | 22.13768 | 22.3718 |
| Q8NEJ9 | Neuroguidir | NGDN | Neuroguidir | 11.09558 | 8.25E-06 | 22.44584 | 22.61401 | 22.56263 |
| Q9NVW2 | E3 ubiquitir | RLIM | E3 ubiquitir | 11.06152 | 2.16E-08 | 23.40145 | 23.33129 | 23.42752 |
| O76094 | Signal recog | SRP72 | Signal recog | 9.096413 | 0.00347 | 22.28365 | 22.07199 | 21.90866 |
| P57088 | Transmemb | TMEM33 | Transmemb | 8.313585 | 0.067835 | 24.1031 | 23.09463 | 23.69636 |
| Q9NPJ8 | NTF2-relate | NXT2 | NTF2-relate | 11.56409 | 1.49E-06 | 24.14063 | 24.00333 | 24.04943 |
| Q9Y5J1 | U3 small nu | UTP18 | U3 small nu | 5.913714 | 0.000613 | 22.85445 | 22.09973 | 21.99507 |
| Q8WXA9 | Splicing reg | SREK1 | Splicing reg | 2.563906 | 0.002867 | 22.47239 | 22.75666 | 23.15789 |
| Q96FV9 | THO compl | THOC1 | THO compl | 9.657246 | 0.000428 | 22.82458 | 22.59628 | 22.70358 |
| Q96GM8 | Target of EG | TOE1 | Target of EG | 11.08574 | 1.97E-07 | 22.70675 | 22.5925 | 22.6193 |
| Q00059 | Transcriptio | TFAM | Transcriptio | 7.977919 | 0.048018 | 23.27085 | 22.26513 | 23.14383 |
| Q92945 | Far upstrear | KHSRP | Far upstrear | 8.286169 | 0.013744 | 23.41248 | 22.12515 | 22.7496 |
| Q13610 | Periodic tryp | PWP1 | Periodic tryp | 10.11343 | 1.60E-06 | 23.0052 | 23.07873 | 22.86051 |
| O43795 | Unconventi | MYO1B | Unconventi | 7.190172 | 0.010966 | 21.88394 | 21.24394 | 23.27099 |
| Q8TEW0 | Partitioning | PARD3 | Partitioning | 10.03491 | 7.53E-07 | 22.1305 | 21.94642 | 21.99852 |
| Q9BRU9 | rRNA-proce | UTP23 | rRNA-proce | 10.62591 | 2.06E-05 | 23.54203 | 23.26342 | 23.60003 |
| Q9BZV1 | UBX domai | UBXN6 | UBX domai | 10.75819 | 9.34E-05 | 22.64636 | 22.7683 | 22.78758 |
| Q8N9T8 | Protein KRI1 | KRI1 | Protein KRI1 | 9.810879 | 6.17E-05 | 22.55494 | 22.02736 | 22.52174 |
| Q6Y7W6 | PERQ amin | GIGYF2 | GRB10-inte | 7.344217 | 0.026097 | 22.25442 | 22.33734 | 22.67467 |
| Q9Y5M8 | Signal recog | SRPRB | Signal recog | 7.433616 | 0.043535 | 22.6946 | 22.36488 | 23.77826 |
| Q9GZR2 | RNA exonuc | REXO4 | RNA exonuc | 9.836797 | 3.80E-05 | 22.90802 | 22.31116 | 22.71697 |
| Q96II8 | Leucine-rich | LRCH3 | DISP comp | 9.122216 | 0.000238 | 22.65938 | 22.6536 | 22.90349 |
| Q96JB6 | Lysyl oxidas | LOXL4 | Lysyl oxidas | 8.688406 | 0.036414 | 23.56512 | 21.94767 | 22.29558 |
| Q99550 | M-phase ph | MPHOSPH9 | M-phase ph | 7.223758 | 0.007599 | 21.25194 | 21.98238 | 22.83636 |
| Q7Z6E9 | E3 ubiquitir | RBBP6 | E3 ubiquitir | 2.745942 | 0.004989 | 23.52229 | 22.04023 | 23.40145 |
| P55084 | Trifunctiona | HADHB | Trifunctiona | 4.0192 | 0.000952 | 22.76444 | 22.68611 | 23.1765 |
| Q9P015 | 39S ribosor | MRPL15 | Large riboso | 8.822741 | 0.015179 | 23.99912 | 21.83596 | 21.8503 |
| Q9Y232 | Chromodor | CDYL | Chromodor | 5.368992 | 0.084849 | 22.75707 | 23.12365 | 22.50218 |
| O95707 | Ribonuclea | POP4 | Ribonuclea | 8.936546 | 0.002534 | 21.28025 | 22.98096 | 23.20482 |

| | | | | | | | | |
|---|---|---|---|---|---|---|---|---|
| Q8N8A6 | ATP-depen( | DDX51 | ATP-depen( | 11.33048 | 0.000131 | 23.02657 | 22.77193 | 22.80138 |
| Q9NY12 | H/ACA ribor | GAR1 | H/ACA ribor | 10.85447 | 9.82E-05 | 23.41287 | 23.25059 | 23.56651 |
| A1L0T0 | Acetolactat( | ILVBL | 2-hydroxya( | -0.43295 | 0.051364 | 22.01192 | 21.89888 | 22.39063 |
| Q9HCL2 | Glycerol-3-| | GPAM | Glycerol-3-| | 10.22912 | 3.54E-06 | 23.4467 | 23.23722 | 22.6511 |
| P49736 | DNA replica | MCM2 | DNA replica | 8.97427 | 0.0277 | 22.876 | 22.22111 | 21.92481 |
| Q9UHB9 | Signal recog | SRP68 | Signal recog | 10.22103 | 6.27E-06 | 22.24435 | 22.27073 | 22.61553 |
| P09001 | 39S ribosor | MRPL3 | Large ribosc | 7.424483 | 0.04686 | 23.09909 | 23.1254 | 23.05406 |
| P05783 | Keratin, typ( | KRT18 | Keratin, typ( | 4.8854 | 0.00032 | 22.28467 | 22.48898 | 22.27042 |
| Q96F07 | Cytoplasmi | CYFIP2 | Cytoplasmi | 10.72119 | 2.26E-06 | 22.95954 | 23.01333 | 23.05783 |
| Q6P161 | 39S ribosor | MRPL54 | Large ribosc | 6.818528 | 0.026484 | 23.73789 | 23.46401 | 23.6826 |
| Q13405 | 39S ribosor | MRPL49 | Large ribosc | 9.882102 | 4.35E-05 | 22.85426 | 22.78083 | 23.14801 |
| Q96GC5 | 39S ribosor | MRPL48 | Large ribosc | 8.681849 | 0.017903 | 23.24355 | 23.21457 | 23.2405 |
| P62979;P6 | Ubiquitin-4( | RPS27A;UE | Ubiquitin-rit | 1.264566 | 0.561376 | 23.26914 | 21.1798 | 21.08141 |
| P30419 | Glycylpepti | NMT1 | Glycylpepti | 9.136278 | 0.012991 | 22.61329 | 22.56939 | 22.6764 |
| Q9H8H2 | Probable AT | DDX31 | Probable AT | 3.482347 | 0.000195 | 22.69192 | 22.32636 | 23.23879 |
| O94992 | Protein HEX | HEXIM1 | Protein HEX | 10.13848 | 0.0004 | 23.08908 | 23.04141 | 23.07468 |
| Q00577 | Transcriptio | PURA | Transcriptio | 10.54459 | 0.001012 | 23.27427 | 23.18903 | 23.49338 |
| Q03135 | Caveolin-1 | CAV1 | Caveolin-1 | 10.65813 | 2.37E-05 | 23.62466 | 22.76474 | 23.69965 |
| P18859 | ATP syntha: | ATP5J | ATP syntha: | 9.730596 | 0.002366 | 23.9227 | 23.59024 | 23.73003 |
| P62879;Q9 | Guanine nu | GNB2;GNB4 | Guanine nu | 8.964156 | 0.013321 | 23.38574 | 23.30828 | 23.48542 |
| P84077;P6 | ADP-ribosyl | ARF1;ARF3 | ADP-ribosyl | 10.33098 | 0.000552 | 23.07485 | 23.2863 | 23.04971 |
| P00558 | Phosphogly | PGK1 | Phosphogly | 6.042381 | 0.210458 | 23.95029 | 19.22419 | 20.0713 |
| P57772 | Selenocyst( | EEFSEC | Selenocyst( | 9.981636 | 4.96E-05 | 22.79309 | 22.65119 | 22.8363 |
| Q9Y6A4 | Cilia- and fl: | CFAP20 | Cilia- and fl: | 8.184477 | 0.015697 | 22.86111 | 22.53298 | 22.99623 |
| P51610 | Host cell fa( | HCFC1 | Host cell fa( | 10.02339 | 7.68E-05 | 22.43431 | 22.17566 | 22.47736 |
| P35606 | Coatomer s | COPB2 | Coatomer s | 9.518079 | 3.55E-05 | 22.68172 | 22.34879 | 22.56118 |
| Q9H6S0 | Probable AT | YTHDC2 | 3-5 RNA hel | 9.235686 | 3.18E-05 | 22.06329 | 22.2338 | 22.3493 |
| P07951 | Tropomyos | TPM2 | Tropomyos | 6.576533 | 0.085338 | 21.58036 | 22.99885 | 24.32629 |
| O95613 | Pericentrin | PCNT | Pericentrin ( | 7.729225 | 0.055565 | 21.72908 | 22.11337 | 22.08359 |
| Q99613;B5 | Eukaryotic t | EIF3C;EIF3 | Eukaryotic t | 6.087687 | 0.031301 | 20.79867 | 22.95541 | 22.60248 |
| Q9NV31 | U3 small nu | IMP3 | U3 small nu | 10.60541 | 0.000201 | 22.69073 | 22.65563 | 22.79237 |
| Q8WUQ7 | Cactin | CACTIN | Splicing fac | 7.949943 | 0.017487 | 23.68955 | 21.86049 | 21.97858 |
| P49711;Q8 | Transcriptio | CTCF;CTCF | Transcriptio | 10.85853 | 2.25E-05 | 23.23968 | 23.0358 | 23.03051 |
| P32322 | Pyrroline-5- | PYCR1 | Pyrroline-5- | 9.576515 | 1.88E-06 | 22.68791 | 22.74538 | 22.60858 |
| Q9NQ29 | Putative RN | LUC7L | Putative RN | 6.875385 | 0.0499 | 23.71062 | 23.70102 | 21.76353 |
| Q1ED39 | Lysine-rich | KNOP1 | Lysine-rich | 7.656124 | 0.069334 | 23.06676 | 22.30109 | 22.28226 |
| Q9Y3D5 | 28S ribosor | MRPS18C | Small ribos | 10.99233 | 5.65E-05 | 23.40691 | 23.34381 | 23.37504 |
| Q71RC2 | La-related p | LARP4 | La-related p | 9.752798 | 9.86E-05 | 22.47231 | 22.36306 | 22.46508 |
| Q96IX5 | Up-regulate | USMG5 | ATP syntha: | 12.49126 | 0.000159 | 23.79491 | 23.49033 | 23.73562 |
| P63279 | SUMO-conj | UBE2I | SUMO-conj | 10.97939 | 2.58E-05 | 23.4545 | 23.24537 | 23.20717 |
| Q9UHB6 | LIM domair | LIMA1 | LIM domair | 6.42888 | 0.013527 | 20.15536 | 21.92231 | 23.2226 |
| O15446 | DNA-directe | CD3EAP | DNA-directe | 10.55583 | 0.000289 | 23.32265 | 23.37172 | 23.53386 |
| Q14165 | Malectin | MLEC | Malectin OS | 8.296317 | 0.007908 | 22.94049 | 22.53604 | 22.60734 |
| L0R819 | | ASNSD1 | ASNSD1 up | 10.30165 | 7.37E-05 | 23.25768 | 22.80455 | 23.22269 |
| O60678 | Protein argi | PRMT3 | Protein argi | 10.36639 | 0.000608 | 22.75252 | 22.63076 | 22.96577 |
| Q6PJT7 | Zinc finger ( | ZC3H14 | Zinc finger ( | 7.497744 | 0.056885 | 22.38195 | 22.40523 | 23.21282 |
| Q58FF8 | Putative hea | HSP90AB2I | Putative hea | 11.06953 | 3.55E-05 | 23.35043 | 23.49375 | 23.57277 |
| P14649 | Myosin ligh | MYL6B | Myosin ligh | 7.376696 | 0.006172 | 22.02817 | 22.35504 | 23.0863 |

| | | | | | | | | |
|---|---|---|---|---|---|---|---|---|
| P09661 | U2 small nu | SNRPA1 | U2 small nu | 3.879279 | 0.000388 | 23.27213 | 22.97433 | 23.31285 |
| Q5SY16 | Polynucleo | NOL9 | Polynucleo | 11.81011 | 2.48E-06 | 23.81235 | 23.64218 | 23.87839 |
| P07203 | Glutathione | GPX1 | Glutathione | 7.933202 | 0.01809 | 21.92568 | 23.39362 | 21.78923 |
| P35659 | Protein DEK | DEK | Protein DEK | 7.008123 | 0.043564 | 22.87738 | 22.96386 | 23.24815 |
| Q96FQ6 | Protein S10 | S100A16 | Protein S10 | 8.872502 | 0.026217 | 22.88757 | 22.90652 | 22.96289 |
| Q96I51 | Williams-Be | WBSCR16 | RCC1-like G | 10.52972 | 4.09E-06 | 23.10265 | 22.95748 | 23.15201 |
| Q96P70 | Importin-9 | IPO9 | Importin-9 ( | 9.915343 | 0.000763 | 22.31788 | 22.18201 | 22.53205 |
| P49959 | Double-stra | MRE11A | Double-stra | 6.459632 | 0.023148 | 22.50472 | 21.44304 | 23.0748 |
| P62875 | DNA-directe | POLR2L | DNA-directe | 10.99251 | 4.77E-06 | 23.37332 | 22.97165 | 23.30466 |
| P61221 | ATP-binding | ABCE1 | ATP-binding | 10.33142 | 0.000761 | 23.00982 | 22.99835 | 23.00975 |
| P62191 | 26S proteas | PSMC1 | 26S proteas | 8.275956 | 0.010262 | 22.63157 | 22.36081 | 22.73374 |
| Q92504 | Zinc transpo | SLC39A7 | Zinc transpo | 11.04855 | 3.58E-05 | 23.65807 | 23.40211 | 23.41015 |
| Q9UN81 | LINE-1 retro | L1RE1 | LINE-1 retro | 10.25737 | 0.00029 | 22.65018 | 22.40343 | 22.92283 |
| Q08379 | Golgin subf | GOLGA2 | Golgin subf | 9.177219 | 1.62E-06 | 22.67747 | 22.29199 | 22.42122 |
| Q9HAH7 | Probable fib | FBRS | Probable fib | 10.53638 | 5.67E-05 | 22.68639 | 22.74096 | 22.819 |
| Q04917 | 14-3-3 prot | YWHAH | 14-3-3 prot | 10.67205 | 4.71E-06 | 22.94616 | 22.54767 | 22.59202 |
| P19447 | TFIIH basal | ERCC3 | General tran | 9.79353 | 1.66E-07 | 22.15749 | 22.13759 | 22.35264 |
| O43823 | A-kinase an | AKAP8 | A-kinase an | 9.584479 | 0.000189 | 22.50215 | 22.21656 | 22.47778 |
| Q9Y262 | Eukaryotic t | EIF3L | Eukaryotic t | 8.113141 | 0.018043 | 22.20908 | 21.92441 | 22.26032 |
| O95602 | DNA-directe | POLR1A | DNA-directe | 10.37898 | 1.97E-05 | 22.18779 | 22.23678 | 22.28348 |
| Q13835 | Plakophilin | PKP1 | Plakophilin | 3.319935 | 0.345843 | 22.9466 | 20.57052 | 21.38543 |
| Q8NBQ5 | Estradiol 17 | HSD17B11 | Estradiol 17 | 10.85858 | 1.62E-05 | 23.15587 | 22.53716 | 22.62658 |
| Q00587 | Cdc42 effec | CDC42EP1 | Cdc42 effec | 6.900079 | 0.052014 | 22.45543 | 22.171 | 22.64216 |
| P22234 | Multifunctic | PAICS | Bifunctiona | 8.775448 | 0.008298 | 22.9996 | 23.04577 | 23.13471 |
| P48730 | Casein kina | CSNK1D | Casein kina | 9.484462 | 0.000243 | 22.93287 | 22.7777 | 23.17776 |
| Q5T160 | Probable ar | RARS2 | Probable ar | -0.22544 | 0.592985 | 22.58896 | 22.59421 | 21.49682 |
| Q6IN84 | rRNA methy | MRM1 | rRNA methy | 11.51916 | 8.62E-05 | 23.02678 | 22.82227 | 23.19411 |
| Q8IZL8 | Proline-, glu | PELP1 | Proline-, glu | 8.04436 | 0.003881 | 22.39383 | 20.72609 | 22.8672 |
| P31946 | 14-3-3 prot | YWHAB | 14-3-3 prot | 7.884943 | 0.046836 | 23.50785 | 23.05452 | 23.19051 |
| Q53EP0 | Fibronectin | FNDC3B | Fibronectin | 7.884681 | 0.024031 | 22.52557 | 22.2105 | 23.98841 |
| Q5T280 | Putative me | C9orf114 | Putative me | 11.25809 | 1.69E-05 | 22.56962 | 22.58068 | 22.71354 |
| Q53EZ4 | Centrosom | CEP55 | Centrosom | 9.098337 | 0.000467 | 22.60316 | 22.47922 | 22.63814 |
| Q9ULW3 | Activator of | ABT1 | Activator of | 1.577902 | 0.06231 | 23.89615 | 22.13605 | 21.9775 |
| P56537 | Eukaryotic t | EIF6 | Eukaryotic t | 7.211243 | 0.043085 | 22.90973 | 22.60815 | 22.88291 |
| P18615 | Negative elo | NELFE | Negative elo | 8.297399 | 0.025578 | 22.87283 | 22.43301 | 22.58891 |
| Q15293 | Reticulocal | RCN1 | Reticulocal | 6.137099 | 0.089377 | 23.05523 | 23.31465 | 21.73302 |
| Q15459 | Splicing fac | SF3A1 | Splicing fac | 2.691903 | 0.002015 | 22.49743 | 22.12607 | 22.68329 |
| Q9H6R0 | Putative ATI | DHX33 | ATP-depend | 9.483301 | 1.57E-05 | 21.96659 | 21.96392 | 22.1181 |
| P22061 | Protein-L-is | PCMT1 | Protein-L-is | 1.254559 | 0.005032 | 22.72232 | 22.7218 | 22.92535 |
| Q8WUM0 | Nuclear por | NUP133 | Nuclear por | 8.003244 | 0.007477 | 21.14725 | 20.97337 | 23.3534 |
| Q9UJV9 | Probable AT | DDX41 | Probable AT | 7.818837 | 0.035165 | 21.61657 | 22.64562 | 22.75776 |
| P04899;P6 | Guanine nu | GNAI2;GNA | Guanine nu | 4.231778 | 0.004126 | 20.9624 | 23.09483 | 23.12485 |
| O15269 | Serine palm | SPTLC1 | Serine palm | 4.53016 | 0.119572 | 21.90734 | 22.1961 | 23.46899 |
| P09234 | U1 small nu | SNRPC | U1 small nu | 10.33748 | 1.39E-05 | 23.3306 | 22.90659 | 23.23908 |
| Q01085 | Nucleolysin | TIAL1 | Nucleolysin | 5.998534 | 0.004283 | 20.23748 | 22.59327 | 23.40211 |
| Q96TA2 | ATP-depend | YME1L1 | ATP-depend | 9.30978 | 0.000399 | 22.308 | 22.24905 | 22.35386 |
| P68366 | Tubulin alp | TUBA4A | Tubulin alp | 7.679327 | 0.103529 | 23.30299 | 22.35568 | 22.46877 |
| Q9H147 | Deoxynucle | DNTTIP1 | Deoxynucle | 10.01104 | 0.000137 | 22.70424 | 22.61881 | 22.86624 |

| | | | | | | | | |
|---|---|---|---|---|---|---|---|---|
| Q15058 | Kinesin-like | KIF14 | Kinesin-like | 9.344988 | 0.000269 | 21.92738 | 21.82147 | 21.98357 |
| Q9UJZ1 | Stomatin-lil | STOML2 | Stomatin-lil | 6.941991 | 0.039882 | 22.31038 | 22.71515 | 22.36407 |
| O14773 | Tripeptidyl-| | TPP1 | Tripeptidyl-| | 11.03457 | 9.90E-05 | 23.27739 | 22.97787 | 23.12465 |
| O43776 | Asparagine- | NARS | Asparagine- | 2.665805 | 0.011114 | 22.80841 | 23.28616 | 21.37565 |
| Q9P031 | Thyroid tran | CCDC59 | Thyroid tran | 8.23505 | 0.010876 | 22.25018 | 22.226 | 22.43576 |
| Q6PD62 | RNA polym | CTR9 | RNA polym | 7.747866 | 0.000867 | 22.12783 | 21.47927 | 22.37196 |
| Q6PI48 | Aspartate--t | DARS2 | Aspartate--t | 9.986991 | 0.000119 | 22.78395 | 22.70761 | 22.75908 |
| Q2PPJ7 | Ral GTPase- | RALGAPA2 | Ral GTPase- | 12.49053 | 5.87E-05 | 23.75256 | 23.78107 | 23.54369 |
| Q9NPI1 | Bromodom | BRD7 | Bromodom | 9.795527 | 3.89E-06 | 22.46021 | 22.25096 | 22.38688 |
| Q9Y6V7 | Probable AT | DDX49 | Probable AT | 10.22525 | 1.40E-06 | 22.21071 | 22.07852 | 22.44852 |
| O00303 | Eukaryotic t | EIF3F | Eukaryotic t | 8.297109 | 0.03624 | 22.13649 | 22.73866 | 23.07886 |
| P28702 | Retinoic aci | RXRB | Retinoic aci | 11.61572 | 0.000106 | 22.35178 | 22.28732 | 22.53331 |
| Q13033 | Striatin-3 | STRN3 | Striatin-3 OS | 10.85461 | 0.005249 | 22.54971 | 22.65626 | 22.63881 |
| Q9H223 | EH domain- | EHD4 | EH domain- | 9.734908 | 2.13E-05 | 22.19991 | 22.28331 | 22.13859 |
| O75940 | Survival of r | SMNDC1 | Survival of r | 6.836364 | 0.01711 | 22.31501 | 22.26937 | 22.65294 |
| P42224 | Signal trans | STAT1 | Signal trans | 4.614027 | 0.001588 | 21.81421 | 22.00027 | 22.83457 |
| Q96GD4 | Aurora kina | AURKB | Aurora kina | 10.46537 | 4.62E-05 | 22.76654 | 22.48203 | 22.69392 |
| Q9Y388 | RNA-bindin | RBMX2 | RNA-bindin | 7.696164 | 0.034025 | 22.27136 | 22.78205 | 23.66871 |
| P13489 | Ribonuclea | RNH1 | Ribonuclea | 8.06859 | 0.043228 | 23.13232 | 22.40346 | 22.69645 |
| O95433 | Activator of | AHSA1 | Activator of | 10.36912 | 0.000107 | 22.75481 | 22.70073 | 22.57342 |
| O75592 | E3 ubiquitir | MYCBP2 | E3 ubiquitir | 7.952686 | 3.35E-07 | 21.47011 | 21.35719 | 21.64588 |
| Q16666 | Gamma-int | IFI16 | Gamma-int | 8.21715 | 0.014872 | 22.34224 | 22.06112 | 21.06979 |
| Q8TDX7;Q9 | Serine/threo | NEK7;NEK6 | Serine/threo | 8.464138 | 0.007312 | 23.01915 | 22.66325 | 22.6333 |
| Q15334 | Lethal(2) gia | LLGL1 | Lethal(2) gia | 7.400321 | 0.049441 | 22.37515 | 22.62127 | 21.84525 |
| P20908 | Collagen al | COL5A1 | Collagen al | 10.17842 | 0.000613 | 22.7536 | 22.95081 | 22.77292 |
| Q6NW34 | Uncharacte | C3orf17 | Nucleolus a | 9.705045 | 0.000101 | 22.60601 | 22.52442 | 22.57464 |
| Q9UI30 | Multifunctic | TRMT112 | Multifunctic | 8.548498 | 0.010158 | 23.49057 | 23.13353 | 22.8171 |
| Q99714 | 3-hydroxyac | HSD17B10 | 3-hydroxyac | 8.942549 | 0.004121 | 22.38398 | 22.24722 | 22.26046 |
| O75818 | Ribonuclea | RPP40 | Ribonuclea | 7.799479 | 0.030204 | 23.63556 | 21.91649 | 22.02422 |
| Q9NVF7 | F-box only p | FBXO28 | F-box only p | 9.873023 | 2.30E-07 | 22.65955 | 22.40072 | 22.47563 |
| Q9H0U3 | Magnesium | MAGT1 | Magnesium | 5.751912 | 0.000958 | 22.95608 | 20.96035 | 22.30354 |
| Q6PJI9 | WD repeat-c | WDR59 | GATOR2 co | 9.394639 | 0.000361 | 22.50567 | 22.72217 | 22.6435 |
| Q13753 | Laminin sul | LAMC2 | Laminin sul | 10.2352 | 0.00017 | 23.75735 | 23.46438 | 23.19589 |
| Q9Y2K7 | Lysine-spec | KDM2A | Lysine-spec | 9.58274 | 0.000837 | 22.25647 | 22.1771 | 22.30936 |
| Q6ZUT6 | Uncharacte | C15orf52 | Coiled-coil | 4.971338 | 0.0612 | 22.72829 | 23.36399 | 21.3819 |
| P55735 | Protein SEC | SEC13 | Protein SEC | 8.320131 | 0.05531 | 22.95378 | 23.73324 | 22.5574 |
| Q15369 | Transcriptio | TCEB1 | Elongin-C O | 3.954676 | 0.007259 | 22.50417 | 23.63677 | 21.09963 |
| P46199 | Translation | MTIF2 | Translation | 10.02902 | 0.000112 | 22.79753 | 22.74794 | 23.0184 |
| P13674 | Prolyl 4-hyc | P4HA1 | Prolyl 4-hyc | 7.984202 | 0.021917 | 21.59409 | 20.98357 | 23.10861 |
| Q8IWI9 | MAX gene-a | MGA | MAX gene-a | 10.27731 | 0.000405 | 22.94342 | 22.53089 | 22.67433 |
| P78316 | Nucleolar p | NOP14 | Nucleolar p | 7.663639 | 0.020257 | 23.08245 | 22.30975 | 21.57325 |
| Q96EY1 | DnaJ homo | DNAJA3 | DnaJ homo | 10.25262 | 6.65E-07 | 22.94555 | 22.6981 | 22.91042 |
| P53597 | Succinyl-Co | SUCLG1 | Succinate--( | 6.580028 | 0.01646 | 23.11073 | 22.88041 | 22.95973 |
| Q9NXS2 | Glutaminyl- | QPCTL | Glutaminyl- | 10.077 | 1.57E-05 | 22.46605 | 22.53433 | 22.80373 |
| O94905;O7 | Erlin-2;Erlin | ERLIN2;ERI | Erlin-2 OS= | 10.61949 | 1.43E-07 | 23.05583 | 22.88998 | 23.18144 |
| Q9UER7 | Death doma | DAXX | Death doma | 11.63717 | 5.08E-05 | 23.7682 | 23.2727 | 23.5852 |
| Q9NR56 | Muscleblin | MBNL1 | Muscleblin | 7.920662 | 0.003976 | 23.24584 | 23.10928 | 23.26284 |
| Q9NXW2 | DnaJ homo | DNAJB12 | DnaJ homo | 6.629069 | 0.005024 | 21.08453 | 22.05165 | 22.37257 |

| | | | | | | | | |
|---|---|---|---|---|---|---|---|---|
| Q13017 | Rho GTPase | ARHGAP5 | Rho GTPase | 9.765251 | 7.75E-08 | 22.39605 | 22.19421 | 22.53664 |
| P45880 | Voltage-dep | VDAC2 | Voltage-dep | 4.298269 | 0.23754 | 23.09952 | 22.31069 | 21.91692 |
| Q8ND56 | Protein LSM | LSM14A | Protein LSM | 2.134064 | 0.071647 | 22.52884 | 21.85228 | 24.14492 |
| O14802 | DNA-directe | POLR3A | DNA-directe | 7.561767 | 0.033381 | 21.23316 | 22.62064 | 21.50877 |
| Q6WKZ4 | Rab11 fami | RAB11FIP1 | Rab11 fami | 3.808189 | 0.022951 | 21.14967 | 20.6796 | 23.96869 |
| Q6P582;Q6 | Mitotic-spin | MZT2A;MZT | Mitotic-spin | 10.29873 | 0.000464 | 22.79062 | 22.50402 | 22.6132 |
| Q9BW92 | Threonine-- | TARS2 | Threonine-- | 7.631115 | 0.010419 | 22.87113 | 22.40364 | 21.34624 |
| Q86W42 | THO compl | THOC6 | THO compl | 10.0104 | 0.000107 | 22.27651 | 22.28184 | 22.53065 |
| Q8IXK0 | Polyhomeo | PHC2 | Polyhomeo | 8.443395 | 0.006407 | 23.23298 | 22.99224 | 23.33252 |
| Q9HC36 | rRNA methy | RNMTL1 | rRNA methy | 9.770165 | 9.97E-05 | 23.09603 | 22.98852 | 23.02494 |
| Q9P035 | Very-long-c | HACD3 | Very-long-c | 10.15875 | 0.000146 | 22.5302 | 22.30772 | 22.65246 |
| Q99460 | 26S proteas | PSMD1 | 26S proteas | 3.923269 | 0.016907 | 23.36613 | 20.30864 | 20.54906 |
| P48681 | Nestin | NES | Nestin OS=I | 7.784787 | 0.047001 | 21.45934 | 21.77016 | 22.57455 |
| Q9H4M9 | EH domain- | EHD1 | EH domain- | 9.953093 | 0.0003 | 22.82388 | 22.46476 | 22.99104 |
| Q9HCN8 | Stromal cel | SDF2L1 | Stromal cel | 11.22741 | 0.00028 | 22.98891 | 23.57623 | 23.28093 |
| Q05048 | Cleavage st | CSTF1 | Cleavage st | 8.45425 | 0.019432 | 23.23153 | 22.81691 | 23.23866 |
| Q9UBM7 | 7-dehydroc | DHCR7 | 7-dehydroc | 7.978512 | 0.024192 | 23.3254 | 22.88618 | 21.96585 |
| P29558;Q1 | RNA-bindin | RBMS1;RBM | RNA-bindin | 10.64777 | 1.78E-05 | 23.1377 | 22.87491 | 23.16276 |
| Q5SRE5 | Nucleopori | NUP188 | Nucleopori | 6.564138 | 0.024505 | 22.05933 | 21.8395 | 21.26762 |
| Q13155 | Aminoacyl t | AIMP2 | Aminoacyl t | 9.030807 | 0.005833 | 23.4472 | 23.15792 | 23.25048 |
| Q96MX3 | Zinc finger p | ZNF48 | Zinc finger p | 10.82132 | 0.000172 | 22.82279 | 22.71396 | 22.78931 |
| Q8IYU8 | Calcium up | MICU2 | Calcium up | 9.159937 | 0.00018 | 22.56674 | 22.66965 | 22.57136 |
| P08134;P6 | Rho-related | RHOC;RHO | Rho-related | 9.267586 | 0.007274 | 23.41596 | 23.20473 | 23.22137 |
| O75475 | PC4 and SF | PSIP1 | PC4 and SF | 8.394095 | 0.005754 | 22.62469 | 22.68744 | 22.8659 |
| Q8NEF9 | Serum resp | SRFBP1 | Serum resp | 10.364 | 0.000167 | 22.99495 | 23.04861 | 22.73694 |
| Q9BQE3 | Tubulin alp | TUBA1C | Tubulin alp | 10.1583 | 0.000288 | 23.20068 | 23.15609 | 23.15379 |
| O14874 | [3-methyl-2 | BCKDK | [3-methyl-2 | 9.513556 | 0.000853 | 22.56892 | 22.18891 | 22.40966 |
| P22392;O6 | Nucleoside | NME2;NME | Nucleoside | 3.092592 | 0.056088 | 23.26112 | 21.19736 | 21.29648 |
| Q8IWS0 | PHD finger p | PHF6 | PHD finger p | 10.85907 | 4.62E-05 | 23.28051 | 23.38522 | 24.0164 |
| Q13613 | Myotubular | MTMR1 | Myotubular | 9.804322 | 2.72E-06 | 22.5705 | 22.71542 | 22.64313 |
| Q53H96 | Pyrroline-5- | PYCRL | Pyrroline-5- | 10.01045 | 2.37E-06 | 22.72775 | 22.52306 | 22.57794 |
| Q9HAU0 | Pleckstrin h | PLEKHA5 | Pleckstrin h | 9.252622 | 2.54E-06 | 22.43578 | 22.21881 | 22.308 |
| O43837 | Isocitrate de | IDH3B | Isocitrate de | 9.930158 | 0.000103 | 22.64427 | 22.36568 | 22.47946 |
| P31944 | Caspase-14 | CASP14 | Caspase-14 | 2.054683 | 0.632987 | 21.05019 | 18.72324 | 19.73962 |
| Q9P032 | NADH dehy | NDUFAF4 | NADH dehy | 7.934299 | 0.012323 | 22.81329 | 21.3208 | 22.83235 |
| P22570 | NADPH:adr | FDXR | NADPH:adr | 9.726654 | 6.82E-06 | 21.86827 | 22.00237 | 21.9693 |
| P09211 | Glutathione | GSTP1 | Glutathione | 5.341747 | 0.228783 | 24.03993 | 19.90395 | 19.61407 |
| Q9BSD7 | Cancer-rela | NTPCR | Cancer-rela | 9.407103 | 0.006866 | 22.52303 | 22.55761 | 22.45098 |
| Q9BU76 | Multiple my | MMTAG2 | Multiple my | 8.685773 | 0.015542 | 23.20198 | 22.82627 | 23.40015 |
| P57740 | Nuclear por | NUP107 | Nuclear por | 3.817706 | 0.018378 | 20.53098 | 20.56969 | 23.51244 |
| P37802 | Transgelin-2 | TAGLN2 | Transgelin-2 | 4.548051 | 0.011181 | 22.85768 | 22.81738 | 20.81198 |
| Q9HCU5 | Prolactin re | PREB | Prolactin re | 9.672768 | 0.000215 | 22.13477 | 21.8651 | 22.19928 |
| O43293 | Death-asso | DAPK3 | Death-asso | 8.69867 | 0.000282 | 22.2262 | 22.10194 | 22.49517 |
| Q99470 | Stromal cel | SDF2 | Stromal cel | 10.87242 | 0.003466 | 23.2417 | 23.28178 | 22.88413 |
| P43358 | Melanoma- | MAGEA4 | Melanoma- | 10.87906 | 0.000396 | 22.74061 | 23.02504 | 23.0776 |
| Q15046 | Lysine--tRN | KARS | Lysine--tRN | 5.339013 | 0.038482 | 22.62812 | 22.20428 | 21.59336 |
| P51153;Q9 | Ras-related | RAB13;RAB | Ras-related | 9.234654 | 0.002484 | 21.72376 | 24.63068 | 20.89349 |
| Q9GZS3 | WD repeat-c | WDR61 | Superkiller | 9.907398 | 3.11E-05 | 22.43911 | 22.6433 | 22.75727 |

| | | | | | | | | |
|---|---|---|---|---|---|---|---|---|
| Q03188 | Centromere | CENPC | Centromere | 9.212286 | 0.00028 | 22.18559 | 21.97614 | 22.38874 |
| O43447 | Peptidyl-pro | PPIH | Peptidyl-pro | 9.257278 | 0.001297 | 22.74854 | 22.37486 | 22.62105 |
| Q07021 | Compleme | C1QBP | Compleme | 10.45379 | 0.00015 | 23.4179 | 23.09986 | 23.42278 |
| P52434 | DNA-directe | POLR2H | DNA-directe | 9.811594 | 6.05E-08 | 22.62223 | 22.49341 | 22.48486 |
| O60701 | UDP-glucos | UGDH | UDP-glucos | 6.161313 | 0.085415 | 23.07302 | 21.69799 | 22.08024 |
| P62491;Q1 | Ras-related | RAB11A;RA | Ras-related | 2.817083 | 0.014341 | 22.77473 | 21.49809 | 20.5383 |
| Q96FK6 | WD repeat-c | WDR89 | WD repeat-c | 9.665686 | 3.18E-05 | 22.6831 | 22.61643 | 22.76235 |
| O43709 | Probable 18 | WBSCR22 | Probable 18 | 7.128393 | 0.044826 | 21.69706 | 22.73254 | 21.70058 |
| Q9HD45 | Transmemb | TM9SF3 | Transmemb | 11.28438 | 1.90E-06 | 23.04961 | 23.15388 | 23.16426 |
| A0A3B3IU4 | RNMT-activa | FAM103A1 | RNA guanin | 7.762708 | 0.039384 | 23.00664 | 23.72109 | 22.72648 |
| Q96DH6 | RNA-bindin | MSI2 | RNA-bindin | 9.744523 | 0.001019 | 22.19081 | 22.09803 | 22.41573 |
| P48444 | Coatomer s | ARCN1 | Coatomer s | 6.979128 | 0.028415 | 22.60513 | 22.42655 | 21.56987 |
| Q7KZF4 | Staphyloco | SND1 | Staphyloco | 5.20481 | 0.088658 | 22.45839 | 22.21392 | 21.20365 |
| P48594 | Serpin B4 | SERPINB4 | Serpin B4 O | 2.678288 | 0.524021 | 20.73265 | 20.67288 | 19.23093 |
| Q8N2K0 | Monoacylgl | ABHD12 | Lysophosp | 7.512067 | 0.050367 | 23.5354 | 22.42409 | 22.65353 |
| Q14160 | Protein scri | SCRIB | Protein scri | 9.108643 | 3.43E-06 | 22.12726 | 22.39511 | 22.61054 |
| Q674X7 | Kazrin | KAZN | Kazrin OS=H | 7.089722 | 0.051863 | 22.26888 | 23.0613 | 22.46996 |
| Q5T2T1 | MAGUK p55 | MPP7 | MAGUK p55 | 7.725431 | 0.029517 | 21.26602 | 21.92817 | 22.81137 |
| P51659 | Peroxisoma | HSD17B4 | Peroxisoma | 1.075715 | 0.385647 | 21.74431 | 21.30129 | 20.93582 |
| P62310 | U6 snRNA-a | LSM3 | U6 snRNA-a | 7.961256 | 0.036387 | 23.20707 | 22.80603 | 22.98278 |
| Q12834 | Cell divisior | CDC20 | Cell divisior | 9.121094 | 4.26E-05 | 22.18129 | 22.05305 | 22.26888 |
| P61160 | Actin-relate | ACTR2 | Actin-relate | 10.6018 | 0.000561 | 22.55262 | 22.63682 | 22.55494 |
| O75461 | Transcriptio | E2F6 | Transcriptio | 9.697863 | 7.97E-07 | 22.37618 | 22.34743 | 22.32133 |
| P08195 | 4F2 cell-sur | SLC3A2 | Amino acid | 10.62555 | 0.000269 | 22.57052 | 22.71194 | 22.98669 |
| P13995 | Bifunctiona | MTHFD2 | Bifunctiona | 8.491055 | 0.016672 | 23.18083 | 22.72655 | 23.11191 |
| O94973 | AP-2 compl | AP2A2 | AP-2 compl | 8.9047 | 0.000195 | 21.86026 | 21.73979 | 22.01185 |
| P56182 | Ribosomal | RRP1 | Ribosomal | 7.983877 | 0.029624 | 22.42939 | 22.12909 | 22.38998 |
| Q15020 | Squamous | SART3 | Squamous | 9.103546 | 0.000193 | 21.78963 | 21.8259 | 21.6823 |
| Q8IXB1 | DnaJ homo | DNAJC10 | DnaJ homo | 9.358086 | 1.14E-05 | 22.20305 | 22.08002 | 22.0082 |
| P07384 | Calpain-1 c | CAPN1 | Calpain-1 c | -0.09814 | 0.881613 | 21.76032 | 21.41266 | 22.01947 |
| O43251;Q9 | RNA binding | RBFOX2;RB | RNA binding | 9.192346 | 1.28E-05 | 22.72561 | 22.31833 | 22.39976 |
| Q99590 | Protein SCA | SCAF11 | Protein SCA | 5.091193 | 0.227266 | 21.65714 | 22.40621 | 21.73409 |
| Q9Y314 | Nitric oxide | NOSIP | Nitric oxide | 7.525934 | 0.036324 | 22.87046 | 22.73866 | 22.08722 |
| Q6P1J9 | Parafibromi | CDC73 | Parafibromi | 7.700177 | 0.051937 | 21.77419 | 21.5134 | 22.65635 |
| P11310 | Medium-ch | ACADM | Medium-ch | 8.337849 | 0.01411 | 22.45141 | 22.41607 | 22.39409 |
| Q7Z7K6 | Centromere | CENPV | Centromere | 9.913505 | 8.74E-06 | 22.42273 | 22.7162 | 22.9255 |
| Q05639 | Elongation | EEF1A2 | Elongation | 9.26387 | 1.08E-05 | 22.86703 | 22.64904 | 22.55 |
| P63000 | Ras-related | RAC1 | Ras-related | 9.117536 | 0.000366 | 22.49982 | 22.04983 | 22.38298 |
| Q9NVJ2 | ADP-ribosyl | ARL8B | ADP-ribosyl | 7.59284 | 0.053427 | 21.08083 | 23.08979 | 23.26699 |
| P31153 | S-adenosyl | MAT2A | S-adenosyl | 10.81321 | 4.67E-06 | 22.5673 | 22.1801 | 22.47321 |
| O00505 | Importin su | KPNA3 | Importin su | 10.07974 | 7.87E-09 | 22.69275 | 22.56076 | 22.56923 |
| P61163 | Alpha-centr | ACTR1A | Alpha-centr | 7.748759 | 0.052697 | 21.804 | 22.79598 | 22.08583 |
| Q8N7H5 | RNA polym | PAF1 | RNA polym | 9.717537 | 0.000158 | 22.26674 | 22.03711 | 22.18435 |
| Q13823 | Nucleolar G | GNL2 | Nucleolar G | 9.147662 | 0.000388 | 22.53231 | 22.56621 | 22.63462 |
| Q86XI2 | Condensin- | NCAPG2 | Condensin- | 9.834772 | 1.59E-05 | 22.40881 | 22.07032 | 22.40891 |
| Q8N3E9 | 1-phospha | PLCD3 | 1-phospha | 0.972323 | 0.032917 | 22.16036 | 21.25649 | 21.25626 |
| Q9H3N1 | Thioredoxin | TMX1 | Thioredoxin | 9.271438 | 0.00036 | 22.24316 | 22.81652 | 23.19477 |
| O75964 | ATP syntha | ATP5L | ATP syntha | 9.58419 | 5.07E-05 | 22.40476 | 22.32781 | 22.27421 |

| | | | | | | | | |
|---|---|---|---|---|---|---|---|---|
| Q9NTK5 | Obg-like ATI | OLA1 | Obg-like ATI | 9.703692 | 3.18E-05 | 22.10543 | 21.90742 | 22.28074 |
| Q9GZL7 | Ribosome b | WDR12 | Ribosome b | 9.913047 | 0.000212 | 22.57662 | 22.08359 | 22.54745 |
| Q6I9Y2 | THO compl | THOC7 | THO compl | 9.7383 | 5.37E-07 | 23.00477 | 22.76721 | 22.72119 |
| Q4G0J3 | La-related p | LARP7 | La-related p | 5.677186 | 0.080389 | 21.78684 | 21.26957 | 22.27446 |
| P51553 | Isocitrate de | IDH3G | Isocitrate de | 6.210519 | 0.107682 | 23.04772 | 22.70938 | 21.75573 |
| Q12824 | SWI/SNF-re | SMARCB1 | SWI/SNF-re | 9.453735 | 0.009339 | 22.96584 | 22.69579 | 23.03398 |
| Q08554 | Desmocolli | DSC1 | Desmocolli | 1.508385 | 0.198631 | 22.54227 | 21.51485 | 21.90991 |
| P85037 | Forkhead bo | FOXK1 | Forkhead bo | 10.51556 | 9.82E-05 | 22.50601 | 22.47791 | 22.75356 |
| P43307 | Translocon | SSR1 | Translocon | 7.146791 | 0.069168 | 22.53229 | 22.58112 | 23.02773 |
| Q8IY17 | Neuropathy | PNPLA6 | Patatin-like | 10.50213 | 4.16E-05 | 21.9397 | 22.12458 | 21.9239 |
| Q96J01 | THO compl | THOC3 | THO compl | 5.559866 | 0.051808 | 21.00765 | 20.7275 | 23.32361 |
| O75131 | Copine-3 | CPNE3 | Copine-3 O | 10.28856 | 0.000139 | 22.86314 | 22.28159 | 22.36094 |
| O95071 | E3 ubiquitir | UBR5 | E3 ubiquitir | 10.05958 | 0.000421 | 21.81695 | 21.79713 | 21.91236 |
| Q9HCE3 | Zinc finger p | ZNF532 | Zinc finger p | 9.298617 | 0.000139 | 22.03032 | 21.78473 | 21.83156 |
| Q9H6Y2 | WD repeat-c | WDR55 | WD repeat-c | 11.02606 | 1.70E-06 | 23.24844 | 22.82303 | 22.9621 |
| Q15428 | Splicing fac | SF3A2 | Splicing fac | 2.920768 | 0.006816 | 21.02905 | 22.2456 | 22.88172 |
| O75691 | Small subu | UTP20 | Small subu | 9.826877 | 0.000569 | 22.3012 | 22.10712 | 22.36848 |
| Q93100 | Phosphoryl | PHKB | Phosphoryl | 9.533547 | 1.53E-05 | 22.1602 | 22.10396 | 22.40481 |
| P01893;P0 | Putative HL | HLA-H;HLA- | Putative HL | 8.333441 | 0.018244 | 23.09062 | 22.43484 | 22.86034 |
| Q15070 | Mitochondr | OXA1L | Mitochondr | 9.830022 | 0.001489 | 22.5919 | 22.45543 | 22.70487 |
| Q9ULV4 | Coronin-1C | CORO1C | Coronin-1C | 9.858359 | 0.000197 | 21.91922 | 21.98527 | 22.31888 |
| Q9NS86 | LanC-like pr | LANCL2 | LanC-like pr | 8.639658 | 0.000111 | 22.15187 | 22.07349 | 22.31992 |
| Q9NYL2 | Mitogen-act | ZAK | Mitogen-act | 7.515535 | 0.056956 | 21.37279 | 21.78524 | 22.64955 |
| P29034 | Protein S10 | S100A2 | Protein S10 | 5.345189 | 0.129087 | 23.59047 | 21.40255 | 20.92636 |
| Q99755 | Phosphatid | PIP5K1A | Phosphatid | 10.15157 | 1.46E-05 | 22.51203 | 22.34884 | 22.52241 |
| Q86W92 | Liprin-beta- | PPFIBP1 | Liprin-beta- | 9.785125 | 0.000142 | 22.54971 | 22.49025 | 22.15146 |
| P19971 | Thymidine p | TYMP | Thymidine p | 1.896189 | 0.64755 | 20.52984 | 18.55108 | 19.04444 |
| Q9H501 | ESF1 homo | ESF1 | ESF1 homo | 5.640975 | 0.036141 | 21.16693 | 22.74888 | 22.97633 |
| Q8WTU2 | Scavenger r | SSC4D | Scavenger r | 10.02294 | 0.0002 | 22.31775 | 22.36912 | 22.26178 |
| Q9Y678;Q9 | Coatomer s | COPG1;CO | Coatomer s | 6.998354 | 0.01328 | 22.15109 | 20.79122 | 22.18062 |
| Q5VWN6 | Protein FAM | FAM208B | Protein TAS | 9.471367 | 4.10E-05 | 22.44448 | 22.17582 | 22.60869 |
| Q9H9P8 | L-2-hydroxy | L2HGDH | L-2-hydroxy | 9.450478 | 0.000134 | 22.08089 | 21.95234 | 22.18062 |
| Q8TED1 | Probable gl | GPX8 | Probable gl | 8.709001 | 3.00E-06 | 22.6154 | 22.59149 | 22.59689 |
| Q86Y07 | Serine/threo | VRK2 | Serine/threo | 9.444641 | 0.000212 | 22.26699 | 22.38335 | 23.21647 |
| Q2TBE0 | CWF19-like | CWF19L2 | CWF19-like | 4.067337 | 0.000538 | 22.39037 | 23.00215 | 22.95202 |
| Q9BRJ7 | Protein syn | NUDT16L1 | Tudor-intera | 9.537442 | 0.00012 | 22.04963 | 21.61095 | 21.93391 |
| Q9P2J5 | Leucine--tR | LARS | Leucine--tR | 8.250239 | 0.004598 | 21.70329 | 21.84226 | 21.78049 |
| P49406 | 39S ribosor | MRPL19 | Large ribosc | 10.43682 | 0.000311 | 22.37175 | 22.42493 | 22.32232 |
| Q02978 | Mitochondr | SLC25A11 | Mitochondr | 8.884926 | 0.000179 | 22.47224 | 22.36536 | 22.48281 |
| Q9Y2T1 | Axin-2 | AXIN2 | Axin-2 OS=I | 8.372307 | 0.018882 | 22.15174 | 21.32926 | 20.47328 |
| Q9P0J0 | NADH dehy | NDUFA13 | NADH dehy | 10.40209 | 0.000321 | 22.36151 | 22.07235 | 22.39984 |
| Q9NRX2 | 39S ribosor | MRPL17 | Large ribosc | 7.911128 | 0.016175 | 22.36997 | 22.1045 | 22.29782 |
| P31949 | Protein S10 | S100A11 | Protein S10 | 0.169233 | 0.929058 | 22.38609 | 19.80474 | 20.35199 |
| O43491 | Band 4.1-lil | EPB41L2 | Band 4.1-lil | 10.40976 | 3.65E-05 | 22.88114 | 22.4052 | 22.47608 |
| Q6DD88 | Atlastin-3 | ATL3 | Atlastin-3 O | 8.934124 | 0.005685 | 22.18495 | 21.96895 | 22.2064 |
| Q5T8P6 | RNA-bindin | RBM26 | RNA-bindin | 8.093021 | 0.012634 | 21.79313 | 21.8353 | 21.74131 |
| Q16537 | Serine/threo | PPP2R5E | Serine/threo | 9.854322 | 9.98E-05 | 22.42073 | 22.25742 | 22.2856 |
| P19387 | DNA-directe | POLR2C | DNA-directe | 10.12635 | 9.15E-05 | 21.8762 | 22.05069 | 22.33478 |

| | | | | | | | | |
|---|---|---|---|---|---|---|---|---|
| Q9NSE4 | Isoleucine-· | IARS2 | Isoleucine-· | 7.823434 | 0.019004 | 23.06902 | 21.06881 | 22.65283 |
| Q8NDT2 | Putative RN | RBM15B | Putative RN | 9.715923 | 0.000211 | 21.72025 | 22.29041 | 22.94079 |
| Q9Y2J2 | Band 4.1-lil | EPB41L3 | Band 4.1-lil | 3.517207 | 0.017469 | 20.62399 | 23.50567 | 21.03698 |
| Q9UG56 | Phosphatid | PISD | Phosphatid | 6.323308 | 0.044557 | 21.49399 | 22.31562 | 21.3175 |
| Q9UET6 | Putative tRN | FTSJ1 | tRNA (cytidi | 10.46725 | 0.000119 | 23.10639 | 22.87461 | 22.99552 |
| Q9HCM4 | Band 4.1-lil | EPB41L5 | Band 4.1-lil | 9.460307 | 0.000157 | 22.70654 | 22.37451 | 22.49003 |
| Q9NXR1;Q9 | Nuclear dist | NDE1;NDEI | Nuclear dist | 6.132473 | 0.144009 | 21.88993 | 21.77177 | 22.00799 |
| Q86W50 | Methyltrans | METTL16 | RNA N6-ade | 10.10436 | 9.33E-05 | 22.12584 | 22.42954 | 22.43573 |
| O75323 | Protein NipS | GBAS | Protein NipS | 8.104502 | 0.055679 | 23.58176 | 22.97264 | 23.03931 |
| Q9NR12 | PDZ and LIN | PDLIM7 | PDZ and LIN | 7.938667 | 0.049479 | 22.77497 | 21.70729 | 22.37788 |
| Q9Y4A5 | Transforma | TRRAP | Transforma | 6.925329 | 0.040477 | 22.11839 | 21.85045 | 21.14083 |
| Q9NWU5 | 39S ribosor | MRPL22 | Large ribosc | 10.22902 | 3.36E-05 | 22.51374 | 21.23351 | 22.18435 |
| P84101 | Small EDRK | SERF2 | Small EDRK | -0.04308 | 0.976542 | 19.26362 | 21.84839 | 22.20637 |
| Q9UBK9 | Protein UXT | UXT | Protein UXT | 4.766533 | 0.109836 | 22.99405 | 22.83136 | 21.35337 |
| Q96G21 | U3 small nu | IMP4 | U3 small nu | 10.37068 | 0.000234 | 22.57108 | 22.33336 | 22.49877 |
| P62070 | Ras-related | RRAS2 | Ras-related | 7.395791 | 0.073901 | 22.06562 | 22.02908 | 22.00402 |
| Q9Y333 | U6 snRNA-a | LSM2 | U6 snRNA-a | 10.55676 | 9.76E-07 | 22.81141 | 22.38753 | 22.43573 |
| Q99584 | Protein S10 | S100A13 | Protein S10 | 5.250826 | 0.099586 | 22.78578 | 22.59019 | 21.65849 |
| Q9NSD9 | Phenylalan | FARSB | Phenylalan | 7.189547 | 0.023675 | 22.00278 | 20.74271 | 21.6793 |
| Q9UL03 | Integrator c | INTS6 | Integrator c | 7.01316 | 0.044876 | 21.2867 | 21.83356 | 21.60621 |
| Q96E29 | Transcriptio | MTERF3 | Transcriptio | 9.909009 | 4.16E-05 | 22.092 | 21.19376 | 21.91254 |
| Q9NV70 | Exocyst cor | EXOC1 | Exocyst cor | 9.606788 | 0.00032 | 22.36485 | 22.16919 | 22.2548 |
| Q9ULR0 | Pre-mRNA-s | ISY1 | Pre-mRNA-s | 9.988585 | 0.000176 | 22.59063 | 21.91754 | 22.59396 |
| P18669;P1 | Phosphogly | PGAM1;PGA | Phosphogly | 5.288978 | 0.24026 | 21.71871 | 19.55127 | 20.70193 |
| P00403 | Cytochrom | MT-CO2 | Cytochrom | 10.4441 | 9.68E-05 | 23.29123 | 22.59161 | 22.27532 |
| Q86SX3 | Uncharacte | C14orf80 | Tubulin eps | 3.773824 | 0.009978 | 22.06992 | 19.85568 | 22.36983 |
| Q14011 | Cold-induc | CIRBP | Cold-induc | 1.913344 | 0.049066 | 21.14364 | 22.03349 | 22.14436 |
| Q96RL1 | BRCA1-A cc | UIMC1 | BRCA1-A cc | 10.021 | 2.55E-06 | 23.14476 | 22.76743 | 22.98642 |
| O43166 | Signal-indu | SIPA1L1 | Signal-indu | 9.041076 | 1.22E-05 | 21.81006 | 21.48409 | 21.75858 |
| O75419 | Cell divisior | CDC45 | Cell divisior | 9.360379 | 0.001171 | 22.10098 | 21.72013 | 21.97334 |
| Q9NWW5 | Ceroid-lipo | CLN6 | Ceroid-lipo | 10.17434 | 1.29E-06 | 22.53924 | 22.46124 | 22.6262 |
| O14893 | Gem-assoc | GEMIN2 | Gem-assoc | 10.25378 | 2.50E-05 | 22.57015 | 22.97846 | 22.6352 |
| Q96DV4 | 39S ribosor | MRPL38 | Large ribosc | 9.483726 | 7.68E-05 | 22.13192 | 21.98235 | 22.03342 |
| Q9Y2S6;A0 | Translation | TMA7;hCG_ | Translation | -0.10335 | 0.940753 | 20.1916 | 20.68646 | 20.63795 |
| P22735 | Protein-glut | TGM1 | Protein-glut | 1.924527 | 0.608968 | 18.0498 | 16.94912 | 18.46479 |
| Q14517 | Protocadhe | FAT1 | Protocadhe | 7.688453 | 0.005723 | 21.91805 | 21.54042 | 21.7643 |
| Q9Y312 | Protein AAR | AAR2 | Protein AAR | 9.512658 | 2.81E-08 | 22.02499 | 21.86963 | 21.90088 |
| O14929 | Histone ace | HAT1 | Histone ace | 9.560125 | 0.000108 | 22.5127 | 22.41862 | 22.54045 |
| Q9NRX1 | RNA-bindin | PNO1 | RNA-bindin | 6.827632 | 0.068994 | 22.54442 | 21.94785 | 22.8806 |
| O60333 | Kinesin-like | KIF1B | Kinesin-like | 9.446615 | 1.09E-06 | 22.48274 | 22.12193 | 22.36954 |
| Q69YQ0 | Cytospin-A | SPECC1L | Cytospin-A | 8.982059 | 0.000472 | 22.61715 | 22.52581 | 22.98739 |
| Q86X95 | Corepresso | CIR1 | Corepresso | 1.10873 | 0.019656 | 22.18683 | 22.0651 | 22.53846 |
| P00742 | Coagulatior | F10 | Coagulatior | 10.27146 | 0.000534 | 23.08364 | 22.8067 | 22.56183 |
| Q9BZK7 | F-box-like/V | TBL1XR1 | F-box-like/V | 9.938347 | 0.000625 | 22.49658 | 22.30566 | 22.53993 |
| Q6P6C2 | RNA demetl | ALKBH5 | RNA demetl | 9.823785 | 2.06E-05 | 22.46809 | 21.95507 | 21.90962 |
| Q14527 | Helicase-lik | HLTF | Helicase-lik | 7.741367 | 0.044511 | 21.75711 | 21.685 | 21.40655 |
| Q9H2P0 | Activity-dep | ADNP | Activity-dep | 5.69686 | 0.046059 | 21.89167 | 21.85068 | 23.03795 |
| Q13084 | 39S ribosor | MRPL28 | Large ribosc | 11.05493 | 3.53E-05 | 22.43532 | 22.49838 | 22.47202 |

| | | | | | | | | |
|---|---|---|---|---|---|---|---|---|
| Q13206 | Probable A1 | DDX10 | Probable A1 | 7.080075 | 0.05599 | 22.78421 | 20.65167 | 21.06769 |
| Q9Y295 | Developme | DRG1 | Developme | 8.121591 | 0.016219 | 21.8018 | 21.16944 | 22.19126 |
| Q9NRL3 | Striatin-4 | STRN4 | Striatin-4 OS | 2.782722 | 0.434551 | 21.96937 | 21.51658 | 22.10162 |
| Q68D10 | Protein SPT | SPTY2D1 | Protein SPT | 6.513438 | 0.057261 | 21.51417 | 21.71607 | 22.30176 |
| Q7L2J0 | 7SK snRNA | MEPCE | 7SK snRNA | 9.424607 | 3.28E-06 | 22.49038 | 22.54583 | 22.39707 |
| P49841 | Glycogen sy | GSK3B | Glycogen sy | 10.16543 | 1.25E-05 | 23.05644 | 22.6132 | 22.89408 |
| Q9BYD2 | 39S ribosor | MRPL9 | Large ribosc | 9.233949 | 6.96E-06 | 21.77338 | 21.5663 | 21.96487 |
| Q53GQ0 | Very-long-c | HSD17B12 | Very-long-c | 7.608675 | 0.034941 | 22.9155 | 21.52645 | 22.80774 |
| Q16540 | 39S ribosor | MRPL23 | Large ribosc | 9.994809 | 6.15E-07 | 22.36097 | 22.10204 | 22.11432 |
| Q8N122 | Regulatory- | RPTOR | Regulatory- | 8.897599 | 0.004419 | 21.56416 | 21.97246 | 22.00148 |
| Q8WUB8 | PHD finger p | PHF10 | PHD finger p | 9.466009 | 0.000705 | 22.17097 | 22.0444 | 22.13185 |
| Q9Y679 | Ancient ubi | AUP1 | Lipid drople | 9.932011 | 0.000138 | 22.40268 | 22.47816 | 22.52006 |
| Q9UIG0 | Tyrosine-pr | BAZ1B | Tyrosine-pr | 9.954956 | 1.25E-06 | 22.54132 | 22.92727 | 22.5866 |
| P61812 | Transformir | TGFB2 | Transformir | 9.782491 | 0.000356 | 21.94267 | 22.11019 | 21.9693 |
| Q14781 | Chromobo: | CBX2 | Chromobo: | 8.403728 | 2.08E-05 | 21.75328 | 21.61324 | 21.44425 |
| Q8WUF5 | RelA-associ | PPP1R13L | RelA-associ | 10.0506 | 0.000102 | 22.30669 | 22.06148 | 22.01733 |
| Q9NWT8 | Aurora kina: | AURKAIP1 | Small ribos | 9.970205 | 0.000158 | 22.2735 | 21.93571 | 22.50826 |
| E9PAV3;Q1 | Nascent po | NACA | Nascent po | 1.619392 | 0.000995 | 22.25332 | 21.96881 | 21.61979 |
| Q7Z4S6 | Kinesin-like | KIF21A | Kinesin-like | 10.58164 | 0.000172 | 22.67698 | 22.31241 | 22.55119 |
| Q9BS16 | Centromere | CENPK | Centromere | 10.49857 | 0.000207 | 21.91306 | 22.08414 | 22.39116 |
| Q05397 | Focal adhes | PTK2 | Focal adhes | 9.907613 | 0.000936 | 22.9132 | 22.4784 | 22.88097 |
| P49755 | Transmemb | TMED10 | Transmemb | 10.36333 | 0.000239 | 22.84146 | 22.36258 | 22.69 |
| Q96SI9 | Spermatid p | STRBP | Spermatid p | 9.466529 | 9.03E-05 | 22.01907 | 21.64478 | 22.04584 |
| Q9Y448 | Small kinet | KNSTRN | Small kinet | 8.302907 | 0.000202 | 21.97684 | 21.83646 | 22.14398 |
| Q9BT40 | Inositol pol | INPP5K | Inositol pol | 9.687486 | 0.000752 | 22.94288 | 22.33164 | 23.24964 |
| Q86US8 | Telomerase | SMG6 | Telomerase | 9.787203 | 9.08E-07 | 21.6123 | 21.39273 | 21.68457 |
| Q00536;Q0 | Cyclin-depe | CDK16;CDI | Cyclin-depe | 8.967716 | 0.000228 | 22.30547 | 22.50079 | 22.81218 |
| Q9NRW3;Q | DNA dC->d | APOBEC3C | DNA dC->d | 8.452907 | 0.020947 | 21.75475 | 23.52169 | 22.21238 |
| P05455 | Lupus La pr | SSB | Lupus La pr | 7.08277 | 0.036804 | 21.99079 | 23.32965 | 21.82489 |
| Q9UGR2 | Zinc finger C | ZC3H7B | Zinc finger C | 8.545544 | 0.001201 | 21.24621 | 21.03765 | 21.3978 |
| P04350 | Tubulin bet | TUBB4A | Tubulin bet | 9.825274 | 1.54E-07 | 22.16272 | 21.91126 | 22.00134 |
| P62312 | U6 snRNA-a | LSM6 | U6 snRNA-a | 8.379291 | 0.017286 | 22.39976 | 22.18513 | 22.00069 |
| Q9P0K7 | Ankycorbin | RAI14 | Ankycorbin | 10.0289 | 0.000722 | 21.59263 | 21.27708 | 23.59503 |
| Q16891 | MICOS com | IMMT | MICOS com | 9.207046 | 0.001684 | 22.1771 | 20.34423 | 21.62778 |
| Q9HAU4;Q9 | E3 ubiquitir | SMURF2;SN | E3 ubiquitir | 6.343418 | 0.145782 | 21.8187 | 21.64944 | 21.88442 |
| P11166 | Solute carri | SLC2A1 | Solute carri | 7.742282 | 0.009143 | 22.31199 | 21.91791 | 22.13292 |
| Q8WUW1 | Protein BRI( | BRK1 | Protein BRI( | 6.829452 | 0.035077 | 23.13261 | 21.80428 | 21.91441 |
| Q9H7E9 | UPF0488 p | C8orf33 | UPF0488 p | 9.065053 | 9.54E-05 | 21.76681 | 21.76162 | 21.74098 |
| Q9Y608 | Leucine-rich | LRRFIP2 | Leucine-rich | 4.915931 | 0.04225 | 22.81353 | 20.6977 | 22.08105 |
| P07195 | L-lactate de | LDHB | L-lactate de | 5.117661 | 0.00283 | 20.03537 | 22.2383 | 22.3118 |
| Q96GP6 | Scavenger r | SCARF2 | Scavenger r | 9.280642 | 5.76E-05 | 22.10422 | 21.79229 | 22.20018 |
| O15372 | Eukaryotic t | EIF3H | Eukaryotic t | 9.926147 | 1.68E-06 | 22.62511 | 22.1363 | 22.26069 |
| O14682 | Ectoderm-n | ENC1 | Ectoderm-n | 8.992299 | 0.000216 | 22.01754 | 21.93352 | 21.86405 |
| Q96C57 | Uncharacte | C12orf43 | Protein CUS | 8.699973 | 0.000183 | 22.01862 | 21.8839 | 22.02466 |
| O75179 | Ankyrin rep | ANKRD17 | Ankyrin rep | 8.992321 | 4.82E-06 | 22.05811 | 21.9502 | 22.1338 |
| Q14137 | Ribosome b | BOP1 | Ribosome b | 9.040382 | 0.000524 | 22.17919 | 21.6511 | 22.26559 |
| P00387 | NADH-cyto | CYB5R3 | NADH-cyto | 2.373971 | 0.003875 | 22.26003 | 21.99047 | 21.9767 |
| O00142 | Thymidine k | TK2 | Thymidine k | 9.531853 | 7.25E-06 | 22.38795 | 22.50407 | 22.46408 |

| | | | | | | | | |
|---|---|---|---|---|---|---|---|---|
| Q7Z6J6 | FERM doma | FRMD5 | FERM doma | 8.478855 | 0.000287 | 21.39566 | 21.77113 | 21.76482 |
| Q8N983 | 39S riboso | MRPL43 | Large ribos | 9.339837 | 3.89E-05 | 22.04624 | 22.16733 | 22.19982 |
| P08574 | Cytochrom | CYC1 | Cytochrom | 7.231482 | 0.026062 | 22.63008 | 21.59851 | 22.44572 |
| Q9BTC8 | Metastasis- | MTA3 | Metastasis- | 9.956537 | 0.00012 | 22.25376 | 22.4303 | 22.0453 |
| Q9Y613 | FH1/FH2 dc | FHOD1 | FH1/FH2 dc | 9.144927 | 0.000166 | 21.36784 | 22.02168 | 21.67831 |
| P49770 | Translation | EIF2B2 | Translation | 8.94094 | 0.000315 | 21.83395 | 21.88346 | 21.974 |
| O14818;Q8 | Proteasome | PSMA7;PSN | Proteasome | 4.233424 | 0.22725 | 22.52212 | 21.90911 | 22.26662 |
| Q13188 | Serine/threo | STK3 | Serine/threo | 10.56727 | 3.31E-06 | 22.51135 | 22.44208 | 22.84548 |
| O15294 | UDP-N-acet | OGT | UDP-N-acet | 9.030172 | 0.001529 | 21.50577 | 21.84831 | 21.65001 |
| Q8IXM3 | 39S riboso | MRPL41 | Large ribos | 6.363798 | 0.115373 | 21.70881 | 21.49648 | 21.81218 |
| O75251 | NADH dehy | NDUFS7 | NADH dehy | 9.874445 | 2.10E-06 | 22.9036 | 22.41736 | 22.78069 |
| Q9UKV8;Q9 | Protein argo | AGO2;AGO | Protein argo | 9.480938 | 0.000355 | 22.3909 | 22.55483 | 22.57916 |
| P78357 | Contactin-a | CNTNAP1 | Contactin-a | 6.681628 | 0.045434 | 21.26688 | 22.11655 | 21.60765 |
| Q01650 | Large neutra | SLC7A5 | Large neutra | 9.925014 | 8.20E-06 | 22.23599 | 22.16895 | 22.259 |
| Q13427 | Peptidyl-pro | PPIG | Peptidyl-pro | 7.6741 | 0.016957 | 21.59268 | 22.27745 | 22.63806 |
| Q9H9Y6 | DNA-directe | POLR1B | DNA-directe | 9.436454 | 9.35E-05 | 21.89929 | 21.98964 | 22.01195 |
| P14625 | Endoplasm | HSP90B1 | Endoplasm | 4.385578 | 0.185298 | 22.39927 | 20.67401 | 21.11273 |
| P84095 | Rho-related | RHOG | Rho-related | 9.948006 | 0.000707 | 22.5271 | 21.92394 | 22.41661 |
| P31040 | Succinate d | SDHA | Succinate d | 7.335965 | 0.00725 | 21.63059 | 21.47303 | 21.7131 |
| Q86YP4 | Transcriptio | GATAD2A | Transcriptio | 6.218661 | 0.076573 | 21.50814 | 21.35332 | 21.90973 |
| Q8NEW0 | Zinc transpo | SLC30A7 | Zinc transpo | 9.097701 | 0.000155 | 21.98318 | 21.75886 | 21.59231 |
| O00192 | Armadillo re | ARVCF | Splicing reg | 9.527014 | 5.37E-05 | 21.30335 | 21.46458 | 21.75507 |
| P48553 | Trafficking p | TRAPPC10 | Trafficking p | 9.325677 | 9.31E-06 | 21.63989 | 21.51735 | 21.79313 |
| Q96A65 | Exocyst cor | EXOC4 | Exocyst cor | 7.904972 | 2.67E-05 | 21.44496 | 21.23152 | 21.53155 |
| Q5C9Z4 | Nucleolar M | NOM1 | Nucleolar M | 9.063479 | 5.47E-07 | 21.24214 | 21.27628 | 21.46274 |
| Q8N5C6 | S1 RNA-bin | SRBD1 | S1 RNA-bin | 6.707419 | 0.124007 | 21.6817 | 21.39587 | 21.53322 |
| P04083 | Annexin A1 | ANXA1 | Annexin A1 | 3.323375 | 0.329546 | 19.97883 | 19.62087 | 19.5806 |
| P49419 | Alpha-amin | ALDH7A1 | Alpha-amin | 6.643447 | 0.047977 | 21.98663 | 21.25482 | 22.02803 |
| P04632 | Calpain sm | CAPNS1 | Calpain sm | 6.965722 | 0.034614 | 22.21848 | 21.84142 | 21.13314 |
| Q5VV42 | Threonylca | CDKAL1 | Threonylca | 8.927602 | 3.66E-07 | 21.70801 | 21.5122 | 21.60436 |
| Q8TCJ2 | Dolichyl-di | STT3B | Dolichyl-di | 9.742867 | 0.000374 | 22.19141 | 21.80211 | 22.05076 |
| P24534 | Elongation | EEF1B2 | Elongation | 7.487461 | 0.056881 | 22.16941 | 21.37729 | 22.9291 |
| O15554 | Intermediat | KCNN4 | Intermediat | 9.239624 | 0.000275 | 21.67956 | 21.21904 | 21.54383 |
| P49915 | GMP syntha | GMPS | GMP syntha | 9.51986 | 1.92E-05 | 21.9904 | 21.57985 | 22.06441 |
| O95639 | Cleavage ar | CPSF4 | Cleavage ar | 9.639614 | 1.79E-06 | 22.55707 | 22.58614 | 22.3577 |
| O75152 | Zinc finger C | ZC3H11A | Zinc finger C | 6.450953 | 0.046124 | 22.04473 | 20.90006 | 21.31926 |
| Q9P0M9 | 39S riboso | MRPL27 | Large ribos | 8.436291 | 0.042997 | 21.99106 | 22.13615 | 22.38324 |
| Q16555 | Dihydropyri | DPYSL2 | Dihydropyri | 7.649676 | 0.018737 | 22.2108 | 21.85833 | 18.11212 |
| Q9H5V9 | UPF0428 p | CXorf56 | STING ER e | 9.199435 | 3.87E-05 | 22.28238 | 22.25972 | 22.0765 |
| Q14146 | Unhealthy r | URB2 | Unhealthy r | 5.264087 | 0.090147 | 21.22258 | 19.28738 | 21.40426 |
| Q96AB3 | Isochorism | ISOC2 | Isochorism | 8.989196 | 0.000665 | 21.64046 | 21.58889 | 22.04083 |
| Q9BT17 | Mitochondr | MTG1 | Mitochondr | 9.609631 | 6.85E-05 | 21.9421 | 22.14395 | 22.37992 |
| P35613 | Basigin | BSG | Basigin OS= | 6.99042 | 0.042598 | 22.72432 | 21.51989 | 21.85821 |
| Q9H2W6 | 39S riboso | MRPL46 | Large ribos | 2.652412 | 0.008487 | 21.9047 | 20.78205 | 22.4546 |
| Q9NPE3 | H/ACA ribo | NOP10 | H/ACA ribo | 8.247341 | 0.01294 | 20.81433 | 21.93611 | 22.38682 |
| Q8TCT9 | Minor histo | HM13 | Minor histo | 8.854575 | 0.000572 | 21.98325 | 21.77161 | 22.07258 |
| O95218 | Zinc finger F | ZRANB2 | Zinc finger F | 1.909037 | 0.026813 | 22.43288 | 21.02107 | 21.74423 |
| Q56P03 | E2F-associ | EAPP | E2F-associ | 9.878377 | 5.53E-06 | 22.59092 | 22.40681 | 22.37981 |

| | | | | | | | | |
|---|---|---|---|---|---|---|---|---|
| Q13501 | Sequestosc | SQSTM1 | Sequestosc | 9.76134 | 5.51E-06 | 22.3983 | 21.92466 | 22.14087 |
| Q9BVI4 | Nucleolar c | NOC4L | Nucleolar c | 5.840048 | 0.047955 | 20.76668 | 23.52049 | 21.01229 |
| P18887 | DNA repair p | XRCC1 | DNA repair p | 9.82341 | 8.28E-05 | 21.84801 | 21.52784 | 21.96077 |
| Q9HD33 | 39S ribosor | MRPL47 | Large ribosc | 9.499304 | 2.57E-05 | 21.66709 | 21.54109 | 21.63414 |
| P60174 | Triosephos | TPI1 | Triosephos | 5.469438 | 0.205963 | 21.91598 | 19.22296 | 20.09319 |
| P08579 | U2 small nu | SNRPB2 | U2 small nu | 6.515693 | 0.025503 | 21.94689 | 22.03049 | 21.94871 |
| P28066 | Proteasome | PSMA5 | Proteasome | 5.061976 | 0.102163 | 22.52874 | 21.84326 | 21.98558 |
| Q9UHR5 | SAP30-binc | SAP30BP | SAP30-binc | 3.838636 | 0.006126 | 20.36944 | 22.29659 | 22.63995 |
| Q96D53 | AarF domai | ADCK4 | Atypical kin | 7.046374 | 0.014588 | 22.38448 | 20.94757 | 21.95975 |
| Q9BPX6 | Calcium up | MICU1 | Calcium up | 9.984846 | 1.22E-06 | 21.68893 | 21.95982 | 22.01393 |
| Q8N5N7 | 39S ribosor | MRPL50 | Large ribosc | 8.492171 | 0.000424 | 21.66036 | 21.47165 | 22.16858 |
| Q9Y2P8 | RNA 3-term | RCL1 | RNA 3-term | 9.33903 | 2.46E-05 | 22.21653 | 21.85083 | 21.90591 |
| Q01831 | DNA repair p | XPC | DNA repair p | 8.695682 | 0.000429 | 22.04943 | 22.50872 | 22.43695 |
| Q9BTD8 | RNA-bindin | RBM42 | RNA-bindin | 9.902205 | 5.54E-05 | 21.97453 | 22.22591 | 22.22391 |
| P29084 | Transcriptio | GTF2E2 | Transcriptio | 9.573398 | 3.40E-05 | 21.92636 | 21.90239 | 22.41864 |
| O60341 | Lysine-spec | KDM1A | Lysine-spec | 9.647404 | 0.001214 | 20.64764 | 21.84169 | 23.03523 |
| Q92947 | Glutaryl-Co | GCDH | Glutaryl-Co | 7.889003 | 0.029621 | 22.6187 | 22.29398 | 22.43 |
| Q6P1M0 | Long-chain | SLC27A4 | Long-chain | 9.233973 | 9.00E-08 | 21.88666 | 21.94757 | 21.65718 |
| Q8IWR0 | Zinc finger C | ZC3H7A | Zinc finger C | 9.102901 | 9.25E-06 | 21.60014 | 21.675 | 21.74554 |
| Q5SYE7 | NHS-like pr | NHSL1 | NHS-like pr | 8.161452 | 0.013994 | 22.02972 | 21.96371 | 21.79891 |
| O95251 | Histone ace | KAT7 | Histone ace | 9.30955 | 7.47E-05 | 22.33312 | 22.05636 | 22.55848 |
| P22695 | Cytochrom | UQCRC2 | Cytochrom | 10.38884 | 0.001517 | 23.06761 | 21.68616 | 21.49907 |
| P08243 | Asparagine | ASNS | Asparagine | 9.986497 | 0.00016 | 21.89482 | 21.54453 | 21.33865 |
| P38398 | Breast canc | BRCA1 | Breast canc | 9.073326 | 0.000537 | 21.76243 | 21.62658 | 21.74259 |
| P55036 | 26S proteas | PSMD4 | 26S proteas | 10.92049 | 0.000398 | 22.34316 | 22.2573 | 22.2235 |
| Q96T60 | Bifunctiona | PNKP | Bifunctiona | 8.220583 | 0.013972 | 21.56495 | 21.32745 | 21.54854 |
| P24941;Q0 | Cyclin-depe | CDK2;CDK3 | Cyclin-depe | 2.507194 | 0.437659 | 21.27742 | 21.42255 | 21.77028 |
| Q9H9T3 | Elongator c | ELP3 | Elongator c | 9.726683 | 5.88E-06 | 21.11401 | 21.39205 | 21.53986 |
| Q13347 | Eukaryotic t | EIF3I | Eukaryotic t | 9.827335 | 0.000153 | 22.02205 | 21.85293 | 21.73979 |
| P43490 | Nicotinamic | NAMPT | Nicotinamic | 7.162951 | 0.029857 | 22.12594 | 21.21596 | 22.02634 |
| P46736 | Lys-63-spe | BRCC3 | Lys-63-spe | 7.144359 | 0.046889 | 20.04958 | 22.50443 | 22.94963 |
| Q9H6S3 | Epidermal g | EPS8L2 | Epidermal g | 3.112179 | 0.278476 | 21.84203 | 21.12155 | 22.00241 |
| Q96MU7 | YTH domair | YTHDC1 | YTH domair | 9.774563 | 1.78E-05 | 22.16757 | 22.1295 | 22.30435 |
| Q5JVF3 | PCI domair | PCID2 | PCI domair | 8.186022 | 0.000902 | 21.23836 | 21.54345 | 21.89848 |
| Q96S15 | WD repeat-c | WDR24 | GATOR2 co | 9.129121 | 3.39E-06 | 21.40728 | 21.41095 | 21.13865 |
| O95159 | Zinc finger p | ZFPL1 | Zinc finger p | 8.469475 | 0.00272 | 21.67698 | 21.37671 | 21.60608 |
| P61803 | Dolichyl-dip | DAD1 | Dolichyl-dip | 6.714603 | 0.044553 | 21.45608 | 20.8423 | 21.97519 |
| P40926 | Malate dehy | MDH2 | Malate dehy | 6.542152 | 0.122312 | 21.77129 | 20.05947 | 22.36215 |
| Q969X5 | Endoplasm | ERGIC1 | Endoplasm | 8.022011 | 0.055365 | 21.65836 | 20.86884 | 22.3497 |
| Q9NZ63 | Uncharacte | C9orf78 | Splicing fac | 8.499869 | 0.000836 | 21.86121 | 21.51234 | 21.92749 |
| Q9BW27 | Nuclear por | NUP85 | Nuclear por | 8.978564 | 0.000588 | 21.46742 | 21.46254 | 21.6454 |
| P09429;B2 | High mobili | HMGB1;HM | High mobili | 0.461037 | 0.604766 | 21.18665 | 21.23491 | 21.58187 |
| Q8N6R0 | Methyltrans | METTL13 | eEF1A lysin | 8.681259 | 0.000118 | 22.22511 | 21.83511 | 21.9365 |
| Q99959 | Plakophilin | PKP2 | Plakophilin | 9.707666 | 0.000786 | 22.06378 | 21.72092 | 21.95504 |
| Q8TEU7 | Rap guanin | RAPGEF6 | Rap guanin | 8.752279 | 0.000217 | 21.19587 | 21.31772 | 21.51075 |
| Q9BXY0 | Protein MAK | MAK16 | Protein MAK | 6.493281 | 0.093172 | 22.19406 | 21.93991 | 21.94464 |
| Q96GA3 | Protein LTV | LTV1 | Protein LTV | 10.27941 | 7.79E-05 | 22.3149 | 22.42306 | 21.92926 |
| Q13889 | General trar | GTF2H3 | General trar | 9.627288 | 5.13E-05 | 22.35116 | 21.96962 | 22.40463 |

| | | | | | | | | |
|---|---|---|---|---|---|---|---|---|
| P78549 | Endonuclea | NTHL1 | Endonuclea | 8.990628 | 1.98E-05 | 21.82116 | 21.49082 | 21.91339 |
| Q96N66 | Lysophosp | MBOAT7 | Lysophosp | 8.882783 | 3.92E-05 | 21.60567 | 21.47194 | 21.90866 |
| Q29RF7 | Sister chron | PDS5A | Sister chron | 8.341691 | 0.028101 | 22.16916 | 21.06579 | 22.14638 |
| P61224;P6 | Ras-related | RAP1B;RAP | Ras-related | 7.529701 | 0.039425 | 21.07627 | 21.87658 | 23.02012 |
| Q96T88 | E3 ubiquitir | UHRF1 | E3 ubiquitir | 9.520694 | 2.29E-05 | 21.88904 | 21.36512 | 21.8753 |
| O96028 | Histone-lys | WHSC1 | Histone-lys | 9.544493 | 8.89E-08 | 21.68979 | 21.58049 | 21.78796 |
| Q8WVX9 | Fatty acyl-C | FAR1 | Fatty acyl-C | 8.810213 | 0.000292 | 21.73318 | 21.48773 | 21.73053 |
| O94921 | Cyclin-depe | CDK14 | Cyclin-depe | 8.410962 | 0.000326 | 21.59427 | 21.75083 | 21.81026 |
| Q8IX18 | Probable A | DHX40 | Probable A | 8.979715 | 0.00047 | 21.62574 | 21.28529 | 21.60241 |
| Q8NB90 | Spermatoge | SPATA5 | ATPase fam | 9.820562 | 4.81E-05 | 21.94041 | 21.8994 | 21.95411 |
| Q92621 | Nuclear por | NUP205 | Nuclear por | 8.706651 | 0.000248 | 21.53445 | 21.36292 | 21.30084 |
| Q8TAD8 | Smad nucle | SNIP1 | Smad nucle | 4.546164 | 0.007481 | 19.67525 | 22.14348 | 22.11191 |
| P61026 | Ras-related | RAB10 | Ras-related | 7.273333 | 0.061663 | 21.6068 | 21.63005 | 21.92723 |
| Q13618 | Cullin-3 | CUL3 | Cullin-3 OS | 7.657704 | 0.00021 | 21.82034 | 21.37369 | 21.71356 |
| P61020 | Ras-related | RAB5B | Ras-related | 6.445798 | 0.067705 | 21.14239 | 21.26757 | 21.0912 |
| Q5QJE6 | Deoxynucle | DNTTIP2 | Deoxynucle | 9.700649 | 0.0002 | 22.13129 | 22.12525 | 22.0439 |
| O60573 | Eukaryotic t | EIF4E2 | Eukaryotic t | 9.748229 | 0.000152 | 21.88267 | 21.3476 | 21.72883 |
| P54132 | Bloom synd | BLM | RecQ-like D | 9.725781 | 2.24E-05 | 21.16288 | 21.15016 | 21.0747 |
| O43815 | Striatin | STRN | Striatin OS= | 8.889673 | 0.001337 | 22.12461 | 21.66058 | 22.14703 |
| P49006 | MARCKS-re | MARCKSL1 | MARCKS-re | 9.169958 | 2.23E-05 | 22.07669 | 21.97106 | 21.76077 |
| O00231 | 26S proteas | PSMD11 | 26S proteas | 9.849392 | 6.55E-07 | 22.01052 | 21.70708 | 21.61221 |
| P26358 | DNA (cytosi | DNMT1 | DNA (cytosi | 8.01152 | 1.14E-05 | 21.23946 | 20.99769 | 21.30519 |
| Q15828 | Cystatin-M | CST6 | Cystatin-M ( | 5.679331 | 0.15118 | 22.59996 | 19.34297 | 20.25084 |
| Q9UPN6;O | Protein SCA | SCAF8;SCA | SR-related a | 8.601511 | 6.95E-05 | 21.76681 | 21.43639 | 21.65923 |
| Q6NZY4 | Zinc finger C | ZCCHC8 | Zinc finger C | 8.44331 | 6.23E-05 | 21.1858 | 21.59914 | 21.08161 |
| Q9Y6W5 | Wiskott-Ald | WASF2 | Actin-bindir | 7.164235 | 0.030652 | 22.17222 | 21.29502 | 22.01499 |
| Q96A35 | 39S ribosor | MRPL24 | Large ribosc | 9.436605 | 3.99E-06 | 22.08534 | 22.16748 | 22.2125 |
| O14974 | Protein pho | PPP1R12A | Protein pho | 9.036507 | 5.60E-05 | 21.47447 | 21.67805 | 21.79539 |
| A6NFI3 | Zinc finger p | ZNF316 | Zinc finger p | 9.547481 | 0.000233 | 21.42578 | 21.5172 | 21.58275 |
| Q8N2M8 | CLK4-assoc | CLASRP | CLK4-assoc | 8.256393 | 6.13E-05 | 21.67887 | 21.50315 | 21.77137 |
| Q9UFW8 | CGG triplet | CGGBP1 | CGG triplet | 9.678306 | 0.000103 | 21.82454 | 21.35933 | 22.12196 |
| Q9Y324 | rRNA-proce | FCF1 | rRNA-proce | 4.711078 | 0.035203 | 22.59605 | 20.20407 | 20.19268 |
| Q96L91 | E1A-binding | EP400 | E1A-binding | 0.613103 | 0.053623 | 21.38036 | 20.60168 | 21.02378 |
| Q8IZ69 | tRNA (uracil | TRMT2A | tRNA (uracil | 8.987748 | 0.000677 | 21.42368 | 21.87406 | 21.92528 |
| Q9BVJ6;Q5 | U3 small nu | UTP14A;UT | U3 small nu | 5.996482 | 0.051912 | 20.70143 | 21.07698 | 22.08018 |
| Q9HCG8 | Pre-mRNA-s | CWC22 | Pre-mRNA-s | 8.685723 | 3.69E-05 | 21.78349 | 21.89545 | 21.89527 |
| Q14331 | Protein FRG | FRG1 | Protein FRG | 2.628772 | 0.073501 | 21.98739 | 20.15869 | 22.29404 |
| Q14739 | Lamin-B rec | LBR | Delta(14)-st | 9.051781 | 0.001707 | 21.80994 | 21.46344 | 21.72092 |
| Q9BXF6 | Rab11 fami | RAB11FIP5 | Rab11 fami | 9.49175 | 0.001384 | 22.17515 | 19.6012 | 22.13176 |
| Q15629 | Translocatir | TRAM1 | Translocatir | 6.89115 | 0.044029 | 21.40385 | 21.18707 | 22.50125 |
| Q9BPX3 | Condensin | NCAPG | Condensin | 9.609474 | 5.61E-05 | 20.85095 | 21.97687 | 22.2122 |
| Q9NVN8 | Guanine nu | GNL3L | Guanine nu | 4.630624 | 0.173893 | 20.50329 | 20.57718 | 22.20497 |
| Q63HN8 | E3 ubiquitir | RNF213 | E3 ubiquitir | 8.091393 | 0.000109 | 20.90095 | 21.40083 | 20.83245 |
| P61604 | 10 kDa hea | HSPE1 | 10 kDa hea | 3.881215 | 0.221249 | 21.56973 | 21.15481 | 21.29266 |
| P67775;P6 | Serine/threc | PPP2CA;PP | Serine/threc | 9.664746 | 0.000449 | 22.54614 | 21.93463 | 20.3926 |
| O00165 | HCLS1-ass | HAX1 | HCLS1-ass | 6.953714 | 0.013165 | 21.3218 | 21.56286 | 21.62154 |
| Q16543 | Hsp90 co-c | CDC37 | Hsp90 co-c | 9.87076 | 5.16E-05 | 21.80562 | 22.35407 | 21.68876 |
| Q9H7H0 | Methyltrans | METTL17 | Methyltrans | 9.508073 | 0.000437 | 21.14096 | 20.97953 | 21.80499 |

| | | | | | | | | |
|---|---|---|---|---|---|---|---|---|
| P28370 | Probable gl | SMARCA1 | Probable gl | 9.202824 | 0.000584 | 21.69969 | 21.45964 | 21.98075 |
| O43324 | Eukaryotic t | EEF1E1 | Eukaryotic t | 9.372207 | 0.000235 | 21.47927 | 21.38211 | 21.69621 |
| Q9UQ80 | Proliferatior | PA2G4 | Proliferatior | 8.521549 | 8.60E-05 | 21.97141 | 21.71641 | 21.4496 |
| Q9NYB0 | Telomeric re | TERF2IP | Telomeric re | 9.080371 | 0.000104 | 21.64891 | 21.84957 | 22.27943 |
| P60981 | Destrin | DSTN | Destrin OS= | 9.842023 | 0.000294 | 22.58522 | 21.94424 | 22.1756 |
| Q01105;PC | Protein SET | SET;SETSIF | Protein SET | 7.14468 | 0.05989 | 21.53084 | 21.71158 | 21.6553 |
| Q69YN2 | CWF19-like | CWF19L1 | CWF19-like | 9.057704 | 5.25E-05 | 21.31424 | 21.35756 | 21.53308 |
| Q6P1K8;Q1 | General trar | GTF2H2C;G | General trar | 9.208736 | 0.000187 | 21.94306 | 21.85456 | 22.14345 |
| Q6IBW4 | Condensin- | NCAPH2 | Condensin- | 6.980524 | 0.06808 | 21.434 | 21.42152 | 21.94063 |
| Q9Y450 | HBS1-like p | HBS1L | HBS1-like p | 8.291972 | 0.00134 | 20.38015 | 21.69247 | 21.77905 |
| O94913 | Pre-mRNA c | PCF11 | Pre-mRNA c | 8.391147 | 0.004356 | 21.82229 | 21.2257 | 21.23894 |
| Q6ZUT1 | Uncharacte | C11orf57 | Uncharacte | 9.304796 | 0.000449 | 21.72422 | 21.57154 | 22.07571 |
| Q15650 | Activating si | TRIP4 | Activating si | 10.24813 | 2.57E-05 | 21.95992 | 21.72941 | 22.2615 |
| Q0ZGT2 | Nexilin | NEXN | Nexilin OS= | 9.919538 | 0.00066 | 21.06677 | 21.30803 | 21.94682 |
| P40616 | ADP-ribosyl | ARL1 | ADP-ribosyl | 8.993266 | 0.000136 | 21.90547 | 21.92172 | 22.028 |
| P52294;O6 | Importin su | KPNA1;KPN | Importin su | 9.244109 | 0.000264 | 21.45989 | 19.56897 | 21.6214 |
| P18754 | Regulator o | RCC1 | Regulator o | 8.540008 | 7.95E-06 | 21.72854 | 21.73566 | 21.56328 |
| Q969V3 | Nicalin | NCLN | BOS compl | 8.569779 | 0.001443 | 21.65242 | 21.73682 | 21.70261 |
| A2RUS2 | DENN dom | DENND3 | DENN dom | 9.061928 | 0.000625 | 21.66458 | 21.89393 | 22.05504 |
| O14744 | Protein argi | PRMT5 | Protein argi | 7.389197 | 0.010454 | 20.87004 | 21.96409 | 21.99621 |
| P52756 | RNA-bindin | RBM5 | RNA-bindin | 8.651585 | 1.35E-06 | 21.50741 | 21.64606 | 21.85064 |
| P30260 | Cell divisior | CDC27 | Cell divisior | 9.534803 | 0.000265 | 21.29793 | 21.16478 | 21.56746 |
| Q9Y6N5 | Sulfide:quir | SQRDL | Sulfide:quir | 7.108478 | 0.050391 | 22.81689 | 21.68162 | 20.95608 |
| Q96EP5 | DAZ-associ | DAZAP1 | DAZ-associ | 8.999406 | 0.001005 | 21.47571 | 21.5092 | 21.90463 |
| O00471 | Exocyst cor | EXOC5 | Exocyst cor | 9.059756 | 5.76E-05 | 22.13182 | 21.42537 | 21.4246 |
| Q14232 | Translation | EIF2B1 | Translation | 7.448419 | 0.000279 | 21.54165 | 21.15035 | 21.44844 |
| Q9NW08 | DNA-directe | POLR3B | DNA-directe | 9.901167 | 0.000927 | 21.36116 | 21.37756 | 21.70573 |
| Q14197 | Peptidyl-tRI | ICT1 | Large ribosc | 8.997485 | 0.000199 | 21.40291 | 21.98569 | 21.52373 |
| Q16658 | Fascin | FSCN1 | Fascin OS=I | 6.777929 | 0.047487 | 22.11674 | 21.01304 | 21.03134 |
| Q9Y5T5 | Ubiquitin ca | USP16 | Ubiquitin ca | 9.358069 | 9.95E-05 | 21.83387 | 20.82182 | 21.35628 |
| Q07352 | Zinc finger p | ZFP36L1 | mRNA deca | 9.377437 | 4.24E-05 | 21.98524 | 21.79213 | 22.05864 |
| O60437 | Periplakin | PPL | Periplakin C | 4.571307 | 0.214865 | 20.69209 | 21.24841 | 19.85567 |
| P23526 | Adenosylhc | AHCY | Adenosylhc | 4.379143 | 0.061497 | 22.63898 | 20.90029 | 20.88889 |
| P52435;Q9 | DNA-directe | POLR2J;POI | DNA-directe | 9.108621 | 4.77E-05 | 21.96684 | 22.04193 | 22.16987 |
| P13073 | Cytochrom | COX4I1 | Cytochrom | 4.937771 | 0.06508 | 21.97477 | 20.23783 | 21.52013 |
| P51531 | Probable gl | SMARCA2 | Probable gl | 9.071538 | 0.000386 | 21.70771 | 21.40442 | 21.53403 |
| Q8WYP5 | Protein ELY! | AHCTF1 | Protein ELY! | 9.188815 | 0.002013 | 21.53659 | 21.31091 | 21.57122 |
| Q00534 | Cyclin-depe | CDK6 | Cyclin-depe | 9.840184 | 2.56E-05 | 21.82159 | 21.66345 | 21.74485 |
| P49790 | Nuclear por | NUP153 | Nuclear por | 9.152752 | 0.002556 | 21.82974 | 21.82943 | 21.7099 |
| P25786 | Proteasome | PSMA1 | Proteasome | 4.298746 | 0.229034 | 22.20514 | 20.69439 | 20.2674 |
| Q9NUQ8 | ATP-binding | ABCF3 | ATP-binding | 5.599879 | 0.045835 | 21.53735 | 20.61189 | 20.56532 |
| Q13415 | Origin recog | ORC1 | Origin recog | 8.554999 | 5.53E-05 | 21.23105 | 21.11044 | 21.31755 |
| Q15022 | Polycomb p | SUZ12 | Polycomb p | 9.36255 | 5.40E-05 | 21.12345 | 21.33805 | 21.99714 |
| Q9Y5S2;Q5 | Serine/threc | CDC42BPB | Serine/threc | 7.238554 | 0.047102 | 21.06428 | 22.18459 | 21.09603 |
| Q96NE9 | FERM doma | FRMD6 | FERM doma | 7.77203 | 2.83E-07 | 21.20658 | 21.11006 | 21.29412 |
| Q13190 | Syntaxin-5 | STX5 | Syntaxin-5 ( | 6.684825 | 0.065409 | 20.4802 | 21.88907 | 21.24458 |
| Q9H814 | Phosphoryl | PHAX | Phosphoryl | 8.462233 | 0.004103 | 21.43781 | 19.43971 | 21.74489 |
| Q8IVF7 | Formin-like | FMNL3 | Formin-like | 8.141868 | 0.000454 | 20.9512 | 20.43868 | 20.81738 |

| | | | | | | | | |
|---|---|---|---|---|---|---|---|---|
| Q13637;O1 | Ras-related | RAB32;RAB | Ras-related | 8.077353 | 2.86E-05 | 21.48925 | 21.2204 | 21.15295 |
| Q9NYY8 | FAST kinase | FASTKD2 | FAST kinase | 8.755575 | 0.000377 | 21.50824 | 21.44253 | 22.15626 |
| P49821 | NADH dehy | NDUFV1 | NADH dehy | 9.150454 | 0.000525 | 21.38395 | 21.37236 | 21.64161 |
| P04183 | Thymidine I | TK1 | Thymidine I | 8.279553 | 0.001872 | 21.12723 | 21.25908 | 21.38158 |
| Q9NR31;Q9 | GTP-binding | SAR1A;SAR | GTP-binding | 6.82208 | 0.030835 | 21.74952 | 21.80821 | 21.56611 |
| Q6DT37 | Serine/threo | CDC42BPG | Serine/threo | 9.921159 | 7.25E-06 | 22.15795 | 21.56295 | 21.77864 |
| Q92759 | General trar | GTF2H4 | General trar | 9.794023 | 1.18E-05 | 21.62876 | 21.92267 | 22.11924 |
| Q96H22 | Centromere | CENPN | Centromere | 8.231862 | 0.000177 | 20.93359 | 21.17713 | 21.76466 |
| Q9UHD2 | Serine/threo | TBK1 | Serine/threo | 9.096448 | 8.22E-06 | 21.00573 | 21.12402 | 21.45819 |
| P78345 | Ribonuclea | RPP38 | Ribonuclea | 9.245879 | 4.16E-06 | 21.80227 | 21.43064 | 21.78277 |
| Q9UKX7 | Nuclear por | NUP50 | Nuclear por | 9.125018 | 0.000651 | 21.30725 | 21.38548 | 21.59778 |
| Q9H7D7 | WD repeat-c | WDR26 | WD repeat-c | 8.697777 | 2.04E-05 | 21.62029 | 21.63174 | 21.91696 |
| Q13356 | Peptidyl-pro | PPIL2 | RING-type E | 8.680114 | 0.000123 | 21.62694 | 21.08952 | 21.6141 |
| O95985;Q1 | DNA topois | TOP3B;TOP | DNA topois | 8.46605 | 0.001268 | 21.24452 | 21.1832 | 21.39441 |
| Q6ICH7 | Aspartate b | ASPHD2 | Aspartate b | 9.674178 | 5.26E-08 | 21.60829 | 21.56904 | 21.61122 |
| O43617 | Trafficking p | TRAPPC3 | Trafficking p | 8.850872 | 0.000424 | 22.17185 | 21.60621 | 20.57191 |
| P49748 | Very long-cl | ACADVL | Very long-cl | 9.001012 | 0.000355 | 22.39488 | 21.68676 | 19.91917 |
| Q9Y5E2;Q9 | Protocadhe | PCDHB7;PC | Protocadhe | 8.576849 | 3.99E-06 | 21.83005 | 21.80959 | 22.02584 |
| P51648 | Fatty aldehy | ALDH3A2 | Aldehyde d | 8.649807 | 3.96E-05 | 21.77656 | 21.28782 | 21.85205 |
| P01111;P0 | GTPase NRa | NRAS;KRAS | GTPase NRa | 7.160751 | 0.028836 | 21.67908 | 20.68919 | 21.90771 |
| P01009 | Alpha-1-ant | SERPINA1 | Alpha-1-ant | 5.329328 | 0.17947 | 21.90014 | 21.94424 | 19.68043 |
| O00487 | 26S proteas | PSMD14 | 26S proteas | 8.613715 | 0.001348 | 20.97407 | 21.27435 | 21.76081 |
| Q9Y294;Q9 | Histone cha | ASF1A;ASF | Histone cha | 5.820358 | 0.07652 | 21.55712 | 21.40182 | 22.02763 |
| Q9NQ50 | 39S riboson | MRPL40 | Large riboso | 6.2752 | 0.044815 | 21.55309 | 20.79899 | 20.66735 |
| Q8NI77 | Kinesin-like | KIF18A | Kinesin-like | 8.559046 | 0.00194 | 20.59578 | 21.50785 | 21.61221 |
| Q9UPN3;Q9 | Microtubule | MACF1;KIA | Microtubule | 8.363998 | 0.000277 | 21.39205 | 20.73769 | 21.49609 |
| P25311 | Zinc-alpha-2 | AZGP1 | Zinc-alpha-2 | 2.442067 | 0.488639 | 18.74678 | 17.37662 | 17.63105 |
| Q8N5L8 | Ribonuclea | RPP25L | Ribonuclea | 9.487107 | 7.24E-05 | 21.77169 | 20.3252 | 21.88372 |
| P20290 | Transcriptio | BTF3 | Transcriptio | 9.313003 | 5.26E-07 | 21.78891 | 21.46767 | 21.71297 |
| O15027 | Protein tran | SEC16A | Protein tran | 9.024612 | 0.000303 | 20.91437 | 21.30257 | 21.44536 |
| Q9NXR7 | BRCA1-A co | BRE | BRISC and I | 8.634128 | 0.000543 | 21.37608 | 21.07659 | 21.58481 |
| Q16342 | Programme | PDCD2 | Programme | 9.368418 | 2.62E-05 | 21.04507 | 20.91437 | 21.16521 |
| Q13131;P5 | 5-AMP-activ | PRKAA1;PR | 5-AMP-activ | 9.558056 | 0.000831 | 21.93442 | 21.89519 | 21.79388 |
| Q8IXI1;Q8I | Mitochondr | RHOT2;RHO | Mitochondr | 8.948443 | 0.000819 | 20.90852 | 21.08848 | 21.35912 |
| P78332 | RNA-bindin | RBM6 | RNA-bindin | 9.4844 | 0.000221 | 21.60359 | 21.47249 | 21.81691 |
| O75923 | Dysferlin | DYSF | Dysferlin OS | 8.103642 | 1.27E-05 | 20.6472 | 20.16864 | 20.42562 |
| P22102 | Trifunctiona | GART | Trifunctiona | 8.519014 | 0.000145 | 21.45357 | 21.44844 | 21.57159 |
| O15460 | Prolyl 4-hyd | P4HA2 | Prolyl 4-hyd | 9.126619 | 0.000491 | 22.08916 | 21.94016 | 19.39425 |
| Q9H2G4 | Testis-speci | TSPYL2 | Testis-speci | 9.276186 | 6.85E-05 | 21.41266 | 21.20383 | 21.28506 |
| P01023;P2 | Alpha-2-ma | A2M;PZP | Alpha-2-ma | 6.653353 | 0.106942 | 21.52688 | 22.67988 | 20.0541 |
| P10599 | Thioredoxin | TXN | Thioredoxin | 4.998852 | 0.169935 | 22.02844 | 20.51133 | 19.88039 |
| P18206 | Vinculin | VCL | Vinculin OS | 7.457937 | 0.031542 | 21.75711 | 21.67655 | 21.8617 |
| O95983 | Methyl-CpG | MBD3 | Methyl-CpG | 5.248509 | 0.090626 | 20.4493 | 20.41782 | 20.97855 |
| Q07866;Q9 | Kinesin ligh | KLC1;KLC2 | Kinesin ligh | 8.827717 | 0.000196 | 21.2752 | 21.52984 | 21.60996 |
| Q5ST30 | Valine--tRNA | VARS2 | Valine--tRNA | 9.460199 | 0.000311 | 21.40572 | 21.41746 | 21.2731 |
| Q92674 | Centromere | CENPI | Centromere | 9.147199 | 9.17E-05 | 21.31479 | 21.43843 | 21.53308 |
| Q53H12 | Acylglycero | AGK | Acylglycero | 7.608298 | 0.027916 | 20.85072 | 20.31915 | 22.23283 |
| Q8N428 | Polypeptide | GALNT16 | Polypeptide | 9.133719 | 2.98E-06 | 21.8567 | 21.9235 | 21.72167 |

| | | | | | | | | |
|---|---|---|---|---|---|---|---|---|
| Q9UDY4 | DnaJ homo | DNAJB4 | DnaJ homo | 8.280659 | 0.001 | 20.96219 | 20.82625 | 21.12811 |
| Q9H5Q4 | Dimethylad | TFB2M | Dimethylad | 8.626551 | 0.000196 | 21.36121 | 21.08499 | 20.92687 |
| P26640 | Valine--tRN | VARS | Valine--tRN | 8.200469 | 0.000448 | 21.58252 | 21.11152 | 21.08557 |
| Q7Z333 | Probable he | SETX | Probable he | 8.195623 | 0.000798 | 21.0831 | 21.45161 | 21.03483 |
| P54803 | Galactocere | GALC | Galactocere | 9.391887 | 8.60E-06 | 20.94149 | 20.76442 | 21.3133 |
| Q96HQ2 | CDKN2AIP | CDKN2AIPI | CDKN2AIP | 9.072107 | 0.000303 | 21.77008 | 21.63834 | 21.81589 |
| Q9NWU2 | Glucose-inc | GID8 | Glucose-inc | 7.590603 | 0.000245 | 20.79162 | 20.97183 | 21.0792 |
| Q9H9L3 | Interferon-s | ISG20L2 | Interferon-s | 8.309185 | 1.24E-07 | 21.34873 | 21.19532 | 21.52215 |
| Q9Y2Z4 | Tyrosine--tF | YARS2 | Tyrosine--tF | 7.66493 | 0.000771 | 21.39273 | 21.74045 | 21.33129 |
| Q96RT8 | Gamma-tub | TUBGCP5 | Gamma-tub | 9.393447 | 0.000116 | 21.44647 | 21.28139 | 21.52703 |
| Q5FWF4 | DNA anneal | ZRANB3 | DNA anneal | 8.775529 | 0.002226 | 21.19556 | 21.17493 | 21.55065 |
| Q9H1I8 | Activating si | ASCC2 | Activating si | 8.063843 | 0.00048 | 20.96564 | 21.11927 | 21.06046 |
| O00299 | Chloride int | CLIC1 | Chloride int | 6.263105 | 0.086201 | 21.57787 | 20.47516 | 20.39916 |
| Q15021 | Condensin | NCAPD2 | Condensin | 9.095539 | 0.000546 | 21.55136 | 21.41302 | 21.84915 |
| Q9H9J2 | 39S ribosor | MRPL44 | Large ribosc | 8.272418 | 0.00097 | 20.98912 | 20.84904 | 21.34863 |
| O95163 | Elongator c | IKBKAP | Elongator c | 10.10998 | 1.67E-05 | 21.65727 | 21.62827 | 21.39692 |
| Q96QA5 | Gasdermin- | GSDMA | Gasdermin- | 5.2905 | 0.126309 | 20.79074 | 21.37475 | 19.93956 |
| Q8N442 | Translation | GUF1 | Translation | 8.711143 | 6.02E-05 | 20.7666 | 20.4802 | 20.80813 |
| Q14318 | Peptidyl-pro | FKBP8 | Peptidyl-pro | 7.423833 | 1.31E-05 | 21.09648 | 21.05993 | 21.28749 |
| Q8N1G4 | Leucine-rich | LRRC47 | Leucine-rich | 8.739627 | 0.000203 | 21.61643 | 21.39142 | 21.5275 |
| Q9Y618 | Nuclear rec | NCOR2 | Nuclear rec | 8.461074 | 0.000465 | 19.89906 | 21.98978 | 21.7984 |
| P23193 | Transcriptio | TCEA1 | Transcriptio | 9.098898 | 0.00063 | 20.84652 | 20.78963 | 21.31838 |
| O43824 | Putative GT | GTPBP6 | Putative GT | 2.116865 | 0.017743 | 21.66353 | 20.07966 | 20.12035 |
| P53992 | Protein tran | SEC24C | Protein tran | 8.644466 | 1.02E-06 | 21.15487 | 20.73892 | 20.83823 |
| P38606 | V-type prot | ATP6V1A | V-type prot | 5.623251 | 0.072538 | 21.0476 | 20.08356 | 20.23935 |
| Q6P1X5 | Transcriptio | TAF2 | Transcriptio | 8.080372 | 0.004326 | 21.58008 | 20.75173 | 20.01141 |
| O43374;C9 | Ras GTPase | RASA4;RAS | Ras GTPase | 7.86246 | 0.00054 | 20.83561 | 20.75605 | 20.2867 |
| P30101 | Protein disu | PDIA3 | Protein disu | 9.050549 | 5.02E-05 | 20.8012 | 20.68372 | 21.00751 |
| Q9NXE8 | Pre-mRNA-s | CWC25 | Pre-mRNA-s | 6.099159 | 0.025522 | 21.26196 | 21.28484 | 21.43258 |
| P37268 | Squalene sy | FDFT1 | Squalene sy | 5.760453 | 0.0353 | 19.68891 | 20.87388 | 20.93179 |
| Q155Q3 | Dixin | DIXDC1 | Dixin OS=H | 7.921575 | 1.94E-05 | 20.76765 | 20.69948 | 20.18362 |
| P51911 | Calponin-1 | CNN1 | Calponin-1 | 7.704211 | 0.070791 | 20.42808 | 20.2018 | 21.56649 |
| Q9UJ70 | N-acetyl-D-g | NAGK | N-acetyl-D-g | 6.934174 | 0.080947 | 20.52908 | 20.04727 | 21.01414 |
| Q8TEX9 | Importin-4 | IPO4 | Importin-4 ( | 9.288873 | 0.000727 | 20.59923 | 20.64059 | 20.84934 |
| P07476 | Involucrin | IVL | Involucrin ( | 4.506853 | 0.142626 | 20.65953 | 19.82831 | 20.02972 |
| Q9Y490 | Talin-1 | TLN1 | Talin-1 OS= | 5.844932 | 0.042032 | 21.36196 | 20.01414 | 20.33451 |
| P55196 | Afadin | MLLT4 | Afadin OS=I | 8.318289 | 8.02E-05 | 20.1928 | 19.50204 | 20.89948 |
| Q460N5 | Poly [ADP-r | PARP14 | Protein mor | 7.745459 | 0.000992 | 20.68209 | 20.35982 | 21.27509 |
| O00232 | 26S proteas | PSMD12 | 26S proteas | 7.324118 | 0.002301 | 20.53991 | 20.25014 | 20.44869 |
| P55072 | Transitional | VCP | Transitional | 2.906358 | 0.277237 | 20.86509 | 20.52133 | 20.48434 |
| O43818 | U3 small nu | RRP9 | U3 small nu | 4.143977 | 0.063279 | 21.64337 | 20.24598 | 20.30586 |
| Q13442 | 28 kDa heat | PDAP1 | 28 kDa heat | 2.303058 | 0.407948 | 19.32435 | 19.38429 | 20.03187 |
| Q8WY36 | HMG box tra | BBX | HMG box tra | 7.433349 | 4.81E-06 | 20.84345 | 20.56374 | 21.05973 |
| Q96P63 | Serpin B12 | SERPINB12 | Serpin B12 | 5.0092 | 0.157251 | 20.23409 | 20.59978 | 20.5838 |
| Q9NUD5 | Zinc finger ( | ZCCHC3 | Zinc finger ( | 7.3532 | 0.000647 | 20.05091 | 21.9911 | 21.58178 |
| Q9UPT5 | Exocyst cor | EXOC7 | Exocyst cor | 7.651727 | 0.00082 | 20.42196 | 21.17323 | 21.10764 |
| Q86TB9 | Protein PAT | PATL1 | Protein PAT | 7.77013 | 1.14E-05 | 20.0442 | 20.47605 | 20.75279 |
| P46020 | Phosphoryl | PHKA1 | Phosphoryl | 7.801389 | 0.000398 | 20.35756 | 20.61153 | 20.34039 |

| | | | | | | | | |
|---|---|---|---|---|---|---|---|---|
| P59998 | Actin-relate | ARPC4 | Actin-relate | 5.652732 | 0.112653 | 21.33963 | 20.17585 | 20.55253 |
| Q53G59 | Kelch-like p | KLHL12 | Kelch-like p | 6.58041 | 0.067475 | 21.55927 | 20.33865 | 20.71749 |
| Q9NYB9 | Abl interact | ABI2 | Abl interact | 8.207504 | 1.87E-05 | 20.96846 | 20.88554 | 21.14046 |
| Q86TI0 | TBC1 doma | TBC1D1 | TBC1 doma | 6.606294 | 0.000261 | 20.31937 | 20.02756 | 20.58316 |
| Q96JG8 | Melanoma- | MAGED4 | Melanoma- | 6.03393 | 0.085281 | 20.87418 | 20.77612 | 20.75075 |
| Q9BVQ7 | Spermatog | SPATA5L1 | ATPase fam | 6.545643 | 0.000107 | 20.63139 | 19.74189 | 20.37624 |
| Q63ZY3 | KN motif an | KANK2 | KN motif an | 8.011079 | 0.001103 | 19.93517 | 20.43919 | 20.60286 |
| Q6AI08 | HEAT repea | HEATR6 | HEAT repea | 8.00481 | 2.13E-05 | 20.8353 | 20.66813 | 20.84192 |
| P78395 | Melanoma | PRAME | Melanoma | 8.020754 | 0.000314 | 20.98753 | 20.75328 | 20.68663 |
| P50995 | Annexin A1 | ANXA11 | Annexin A1 | 6.076425 | 0.076609 | 20.28828 | 20.77258 | 20.7446 |
| Q9Y2W2 | WW domai | WBP11 | WW domai | 6.902683 | 0.053523 | 21.27321 | 20.31893 | 20.75939 |
| Q14188 | Transcriptio | TFDP2 | Transcriptio | 7.518718 | 2.96E-06 | 20.64922 | 20.61585 | 20.58141 |
| P00367 | Glutamate | GLUD1 | Glutamate | 8.860742 | 2.37E-05 | 20.9047 | 20.68987 | 20.93647 |
| O43707 | Alpha-actin | ACTN4 | Alpha-actin | 5.918564 | 0.094385 | 20.68808 | 20.15314 | 20.05669 |
| P32780 | General tra | GTF2H1 | General tra | 7.156337 | 0.000726 | 20.59231 | 20.1168 | 20.49121 |
| Q12894 | Interferon-r | IFRD2 | Interferon-r | 8.194525 | 0.000346 | 21.26934 | 20.26588 | 21.36234 |
| O75147 | Obscurin-li | OBSL1 | Obscurin-li | 7.644627 | 6.77E-05 | 19.86591 | 20.96155 | 20.16325 |
| O14965 | Aurora kina | AURKA | Aurora kina | 8.489697 | 9.53E-05 | 20.92991 | 20.65403 | 20.48827 |
| Q6P2E9 | Enhancer o | EDC4 | Enhancer o | 8.041941 | 8.91E-05 | 19.93805 | 19.64678 | 20.15993 |
| Q6PML9 | Zinc transp | SLC30A9 | Proton-cou | 7.741879 | 0.000324 | 19.93905 | 19.92812 | 20.7766 |
| O75530 | Polycomb | EED | Polycomb | 8.227685 | 0.000612 | 19.76417 | 21.35606 | 21.059 |
| Q9BYE7 | Polycomb | PCGF6 | Polycomb | 6.009388 | 0.0014 | 19.81239 | 18.81702 | 20.13896 |
| P42357 | Histidine an | HAL | Histidine an | 5.066334 | 0.102122 | 19.58068 | 19.62608 | 19.72234 |
| Q92797 | Symplekin | SYMPK | Symplekin | 5.807572 | 0.096407 | 20.24016 | 20.67808 | 20.563 |
| Q15773 | Myeloid leu | MLF2 | Myeloid leu | 8.422208 | 7.46E-06 | 20.36677 | 20.49629 | 20.89112 |
| Q92979 | Ribosomal | EMG1 | Ribosomal | 5.558922 | 0.053975 | 20.45704 | 21.1194 | 20.18684 |
| P42345 | Serine/thre | MTOR | Serine/thre | 7.022154 | 0.000514 | 20.13283 | 20.13195 | 19.97994 |
| O75223 | Gamma-glu | GGCT | Gamma-glu | 4.681152 | 0.142487 | 20.01686 | 19.22973 | 20.51282 |
| Q9Y5Y2 | Cytosolic F | NUBP2 | Cytosolic F | 4.931972 | 0.055032 | 19.32843 | 19.52064 | 20.64676 |
| Q9NXH8 | Torsin-4A | TOR4A | Torsin-4A O | 7.615653 | 0.002124 | 20.96346 | 19.6052 | 20.85582 |
| P61225;Q9 | Ras-related | RAP2B;RAP | Ras-related | 7.594161 | 0.000531 | 20.35337 | 19.85517 | 20.59222 |
| Q8WU90 | Zinc finger C | ZC3H15 | Zinc finger C | 6.894705 | 0.003319 | 20.05411 | 20.16509 | 20.25811 |

| Log 2 LFQ intensity IgG R1 | Log2 LFQ intensity IgG R2 | Log2 LFQ intensity IgG R3 | Peptides | Razor + unique peptides | Unique peptides | Sequence coverage [%] | Intensity |
|---|---|---|---|---|---|---|---|
| 23.59582 | 25.4972 | 25.82148 | 192 | 192 | 164 | 64.6 | 8.12E+10 |
| 25.4224 | 27.3652 | 27.29149 | 27 | 27 | 10 | 65.6 | 6.64E+10 |
| 18.47991 | 21.18156 | 18.93672 | 56 | 56 | 53 | 81.1 | 3.25E+10 |
| 23.46314 | 24.32196 | 21.1687 | 13 | 13 | 13 | 42.9 | 3.23E+10 |
| 22.90621 | 24.92348 | 23.60388 | 15 | 15 | 15 | 62.1 | 2.55E+10 |
| 19.42536 | 21.862 | 18.27403 | 24 | 24 | 24 | 53.6 | 2.30E+10 |
| 12.92216 | 10.86674 | 12.42288 | 45 | 45 | 45 | 51.3 | 2.30E+10 |
| 23.54203 | 25.16741 | 23.32581 | 19 | 19 | 7 | 43.7 | 2.23E+10 |
| 12.54313 | 14.61804 | 14.71639 | 139 | 139 | 139 | 55.2 | 2.11E+10 |
| 11.4994 | 16.16748 | 15.78892 | 38 | 38 | 38 | 57.3 | 2.05E+10 |
| 20.91998 | 23.00441 | 21.53924 | 23 | 23 | 5 | 58.8 | 1.91E+10 |
| 14.5842 | 18.54518 | 14.74693 | 34 | 34 | 33 | 71.1 | 1.87E+10 |
| 13.40846 | 14.03179 | 12.61218 | 114 | 114 | 114 | 56.6 | 1.83E+10 |
| 18.89226 | 22.37188 | 19.44407 | 21 | 21 | 21 | 64.3 | 1.79E+10 |
| 17.1205 | 14.98268 | 12.34892 | 21 | 21 | 21 | 46.9 | 1.75E+10 |
| 21.1359 | 23.46076 | 21.79142 | 26 | 26 | 0 | 58.3 | 1.69E+10 |
| 13.7456 | 17.03928 | 13.70383 | 41 | 41 | 41 | 55.6 | 1.50E+10 |
| 21.30919 | 24.08205 | 21.50698 | 25 | 25 | 25 | 61.2 | 1.49E+10 |
| 17.84298 | 20.50019 | 19.37647 | 27 | 27 | 27 | 44.7 | 1.45E+10 |
| 15.09139 | 13.91317 | 12.02566 | 38 | 38 | 38 | 61.9 | 1.42E+10 |
| 20.84529 | 23.88725 | 21.60273 | 9 | 9 | 3 | 51.6 | 1.37E+10 |
| 18.63548 | 20.7729 | 19.85307 | 49 | 49 | 49 | 58.5 | 1.29E+10 |
| 14.91896 | 14.34651 | 13.12799 | 115 | 115 | 115 | 54.8 | 1.23E+10 |
| 19.31216 | 22.96049 | 19.821 | 15 | 15 | 15 | 69.9 | 1.19E+10 |
| 18.69836 | 22.55452 | 20.72067 | 20 | 20 | 20 | 54 | 1.13E+10 |
| 17.44265 | 20.9307 | 18.81719 | 25 | 25 | 25 | 56.6 | 1.13E+10 |
| 18.90971 | 21.91787 | 19.5945 | 18 | 18 | 18 | 54.1 | 1.12E+10 |
| 15.86373 | 18.9757 | 11.28279 | 52 | 52 | 51 | 58 | 1.07E+10 |
| 18.3304 | 22.80984 | 19.3363 | 25 | 25 | 25 | 63.3 | 1.05E+10 |
| 18.97006 | 21.08803 | 19.27842 | 21 | 21 | 21 | 53.2 | 1.04E+10 |
| 17.62435 | 21.01944 | 18.9043 | 17 | 17 | 17 | 45.9 | 1.03E+10 |
| 19.70361 | 21.28048 | 20.94457 | 10 | 10 | 10 | 55.1 | 9.85E+09 |
| 19.88938 | 21.79646 | 20.96902 | 12 | 12 | 12 | 45.7 | 9.82E+09 |
| 21.3641 | 24.76618 | 21.91576 | 7 | 7 | 2 | 35.9 | 9.80E+09 |
| 18.15679 | 22.34055 | 18.28582 | 31 | 31 | 31 | 35.3 | 9.62E+09 |
| 22.76903 | 25.0436 | 23.01545 | 27 | 27 | 27 | 61.1 | 9.54E+09 |
| 23.05041 | 25.01448 | 25.3413 | 9 | 9 | 7 | 56.3 | 9.15E+09 |
| 15.53394 | 19.65192 | 17.45467 | 29 | 29 | 29 | 52.4 | 9.01E+09 |
| 17.04356 | 20.58527 | 19.97771 | 11 | 11 | 11 | 46.3 | 8.89E+09 |
| 18.21869 | 21.26276 | 19.25613 | 17 | 17 | 17 | 53.6 | 8.67E+09 |
| 15.62362 | 18.67456 | 16.08568 | 47 | 47 | 47 | 65.1 | 8.33E+09 |
| 24.00067 | 26.6102 | 24.14989 | 15 | 15 | 6 | 40.3 | 8.17E+09 |
| 17.89006 | 19.282 | 18.5998 | 16 | 16 | 16 | 45.1 | 8.11E+09 |
| 11.29428 | 13.59887 | 12.53019 | 44 | 44 | 44 | 46.7 | 8.06E+09 |

| | | | | | | | |
|---|---|---|---|---|---|---|---|
| 17.74459 | 21.82369 | 18.50133 | 28 | 28 | 28 | 55.5 | 7.85E+09 |
| 19.73856 | 21.84076 | 20.14681 | 13 | 13 | 13 | 58.3 | 7.76E+09 |
| 24.10462 | 26.49863 | 24.24445 | 19 | 19 | 19 | 62.5 | 7.74E+09 |
| 19.99852 | 22.64625 | 20.22423 | 14 | 14 | 14 | 69.2 | 7.73E+09 |
| 17.09063 | 20.09828 | 19.1279 | 14 | 14 | 14 | 51 | 7.71E+09 |
| 25.65696 | 29.89682 | 27.19442 | 83 | 83 | 78 | 22.3 | 7.70E+09 |
| 15.50342 | 18.70523 | 16.59845 | 40 | 40 | 29 | 62.4 | 7.44E+09 |
| 16.8681 | 21.91627 | 18.00859 | 33 | 33 | 33 | 58.2 | 7.30E+09 |
| 18.756 | 22.1121 | 20.32553 | 18 | 18 | 6 | 43.3 | 7.19E+09 |
| 12.32317 | 13.88369 | 12.48776 | 46 | 46 | 46 | 45.5 | 6.96E+09 |
| 17.89847 | 23.7348 | 20.10457 | 6 | 6 | 2 | 27.2 | 6.91E+09 |
| 18.98053 | 22.79237 | 18.8119 | 26 | 25 | 25 | 52.7 | 6.89E+09 |
| 23.09094 | 23.94887 | 22.32158 | 11 | 11 | 11 | 32 | 6.86E+09 |
| 20.22235 | 21.84521 | 20.52315 | 50 | 50 | 50 | 39.5 | 6.86E+09 |
| 20.8674 | 29.72061 | 22.62883 | 30 | 30 | 30 | 37.7 | 6.66E+09 |
| 16.23678 | 19.68267 | 14.9177 | 16 | 16 | 16 | 39.8 | 6.54E+09 |
| 16.54046 | 22.48618 | 16.9704 | 22 | 22 | 22 | 72 | 6.49E+09 |
| 19.52309 | 21.56579 | 20.64148 | 10 | 10 | 10 | 37.4 | 6.48E+09 |
| 21.43487 | 24.18767 | 24.41661 | 12 | 12 | 5 | 55 | 6.37E+09 |
| 18.86852 | 21.89475 | 19.1538 | 39 | 39 | 34 | 50.7 | 6.32E+09 |
| 13.9796 | 13.81868 | 13.63152 | 93 | 93 | 93 | 49.8 | 6.05E+09 |
| 12.794 | 11.21102 | 14.85326 | 31 | 31 | 31 | 51.6 | 6.01E+09 |
| 18.71055 | 21.49473 | 19.59842 | 10 | 10 | 10 | 31.3 | 5.70E+09 |
| 18.41951 | 20.88621 | 19.32613 | 10 | 10 | 9 | 95.7 | 5.61E+09 |
| 20.35445 | 21.32312 | 20.46931 | 13 | 13 | 13 | 45.1 | 5.46E+09 |
| 17.00295 | 19.14113 | 17.57555 | 50 | 50 | 50 | 57.9 | 5.05E+09 |
| 20.79559 | 23.13256 | 20.55525 | 16 | 16 | 16 | 51.9 | 5.05E+09 |
| 21.24685 | 22.5813 | 21.18223 | 14 | 14 | 14 | 39.3 | 5.01E+09 |
| 22.26479 | 24.61818 | 22.78674 | 9 | 9 | 9 | 48 | 5.00E+09 |
| 16.59663 | 19.62888 | 19.1792 | 11 | 11 | 11 | 48.2 | 4.94E+09 |
| 14.2048 | 17.29661 | 13.70466 | 23 | 23 | 23 | 40.7 | 4.79E+09 |
| 19.93155 | 23.92488 | 17.54019 | 17 | 17 | 17 | 66.5 | 4.70E+09 |
| 29.45138 | 21.41978 | 22.92539 | 29 | 29 | 29 | 30.8 | 4.69E+09 |
| 21.61005 | 22.96031 | 21.7645 | 31 | 30 | 24 | 46.9 | 4.68E+09 |
| 13.13252 | 15.84823 | 13.6049 | 17 | 17 | 17 | 42.4 | 4.50E+09 |
| 18.64832 | 21.42465 | 19.39496 | 8 | 8 | 8 | 38.8 | 4.48E+09 |
| 14.39238 | 16.61813 | 14.86988 | 44 | 44 | 44 | 48.8 | 4.35E+09 |
| 13.69621 | 13.96669 | 13.15436 | 119 | 119 | 119 | 42.9 | 4.28E+09 |
| 17.99314 | 19.34041 | 17.08213 | 9 | 9 | 9 | 47.9 | 4.24E+09 |
| 14.45469 | 17.30626 | 16.82451 | 87 | 87 | 86 | 37.5 | 4.23E+09 |
| 22.08651 | 23.83132 | 21.98919 | 8 | 8 | 8 | 38.2 | 4.23E+09 |
| 19.55476 | 21.54062 | 18.15813 | 8 | 8 | 8 | 33.1 | 4.17E+09 |
| 14.33783 | 14.63911 | 14.19722 | 11 | 11 | 11 | 30.7 | 4.10E+09 |
| 21.68539 | 24.39846 | 21.94853 | 9 | 9 | 9 | 57 | 4.09E+09 |
| 12.25352 | 14.83457 | 14.27401 | 39 | 39 | 37 | 45.7 | 4.08E+09 |
| 15.60247 | 17.18747 | 16.61655 | 29 | 29 | 29 | 28.9 | 4.02E+09 |
| 11.29267 | 13.50841 | 11.11667 | 35 | 35 | 24 | 54.1 | 3.97E+09 |
| 14.19837 | 18.25114 | 15.98114 | 12 | 12 | 2 | 52.4 | 3.90E+09 |

| | | | | | | | |
|---|---|---|---|---|---|---|---|
| 18.4924 | 20.76012 | 19.5461 | 12 | 12 | 12 | 44.8 | 3.87E+09 |
| 13.54255 | 14.45687 | 10.6989 | 28 | 28 | 28 | 48.8 | 3.85E+09 |
| 19.71793 | 23.4805 | 20.90588 | 28 | 28 | 28 | 54.6 | 3.84E+09 |
| 12.58827 | 14.83348 | 13.86486 | 10 | 10 | 10 | 55.5 | 3.84E+09 |
| 15.07439 | 15.34287 | 14.66772 | 25 | 25 | 25 | 33.8 | 3.81E+09 |
| 18.54181 | 21.82439 | 19.25854 | 11 | 11 | 11 | 38 | 3.73E+09 |
| 14.02306 | 15.62122 | 13.96921 | 18 | 18 | 12 | 68.8 | 3.65E+09 |
| 15.79741 | 19.82779 | 16.66122 | 19 | 19 | 19 | 55.2 | 3.62E+09 |
| 11.80731 | 16.57411 | 11.42022 | 121 | 121 | 121 | 31.1 | 3.61E+09 |
| 18.78721 | 21.77133 | 18.764 | 11 | 11 | 11 | 43 | 3.60E+09 |
| 19.66412 | 22.34511 | 20.12187 | 9 | 9 | 9 | 48.8 | 3.59E+09 |
| 17.60849 | 20.54104 | 18.59602 | 5 | 5 | 5 | 22.2 | 3.57E+09 |
| 16.97612 | 18.93462 | 16.83861 | 42 | 42 | 41 | 42.1 | 3.56E+09 |
| 19.91863 | 21.73929 | 20.54764 | 7 | 7 | 7 | 33.5 | 3.53E+09 |
| 18.8652 | 20.85506 | 19.84353 | 4 | 4 | 4 | 33.3 | 3.50E+09 |
| 12.74161 | 14.32052 | 12.99903 | 36 | 36 | 32 | 47.1 | 3.47E+09 |
| 16.86737 | 19.41229 | 17.84531 | 73 | 73 | 73 | 33 | 3.45E+09 |
| 16.02464 | 17.6768 | 11.50547 | 18 | 18 | 18 | 44.8 | 3.38E+09 |
| 19.13874 | 21.36255 | 21.61239 | 107 | 89 | 86 | 49.8 | 3.34E+09 |
| 12.59107 | 16.80574 | 11.91826 | 19 | 19 | 19 | 66.7 | 3.33E+09 |
| 20.05179 | 22.01182 | 20.74427 | 6 | 6 | 6 | 34.3 | 3.29E+09 |
| 21.47981 | 24.94668 | 21.96127 | 31 | 31 | 30 | 47.9 | 3.27E+09 |
| 23.48714 | 26.57267 | 23.20361 | 17 | 17 | 17 | 69.7 | 3.23E+09 |
| 19.00766 | 20.22459 | 20.33353 | 29 | 29 | 27 | 53.8 | 3.19E+09 |
| 14.57134 | 15.70871 | 14.37511 | 14 | 14 | 10 | 34.9 | 3.19E+09 |
| 21.67293 | 22.59987 | 21.92586 | 36 | 36 | 35 | 55.7 | 3.17E+09 |
| 17.48731 | 19.63222 | 17.94792 | 11 | 11 | 11 | 38.6 | 3.12E+09 |
| 19.18141 | 23.37491 | 19.66955 | 20 | 20 | 20 | 52.9 | 3.08E+09 |
| 18.69347 | 21.85776 | 18.25709 | 7 | 7 | 7 | 36.1 | 3.05E+09 |
| 13.60119 | 16.34748 | 11.60797 | 15 | 15 | 13 | 44.8 | 3.04E+09 |
| 13.10388 | 15.08493 | 14.31217 | 29 | 22 | 22 | 44.4 | 2.80E+09 |
| 15.32119 | 14.41237 | 14.01489 | 28 | 28 | 28 | 54.1 | 2.78E+09 |
| 11.66596 | 16.61712 | 14.84647 | 29 | 29 | 29 | 31.2 | 2.69E+09 |
| 16.97085 | 17.02605 | 18.47931 | 6 | 6 | 6 | 33.3 | 2.68E+09 |
| 17.89409 | 20.73562 | 17.83028 | 12 | 12 | 12 | 46.5 | 2.68E+09 |
| 19.80185 | 22.05563 | 20.48168 | 27 | 27 | 14 | 43.5 | 2.66E+09 |
| 13.99373 | 15.59933 | 14.24154 | 90 | 90 | 90 | 50.5 | 2.60E+09 |
| 20.42132 | 23.37637 | 16.9798 | 16 | 9 | 9 | 43.7 | 2.59E+09 |
| 19.10753 | 21.01964 | 19.78995 | 4 | 4 | 4 | 17.6 | 2.59E+09 |
| 16.13827 | 17.16703 | 16.49473 | 47 | 47 | 47 | 43.8 | 2.56E+09 |
| 14.78928 | 17.33638 | 15.19641 | 30 | 30 | 23 | 53 | 2.56E+09 |
| 16.63268 | 19.66504 | 17.29401 | 37 | 37 | 37 | 52.1 | 2.54E+09 |
| 22.03919 | 25.07806 | 24.46824 | 76 | 68 | 68 | 45.4 | 2.53E+09 |
| 11.24757 | 22.6174 | 10.61792 | 23 | 6 | 1 | 58.7 | 2.46E+09 |
| 16.06808 | 15.51653 | 13.95919 | 47 | 47 | 47 | 62.8 | 2.45E+09 |
| 14.29842 | 14.22799 | 13.61252 | 31 | 31 | 31 | 40.4 | 2.41E+09 |
| 10.41863 | 12.58999 | 10.80088 | 20 | 20 | 20 | 47.5 | 2.36E+09 |
| 12.80641 | 15.8269 | 11.65748 | 54 | 54 | 54 | 31.6 | 2.35E+09 |

| | | | | | | | |
|---|---|---|---|---|---|---|---|
| 16.87051 | 22.25254 | 17.02562 | 21 | 21 | 21 | 51.7 | 2.34E+09 |
| 14.09036 | 18.00459 | 16.34544 | 23 | 23 | 23 | 57.1 | 2.32E+09 |
| 13.05066 | 15.07322 | 13.51299 | 41 | 41 | 41 | 39.9 | 2.31E+09 |
| 9.99903 | 19.86354 | 19.2616 | 18 | 18 | 15 | 60.4 | 2.31E+09 |
| 13.74651 | 17.48298 | 13.98014 | 14 | 14 | 13 | 30.7 | 2.30E+09 |
| 19.07755 | 21.31058 | 18.22157 | 8 | 8 | 8 | 47.8 | 2.27E+09 |
| 13.99289 | 14.3384 | 11.95107 | 3 | 3 | 3 | 27 | 2.27E+09 |
| 14.70531 | 14.17843 | 10.70053 | 9 | 9 | 9 | 47.4 | 2.19E+09 |
| 10.24581 | 14.8814 | 13.60258 | 17 | 17 | 17 | 51.8 | 2.18E+09 |
| 16.72161 | 20.24516 | 19.20221 | 6 | 6 | 6 | 23.3 | 2.14E+09 |
| 11.61187 | 19.93618 | 18.1165 | 4 | 4 | 4 | 26.1 | 2.10E+09 |
| 20.75971 | 24.46582 | 21.06467 | 28 | 27 | 27 | 56.5 | 2.09E+09 |
| 14.67986 | 15.45943 | 13.90802 | 21 | 21 | 21 | 48.5 | 2.08E+09 |
| 20.29749 | 24.43892 | 20.03537 | 7 | 7 | 7 | 32.8 | 2.08E+09 |
| 16.18503 | 18.55962 | 13.92416 | 42 | 42 | 42 | 44.6 | 2.08E+09 |
| 12.03663 | 19.16342 | 11.49317 | 16 | 16 | 16 | 28.6 | 2.07E+09 |
| 11.62678 | 12.31305 | 13.43267 | 30 | 30 | 28 | 53.6 | 2.05E+09 |
| 13.2652 | 19.44374 | 14.68978 | 29 | 28 | 28 | 47.7 | 2.00E+09 |
| 12.8853 | 14.3286 | 11.58829 | 14 | 14 | 13 | 57.2 | 1.99E+09 |
| 17.39331 | 17.40989 | 13.21992 | 34 | 33 | 33 | 40.2 | 1.96E+09 |
| 15.92623 | 15.84426 | 15.80974 | 60 | 60 | 60 | 25 | 1.96E+09 |
| 14.9506 | 17.07106 | 15.6868 | 38 | 38 | 38 | 49.2 | 1.95E+09 |
| 14.30955 | 16.48886 | 11.70608 | 32 | 32 | 32 | 43.1 | 1.89E+09 |
| 11.73334 | 19.95617 | 18.01531 | 11 | 9 | 7 | 34.3 | 1.89E+09 |
| 21.33903 | 22.49001 | 22.06881 | 28 | 28 | 28 | 64 | 1.87E+09 |
| 12.15046 | 19.39741 | 16.22395 | 8 | 7 | 7 | 41.5 | 1.85E+09 |
| 20.54623 | 20.72 | 19.53184 | 6 | 6 | 6 | 81.4 | 1.85E+09 |
| 12.04545 | 13.09268 | 10.87781 | 22 | 22 | 22 | 53 | 1.84E+09 |
| 21.28071 | 23.38021 | 23.59719 | 28 | 17 | 13 | 62.3 | 1.83E+09 |
| 19.74092 | 22.5858 | 15.88147 | 5 | 5 | 5 | 52.9 | 1.82E+09 |
| 12.20714 | 14.09218 | 15.5136 | 19 | 19 | 19 | 55.4 | 1.82E+09 |
| 12.69765 | 15.05295 | 13.46217 | 32 | 32 | 31 | 47.3 | 1.82E+09 |
| 19.25893 | 17.74169 | 19.53733 | 33 | 23 | 21 | 49.7 | 1.81E+09 |
| 12.04774 | 12.68563 | 14.115 | 38 | 38 | 38 | 28.4 | 1.80E+09 |
| 16.66539 | 18.25714 | 15.20075 | 6 | 6 | 6 | 37.5 | 1.79E+09 |
| 14.39681 | 19.68061 | 16.61913 | 29 | 29 | 29 | 47.1 | 1.78E+09 |
| 12.69157 | 14.99691 | 14.02219 | 46 | 46 | 46 | 30.7 | 1.77E+09 |
| 15.84142 | 19.95419 | 15.74567 | 23 | 23 | 23 | 54.9 | 1.76E+09 |
| 20.89815 | 24.11188 | 19.93301 | 10 | 10 | 10 | 57.8 | 1.76E+09 |
| 15.70244 | 17.32061 | 15.65732 | 35 | 35 | 34 | 43.3 | 1.75E+09 |
| 11.2465 | 14.69218 | 15.85309 | 57 | 57 | 57 | 25.9 | 1.73E+09 |
| 20.68123 | 22.20171 | 21.45342 | 10 | 10 | 8 | 36.2 | 1.73E+09 |
| 11.70118 | 14.48375 | 11.67424 | 53 | 53 | 53 | 57.3 | 1.72E+09 |
| 16.3532 | 20.12591 | 16.87412 | 15 | 15 | 15 | 52.4 | 1.72E+09 |
| 11.75089 | 17.44621 | 17.45459 | 42 | 42 | 42 | 31.1 | 1.71E+09 |
| 12.45322 | 15.52402 | 12.74001 | 27 | 27 | 27 | 45.5 | 1.70E+09 |
| 13.9405 | 18.21622 | 13.80221 | 29 | 24 | 24 | 36.2 | 1.70E+09 |
| 13.25301 | 17.22185 | 14.52062 | 19 | 19 | 16 | 43.3 | 1.70E+09 |

| | | | | | | | |
|---|---|---|---|---|---|---|---|
| 23.24493 | 25.42541 | 21.2064 | 8 | 8 | 8 | 40.1 | 1.69E+09 |
| 10.49717 | 17.45129 | 13.82028 | 28 | 28 | 28 | 57.1 | 1.69E+09 |
| 15.69444 | 20.78397 | 18.27753 | 6 | 6 | 6 | 54.5 | 1.67E+09 |
| 14.04644 | 14.55147 | 14.0175 | 39 | 39 | 39 | 44.3 | 1.67E+09 |
| 19.43884 | 20.84506 | 19.42008 | 8 | 2 | 2 | 50 | 1.65E+09 |
| 12.51274 | 16.90207 | 12.93402 | 16 | 16 | 16 | 40.8 | 1.64E+09 |
| 20.72409 | 21.24731 | 21.26218 | 13 | 13 | 13 | 31.6 | 1.64E+09 |
| 22.38068 | 25.54153 | 22.11312 | 11 | 11 | 11 | 57 | 1.63E+09 |
| 18.9455 | 20.28478 | 18.92286 | 47 | 47 | 47 | 48.1 | 1.63E+09 |
| 15.19063 | 14.29498 | 14.52497 | 16 | 16 | 10 | 8.7 | 1.61E+09 |
| 11.99707 | 16.2452 | 13.30276 | 23 | 20 | 20 | 38.2 | 1.61E+09 |
| 16.60481 | 15.43502 | 17.32351 | 72 | 72 | 72 | 35.1 | 1.60E+09 |
| 18.0654 | 19.16759 | 16.40556 | 26 | 26 | 26 | 31.9 | 1.58E+09 |
| 13.28586 | 13.17415 | 15.93765 | 23 | 23 | 23 | 45.6 | 1.57E+09 |
| 12.4159 | 17.62421 | 13.37082 | 22 | 22 | 22 | 51.9 | 1.56E+09 |
| 12.86486 | 15.04247 | 12.80361 | 26 | 26 | 26 | 38.3 | 1.55E+09 |
| 20.60241 | 22.03855 | 19.73089 | 17 | 5 | 5 | 38.4 | 1.53E+09 |
| 11.48777 | 15.93088 | 14.34942 | 52 | 52 | 52 | 29.1 | 1.51E+09 |
| 20.03241 | 23.3692 | 22.69855 | 4 | 4 | 4 | 14.9 | 1.50E+09 |
| 20.41854 | 21.46603 | 21.06546 | 35 | 35 | 35 | 51.3 | 1.48E+09 |
| 20.14071 | 13.69031 | 12.04383 | 3 | 3 | 3 | 2.9 | 1.44E+09 |
| 13.86079 | 18.4931 | 16.59959 | 24 | 24 | 24 | 38.9 | 1.43E+09 |
| 11.83641 | 14.9376 | 12.16914 | 32 | 32 | 32 | 47.7 | 1.42E+09 |
| 17.4578 | 20.4715 | 19.01985 | 49 | 49 | 44 | 26.5 | 1.38E+09 |
| 13.37436 | 14.85141 | 12.5631 | 60 | 60 | 60 | 35.4 | 1.37E+09 |
| 12.95037 | 16.67659 | 13.96244 | 21 | 21 | 21 | 27.3 | 1.37E+09 |
| 11.32227 | 12.7148 | 11.68891 | 24 | 24 | 24 | 44.6 | 1.35E+09 |
| 14.05926 | 11.31032 | 10.92533 | 17 | 14 | 14 | 49.4 | 1.35E+09 |
| 14.8013 | 18.31279 | 11.19125 | 21 | 14 | 10 | 51.1 | 1.34E+09 |
| 13.46817 | 15.26455 | 16.31129 | 20 | 20 | 20 | 36.2 | 1.34E+09 |
| 17.67072 | 21.12263 | 15.29444 | 12 | 12 | 12 | 38.7 | 1.33E+09 |
| 12.27525 | 17.56785 | 12.12738 | 9 | 9 | 8 | 30.7 | 1.32E+09 |
| 13.59444 | 17.47048 | 17.62364 | 18 | 18 | 18 | 43.5 | 1.32E+09 |
| 19.92554 | 22.42795 | 18.24321 | 17 | 17 | 17 | 27.2 | 1.32E+09 |
| 17.11217 | 18.56863 | 18.22034 | 19 | 19 | 19 | 44.6 | 1.29E+09 |
| 15.14685 | 18.41897 | 19.36468 | 37 | 37 | 37 | 43.3 | 1.28E+09 |
| 21.43715 | 21.13521 | 21.84161 | 27 | 27 | 27 | 34 | 1.27E+09 |
| 13.31784 | 13.81658 | 15.52659 | 60 | 60 | 40 | 27.9 | 1.26E+09 |
| 22.92825 | 23.67088 | 23.41648 | 26 | 26 | 26 | 29.8 | 1.26E+09 |
| 13.3253 | 14.95488 | 14.28757 | 35 | 35 | 35 | 29.2 | 1.24E+09 |
| 20.84751 | 21.85274 | 21.30117 | 6 | 6 | 6 | 44.6 | 1.24E+09 |
| 11.12585 | 18.65459 | 11.23915 | 13 | 13 | 13 | 42.2 | 1.24E+09 |
| 12.03246 | 13.79279 | 13.17458 | 44 | 44 | 44 | 27.8 | 1.23E+09 |
| 18.36965 | 21.0625 | 21.04166 | 26 | 26 | 26 | 45.8 | 1.23E+09 |
| 20.16103 | 17.60052 | 20.20717 | 58 | 58 | 58 | 27.5 | 1.22E+09 |
| 17.07075 | 16.90301 | 12.56125 | 50 | 50 | 50 | 36.5 | 1.21E+09 |
| 25.01996 | 25.24561 | 25.20314 | 14 | 14 | 14 | 32.6 | 1.20E+09 |
| 15.48699 | 17.73211 | 13.90359 | 30 | 30 | 30 | 36.3 | 1.18E+09 |

| | | | | | | | |
|---|---|---|---|---|---|---|---|
| 16.42879 | 13.79462 | 12.31699 | 7 | 7 | 7 | 25.4 | 1.16E+09 |
| 12.37928 | 14.99475 | 12.91889 | 21 | 21 | 19 | 40.4 | 1.16E+09 |
| 22.35595 | 23.32553 | 22.98867 | 40 | 40 | 40 | 41.5 | 1.14E+09 |
| 20.93005 | 22.53017 | 20.9502 | 8 | 8 | 7 | 61.4 | 1.13E+09 |
| 11.29167 | 17.5648 | 11.10889 | 13 | 13 | 12 | 43.9 | 1.13E+09 |
| 22.88867 | 23.62556 | 23.47209 | 27 | 27 | 27 | 28.2 | 1.13E+09 |
| 17.58586 | 16.8343 | 17.54751 | 28 | 28 | 11 | 47.5 | 1.12E+09 |
| 24.95934 | 27.42374 | 24.91159 | 24 | 24 | 24 | 41.1 | 1.12E+09 |
| 23.28093 | 25.39352 | 22.93058 | 4 | 4 | 4 | 9 | 1.10E+09 |
| 20.48267 | 22.89778 | 18.53389 | 20 | 20 | 20 | 27.8 | 1.09E+09 |
| 17.55037 | 17.4492 | 17.58057 | 19 | 19 | 19 | 50 | 1.09E+09 |
| 22.83655 | 23.59764 | 23.42816 | 15 | 15 | 15 | 36.4 | 1.07E+09 |
| 17.17104 | 20.69422 | 16.29009 | 62 | 62 | 62 | 23.9 | 1.06E+09 |
| 12.08849 | 14.91882 | 12.25677 | 27 | 27 | 27 | 37.9 | 1.06E+09 |
| 13.80514 | 20.8353 | 14.80937 | 27 | 27 | 27 | 55.3 | 1.06E+09 |
| 14.44514 | 14.71113 | 11.71694 | 15 | 15 | 15 | 61.9 | 1.05E+09 |
| 14.6052 | 18.7796 | 13.62023 | 8 | 8 | 8 | 38.9 | 1.05E+09 |
| 15.07209 | 17.75753 | 16.37843 | 42 | 42 | 32 | 32.7 | 1.05E+09 |
| 19.37742 | 22.94601 | 19.53285 | 9 | 9 | 9 | 37.3 | 1.04E+09 |
| 10.84872 | 18.10226 | 15.50621 | 17 | 17 | 17 | 42.9 | 1.03E+09 |
| 16.38163 | 19.88777 | 18.54034 | 14 | 14 | 12 | 33.8 | 1.03E+09 |
| 13.10281 | 13.65161 | 12.08471 | 44 | 44 | 44 | 36.2 | 1.02E+09 |
| 16.28733 | 14.92514 | 12.93468 | 52 | 52 | 52 | 14.5 | 1.02E+09 |
| 22.20717 | 22.99155 | 22.56544 | 12 | 12 | 7 | 27.8 | 1.01E+09 |
| 22.86452 | 24.13782 | 26.97717 | 18 | 18 | 18 | 23.9 | 1.01E+09 |
| 18.18607 | 20.14183 | 18.67225 | 33 | 33 | 33 | 34.6 | 1.01E+09 |
| 18.31973 | 21.0347 | 19.43394 | 22 | 22 | 22 | 44.5 | 9.95E+08 |
| 13.80201 | 17.96787 | 14.79775 | 4 | 4 | 3 | 31 | 9.92E+08 |
| 16.08466 | 14.8519 | 14.39781 | 41 | 41 | 41 | 40.4 | 9.92E+08 |
| 22.67607 | 25.34557 | 22.0309 | 12 | 12 | 10 | 21.1 | 9.78E+08 |
| 16.5093 | 21.93106 | 18.93882 | 40 | 40 | 31 | 27.9 | 9.71E+08 |
| 13.79126 | 12.07665 | 13.09653 | 41 | 41 | 28 | 27.2 | 9.68E+08 |
| 16.43288 | 13.21118 | 12.85893 | 38 | 38 | 38 | 38.4 | 9.67E+08 |
| 12.2568 | 16.48668 | 12.58339 | 36 | 36 | 36 | 33.4 | 9.54E+08 |
| 16.85246 | 18.58186 | 17.28111 | 20 | 20 | 20 | 36.2 | 9.47E+08 |
| 21.70885 | 22.91523 | 19.92855 | 7 | 7 | 6 | 39.8 | 9.29E+08 |
| 11.1153 | 16.1205 | 16.01003 | 28 | 28 | 28 | 47.3 | 9.20E+08 |
| 18.66324 | 21.62609 | 19.40179 | 25 | 22 | 13 | 43.5 | 9.20E+08 |
| 18.58556 | 19.38995 | 13.03726 | 27 | 27 | 27 | 37.9 | 8.93E+08 |
| 23.05676 | 27.60421 | 22.64194 | 9 | 9 | 9 | 19.7 | 8.90E+08 |
| 12.92511 | 14.1217 | 13.38627 | 21 | 21 | 21 | 39.7 | 8.89E+08 |
| 12.3606 | 12.22525 | 11.11968 | 17 | 17 | 17 | 28.4 | 8.86E+08 |
| 17.8541 | 15.24681 | 16.33682 | 14 | 14 | 14 | 36.9 | 8.81E+08 |
| 13.8575 | 12.82077 | 13.80443 | 28 | 28 | 28 | 34 | 8.81E+08 |
| 19.66452 | 22.54882 | 11.22562 | 6 | 6 | 6 | 44.9 | 8.79E+08 |
| 13.78913 | 13.7441 | 13.99418 | 35 | 35 | 35 | 40.4 | 8.75E+08 |
| 18.08301 | 25.43584 | 17.85745 | 61 | 61 | 61 | 22.4 | 8.70E+08 |
| 11.11607 | 19.22235 | 13.47526 | 10 | 8 | 8 | 20.7 | 8.67E+08 |

| | | | | | | | |
|---|---|---|---|---|---|---|---|
| 12.52214 | 17.72287 | 13.42515 | 9 | 9 | 9 | 25.6 | 8.61E+08 |
| 13.52943 | 14.72046 | 13.18162 | 39 | 39 | 34 | 30.6 | 8.47E+08 |
| 19.22748 | 13.08469 | 15.35435 | 14 | 14 | 14 | 42.5 | 8.41E+08 |
| 19.89828 | 22.18982 | 22.84197 | 2 | 2 | 2 | 8.2 | 8.36E+08 |
| 16.98025 | 21.34169 | 17.89071 | 10 | 10 | 10 | 33.4 | 8.34E+08 |
| 12.42642 | 12.68758 | 11.48635 | 21 | 21 | 21 | 36.1 | 8.31E+08 |
| 11.95245 | 13.78376 | 11.00775 | 10 | 10 | 9 | 34.7 | 8.21E+08 |
| 10.79903 | 13.3319 | 14.30549 | 11 | 11 | 11 | 50.1 | 8.20E+08 |
| 17.93145 | 19.28062 | 16.14079 | 13 | 13 | 13 | 47.9 | 8.18E+08 |
| 14.86887 | 15.50491 | 14.77839 | 40 | 40 | 36 | 25 | 8.09E+08 |
| 13.65107 | 20.50106 | 17.47199 | 13 | 13 | 13 | 28.2 | 8.06E+08 |
| 11.04596 | 12.69415 | 11.84687 | 11 | 10 | 10 | 29.4 | 7.98E+08 |
| 13.95186 | 19.06779 | 17.19806 | 26 | 26 | 26 | 33.2 | 7.95E+08 |
| 15.58349 | 19.0104 | 16.05835 | 24 | 23 | 23 | 32.9 | 7.73E+08 |
| 17.17358 | 19.54732 | 17.55487 | 11 | 11 | 11 | 18.1 | 7.70E+08 |
| 11.9863 | 20.47358 | 12.28293 | 26 | 26 | 26 | 29.1 | 7.69E+08 |
| 13.34639 | 12.26475 | 16.65088 | 30 | 30 | 30 | 41.8 | 7.67E+08 |
| 16.3338 | 23.25422 | 18.02751 | 19 | 19 | 19 | 58.2 | 7.64E+08 |
| 18.16742 | 21.56001 | 18.07885 | 7 | 7 | 7 | 66.1 | 7.62E+08 |
| 17.74852 | 17.12121 | 17.4351 | 39 | 39 | 39 | 23.5 | 7.60E+08 |
| 14.00685 | 15.27747 | 14.59234 | 16 | 16 | 16 | 39.8 | 7.58E+08 |
| 15.7621 | 12.66236 | 11.43932 | 19 | 19 | 19 | 31.9 | 7.53E+08 |
| 11.7252 | 14.34394 | 13.53041 | 27 | 27 | 27 | 38.3 | 7.53E+08 |
| 13.97773 | 13.71091 | 13.84107 | 25 | 25 | 25 | 37.2 | 7.52E+08 |
| 12.53985 | 13.83506 | 17.48259 | 30 | 30 | 30 | 40.9 | 7.50E+08 |
| 23.35971 | 24.454 | 23.12425 | 2 | 2 | 2 | 23.5 | 7.50E+08 |
| 12.86192 | 14.94901 | 12.53807 | 15 | 15 | 11 | 39.8 | 7.46E+08 |
| 19.86543 | 21.64205 | 20.27833 | 10 | 10 | 10 | 33.3 | 7.44E+08 |
| 12.9329 | 16.78558 | 15.12242 | 35 | 35 | 35 | 32.7 | 7.44E+08 |
| 11.8217 | 19.93111 | 16.4767 | 15 | 15 | 13 | 44.2 | 7.35E+08 |
| 26.57066 | 22.52203 | 26.83702 | 9 | 9 | 9 | 19.5 | 7.26E+08 |
| 15.89503 | 15.93196 | 12.01946 | 11 | 11 | 11 | 28.2 | 7.21E+08 |
| 15.0228 | 21.2619 | 16.72614 | 17 | 14 | 14 | 43.8 | 7.17E+08 |
| 11.4922 | 14.4486 | 12.62408 | 21 | 21 | 21 | 29 | 7.14E+08 |
| 19.31104 | 20.03416 | 19.2163 | 19 | 19 | 19 | 21.6 | 7.08E+08 |
| 20.02567 | 18.79586 | 12.64252 | 22 | 22 | 21 | 30.9 | 7.08E+08 |
| 11.32165 | 15.20293 | 14.80277 | 20 | 20 | 20 | 28.6 | 7.08E+08 |
| 22.17118 | 22.47341 | 20.47843 | 11 | 11 | 11 | 43.9 | 7.02E+08 |
| 17.24943 | 19.52684 | 16.5446 | 3 | 3 | 3 | 4.5 | 7.01E+08 |
| 11.35007 | 17.10697 | 16.48014 | 23 | 23 | 23 | 51.2 | 6.98E+08 |
| 20.64447 | 24.05275 | 21.63781 | 14 | 14 | 14 | 86.8 | 6.84E+08 |
| 16.65172 | 23.66861 | 16.38564 | 24 | 24 | 21 | 35.4 | 6.83E+08 |
| 18.08405 | 22.45051 | 18.03848 | 19 | 19 | 19 | 23.4 | 6.82E+08 |
| 10.42093 | 14.38869 | 14.28793 | 7 | 7 | 7 | 33.4 | 6.81E+08 |
| 13.13012 | 13.10028 | 12.70503 | 11 | 11 | 11 | 18.7 | 6.75E+08 |
| 11.01752 | 15.09861 | 13.16826 | 21 | 21 | 21 | 36.8 | 6.73E+08 |
| 20.90264 | 23.06502 | 20.00402 | 9 | 9 | 9 | 34.2 | 6.71E+08 |
| 12.05056 | 11.44048 | 12.84404 | 15 | 15 | 15 | 37.7 | 6.71E+08 |

| | | | | | | | |
|---|---|---|---|---|---|---|---|
| 18.03047 | 20.52707 | 18.47354 | 13 | 13 | 13 | 31.4 | 6.69E+08 |
| 19.47187 | 23.68603 | 19.01808 | 11 | 11 | 11 | 57.3 | 6.67E+08 |
| 16.39918 | 13.79726 | 12.83762 | 21 | 21 | 21 | 47.7 | 6.66E+08 |
| 11.92754 | 16.73952 | 10.38832 | 29 | 29 | 16 | 25.5 | 6.65E+08 |
| 20.36773 | 22.83086 | 22.9174 | 27 | 27 | 26 | 24.8 | 6.63E+08 |
| 17.46284 | 20.34743 | 21.12666 | 28 | 28 | 28 | 41.9 | 6.63E+08 |
| 15.70782 | 14.2916 | 13.23554 | 12 | 12 | 12 | 47.7 | 6.56E+08 |
| 22.86965 | 24.98681 | 21.98266 | 2 | 2 | 2 | 39.3 | 6.52E+08 |
| 10.83443 | 14.24429 | 13.49069 | 7 | 6 | 6 | 32 | 6.51E+08 |
| 15.28851 | 20.58463 | 17.85386 | 11 | 11 | 11 | 33.9 | 6.51E+08 |
| 11.36075 | 13.92457 | 11.51713 | 5 | 5 | 5 | 13.5 | 6.50E+08 |
| 13.57814 | 11.67552 | 14.61534 | 27 | 27 | 27 | 19.4 | 6.45E+08 |
| 13.66711 | 14.03523 | 12.72494 | 35 | 30 | 30 | 28.5 | 6.44E+08 |
| 13.45185 | 15.88998 | 14.20029 | 25 | 25 | 25 | 60.6 | 6.39E+08 |
| 15.47637 | 16.26266 | 12.81536 | 13 | 13 | 13 | 49.1 | 6.37E+08 |
| 17.64075 | 24.6774 | 16.362 | 12 | 12 | 8 | 51.8 | 6.35E+08 |
| 11.24211 | 15.036 | 14.48111 | 24 | 24 | 18 | 31.6 | 6.32E+08 |
| 14.1309 | 17.54163 | 13.76966 | 13 | 13 | 13 | 42.8 | 6.26E+08 |
| 12.12348 | 13.12806 | 12.40719 | 15 | 15 | 15 | 56.5 | 6.25E+08 |
| 19.94521 | 20.88569 | 20.19748 | 21 | 20 | 20 | 41.7 | 6.21E+08 |
| 13.27263 | 15.98764 | 12.75967 | 10 | 10 | 10 | 33.5 | 6.15E+08 |
| 15.49022 | 15.7272 | 12.69397 | 12 | 12 | 12 | 33.6 | 6.10E+08 |
| 16.98237 | 15.71829 | 16.16044 | 18 | 18 | 18 | 49.4 | 6.09E+08 |
| 15.22506 | 19.57426 | 17.92463 | 4 | 4 | 4 | 34.1 | 6.08E+08 |
| 11.66043 | 12.50767 | 18.40694 | 12 | 12 | 12 | 37.2 | 6.07E+08 |
| 14.92532 | 13.90255 | 12.57685 | 27 | 27 | 27 | 25.7 | 6.06E+08 |
| 20.5976 | 21.60725 | 20.96317 | 9 | 9 | 9 | 28.8 | 6.06E+08 |
| 16.35001 | 15.48687 | 15.77777 | 27 | 27 | 27 | 30.5 | 6.00E+08 |
| 10.47004 | 15.8851 | 11.55558 | 10 | 10 | 7 | 23 | 5.97E+08 |
| 17.12374 | 18.37958 | 10.96043 | 29 | 29 | 29 | 40.7 | 5.93E+08 |
| 11.54136 | 14.21394 | 12.92678 | 16 | 16 | 4 | 46.2 | 5.91E+08 |
| 12.3236 | 16.96961 | 13.73355 | 25 | 25 | 25 | 28.2 | 5.89E+08 |
| 13.2661 | 15.73778 | 16.33158 | 14 | 14 | 14 | 42.6 | 5.89E+08 |
| 13.06283 | 13.06126 | 18.12055 | 16 | 16 | 16 | 50.4 | 5.88E+08 |
| 13.93537 | 13.13838 | 13.60479 | 12 | 12 | 12 | 34.8 | 5.87E+08 |
| 18.31968 | 22.3541 | 15.72302 | 12 | 12 | 12 | 40.1 | 5.85E+08 |
| 13.39044 | 17.58659 | 14.64532 | 21 | 21 | 21 | 27.4 | 5.85E+08 |
| 13.62735 | 17.21181 | 11.30181 | 13 | 13 | 13 | 34.1 | 5.83E+08 |
| 13.84745 | 15.15853 | 13.75018 | 19 | 19 | 17 | 38.6 | 5.83E+08 |
| 12.22339 | 14.17633 | 11.30482 | 18 | 18 | 18 | 30.9 | 5.78E+08 |
| 14.25244 | 12.91603 | 12.47724 | 23 | 23 | 23 | 28.3 | 5.75E+08 |
| 16.98304 | 14.85037 | 12.31838 | 15 | 15 | 15 | 49.4 | 5.74E+08 |
| 11.84556 | 11.1383 | 15.18038 | 10 | 10 | 10 | 55.8 | 5.72E+08 |
| 11.22459 | 19.66028 | 10.91956 | 9 | 9 | 9 | 52.8 | 5.70E+08 |
| 15.07138 | 14.94297 | 18.74924 | 19 | 19 | 19 | 25.4 | 5.64E+08 |
| 14.89903 | 15.56772 | 16.70968 | 33 | 33 | 32 | 28.3 | 5.63E+08 |
| 11.40235 | 14.01304 | 10.43504 | 20 | 19 | 19 | 19.4 | 5.61E+08 |
| 18.19378 | 19.09841 | 18.20333 | 4 | 4 | 4 | 26.8 | 5.60E+08 |

| | | | | | | | |
|---|---|---|---|---|---|---|---|
| 11.97387 | 14.34587 | 11.25159 | 7 | 7 | 2 | 32.1 | 5.60E+08 |
| 18.37704 | 22.65661 | 22.30991 | 6 | 6 | 3 | 18.7 | 5.58E+08 |
| 19.19302 | 16.50987 | 19.3459 | 27 | 27 | 27 | 38.6 | 5.57E+08 |
| 12.22883 | 16.03071 | 11.33221 | 25 | 25 | 25 | 25.1 | 5.57E+08 |
| 14.23684 | 15.47792 | 14.30278 | 10 | 10 | 10 | 29.5 | 5.54E+08 |
| 19.02645 | 22.08154 | 20.67107 | 16 | 16 | 8 | 27.6 | 5.53E+08 |
| 23.905 | 25.0814 | 24.46669 | 4 | 4 | 4 | 29.7 | 5.52E+08 |
| 13.08515 | 12.43194 | 10.63414 | 17 | 17 | 17 | 27.2 | 5.49E+08 |
| 14.82918 | 16.45722 | 13.89283 | 18 | 18 | 18 | 32.8 | 5.47E+08 |
| 13.47206 | 14.25392 | 13.05269 | 27 | 27 | 27 | 36.1 | 5.47E+08 |
| 12.12874 | 13.18038 | 12.16249 | 22 | 22 | 22 | 38.2 | 5.42E+08 |
| 14.60687 | 15.7306 | 14.7151 | 21 | 21 | 21 | 39.5 | 5.41E+08 |
| 14.41825 | 18.71629 | 13.91756 | 21 | 21 | 21 | 28.6 | 5.41E+08 |
| 16.44736 | 16.16577 | 16.50044 | 17 | 17 | 17 | 39.5 | 5.41E+08 |
| 11.69033 | 13.84523 | 11.39897 | 18 | 18 | 18 | 33.7 | 5.40E+08 |
| 15.68467 | 19.67206 | 10.86552 | 26 | 26 | 26 | 34.1 | 5.38E+08 |
| 14.20006 | 16.52751 | 12.7535 | 7 | 7 | 3 | 25.1 | 5.34E+08 |
| 14.20404 | 15.17979 | 13.18593 | 12 | 12 | 12 | 65.3 | 5.33E+08 |
| 11.61297 | 16.48413 | 11.87247 | 10 | 10 | 10 | 42.5 | 5.32E+08 |
| 22.47006 | 22.08291 | 22.27813 | 8 | 8 | 8 | 9.1 | 5.22E+08 |
| 10.73721 | 18.63101 | 11.22287 | 16 | 16 | 16 | 29.2 | 5.20E+08 |
| 13.10232 | 13.73517 | 13.78688 | 20 | 20 | 20 | 37.5 | 5.17E+08 |
| 11.61842 | 14.05016 | 11.89078 | 12 | 12 | 12 | 46.4 | 5.15E+08 |
| 11.4608 | 12.44623 | 10.93539 | 14 | 14 | 14 | 41.1 | 5.08E+08 |
| 11.35076 | 14.82197 | 13.3062 | 35 | 35 | 35 | 19.9 | 4.99E+08 |
| 11.79055 | 18.89888 | 21.10386 | 21 | 21 | 21 | 45.7 | 4.97E+08 |
| 18.81735 | 24.13907 | 19.1167 | 14 | 14 | 14 | 48.7 | 4.96E+08 |
| 14.55258 | 19.53353 | 13.16335 | 8 | 8 | 8 | 42.4 | 4.93E+08 |
| 12.97322 | 15.47788 | 12.76698 | 29 | 29 | 29 | 36.9 | 4.88E+08 |
| 11.94979 | 13.93765 | 11.53538 | 18 | 18 | 18 | 54.4 | 4.87E+08 |
| 13.07484 | 17.78558 | 11.03841 | 26 | 26 | 26 | 26.6 | 4.87E+08 |
| 19.63793 | 19.39611 | 15.54055 | 12 | 12 | 12 | 29.7 | 4.87E+08 |
| 16.11502 | 16.78098 | 16.34658 | 12 | 12 | 11 | 33.5 | 4.86E+08 |
| 12.31708 | 17.77043 | 12.55487 | 16 | 16 | 16 | 41.1 | 4.85E+08 |
| 10.55371 | 26.91639 | 19.22376 | 9 | 9 | 9 | 67.5 | 4.82E+08 |
| 16.24741 | 18.25091 | 17.05696 | 4 | 3 | 3 | 31 | 4.81E+08 |
| 14.59368 | 15.92379 | 14.57152 | 31 | 31 | 31 | 26.3 | 4.78E+08 |
| 10.6695 | 13.12797 | 15.96522 | 17 | 17 | 17 | 22.6 | 4.78E+08 |
| 11.93857 | 17.32922 | 12.22154 | 15 | 15 | 15 | 37.1 | 4.75E+08 |
| 11.47578 | 16.96477 | 19.22232 | 19 | 19 | 19 | 23.2 | 4.74E+08 |
| 11.64382 | 15.8528 | 14.19845 | 11 | 11 | 11 | 29.5 | 4.74E+08 |
| 13.47358 | 13.68595 | 13.36701 | 22 | 22 | 22 | 21.4 | 4.72E+08 |
| 12.23455 | 11.85436 | 13.03806 | 33 | 33 | 33 | 23.1 | 4.72E+08 |
| 11.62613 | 14.42282 | 11.62628 | 22 | 22 | 22 | 26.7 | 4.72E+08 |
| 16.62357 | 11.43059 | 16.26224 | 25 | 25 | 25 | 28.3 | 4.68E+08 |
| 12.12098 | 15.29282 | 11.89835 | 10 | 10 | 10 | 35.1 | 4.68E+08 |
| 16.80675 | 13.18016 | 16.78021 | 17 | 17 | 17 | 12.4 | 4.67E+08 |
| 13.44825 | 14.64425 | 14.31777 | 16 | 16 | 13 | 28.5 | 4.65E+08 |

| | | | | | | | |
|---|---|---|---|---|---|---|---|
| 11.60329 | 13.56673 | 11.55983 | 12 | 12 | 12 | 72.5 | 4.61E+08 |
| 12.50931 | 15.75557 | 12.79324 | 25 | 25 | 25 | 24.1 | 4.61E+08 |
| 12.33347 | 14.91462 | 12.28002 | 8 | 8 | 8 | 44.9 | 4.60E+08 |
| 22.67549 | 23.43644 | 23.35165 | 12 | 12 | 12 | 28.3 | 4.59E+08 |
| 17.42119 | 15.2688 | 18.01471 | 29 | 29 | 29 | 19.6 | 4.58E+08 |
| 15.71169 | 18.46905 | 16.69602 | 14 | 14 | 14 | 36.8 | 4.57E+08 |
| 11.20788 | 18.7339 | 14.18637 | 9 | 9 | 9 | 26.6 | 4.56E+08 |
| 20.1728 | 13.69681 | 19.24651 | 30 | 30 | 30 | 18 | 4.53E+08 |
| 12.61576 | 14.05511 | 12.86933 | 21 | 21 | 20 | 32.5 | 4.51E+08 |
| 12.35697 | 16.40458 | 11.01289 | 19 | 19 | 19 | 37.7 | 4.49E+08 |
| 12.46183 | 16.03131 | 10.60532 | 14 | 14 | 14 | 46.3 | 4.48E+08 |
| 11.96634 | 20.43329 | 11.56247 | 21 | 21 | 21 | 35.4 | 4.47E+08 |
| 17.88959 | 15.3348 | 16.44616 | 21 | 21 | 21 | 21 | 4.47E+08 |
| 11.57263 | 16.50185 | 11.32004 | 13 | 13 | 13 | 53.6 | 4.46E+08 |
| 12.1852 | 14.93231 | 13.64506 | 29 | 29 | 28 | 26.9 | 4.43E+08 |
| 14.98157 | 14.74257 | 12.2729 | 19 | 19 | 19 | 26.9 | 4.40E+08 |
| 12.5768 | 18.98904 | 18.30759 | 14 | 14 | 14 | 21.4 | 4.40E+08 |
| 16.55869 | 18.2837 | 16.16107 | 21 | 21 | 21 | 43.4 | 4.40E+08 |
| 10.75873 | 17.22449 | 10.56258 | 12 | 12 | 12 | 46.7 | 4.40E+08 |
| 15.28388 | 12.97004 | 13.51193 | 23 | 23 | 23 | 27.5 | 4.38E+08 |
| 14.10222 | 17.02929 | 15.69163 | 16 | 16 | 16 | 37 | 4.38E+08 |
| 14.60877 | 16.04446 | 13.5813 | 22 | 22 | 22 | 17.8 | 4.38E+08 |
| 12.3888 | 12.97023 | 18.86181 | 5 | 5 | 5 | 52.7 | 4.37E+08 |
| 21.02763 | 22.49599 | 22.40691 | 5 | 3 | 3 | 31.2 | 4.37E+08 |
| 12.42305 | 17.3699 | 12.91715 | 19 | 19 | 19 | 26.7 | 4.35E+08 |
| 25.53872 | 25.75387 | 25.44199 | 9 | 9 | 9 | 6.6 | 4.34E+08 |
| 11.48142 | 26.92184 | 18.2491 | 10 | 10 | 10 | 70.6 | 4.33E+08 |
| 12.49628 | 19.56471 | 21.19328 | 15 | 15 | 13 | 25.6 | 4.32E+08 |
| 12.74062 | 16.46183 | 14.86452 | 30 | 30 | 29 | 28.5 | 4.27E+08 |
| 18.01634 | 18.54197 | 18.41514 | 11 | 11 | 11 | 38.1 | 4.27E+08 |
| 17.39916 | 19.17555 | 17.21769 | 7 | 5 | 5 | 20.6 | 4.26E+08 |
| 16.94809 | 16.15108 | 13.43788 | 8 | 8 | 8 | 40.2 | 4.25E+08 |
| 17.20801 | 19.45836 | 17.90624 | 9 | 7 | 6 | 19.2 | 4.23E+08 |
| 23.79719 | 23.97299 | 24.08359 | 9 | 9 | 9 | 35.2 | 4.20E+08 |
| 18.87054 | 19.17309 | 19.63952 | 22 | 22 | 14 | 51.2 | 4.20E+08 |
| 10.37791 | 17.57319 | 12.37556 | 15 | 15 | 15 | 42.7 | 4.18E+08 |
| 15.86104 | 17.07984 | 15.87726 | 19 | 19 | 19 | 42.9 | 4.18E+08 |
| 15.42098 | 15.72401 | 15.77283 | 14 | 14 | 14 | 41.3 | 4.18E+08 |
| 15.73426 | 17.7184 | 12.37206 | 13 | 13 | 13 | 36.4 | 4.18E+08 |
| 15.69428 | 19.06092 | 16.60103 | 15 | 15 | 15 | 25.4 | 4.15E+08 |
| 15.71459 | 12.58176 | 14.06643 | 11 | 11 | 11 | 24.7 | 4.14E+08 |
| 13.96507 | 20.0434 | 11.80619 | 3 | 3 | 3 | 27 | 4.14E+08 |
| 16.74465 | 17.39891 | 17.25949 | 21 | 21 | 19 | 31.6 | 4.13E+08 |
| 20.05643 | 21.3424 | 19.81328 | 3 | 3 | 3 | 29.3 | 4.10E+08 |
| 22.78377 | 22.87442 | 23.12308 | 6 | 6 | 6 | 18.2 | 4.07E+08 |
| 16.23616 | 15.24165 | 17.0116 | 13 | 13 | 13 | 29.9 | 4.01E+08 |
| 11.45722 | 16.53652 | 11.2181 | 15 | 15 | 14 | 18.5 | 4.01E+08 |
| 11.67877 | 18.3071 | 12.35851 | 15 | 15 | 15 | 33.8 | 3.97E+08 |

| | | | | | | | |
|---|---|---|---|---|---|---|---|
| 13.53584 | 13.71317 | 14.23032 | 26 | 26 | 26 | 24.7 | 3.95E+08 |
| 12.58322 | 17.02799 | 12.20877 | 10 | 10 | 10 | 39.6 | 3.95E+08 |
| 12.70154 | 14.59304 | 12.25739 | 23 | 23 | 23 | 28.7 | 3.93E+08 |
| 12.47653 | 12.17835 | 12.64328 | 26 | 26 | 16 | 26.2 | 3.93E+08 |
| 11.71557 | 14.71377 | 11.79359 | 17 | 17 | 17 | 36.7 | 3.93E+08 |
| 20.78588 | 21.4227 | 21.1432 | 4 | 4 | 4 | 18.8 | 3.93E+08 |
| 16.50003 | 19.25659 | 10.80096 | 9 | 9 | 8 | 32.4 | 3.92E+08 |
| 13.07257 | 15.79198 | 12.00032 | 18 | 18 | 18 | 29.8 | 3.90E+08 |
| 12.70037 | 16.83133 | 13.51911 | 10 | 10 | 10 | 43.1 | 3.88E+08 |
| 15.08746 | 14.86191 | 10.83326 | 11 | 11 | 11 | 24.4 | 3.85E+08 |
| 11.95049 | 14.79344 | 16.97231 | 18 | 18 | 18 | 35 | 3.85E+08 |
| 17.48684 | 15.60154 | 11.84961 | 17 | 17 | 17 | 23.9 | 3.85E+08 |
| 18.61418 | 21.63032 | 20.22447 | 23 | 14 | 12 | 50.4 | 3.83E+08 |
| 21.26911 | 23.12011 | 22.51372 | 5 | 5 | 5 | 35.2 | 3.82E+08 |
| 12.86535 | 13.45053 | 13.27477 | 10 | 10 | 10 | 41 | 3.81E+08 |
| 10.96038 | 12.91303 | 11.49738 | 10 | 10 | 10 | 15.8 | 3.79E+08 |
| 11.91605 | 15.31603 | 10.43816 | 13 | 13 | 13 | 41.5 | 3.79E+08 |
| 11.02951 | 15.78887 | 13.64504 | 16 | 16 | 16 | 26.3 | 3.77E+08 |
| 15.99841 | 16.93168 | 16.33645 | 24 | 23 | 23 | 29.1 | 3.71E+08 |
| 21.65508 | 22.46533 | 22.17405 | 13 | 13 | 13 | 34 | 3.71E+08 |
| 12.21785 | 20.92579 | 17.84922 | 20 | 7 | 7 | 52.3 | 3.70E+08 |
| 17.00536 | 19.80022 | 18.28392 | 9 | 9 | 9 | 31.1 | 3.67E+08 |
| 16.11744 | 17.59987 | 14.09259 | 11 | 11 | 11 | 30.8 | 3.67E+08 |
| 17.17143 | 15.78341 | 17.76442 | 7 | 7 | 7 | 43.6 | 3.66E+08 |
| 18.75502 | 19.46364 | 19.05144 | 8 | 8 | 8 | 49.1 | 3.66E+08 |
| 11.37969 | 13.20378 | 10.78972 | 6 | 6 | 6 | 43.7 | 3.66E+08 |
| 15.84176 | 15.79292 | 12.2514 | 18 | 18 | 18 | 21.7 | 3.65E+08 |
| 12.67727 | 20.04233 | 12.22222 | 10 | 10 | 9 | 40.8 | 3.64E+08 |
| 11.75861 | 16.30453 | 11.82424 | 22 | 22 | 22 | 31.1 | 3.64E+08 |
| 20.1928 | 22.40598 | 19.65279 | 6 | 5 | 5 | 64.8 | 3.60E+08 |
| 11.37806 | 15.81007 | 11.20122 | 13 | 13 | 13 | 38.7 | 3.60E+08 |
| 13.50819 | 13.22985 | 12.30492 | 20 | 18 | 18 | 31.9 | 3.60E+08 |
| 12.57307 | 13.26563 | 12.09672 | 3 | 3 | 3 | 25.7 | 3.59E+08 |
| 24.60286 | 12.93814 | 11.4508 | 9 | 8 | 8 | 7.2 | 3.58E+08 |
| 13.29514 | 19.96776 | 12.1682 | 9 | 9 | 9 | 52.7 | 3.57E+08 |
| 20.55899 | 21.23392 | 18.92833 | 17 | 17 | 17 | 13.4 | 3.57E+08 |
| 12.02634 | 14.09391 | 18.73685 | 15 | 15 | 15 | 54.2 | 3.57E+08 |
| 10.69395 | 16.90089 | 14.57009 | 16 | 16 | 16 | 30.6 | 3.56E+08 |
| 17.05378 | 17.44354 | 16.81353 | 9 | 9 | 9 | 20 | 3.55E+08 |
| 13.98495 | 16.03045 | 14.21803 | 12 | 12 | 12 | 37.9 | 3.55E+08 |
| 10.81498 | 13.0287 | 12.45084 | 22 | 22 | 22 | 15.5 | 3.54E+08 |
| 10.81198 | 18.85583 | 13.12574 | 7 | 7 | 2 | 12.6 | 3.52E+08 |
| 11.37021 | 19.2023 | 17.36359 | 14 | 14 | 14 | 43 | 3.51E+08 |
| 12.92481 | 14.99722 | 15.3549 | 29 | 29 | 29 | 21.2 | 3.49E+08 |
| 13.33617 | 17.23285 | 12.49646 | 20 | 20 | 20 | 29.1 | 3.49E+08 |
| 11.0337 | 16.20562 | 13.16579 | 21 | 21 | 21 | 24.4 | 3.46E+08 |
| 13.84471 | 14.81047 | 12.77186 | 19 | 19 | 19 | 21.8 | 3.42E+08 |
| 12.41901 | 16.47596 | 11.05849 | 19 | 19 | 19 | 22.7 | 3.41E+08 |

| | | | | | | | |
|---|---|---|---|---|---|---|---|
| 10.60533 | 18.52868 | 10.24885 | 17 | 17 | 17 | 37.3 | 3.41E+08 |
| 18.32202 | 22.51128 | 17.25072 | 10 | 10 | 9 | 43.7 | 3.39E+08 |
| 21.23298 | 21.12937 | 21.20741 | 18 | 18 | 18 | 17.5 | 3.38E+08 |
| 11.56358 | 13.04689 | 10.57343 | 23 | 23 | 10 | 16.4 | 3.37E+08 |
| 12.10324 | 17.69125 | 16.13563 | 16 | 16 | 16 | 8.3 | 3.37E+08 |
| 12.42073 | 17.01585 | 15.21117 | 8 | 8 | 1 | 64.2 | 3.34E+08 |
| 12.12858 | 17.28735 | 13.18903 | 11 | 11 | 11 | 42.8 | 3.34E+08 |
| 24.26871 | 23.96261 | 23.97009 | 12 | 12 | 12 | 33.8 | 3.34E+08 |
| 16.27292 | 17.31708 | 13.50097 | 18 | 18 | 18 | 24.5 | 3.32E+08 |
| 18.8363 | 16.52532 | 19.25142 | 15 | 15 | 15 | 23.5 | 3.31E+08 |
| 18.97477 | 20.96423 | 11.40665 | 9 | 9 | 9 | 38 | 3.30E+08 |
| 13.26634 | 17.89569 | 12.13876 | 26 | 26 | 26 | 26.2 | 3.29E+08 |
| 12.89205 | 17.31858 | 15.85309 | 18 | 6 | 5 | 49.3 | 3.29E+08 |
| 11.38075 | 16.51317 | 14.0676 | 15 | 9 | 9 | 57.5 | 3.29E+08 |
| 22.8486 | 23.11399 | 23.18794 | 10 | 10 | 10 | 32.1 | 3.29E+08 |
| 15.20163 | 15.9514 | 14.22302 | 20 | 20 | 20 | 44.1 | 3.27E+08 |
| 20.48326 | 19.88545 | 20.61207 | 20 | 20 | 20 | 28.6 | 3.26E+08 |
| 16.56035 | 14.14354 | 12.28396 | 11 | 11 | 11 | 28 | 3.26E+08 |
| 21.74874 | 24.26849 | 20.63076 | 6 | 6 | 6 | 13.1 | 3.25E+08 |
| 16.50499 | 16.78021 | 16.68223 | 26 | 26 | 26 | 25.3 | 3.23E+08 |
| 13.68166 | 18.62392 | 12.53663 | 4 | 4 | 4 | 31.1 | 3.21E+08 |
| 12.44164 | 12.74209 | 13.04968 | 12 | 12 | 12 | 50.8 | 3.20E+08 |
| 19.15046 | 16.79335 | 20.11388 | 9 | 9 | 9 | 31.8 | 3.20E+08 |
| 14.66245 | 16.10259 | 14.3768 | 10 | 10 | 10 | 33.2 | 3.18E+08 |
| 12.04123 | 11.82683 | 12.15308 | 21 | 21 | 21 | 21 | 3.18E+08 |
| 23.65349 | 23.7543 | 22.32814 | 6 | 6 | 6 | 25.6 | 3.17E+08 |
| 12.12668 | 19.37454 | 12.65304 | 4 | 4 | 4 | 40.7 | 3.17E+08 |
| 18.35651 | 13.35127 | 13.38986 | 19 | 19 | 19 | 26.8 | 3.16E+08 |
| 10.99554 | 19.4502 | 12.20632 | 11 | 11 | 11 | 15.2 | 3.15E+08 |
| 10.98113 | 13.19455 | 10.40621 | 9 | 9 | 9 | 48.7 | 3.14E+08 |
| 12.01895 | 14.26123 | 11.31788 | 7 | 7 | 7 | 19.7 | 3.14E+08 |
| 13.29878 | 13.07312 | 14.89907 | 11 | 11 | 11 | 41.3 | 3.14E+08 |
| 15.99932 | 18.87228 | 18.76914 | 10 | 10 | 10 | 43.5 | 3.13E+08 |
| 23.46164 | 24.01018 | 23.33838 | 4 | 4 | 4 | 21.9 | 3.10E+08 |
| 12.3668 | 16.17623 | 12.32284 | 16 | 16 | 16 | 23.1 | 3.09E+08 |
| 9.780993 | 20.16165 | 13.09531 | 16 | 16 | 16 | 26.2 | 3.07E+08 |
| 13.07678 | 26.32922 | 15.85426 | 2 | 2 | 2 | 25.8 | 3.05E+08 |
| 18.6352 | 19.05849 | 15.34966 | 15 | 15 | 15 | 22.9 | 3.04E+08 |
| 12.01111 | 15.55842 | 12.28897 | 17 | 17 | 17 | 13.6 | 3.02E+08 |
| 12.14499 | 16.85781 | 12.90633 | 8 | 8 | 8 | 35.8 | 3.02E+08 |
| 19.0351 | 13.42102 | 14.03601 | 8 | 8 | 8 | 25.9 | 3.01E+08 |
| 14.4485 | 16.29878 | 13.26683 | 20 | 20 | 20 | 21 | 2.97E+08 |
| 19.8828 | 17.85678 | 10.66925 | 12 | 12 | 12 | 34.8 | 2.95E+08 |
| 18.03848 | 15.9478 | 13.17351 | 16 | 16 | 15 | 23.7 | 2.94E+08 |
| 12.78986 | 15.68387 | 14.36482 | 8 | 8 | 6 | 22.5 | 2.93E+08 |
| 18.66799 | 18.63729 | 18.67238 | 17 | 17 | 17 | 26.2 | 2.92E+08 |
| 12.88315 | 11.91516 | 12.09716 | 11 | 9 | 9 | 32.2 | 2.91E+08 |
| 13.1288 | 13.69116 | 18.10185 | 18 | 18 | 17 | 7.6 | 2.90E+08 |

| | | | | | | | |
|---|---|---|---|---|---|---|---|
| 12.77194 | 11.55564 | 12.71191 | 15 | 15 | 13 | 28.6 | 2.89E+08 |
| 18.03397 | 16.68647 | 13.43412 | 11 | 11 | 11 | 41.9 | 2.88E+08 |
| 13.23022 | 19.37029 | 12.62176 | 12 | 12 | 12 | 39.8 | 2.87E+08 |
| 16.78839 | 20.42234 | 16.46346 | 14 | 14 | 14 | 16.3 | 2.86E+08 |
| 10.36814 | 18.4586 | 18.1921 | 14 | 14 | 14 | 21.1 | 2.84E+08 |
| 11.78189 | 13.72806 | 12.95465 | 19 | 19 | 19 | 13.9 | 2.83E+08 |
| 13.30644 | 13.60509 | 12.36342 | 12 | 12 | 12 | 25.5 | 2.82E+08 |
| 13.85313 | 17.20151 | 11.24047 | 10 | 10 | 4 | 32 | 2.82E+08 |
| 13.89273 | 13.33692 | 13.53296 | 8 | 8 | 8 | 31.6 | 2.81E+08 |
| 20.59887 | 19.77791 | 20.57052 | 19 | 19 | 19 | 30.5 | 2.81E+08 |
| 17.54064 | 19.0665 | 18.01803 | 17 | 17 | 17 | 24.7 | 2.79E+08 |
| 18.44415 | 17.84604 | 12.96476 | 16 | 16 | 16 | 25.6 | 2.78E+08 |
| 12.18886 | 13.3528 | 13.13205 | 15 | 15 | 15 | 36.2 | 2.77E+08 |
| 14.1437 | 14.25444 | 13.99028 | 15 | 15 | 15 | 30.9 | 2.77E+08 |
| 22.93355 | 23.93526 | 23.06776 | 19 | 5 | 3 | 33.2 | 2.76E+08 |
| 21.90205 | 22.86712 | 22.72873 | 5 | 5 | 5 | 20.7 | 2.75E+08 |
| 11.13572 | 17.0898 | 12.31534 | 13 | 13 | 13 | 30.7 | 2.75E+08 |
| 20.07561 | 18.85881 | 20.14706 | 8 | 8 | 6 | 5.6 | 2.75E+08 |
| 15.7402 | 19.24056 | 11.64756 | 6 | 6 | 6 | 30.2 | 2.74E+08 |
| 13.88974 | 15.35669 | 12.91206 | 15 | 15 | 15 | 25.8 | 2.73E+08 |
| 13.441 | 13.75048 | 13.50804 | 5 | 5 | 5 | 29.9 | 2.73E+08 |
| 14.08828 | 17.26215 | 13.39046 | 20 | 20 | 20 | 16.8 | 2.72E+08 |
| 12.59629 | 21.7766 | 12.2847 | 7 | 2 | 2 | 35.4 | 2.71E+08 |
| 12.48824 | 15.71478 | 12.88178 | 10 | 10 | 10 | 28.3 | 2.71E+08 |
| 13.40569 | 15.18843 | 12.73527 | 19 | 5 | 5 | 51.6 | 2.68E+08 |
| 11.71146 | 15.55504 | 11.53941 | 17 | 17 | 15 | 32.6 | 2.68E+08 |
| 22.70299 | 23.6231 | 11.96892 | 7 | 7 | 7 | 14.9 | 2.67E+08 |
| 11.26784 | 16.90266 | 12.80565 | 17 | 17 | 17 | 22.3 | 2.66E+08 |
| 16.08023 | 18.66376 | 17.04025 | 13 | 13 | 13 | 37.2 | 2.65E+08 |
| 12.9326 | 14.85687 | 12.59739 | 23 | 23 | 23 | 24.5 | 2.65E+08 |
| 12.27301 | 14.82302 | 12.2948 | 22 | 22 | 22 | 20.6 | 2.64E+08 |
| 20.99776 | 23.36172 | 20.38121 | 6 | 6 | 6 | 42.8 | 2.64E+08 |
| 15.03169 | 14.059 | 13.64726 | 15 | 15 | 15 | 22.8 | 2.63E+08 |
| 12.23235 | 21.6105 | 12.65438 | 2 | 2 | 2 | 15.1 | 2.62E+08 |
| 11.46301 | 18.8579 | 17.03273 | 11 | 11 | 11 | 31.7 | 2.61E+08 |
| 20.09816 | 20.29804 | 19.73175 | 12 | 12 | 12 | 19.9 | 2.58E+08 |
| 11.62353 | 13.73987 | 17.64885 | 15 | 15 | 14 | 19.7 | 2.58E+08 |
| 10.69823 | 17.49076 | 10.9819 | 19 | 19 | 19 | 30.7 | 2.58E+08 |
| 12.70434 | 12.60745 | 13.574 | 25 | 17 | 15 | 42 | 2.57E+08 |
| 12.07455 | 17.70941 | 11.70922 | 11 | 11 | 11 | 35 | 2.57E+08 |
| 10.97621 | 15.50618 | 11.53732 | 9 | 9 | 9 | 23 | 2.57E+08 |
| 18.55093 | 11.80884 | 11.78627 | 16 | 14 | 14 | 32.3 | 2.57E+08 |
| 14.7158 | 14.09688 | 13.55447 | 10 | 10 | 10 | 31.9 | 2.56E+08 |
| 16.55223 | 17.6933 | 16.6622 | 8 | 8 | 5 | 56.7 | 2.55E+08 |
| 20.03591 | 11.94997 | 20.16717 | 7 | 2 | 2 | 32.1 | 2.55E+08 |
| 12.70114 | 18.60447 | 12.04249 | 7 | 7 | 6 | 36.5 | 2.54E+08 |
| 17.11837 | 20.27583 | 11.63194 | 4 | 4 | 4 | 34.1 | 2.53E+08 |
| 12.15541 | 19.65032 | 12.6265 | 8 | 8 | 5 | 18.8 | 2.50E+08 |

| | | | | | | | |
|---|---|---|---|---|---|---|---|
| 12.16991 | 12.36078 | 12.43214 | 17 | 17 | 17 | 28.7 | 2.49E+08 |
| 16.25292 | 14.7909 | 11.90208 | 12 | 12 | 12 | 28.9 | 2.48E+08 |
| 10.0481 | 13.38887 | 16.94467 | 5 | 5 | 5 | 22.9 | 2.47E+08 |
| 21.16447 | 21.67137 | 21.6413 | 9 | 9 | 9 | 16.3 | 2.47E+08 |
| 11.91645 | 13.60328 | 13.38129 | 12 | 12 | 12 | 32.2 | 2.46E+08 |
| 11.96445 | 13.87117 | 17.71129 | 18 | 18 | 18 | 21.8 | 2.45E+08 |
| 17.3265 | 15.09857 | 17.98499 | 4 | 4 | 4 | 41.9 | 2.44E+08 |
| 18.9356 | 16.66386 | 18.78807 | 15 | 15 | 15 | 19.1 | 2.43E+08 |
| 20.68731 | 20.9612 | 17.77301 | 14 | 14 | 14 | 30.4 | 2.43E+08 |
| 12.88348 | 14.04795 | 17.55225 | 12 | 12 | 12 | 16.4 | 2.43E+08 |
| 19.99466 | 20.74559 | 20.51258 | 8 | 8 | 8 | 17.8 | 2.43E+08 |
| 19.34425 | 16.97242 | 12.26745 | 23 | 23 | 23 | 22.8 | 2.42E+08 |
| 15.33525 | 15.35903 | 12.03409 | 20 | 20 | 20 | 14.6 | 2.40E+08 |
| 11.90429 | 17.50013 | 12.64693 | 12 | 12 | 12 | 22.1 | 2.40E+08 |
| 14.97338 | 15.21257 | 13.94995 | 11 | 11 | 11 | 5 | 2.39E+08 |
| 12.6953 | 14.51812 | 13.35257 | 12 | 12 | 12 | 32.3 | 2.37E+08 |
| 13.84904 | 19.54721 | 11.37881 | 10 | 10 | 10 | 39.1 | 2.37E+08 |
| 13.54633 | 19.05001 | 10.48442 | 12 | 12 | 9 | 24.6 | 2.36E+08 |
| 17.12979 | 19.47508 | 18.08665 | 4 | 4 | 4 | 40.6 | 2.36E+08 |
| 12.29384 | 14.4384 | 11.9039 | 14 | 14 | 14 | 22.3 | 2.33E+08 |
| 11.91145 | 18.37046 | 12.29286 | 4 | 4 | 4 | 27.1 | 2.31E+08 |
| 12.25036 | 12.08276 | 11.8825 | 6 | 6 | 6 | 23.5 | 2.30E+08 |
| 11.81996 | 15.80642 | 13.283 | 14 | 14 | 14 | 15.1 | 2.30E+08 |
| 13.58856 | 15.99696 | 11.16443 | 12 | 12 | 8 | 12.3 | 2.30E+08 |
| 16.81265 | 19.68779 | 17.75075 | 7 | 7 | 7 | 42.7 | 2.29E+08 |
| 15.1547 | 19.07535 | 12.28618 | 18 | 18 | 18 | 15.1 | 2.28E+08 |
| 12.11618 | 20.15746 | 18.60274 | 7 | 7 | 7 | 25.7 | 2.28E+08 |
| 12.08877 | 16.03881 | 13.12472 | 11 | 11 | 11 | 38.4 | 2.28E+08 |
| 10.81237 | 14.82023 | 13.14522 | 13 | 13 | 13 | 33.1 | 2.27E+08 |
| 12.07253 | 19.84924 | 10.27577 | 8 | 5 | 5 | 26.7 | 2.27E+08 |
| 18.2963 | 16.28361 | 14.97634 | 15 | 15 | 15 | 20 | 2.26E+08 |
| 21.19334 | 23.65895 | 13.44453 | 5 | 4 | 4 | 29.2 | 2.26E+08 |
| 13.17033 | 13.0247 | 17.9783 | 8 | 8 | 8 | 42.9 | 2.26E+08 |
| 16.93157 | 16.44476 | 11.32683 | 16 | 16 | 16 | 28.8 | 2.25E+08 |
| 12.44249 | 23.64625 | 20.51509 | 6 | 5 | 5 | 45.6 | 2.25E+08 |
| 12.57212 | 17.67473 | 13.09009 | 5 | 5 | 5 | 30.5 | 2.25E+08 |
| 14.85973 | 16.36968 | 15.20888 | 13 | 13 | 13 | 37.1 | 2.24E+08 |
| 19.11217 | 14.27919 | 16.5301 | 6 | 6 | 6 | 24 | 2.23E+08 |
| 17.42865 | 20.77555 | 17.75427 | 10 | 6 | 6 | 44.1 | 2.23E+08 |
| 11.9756 | 12.0994 | 15.98444 | 19 | 19 | 19 | 13.2 | 2.22E+08 |
| 18.58009 | 18.32056 | 20.01713 | 6 | 6 | 6 | 16 | 2.21E+08 |
| 12.36096 | 13.75228 | 10.95504 | 7 | 7 | 7 | 20.8 | 2.20E+08 |
| 17.15521 | 16.91788 | 16.70914 | 18 | 18 | 18 | 20.2 | 2.20E+08 |
| 13.80725 | 14.76503 | 10.83107 | 13 | 13 | 13 | 32.2 | 2.20E+08 |
| 12.15427 | 16.12422 | 11.57687 | 13 | 13 | 13 | 19.7 | 2.19E+08 |
| 12.82802 | 16.84009 | 11.19814 | 8 | 8 | 8 | 48 | 2.19E+08 |
| 12.1837 | 14.52901 | 11.00792 | 8 | 8 | 8 | 26.5 | 2.19E+08 |
| 20.75124 | 12.36732 | 13.43498 | 17 | 17 | 17 | 21.6 | 2.17E+08 |

| | | | | | | | |
|---|---|---|---|---|---|---|---|
| 15.3527 | 18.53886 | 16.80952 | 7 | 7 | 7 | 31.9 | 2.16E+08 |
| 15.50544 | 15.56323 | 12.73437 | 9 | 9 | 9 | 22.4 | 2.16E+08 |
| 10.72332 | 13.63898 | 12.3904 | 15 | 15 | 15 | 19.7 | 2.15E+08 |
| 11.54962 | 24.8906 | 12.26753 | 6 | 6 | 6 | 20.1 | 2.15E+08 |
| 14.37923 | 18.69377 | 13.10212 | 16 | 13 | 13 | 25 | 2.14E+08 |
| 16.83306 | 17.38279 | 16.69466 | 7 | 7 | 5 | 18.7 | 2.14E+08 |
| 11.97613 | 13.154 | 12.19992 | 13 | 9 | 9 | 21.9 | 2.13E+08 |
| 12.87676 | 14.8027 | 10.89286 | 6 | 2 | 0 | 35.1 | 2.12E+08 |
| 11.95589 | 19.83424 | 15.42663 | 11 | 11 | 11 | 53.7 | 2.12E+08 |
| 18.40532 | 13.55523 | 13.30153 | 6 | 6 | 6 | 26.9 | 2.12E+08 |
| 19.20729 | 14.98877 | 11.5366 | 11 | 11 | 10 | 26.5 | 2.11E+08 |
| 11.2392 | 12.11569 | 11.79431 | 6 | 6 | 6 | 36.6 | 2.10E+08 |
| 11.63956 | 13.68725 | 12.73115 | 10 | 10 | 10 | 26.9 | 2.10E+08 |
| 12.91193 | 15.71105 | 10.73 | 8 | 8 | 8 | 51.5 | 2.09E+08 |
| 13.08943 | 15.47564 | 12.72875 | 7 | 7 | 7 | 25.9 | 2.09E+08 |
| 12.6567 | 13.71994 | 12.34819 | 14 | 14 | 14 | 17.5 | 2.08E+08 |
| 19.48581 | 19.34241 | 19.15242 | 8 | 8 | 8 | 31.7 | 2.08E+08 |
| 10.51838 | 16.53397 | 17.33106 | 15 | 15 | 15 | 20.9 | 2.07E+08 |
| 14.7703 | 15.37429 | 11.87864 | 12 | 12 | 12 | 17.6 | 2.07E+08 |
| 11.89342 | 13.78167 | 18.11345 | 8 | 8 | 8 | 17.9 | 2.07E+08 |
| 11.14588 | 11.91224 | 12.65913 | 5 | 5 | 5 | 34.1 | 2.07E+08 |
| 16.62599 | 13.24136 | 17.09177 | 13 | 13 | 13 | 37.8 | 2.07E+08 |
| 16.93997 | 15.86227 | 16.61641 | 20 | 20 | 20 | 20.8 | 2.02E+08 |
| 13.87887 | 18.9757 | 15.53199 | 9 | 9 | 9 | 26 | 2.02E+08 |
| 18.67421 | 14.05427 | 11.47962 | 21 | 21 | 21 | 17.7 | 2.02E+08 |
| 14.38799 | 20.89519 | 19.5656 | 7 | 7 | 7 | 39.9 | 2.01E+08 |
| 13.4222 | 17.61784 | 17.26682 | 10 | 10 | 10 | 24.3 | 2.01E+08 |
| 11.84827 | 17.90566 | 12.69471 | 12 | 12 | 12 | 26.4 | 2.01E+08 |
| 12.5133 | 14.7139 | 12.47872 | 11 | 9 | 9 | 51.4 | 2.00E+08 |
| 17.9259 | 20.69668 | 19.34601 | 7 | 7 | 7 | 18.5 | 1.99E+08 |
| 11.32156 | 12.65099 | 13.25819 | 10 | 10 | 10 | 26.5 | 1.99E+08 |
| 13.21849 | 21.58412 | 11.43262 | 3 | 3 | 3 | 22.3 | 1.98E+08 |
| 14.20697 | 11.63918 | 15.95087 | 8 | 8 | 8 | 31.7 | 1.98E+08 |
| 12.65 | 13.75217 | 14.73983 | 21 | 15 | 15 | 10.1 | 1.97E+08 |
| 16.76927 | 12.16881 | 12.377 | 17 | 17 | 17 | 17.7 | 1.97E+08 |
| 11.19486 | 12.67224 | 11.95845 | 8 | 8 | 8 | 20.1 | 1.97E+08 |
| 11.31606 | 13.40992 | 12.01378 | 11 | 11 | 11 | 19.1 | 1.97E+08 |
| 16.49149 | 12.66832 | 12.25376 | 12 | 12 | 12 | 27.1 | 1.96E+08 |
| 11.43793 | 15.93556 | 12.66335 | 4 | 4 | 4 | 29.7 | 1.95E+08 |
| 11.37172 | 13.47056 | 14.00945 | 4 | 4 | 4 | 29.5 | 1.95E+08 |
| 17.63254 | 18.18694 | 11.18364 | 21 | 21 | 21 | 13.7 | 1.95E+08 |
| 11.31403 | 18.99515 | 12.68499 | 23 | 5 | 1 | 58.7 | 1.95E+08 |
| 12.26529 | 19.02467 | 11.83465 | 11 | 11 | 10 | 42.1 | 1.95E+08 |
| 10.98437 | 14.55399 | 11.27682 | 12 | 12 | 12 | 9 | 1.93E+08 |
| 21.70982 | 17.1135 | 12.3817 | 9 | 9 | 9 | 22.2 | 1.93E+08 |
| 14.64836 | 15.75045 | 16.03368 | 7 | 7 | 7 | 20.7 | 1.92E+08 |
| 10.84756 | 18.31265 | 14.89088 | 13 | 13 | 13 | 19.8 | 1.92E+08 |
| 13.21417 | 12.86039 | 12.33232 | 13 | 13 | 13 | 26.9 | 1.91E+08 |

| | | | | | | | |
|---|---|---|---|---|---|---|---|
| 12.79494 | 17.76584 | 17.6683 | 17 | 16 | 16 | 14.6 | 1.90E+08 |
| 13.21785 | 18.37254 | 12.64561 | 14 | 14 | 14 | 23 | 1.89E+08 |
| 11.51804 | 13.18051 | 13.24259 | 3 | 3 | 3 | 5.6 | 1.88E+08 |
| 16.62614 | 17.89841 | 16.49816 | 9 | 3 | 3 | 27.3 | 1.88E+08 |
| 19.12449 | 18.213 | 14.9443 | 18 | 18 | 18 | 26 | 1.87E+08 |
| 10.70193 | 16.67563 | 11.87743 | 10 | 10 | 10 | 25.5 | 1.87E+08 |
| 13.08448 | 17.07169 | 12.08117 | 9 | 9 | 9 | 14.7 | 1.87E+08 |
| 14.89964 | 16.95356 | 13.86308 | 9 | 9 | 9 | 28.4 | 1.86E+08 |
| 12.37231 | 15.22536 | 13.0336 | 15 | 13 | 13 | 27.3 | 1.86E+08 |
| 12.26446 | 18.91628 | 11.65654 | 8 | 8 | 8 | 26.6 | 1.86E+08 |
| 17.12727 | 18.28234 | 17.32166 | 10 | 10 | 10 | 26.4 | 1.85E+08 |
| 15.98081 | 20.15016 | 16.01435 | 15 | 15 | 15 | 37.9 | 1.85E+08 |
| 12.83364 | 16.74123 | 13.0499 | 9 | 9 | 6 | 58.8 | 1.85E+08 |
| 13.18919 | 16.46639 | 12.75907 | 10 | 10 | 10 | 28.7 | 1.85E+08 |
| 12.74582 | 15.08032 | 12.56162 | 7 | 7 | 7 | 23.4 | 1.84E+08 |
| 11.43112 | 19.78322 | 10.82524 | 6 | 6 | 6 | 20.9 | 1.84E+08 |
| 13.2223 | 13.82693 | 14.21167 | 4 | 4 | 4 | 43.1 | 1.84E+08 |
| 10.97785 | 13.25224 | 12.33975 | 12 | 12 | 12 | 27.4 | 1.84E+08 |
| 16.27879 | 19.39333 | 19.51019 | 10 | 3 | 3 | 50.6 | 1.83E+08 |
| 14.10058 | 11.88414 | 11.53019 | 15 | 15 | 15 | 18.6 | 1.81E+08 |
| 11.49594 | 16.32829 | 12.77113 | 17 | 17 | 17 | 15.6 | 1.81E+08 |
| 13.82926 | 18.28965 | 12.47322 | 10 | 10 | 10 | 28.6 | 1.81E+08 |
| 12.32939 | 19.6294 | 11.95619 | 8 | 8 | 8 | 35.7 | 1.80E+08 |
| 17.81215 | 21.17475 | 18.17933 | 8 | 5 | 5 | 4.9 | 1.80E+08 |
| 13.70919 | 15.2109 | 11.72667 | 9 | 9 | 9 | 22 | 1.80E+08 |
| 12.67666 | 24.19776 | 17.82146 | 9 | 6 | 6 | 29.4 | 1.80E+08 |
| 11.74212 | 12.78187 | 12.45319 | 4 | 4 | 4 | 45.3 | 1.79E+08 |
| 11.62156 | 14.05394 | 12.64395 | 12 | 12 | 12 | 25.2 | 1.78E+08 |
| 14.75045 | 14.21977 | 14.97361 | 10 | 10 | 10 | 19.3 | 1.77E+08 |
| 13.26061 | 20.19448 | 12.50887 | 8 | 8 | 8 | 33.1 | 1.76E+08 |
| 20.09468 | 12.96321 | 15.31494 | 12 | 12 | 12 | 17.5 | 1.76E+08 |
| 13.14227 | 13.9074 | 11.40446 | 14 | 14 | 14 | 6.4 | 1.75E+08 |
| 11.91983 | 14.35291 | 11.59987 | 10 | 10 | 10 | 50.8 | 1.75E+08 |
| 11.30988 | 14.95801 | 11.80076 | 12 | 12 | 12 | 23.2 | 1.75E+08 |
| 18.3276 | 20.59268 | 20.65176 | 12 | 12 | 12 | 21.3 | 1.75E+08 |
| 19.72447 | 17.6082 | 12.09204 | 7 | 7 | 7 | 23 | 1.74E+08 |
| 11.84615 | 11.76474 | 11.53009 | 14 | 14 | 14 | 18.7 | 1.74E+08 |
| 10.73024 | 13.637 | 11.43728 | 10 | 10 | 10 | 32.5 | 1.74E+08 |
| 20.83961 | 12.38618 | 9.631221 | 8 | 8 | 8 | 25.3 | 1.73E+08 |
| 13.2832 | 15.23934 | 13.79675 | 7 | 7 | 7 | 30.5 | 1.73E+08 |
| 17.82693 | 15.66783 | 10.56813 | 9 | 9 | 9 | 25.1 | 1.73E+08 |
| 18.79902 | 25.79454 | 18.67538 | 6 | 6 | 6 | 30.7 | 1.72E+08 |
| 9.904526 | 24.08253 | 22.56407 | 2 | 2 | 2 | 18.9 | 1.72E+08 |
| 10.72176 | 13.52997 | 13.64491 | 9 | 9 | 9 | 18.2 | 1.72E+08 |
| 12.83647 | 19.91718 | 10.24959 | 5 | 5 | 5 | 45.3 | 1.72E+08 |
| 12.66933 | 12.8779 | 11.76671 | 14 | 14 | 14 | 20 | 1.70E+08 |
| 12.74578 | 12.96812 | 13.02606 | 9 | 9 | 9 | 16.9 | 1.70E+08 |
| 11.29859 | 18.15512 | 12.88502 | 9 | 9 | 9 | 28 | 1.70E+08 |

| | | | | | | | |
|---|---|---|---|---|---|---|---|
| 17.45884 | 17.7127 | 13.48356 | 8 | 8 | 8 | 23.5 | 1.70E+08 |
| 14.55213 | 15.28211 | 12.28295 | 13 | 13 | 13 | 18 | 1.70E+08 |
| 11.67425 | 17.70334 | 11.74682 | 6 | 6 | 6 | 40 | 1.70E+08 |
| 14.26959 | 11.80492 | 11.61662 | 12 | 12 | 12 | 35.2 | 1.69E+08 |
| 19.96832 | 20.63316 | 12.63188 | 10 | 10 | 10 | 20.4 | 1.69E+08 |
| 11.86758 | 17.76137 | 17.22543 | 9 | 9 | 9 | 26.7 | 1.69E+08 |
| 11.86832 | 12.88366 | 11.89544 | 10 | 10 | 10 | 26.7 | 1.68E+08 |
| 10.83813 | 13.71263 | 12.13963 | 17 | 16 | 16 | 20.4 | 1.68E+08 |
| 13.34425 | 17.40449 | 12.55321 | 12 | 12 | 12 | 20.4 | 1.68E+08 |
| 20.84353 | 23.39506 | 21.5442 | 7 | 7 | 7 | 45.7 | 1.67E+08 |
| 11.6205 | 12.94795 | 13.62378 | 3 | 3 | 3 | 5 | 1.67E+08 |
| 19.69744 | 14.01602 | 11.87956 | 4 | 4 | 4 | 14.6 | 1.66E+08 |
| 13.50701 | 13.55937 | 12.00339 | 11 | 11 | 11 | 10.1 | 1.66E+08 |
| 11.46775 | 12.92912 | 12.77363 | 11 | 11 | 11 | 20.8 | 1.65E+08 |
| 13.33493 | 18.9946 | 11.74106 | 3 | 3 | 3 | 37.5 | 1.65E+08 |
| 12.41519 | 16.96172 | 11.87436 | 9 | 9 | 9 | 23.9 | 1.65E+08 |
| 12.41786 | 24.94552 | 16.15658 | 10 | 10 | 10 | 74.7 | 1.65E+08 |
| 14.61506 | 16.77867 | 15.72486 | 17 | 17 | 17 | 30.4 | 1.65E+08 |
| 12.16758 | 16.96274 | 16.68072 | 6 | 6 | 6 | 26.1 | 1.64E+08 |
| 11.8751 | 13.81626 | 12.02673 | 6 | 6 | 6 | 52.6 | 1.64E+08 |
| 13.98886 | 14.58895 | 12.65504 | 13 | 13 | 13 | 15.9 | 1.64E+08 |
| 17.4949 | 16.92347 | 14.68174 | 12 | 12 | 12 | 15.5 | 1.64E+08 |
| 11.96138 | 17.24042 | 15.2977 | 8 | 8 | 8 | 7.3 | 1.63E+08 |
| 11.79205 | 12.26855 | 11.77766 | 4 | 4 | 4 | 21 | 1.63E+08 |
| 12.80366 | 17.08597 | 16.918 | 10 | 10 | 10 | 48.3 | 1.63E+08 |
| 10.67764 | 13.1008 | 12.72118 | 10 | 10 | 10 | 8.8 | 1.62E+08 |
| 12.25615 | 16.36338 | 12.24415 | 16 | 16 | 16 | 21.9 | 1.62E+08 |
| 11.56101 | 18.79589 | 16.29915 | 9 | 9 | 9 | 56.7 | 1.62E+08 |
| 13.60955 | 14.56416 | 12.42261 | 17 | 16 | 16 | 14.2 | 1.62E+08 |
| 16.81303 | 14.20831 | 14.18898 | 8 | 8 | 8 | 36.7 | 1.61E+08 |
| 12.26048 | 13.80319 | 12.17653 | 14 | 14 | 14 | 16.7 | 1.61E+08 |
| 12.93892 | 13.69225 | 12.7046 | 7 | 7 | 7 | 28.7 | 1.60E+08 |
| 14.89614 | 13.83121 | 12.70594 | 7 | 7 | 7 | 56 | 1.60E+08 |
| 12.2361 | 12.13948 | 12.97983 | 5 | 5 | 5 | 17.7 | 1.60E+08 |
| 10.78934 | 13.14046 | 10.21602 | 13 | 13 | 13 | 15.5 | 1.59E+08 |
| 10.41067 | 20.04327 | 11.10958 | 22 | 12 | 12 | 15.5 | 1.57E+08 |
| 13.5541 | 16.38854 | 16.60661 | 8 | 8 | 8 | 30.7 | 1.56E+08 |
| 13.13576 | 15.38114 | 11.43157 | 23 | 15 | 12 | 14.2 | 1.55E+08 |
| 10.34906 | 19.6857 | 19.34317 | 15 | 15 | 15 | 12.4 | 1.55E+08 |
| 11.70506 | 12.42369 | 13.0671 | 6 | 6 | 6 | 28.8 | 1.55E+08 |
| 14.02349 | 13.45748 | 12.60317 | 8 | 8 | 8 | 17.1 | 1.55E+08 |
| 12.11965 | 19.58771 | 10.6608 | 6 | 5 | 5 | 20.6 | 1.55E+08 |
| 15.09433 | 17.43583 | 14.38526 | 20 | 20 | 20 | 6.2 | 1.55E+08 |
| 13.15358 | 19.30198 | 18.15605 | 15 | 15 | 15 | 21.4 | 1.54E+08 |
| 11.55592 | 20.3186 | 18.10682 | 7 | 7 | 7 | 40.3 | 1.54E+08 |
| 12.11938 | 20.49189 | 12.44306 | 15 | 15 | 15 | 9.4 | 1.54E+08 |
| 12.07735 | 17.93128 | 12.90713 | 11 | 11 | 11 | 20.6 | 1.54E+08 |
| 17.91099 | 20.92868 | 18.33176 | 9 | 9 | 9 | 32.1 | 1.54E+08 |

| | | | | | | | |
|---|---|---|---|---|---|---|---|
| 18.89897 | 17.6875 | 15.84992 | 16 | 16 | 16 | 14.8 | 1.53E+08 |
| 11.53551 | 19.7286 | 18.73932 | 5 | 5 | 5 | 25 | 1.53E+08 |
| 15.47659 | 16.68127 | 12.53976 | 4 | 4 | 4 | 25.4 | 1.53E+08 |
| 10.61201 | 11.78388 | 12.00618 | 7 | 7 | 7 | 17.3 | 1.52E+08 |
| 18.16575 | 23.55262 | 18.22095 | 9 | 9 | 9 | 26.2 | 1.52E+08 |
| 12.64903 | 13.92274 | 12.07805 | 7 | 7 | 7 | 23.3 | 1.52E+08 |
| 14.08978 | 15.08154 | 14.25068 | 14 | 14 | 14 | 37.8 | 1.50E+08 |
| 10.32782 | 12.24281 | 11.47267 | 7 | 7 | 7 | 42.7 | 1.50E+08 |
| 13.04737 | 18.15882 | 15.89851 | 9 | 9 | 9 | 20.6 | 1.50E+08 |
| 15.10259 | 15.88772 | 13.04447 | 11 | 11 | 11 | 16.8 | 1.49E+08 |
| 11.60811 | 13.63618 | 12.84502 | 6 | 6 | 6 | 39.2 | 1.48E+08 |
| 11.61794 | 17.09949 | 15.2911 | 6 | 6 | 6 | 35.3 | 1.48E+08 |
| 11.80513 | 12.4211 | 12.43794 | 15 | 15 | 15 | 11.3 | 1.48E+08 |
| 11.29751 | 15.40228 | 12.60782 | 12 | 12 | 12 | 14.9 | 1.45E+08 |
| 12.53853 | 16.39724 | 12.7756 | 9 | 9 | 9 | 17.2 | 1.45E+08 |
| 10.84536 | 17.17981 | 12.43405 | 13 | 13 | 13 | 29.3 | 1.44E+08 |
| 12.35616 | 17.56138 | 15.48972 | 8 | 8 | 8 | 16.1 | 1.44E+08 |
| 13.60207 | 18.21357 | 13.2961 | 12 | 12 | 12 | 27.4 | 1.44E+08 |
| 12.037 | 13.80657 | 16.71679 | 5 | 5 | 5 | 17.9 | 1.42E+08 |
| 11.76685 | 20.96529 | 12.1701 | 10 | 10 | 10 | 22.4 | 1.42E+08 |
| 12.57752 | 14.72132 | 10.82144 | 5 | 5 | 5 | 16.9 | 1.42E+08 |
| 10.86815 | 14.40178 | 14.16137 | 4 | 4 | 4 | 22.2 | 1.42E+08 |
| 18.35716 | 11.83804 | 18.51788 | 5 | 5 | 5 | 30.1 | 1.41E+08 |
| 11.29772 | 14.93509 | 13.88045 | 15 | 15 | 15 | 15.2 | 1.41E+08 |
| 19.16271 | 19.08618 | 19.32635 | 5 | 4 | 4 | 14.5 | 1.41E+08 |
| 17.83356 | 14.18769 | 13.0011 | 8 | 8 | 8 | 26.3 | 1.40E+08 |
| 13.57542 | 18.81309 | 11.6345 | 9 | 9 | 9 | 12.3 | 1.40E+08 |
| 12.21442 | 18.45647 | 12.50621 | 10 | 9 | 9 | 18.5 | 1.40E+08 |
| 12.10583 | 17.72986 | 12.84368 | 7 | 7 | 7 | 24.2 | 1.40E+08 |
| 13.48671 | 12.96701 | 16.58595 | 10 | 6 | 6 | 11.1 | 1.38E+08 |
| 11.7822 | 21.78831 | 12.94798 | 4 | 4 | 4 | 35 | 1.38E+08 |
| 10.95191 | 17.66574 | 12.40611 | 12 | 12 | 9 | 20.6 | 1.38E+08 |
| 15.95233 | 18.52936 | 16.79829 | 8 | 8 | 8 | 37.7 | 1.37E+08 |
| 13.12064 | 13.63159 | 11.65654 | 2 | 2 | 2 | 3.6 | 1.37E+08 |
| 12.65117 | 11.13704 | 13.17613 | 3 | 3 | 3 | 32.2 | 1.37E+08 |
| 11.46088 | 11.91728 | 13.38899 | 9 | 9 | 9 | 29.5 | 1.37E+08 |
| 13.1121 | 15.86351 | 11.83477 | 8 | 8 | 8 | 35.8 | 1.37E+08 |
| 13.97879 | 17.41757 | 12.08874 | 10 | 10 | 10 | 18.6 | 1.37E+08 |
| 11.15844 | 13.19654 | 12.09017 | 8 | 8 | 8 | 14.5 | 1.36E+08 |
| 15.74502 | 14.54007 | 17.1436 | 6 | 6 | 6 | 33.2 | 1.36E+08 |
| 10.1729 | 15.60918 | 12.6409 | 12 | 10 | 10 | 21.1 | 1.35E+08 |
| 11.96783 | 15.07445 | 11.8928 | 10 | 10 | 10 | 22 | 1.34E+08 |
| 11.65691 | 18.77941 | 11.41303 | 4 | 4 | 4 | 22.8 | 1.34E+08 |
| 17.55112 | 18.69598 | 13.59071 | 4 | 4 | 4 | 39.5 | 1.34E+08 |
| 11.58679 | 13.57498 | 11.9481 | 13 | 13 | 13 | 15.7 | 1.33E+08 |
| 14.77309 | 14.52717 | 13.64667 | 8 | 8 | 8 | 16 | 1.33E+08 |
| 11.02178 | 12.67684 | 14.56979 | 4 | 4 | 4 | 22.4 | 1.33E+08 |
| 12.55743 | 14.96628 | 13.17892 | 12 | 12 | 12 | 32.6 | 1.33E+08 |

| | | | | | | | |
|---|---|---|---|---|---|---|---|
| 11.23366 | 12.99203 | 13.99353 | 9 | 9 | 9 | 23.5 | 1.33E+08 |
| 12.39859 | 11.92604 | 12.99997 | 10 | 10 | 8 | 6.7 | 1.33E+08 |
| 11.72298 | 20.47279 | 11.91354 | 7 | 3 | 3 | 26.1 | 1.33E+08 |
| 13.70706 | 20.6289 | 13.15782 | 5 | 5 | 2 | 36.3 | 1.32E+08 |
| 21.1161 | 21.4889 | 21.06039 | 11 | 11 | 11 | 21.5 | 1.32E+08 |
| 11.40549 | 18.82258 | 11.48066 | 10 | 10 | 10 | 24.6 | 1.31E+08 |
| 11.92614 | 12.59685 | 11.95677 | 6 | 6 | 6 | 28.2 | 1.31E+08 |
| 12.42195 | 12.50645 | 12.48369 | 7 | 7 | 7 | 14.4 | 1.31E+08 |
| 13.81384 | 17.88071 | 10.2699 | 5 | 5 | 5 | 31.2 | 1.30E+08 |
| 17.65438 | 25.84628 | 19.63151 | 4 | 4 | 4 | 20.7 | 1.30E+08 |
| 12.77372 | 14.92629 | 11.97213 | 9 | 9 | 9 | 19.4 | 1.29E+08 |
| 12.95934 | 11.86102 | 12.98996 | 5 | 5 | 5 | 50.6 | 1.29E+08 |
| 12.33239 | 19.79193 | 19.94135 | 9 | 9 | 9 | 16 | 1.29E+08 |
| 14.30948 | 16.52795 | 15.10292 | 12 | 12 | 9 | 13.1 | 1.29E+08 |
| 13.58402 | 14.96034 | 12.79516 | 10 | 6 | 6 | 27.6 | 1.29E+08 |
| 12.61382 | 21.80207 | 17.92138 | 7 | 7 | 7 | 17.6 | 1.27E+08 |
| 11.95784 | 12.5421 | 12.37342 | 5 | 5 | 5 | 23.5 | 1.27E+08 |
| 15.80811 | 14.52913 | 11.98103 | 11 | 11 | 11 | 11.7 | 1.27E+08 |
| 19.51609 | 14.51378 | 12.01343 | 7 | 7 | 7 | 16.2 | 1.27E+08 |
| 16.4699 | 17.91893 | 16.46931 | 6 | 6 | 6 | 12.6 | 1.26E+08 |
| 12.56234 | 16.39052 | 15.09996 | 7 | 7 | 7 | 25.7 | 1.25E+08 |
| 13.63182 | 15.26557 | 13.22778 | 6 | 6 | 6 | 37.5 | 1.25E+08 |
| 12.19325 | 16.20214 | 15.52194 | 11 | 11 | 11 | 17 | 1.25E+08 |
| 12.55771 | 17.13531 | 14.0962 | 7 | 7 | 7 | 38.3 | 1.25E+08 |
| 11.61273 | 16.06672 | 15.16965 | 12 | 12 | 12 | 19.6 | 1.24E+08 |
| 11.76547 | 17.86942 | 12.88706 | 9 | 9 | 9 | 26.2 | 1.23E+08 |
| 11.63306 | 11.18139 | 11.24101 | 6 | 6 | 6 | 15.5 | 1.23E+08 |
| 15.86974 | 12.05649 | 16.82588 | 10 | 10 | 10 | 28.8 | 1.22E+08 |
| 13.07745 | 17.15076 | 13.584 | 12 | 12 | 12 | 12.4 | 1.22E+08 |
| 15.82075 | 15.89212 | 15.97806 | 8 | 8 | 8 | 31.4 | 1.22E+08 |
| 15.75549 | 16.55808 | 12.56951 | 4 | 4 | 4 | 32.3 | 1.22E+08 |
| 10.86831 | 20.34495 | 12.94738 | 3 | 3 | 3 | 29.6 | 1.21E+08 |
| 10.98183 | 19.76437 | 13.0808 | 7 | 7 | 7 | 32 | 1.21E+08 |
| 15.67923 | 18.82283 | 16.3809 | 9 | 9 | 9 | 17.7 | 1.20E+08 |
| 11.36205 | 18.79848 | 12.98817 | 9 | 9 | 9 | 20 | 1.19E+08 |
| 12.20905 | 18.34888 | 12.59069 | 9 | 9 | 9 | 34.2 | 1.19E+08 |
| 13.51835 | 20.82065 | 20.55992 | 3 | 3 | 3 | 42.7 | 1.18E+08 |
| 14.0022 | 16.15393 | 12.85576 | 14 | 14 | 14 | 26.4 | 1.18E+08 |
| 14.08843 | 13.86504 | 11.6423 | 12 | 12 | 12 | 14.8 | 1.18E+08 |
| 13.74598 | 16.32516 | 11.69271 | 7 | 7 | 7 | 14.6 | 1.18E+08 |
| 12.02315 | 16.90677 | 12.04568 | 7 | 7 | 7 | 28.3 | 1.17E+08 |
| 16.06827 | 21.31413 | 18.689 | 23 | 19 | 17 | 11.8 | 1.17E+08 |
| 17.37823 | 13.99766 | 12.34471 | 9 | 9 | 9 | 24.8 | 1.17E+08 |
| 12.34086 | 13.87026 | 13.4689 | 6 | 6 | 6 | 29.4 | 1.17E+08 |
| 12.47443 | 18.69995 | 12.72144 | 8 | 8 | 7 | 25.3 | 1.17E+08 |
| 10.88471 | 12.97959 | 12.45977 | 11 | 11 | 11 | 18 | 1.17E+08 |
| 12.84226 | 19.29757 | 13.36447 | 10 | 10 | 10 | 30.3 | 1.17E+08 |
| 13.35825 | 13.27561 | 12.90327 | 7 | 7 | 7 | 27.7 | 1.15E+08 |

| | | | | | | | |
|---|---|---|---|---|---|---|---|
| 17.96595 | 16.50109 | 11.05871 | 3 | 3 | 3 | 14.5 | 1.15E+08 |
| 18.16182 | 18.71683 | 18.26513 | 13 | 13 | 13 | 13.9 | 1.15E+08 |
| 11.3578 | 14.08005 | 11.99594 | 6 | 6 | 6 | 17.6 | 1.14E+08 |
| 11.04329 | 18.4771 | 12.03851 | 8 | 7 | 6 | 36.4 | 1.14E+08 |
| 13.54538 | 12.74547 | 11.12644 | 9 | 9 | 9 | 23.7 | 1.13E+08 |
| 16.77841 | 17.25857 | 16.9642 | 9 | 9 | 9 | 34.7 | 1.13E+08 |
| 11.25972 | 17.16103 | 15.86021 | 5 | 4 | 4 | 26.3 | 1.12E+08 |
| 20.26808 | 19.42673 | 18.37806 | 6 | 6 | 6 | 12.3 | 1.12E+08 |
| 12.27685 | 13.16855 | 10.54792 | 6 | 6 | 6 | 53.4 | 1.12E+08 |
| 11.2188 | 13.80272 | 12.74738 | 2 | 2 | 2 | 25.5 | 1.12E+08 |
| 12.84844 | 17.55052 | 12.17845 | 10 | 10 | 9 | 5.2 | 1.12E+08 |
| 13.9305 | 12.97426 | 12.6242 | 8 | 7 | 7 | 14.5 | 1.11E+08 |
| 10.47549 | 20.07352 | 11.90855 | 2 | 2 | 2 | 10.4 | 1.11E+08 |
| 12.73962 | 13.8434 | 10.47302 | 6 | 6 | 3 | 9.3 | 1.11E+08 |
| 11.95041 | 12.33084 | 18.91675 | 11 | 11 | 11 | 23.3 | 1.11E+08 |
| 11.19962 | 12.89011 | 11.82172 | 10 | 10 | 10 | 17.9 | 1.11E+08 |
| 20.51451 | 20.7917 | 20.31805 | 6 | 6 | 6 | 17 | 1.11E+08 |
| 11.45294 | 12.98143 | 11.27253 | 7 | 7 | 7 | 18.7 | 1.10E+08 |
| 10.80875 | 13.32809 | 12.86482 | 4 | 4 | 4 | 21.8 | 1.10E+08 |
| 12.98457 | 17.60719 | 10.72018 | 12 | 12 | 12 | 6.6 | 1.09E+08 |
| 20.15227 | 23.43949 | 23.44973 | 5 | 5 | 2 | 11.2 | 1.09E+08 |
| 16.39594 | 19.20146 | 13.8487 | 6 | 6 | 6 | 14.3 | 1.09E+08 |
| 11.84461 | 13.03479 | 16.81678 | 12 | 12 | 12 | 21.6 | 1.09E+08 |
| 13.05035 | 14.58678 | 14.00536 | 11 | 11 | 11 | 14.9 | 1.08E+08 |
| 22.22311 | 23.08926 | 22.76286 | 4 | 4 | 4 | 33.5 | 1.08E+08 |
| 14.23974 | 22.11518 | 17.33752 | 7 | 4 | 4 | 27.5 | 1.08E+08 |
| 12.43309 | 17.27812 | 12.71312 | 7 | 7 | 7 | 15.7 | 1.08E+08 |
| 12.91216 | 17.10677 | 10.51004 | 7 | 7 | 4 | 18.2 | 1.08E+08 |
| 18.55142 | 12.24347 | 11.25319 | 5 | 5 | 5 | 12.5 | 1.07E+08 |
| 13.36356 | 19.34549 | 12.14507 | 8 | 8 | 8 | 13.6 | 1.07E+08 |
| 9.773619 | 13.17562 | 13.59938 | 6 | 6 | 6 | 14.6 | 1.07E+08 |
| 17.20304 | 21.74275 | 16.98359 | 14 | 14 | 9 | 18.8 | 1.07E+08 |
| 12.25628 | 13.22359 | 11.17443 | 6 | 6 | 6 | 14 | 1.07E+08 |
| 10.89115 | 15.02488 | 12.54164 | 6 | 6 | 6 | 17.6 | 1.06E+08 |
| 10.85883 | 13.48546 | 14.40931 | 3 | 3 | 3 | 7 | 1.06E+08 |
| 12.92042 | 13.37499 | 13.77582 | 8 | 8 | 8 | 11.1 | 1.05E+08 |
| 11.16514 | 14.9756 | 13.82597 | 9 | 9 | 9 | 19.8 | 1.05E+08 |
| 19.04097 | 20.04073 | 19.61954 | 3 | 3 | 3 | 10.8 | 1.05E+08 |
| 11.24539 | 14.37599 | 17.20122 | 6 | 6 | 6 | 10.9 | 1.05E+08 |
| 11.92935 | 19.56387 | 10.62576 | 3 | 3 | 3 | 51.8 | 1.05E+08 |
| 13.64636 | 13.24114 | 12.92603 | 8 | 8 | 5 | 13.7 | 1.04E+08 |
| 12.28352 | 13.03473 | 13.07619 | 6 | 6 | 6 | 20.7 | 1.04E+08 |
| 11.52417 | 14.59528 | 12.81178 | 6 | 6 | 6 | 13 | 1.04E+08 |
| 12.37977 | 12.68236 | 21.49434 | 2 | 2 | 2 | 3.3 | 1.04E+08 |
| 13.81408 | 13.01393 | 13.1751 | 7 | 7 | 7 | 12.5 | 1.03E+08 |
| 13.43117 | 13.35805 | 12.19246 | 7 | 7 | 7 | 4.9 | 1.03E+08 |
| 18.15472 | 20.55478 | 12.86923 | 7 | 7 | 7 | 29.5 | 1.03E+08 |
| 13.38856 | 13.85359 | 12.01036 | 8 | 6 | 3 | 20.3 | 1.02E+08 |

| | | | | | | | |
|---|---|---|---|---|---|---|---|
| 19.89263 | 14.58231 | 20.0988 | 9 | 9 | 9 | 9.6 | 1.02E+08 |
| 11.50661 | 14.90427 | 13.65545 | 8 | 8 | 8 | 29.4 | 1.02E+08 |
| 12.16853 | 15.41521 | 12.12176 | 7 | 7 | 7 | 18.8 | 1.02E+08 |
| 11.83454 | 17.97466 | 12.04792 | 10 | 10 | 10 | 12.7 | 1.01E+08 |
| 15.77126 | 14.81628 | 13.81318 | 10 | 10 | 10 | 9.8 | 1.01E+08 |
| 17.4399 | 23.44708 | 16.90748 | 6 | 6 | 6 | 9.2 | 1.01E+08 |
| 12.88426 | 12.85048 | 10.69257 | 4 | 4 | 4 | 33.8 | 1.00E+08 |
| 15.13643 | 12.43532 | 13.51574 | 13 | 13 | 13 | 11.5 | 1.00E+08 |
| 13.72722 | 14.10915 | 12.39299 | 7 | 7 | 7 | 17 | 1.00E+08 |
| 17.86967 | 11.72377 | 13.23721 | 13 | 12 | 12 | 8.8 | 1.00E+08 |
| 18.00805 | 13.36974 | 11.54862 | 8 | 8 | 8 | 14.4 | 1.00E+08 |
| 14.29182 | 15.40045 | 11.77299 | 5 | 5 | 5 | 11.2 | 1.00E+08 |
| 11.97966 | 14.61419 | 13.56538 | 5 | 5 | 5 | 15 | 1.00E+08 |
| 12.39825 | 14.23182 | 10.73693 | 18 | 18 | 18 | 5.9 | 1.00E+08 |
| 11.58189 | 14.59341 | 12.39536 | 7 | 7 | 4 | 17.5 | 9.96E+07 |
| 13.47724 | 14.17779 | 13.12776 | 4 | 4 | 4 | 17.5 | 9.95E+07 |
| 16.25449 | 16.19637 | 16.5086 | 3 | 3 | 2 | 4.2 | 9.95E+07 |
| 13.41083 | 14.00281 | 13.58021 | 3 | 3 | 3 | 15.4 | 9.92E+07 |
| 12.7593 | 14.08899 | 16.3135 | 9 | 9 | 9 | 8.4 | 9.88E+07 |
| 12.61205 | 13.19026 | 13.2529 | 9 | 9 | 9 | 19.2 | 9.87E+07 |
| 11.28458 | 12.08444 | 17.5389 | 9 | 9 | 9 | 23.7 | 9.86E+07 |
| 12.45258 | 11.77385 | 13.7259 | 8 | 8 | 8 | 19.6 | 9.84E+07 |
| 19.69575 | 19.89636 | 20.14793 | 14 | 14 | 14 | 12.9 | 9.81E+07 |
| 12.74148 | 23.22672 | 12.0831 | 5 | 5 | 5 | 16.9 | 9.80E+07 |
| 13.69369 | 13.48705 | 11.58622 | 8 | 8 | 8 | 11.3 | 9.79E+07 |
| 19.67756 | 18.73975 | 19.77116 | 8 | 8 | 8 | 11.2 | 9.78E+07 |
| 18.27826 | 23.4353 | 18.43941 | 7 | 7 | 6 | 59.7 | 9.78E+07 |
| 11.94721 | 17.90906 | 12.7458 | 8 | 8 | 8 | 13.1 | 9.75E+07 |
| 12.69859 | 17.97298 | 12.76067 | 6 | 6 | 6 | 33.5 | 9.73E+07 |
| 12.09035 | 14.79893 | 13.17455 | 8 | 8 | 8 | 26 | 9.70E+07 |
| 12.46136 | 13.0903 | 12.48138 | 4 | 4 | 4 | 28.8 | 9.66E+07 |
| 10.94293 | 15.61252 | 16.64414 | 7 | 7 | 7 | 12.7 | 9.66E+07 |
| 12.54647 | 19.06647 | 12.67563 | 7 | 7 | 7 | 24.2 | 9.64E+07 |
| 17.95901 | 19.20278 | 18.54699 | 2 | 2 | 2 | 13.6 | 9.63E+07 |
| 12.3661 | 13.67027 | 16.12707 | 11 | 11 | 11 | 15.4 | 9.63E+07 |
| 14.05889 | 17.77623 | 10.42597 | 4 | 4 | 4 | 28.2 | 9.60E+07 |
| 13.1492 | 15.95707 | 13.34045 | 15 | 15 | 15 | 6.6 | 9.58E+07 |
| 11.54737 | 13.81148 | 9.862206 | 8 | 8 | 8 | 9.4 | 9.47E+07 |
| 13.34542 | 12.6089 | 12.54502 | 6 | 6 | 6 | 7.4 | 9.46E+07 |
| 10.34588 | 12.79001 | 12.35434 | 2 | 2 | 2 | 12 | 9.45E+07 |
| 12.10483 | 12.83272 | 12.02676 | 10 | 10 | 10 | 10.4 | 9.45E+07 |
| 16.91975 | 16.88774 | 16.72134 | 12 | 12 | 12 | 7.1 | 9.41E+07 |
| 19.08724 | 19.61081 | 19.37865 | 7 | 7 | 7 | 16.2 | 9.40E+07 |
| 12.6802 | 16.82252 | 16.15464 | 11 | 11 | 11 | 19.8 | 9.39E+07 |
| 12.88001 | 19.09762 | 12.63674 | 8 | 5 | 4 | 31.2 | 9.37E+07 |
| 10.85746 | 20.11769 | 11.66931 | 6 | 5 | 5 | 12.4 | 9.35E+07 |
| 13.09606 | 17.46228 | 13.52617 | 7 | 7 | 7 | 15.1 | 9.33E+07 |
| 15.80514 | 16.81666 | 13.22637 | 13 | 5 | 5 | 48.9 | 9.33E+07 |

| | | | | | | | |
|---|---|---|---|---|---|---|---|
| 12.21559 | 12.47448 | 11.37489 | 11 | 11 | 11 | 6 | 9.33E+07 |
| 13.75407 | 19.02524 | 12.89311 | 9 | 9 | 9 | 12.9 | 9.29E+07 |
| 11.01841 | 20.10072 | 9.662714 | 7 | 7 | 7 | 42.9 | 9.28E+07 |
| 10.0445 | 18.49911 | 12.11212 | 5 | 5 | 4 | 19.1 | 9.25E+07 |
| 11.58697 | 14.20337 | 13.9692 | 7 | 7 | 7 | 28.7 | 9.24E+07 |
| 10.75731 | 13.19408 | 12.49037 | 8 | 6 | 6 | 14.7 | 9.14E+07 |
| 15.25078 | 14.72462 | 16.03004 | 11 | 11 | 11 | 25 | 9.14E+07 |
| 11.94734 | 13.25548 | 10.91559 | 9 | 9 | 9 | 10.4 | 9.11E+07 |
| 17.17933 | 18.32307 | 17.01498 | 17 | 13 | 13 | 10.7 | 9.02E+07 |
| 12.72293 | 21.42044 | 22.86757 | 2 | 2 | 2 | 20.2 | 9.00E+07 |
| 19.87099 | 13.89991 | 17.5885 | 8 | 8 | 8 | 8.8 | 8.95E+07 |
| 11.93788 | 19.88083 | 11.78233 | 8 | 8 | 8 | 9.4 | 8.95E+07 |
| 19.53298 | 11.97375 | 22.55328 | 6 | 6 | 6 | 17.6 | 8.95E+07 |
| 11.47754 | 18.18413 | 12.91042 | 2 | 2 | 2 | 11.2 | 8.93E+07 |
| 11.80013 | 17.75532 | 13.11366 | 7 | 7 | 7 | 18.3 | 8.91E+07 |
| 18.92706 | 22.02871 | 18.96415 | 7 | 7 | 7 | 10.4 | 8.90E+07 |
| 10.87051 | 15.22298 | 12.85733 | 7 | 7 | 7 | 6.8 | 8.86E+07 |
| 18.05579 | 20.39645 | 11.99041 | 5 | 5 | 5 | 14.5 | 8.83E+07 |
| 15.54801 | 18.204 | 13.50233 | 9 | 9 | 8 | 12.5 | 8.81E+07 |
| 12.19079 | 13.3823 | 12.24851 | 4 | 4 | 4 | 17 | 8.79E+07 |
| 12.61065 | 11.59441 | 11.7923 | 2 | 2 | 2 | 4.8 | 8.73E+07 |
| 9.913073 | 13.45838 | 12.99518 | 5 | 5 | 5 | 50.6 | 8.70E+07 |
| 11.85417 | 19.61288 | 13.18727 | 8 | 8 | 8 | 15.2 | 8.69E+07 |
| 11.8313 | 17.83664 | 11.63254 | 3 | 3 | 3 | 13.4 | 8.66E+07 |
| 13.56009 | 20.73769 | 12.16938 | 8 | 8 | 8 | 7.6 | 8.63E+07 |
| 14.67165 | 12.92232 | 16.70063 | 10 | 10 | 10 | 9.3 | 8.62E+07 |
| 11.47171 | 14.63916 | 11.70641 | 8 | 8 | 8 | 7.9 | 8.61E+07 |
| 12.79444 | 12.68266 | 11.09598 | 11 | 11 | 11 | 19 | 8.59E+07 |
| 10.33013 | 19.75053 | 20.51865 | 3 | 3 | 2 | 23.5 | 8.58E+07 |
| 20.02378 | 14.20785 | 19.45724 | 6 | 6 | 6 | 16.4 | 8.57E+07 |
| 13.28639 | 14.2927 | 13.50473 | 3 | 3 | 3 | 13 | 8.56E+07 |
| 10.86229 | 13.48149 | 14.00571 | 4 | 4 | 4 | 34 | 8.55E+07 |
| 11.6091 | 12.16892 | 11.54568 | 5 | 5 | 5 | 14.3 | 8.51E+07 |
| 12.67789 | 23.05876 | 10.95984 | 10 | 10 | 10 | 31.8 | 8.51E+07 |
| 11.81972 | 14.59033 | 12.71052 | 4 | 4 | 4 | 21.6 | 8.50E+07 |
| 13.91496 | 14.90796 | 11.58423 | 7 | 7 | 7 | 10.7 | 8.44E+07 |
| 11.68907 | 15.26958 | 18.713 | 6 | 6 | 6 | 32.1 | 8.41E+07 |
| 12.44353 | 20.92904 | 17.84003 | 3 | 3 | 3 | 23.8 | 8.36E+07 |
| 13.75705 | 13.23907 | 13.62162 | 7 | 7 | 7 | 28.6 | 8.29E+07 |
| 18.07796 | 18.72657 | 11.33694 | 6 | 6 | 5 | 7.2 | 8.27E+07 |
| 13.25807 | 12.76463 | 12.62779 | 9 | 9 | 9 | 8.1 | 8.22E+07 |
| 18.78411 | 13.61388 | 19.05854 | 7 | 7 | 7 | 9.6 | 8.14E+07 |
| 13.8856 | 23.19408 | 15.13218 | 9 | 9 | 9 | 32.7 | 8.13E+07 |
| 11.71746 | 13.89262 | 12.4183 | 4 | 4 | 3 | 28.2 | 8.11E+07 |
| 16.49787 | 16.25061 | 12.76551 | 7 | 7 | 7 | 16.4 | 8.10E+07 |
| 13.22707 | 17.28996 | 13.30988 | 7 | 4 | 4 | 19.6 | 8.02E+07 |
| 19.22147 | 23.55555 | 19.37729 | 7 | 6 | 6 | 35.9 | 8.02E+07 |
| 11.77967 | 17.461 | 11.34628 | 5 | 5 | 5 | 10 | 8.01E+07 |

| | | | | | | | |
|---|---|---|---|---|---|---|---|
| 14.58061 | 14.45513 | 12.55616 | 7 | 7 | 7 | 17.5 | 7.99E+07 |
| 11.20171 | 10.92955 | 12.48464 | 5 | 5 | 5 | 23.6 | 7.95E+07 |
| 11.64289 | 13.78914 | 13.09373 | 6 | 6 | 6 | 12 | 7.89E+07 |
| 15.63931 | 14.15452 | 12.30657 | 12 | 12 | 12 | 5 | 7.89E+07 |
| 13.69351 | 17.25654 | 10.64361 | 7 | 7 | 7 | 9.4 | 7.89E+07 |
| 11.64519 | 20.14718 | 10.99429 | 3 | 3 | 3 | 12.9 | 7.85E+07 |
| 16.78507 | 17.28988 | 17.57555 | 5 | 5 | 5 | 15.3 | 7.85E+07 |
| 20.65631 | 13.31308 | 22.807 | 8 | 8 | 6 | 11.6 | 7.84E+07 |
| 12.17924 | 13.73093 | 12.29029 | 5 | 5 | 5 | 27.6 | 7.80E+07 |
| 13.83545 | 12.55045 | 10.13573 | 4 | 4 | 4 | 45.7 | 7.80E+07 |
| 12.92387 | 19.57804 | 12.47519 | 6 | 6 | 6 | 20.9 | 7.77E+07 |
| 19.92281 | 18.74557 | 12.49104 | 7 | 7 | 7 | 18.7 | 7.70E+07 |
| 12.98841 | 15.67934 | 10.0252 | 10 | 10 | 10 | 5.4 | 7.67E+07 |
| 12.4544 | 11.38939 | 13.47034 | 8 | 8 | 8 | 10.7 | 7.66E+07 |
| 17.76137 | 17.54125 | 13.48838 | 9 | 9 | 9 | 32.1 | 7.66E+07 |
| 13.1059 | 13.05524 | 12.5143 | 7 | 7 | 7 | 15.4 | 7.62E+07 |
| 12.95008 | 12.81615 | 11.86167 | 4 | 4 | 4 | 11.2 | 7.62E+07 |
| 13.04739 | 18.88956 | 13.26433 | 2 | 2 | 2 | 20.1 | 7.58E+07 |
| 21.90065 | 22.02405 | 21.92904 | 6 | 6 | 6 | 13.6 | 7.56E+07 |
| 11.1548 | 17.56347 | 12.61975 | 8 | 8 | 8 | 14.7 | 7.56E+07 |
| 13.85614 | 13.81435 | 11.73087 | 10 | 7 | 7 | 19.7 | 7.53E+07 |
| 11.71441 | 12.55874 | 13.16717 | 6 | 6 | 5 | 15.3 | 7.52E+07 |
| 12.3222 | 13.08452 | 11.66342 | 8 | 8 | 8 | 31.8 | 7.50E+07 |
| 12.95714 | 12.60864 | 11.52378 | 6 | 6 | 6 | 7.2 | 7.47E+07 |
| 12.94516 | 13.22918 | 13.77529 | 7 | 7 | 7 | 10.9 | 7.46E+07 |
| 11.53861 | 12.08718 | 12.91484 | 8 | 8 | 8 | 24.8 | 7.42E+07 |
| 12.82343 | 13.38738 | 11.96156 | 3 | 3 | 3 | 18.7 | 7.41E+07 |
| 10.23984 | 17.11777 | 12.53386 | 8 | 8 | 8 | 26.3 | 7.38E+07 |
| 12.9775 | 13.38086 | 12.16136 | 9 | 9 | 9 | 29.3 | 7.37E+07 |
| 19.08869 | 18.76237 | 18.81594 | 5 | 5 | 5 | 6.2 | 7.36E+07 |
| 15.51465 | 14.48596 | 13.09688 | 6 | 6 | 6 | 16.5 | 7.34E+07 |
| 11.3632 | 13.06982 | 13.69746 | 7 | 7 | 7 | 15.3 | 7.31E+07 |
| 12.11415 | 14.16188 | 11.57794 | 5 | 5 | 5 | 31.7 | 7.30E+07 |
| 10.99362 | 14.93099 | 13.1366 | 4 | 4 | 4 | 42.9 | 7.29E+07 |
| 11.24978 | 13.12528 | 12.23711 | 9 | 9 | 9 | 17 | 7.29E+07 |
| 12.07385 | 12.99019 | 13.34645 | 6 | 6 | 6 | 19.5 | 7.28E+07 |
| 10.80515 | 14.03502 | 13.15205 | 8 | 8 | 8 | 25.2 | 7.25E+07 |
| 16.8538 | 16.90619 | 11.79111 | 7 | 7 | 7 | 8.1 | 7.24E+07 |
| 17.80744 | 17.72294 | 17.52528 | 3 | 3 | 3 | 10.5 | 7.23E+07 |
| 12.42862 | 12.42454 | 11.78286 | 7 | 7 | 7 | 13.7 | 7.22E+07 |
| 12.26502 | 15.56546 | 12.30046 | 7 | 7 | 7 | 13 | 7.22E+07 |
| 12.79057 | 13.41883 | 12.90942 | 6 | 6 | 6 | 17 | 7.18E+07 |
| 17.33821 | 18.08794 | 17.53191 | 7 | 7 | 7 | 22.8 | 7.16E+07 |
| 12.88777 | 12.66409 | 13.01936 | 9 | 9 | 9 | 11.7 | 7.14E+07 |
| 13.08284 | 13.57085 | 11.69176 | 6 | 6 | 6 | 10.4 | 7.13E+07 |
| 13.35672 | 14.36796 | 12.95114 | 5 | 5 | 5 | 16.4 | 7.12E+07 |
| 12.69293 | 17.9718 | 12.15428 | 7 | 7 | 7 | 12.2 | 7.12E+07 |
| 17.11136 | 16.92474 | 16.71706 | 8 | 8 | 8 | 8.8 | 7.10E+07 |

| | | | | | | | |
|---|---|---|---|---|---|---|---|
| 12.85187 | 19.06497 | 13.3493 | 7 | 7 | 7 | 18 | 7.08E+07 |
| 17.64555 | 18.08171 | 17.66178 | 5 | 5 | 5 | 14.6 | 7.07E+07 |
| 14.24742 | 14.69717 | 13.16737 | 6 | 2 | 2 | 20.7 | 7.07E+07 |
| 16.39247 | 16.61999 | 12.90661 | 6 | 6 | 6 | 11.1 | 7.06E+07 |
| 12.93032 | 18.22976 | 14.03649 | 3 | 3 | 3 | 21.1 | 7.05E+07 |
| 12.90736 | 12.7446 | 12.20709 | 2 | 2 | 2 | 29.3 | 7.05E+07 |
| 12.5362 | 13.53649 | 10.51255 | 6 | 6 | 6 | 7.3 | 7.01E+07 |
| 11.5706 | 13.28481 | 11.60209 | 7 | 7 | 6 | 5 | 7.01E+07 |
| 14.45089 | 13.43515 | 13.71607 | 8 | 8 | 8 | 22.5 | 6.98E+07 |
| 12.7627 | 14.2087 | 12.69223 | 7 | 7 | 7 | 13.8 | 6.98E+07 |
| 13.74362 | 14.90176 | 13.44889 | 6 | 6 | 6 | 12.5 | 6.96E+07 |
| 11.7673 | 12.74435 | 18.88417 | 8 | 8 | 8 | 8.3 | 6.93E+07 |
| 12.70537 | 15.21054 | 12.62039 | 7 | 7 | 7 | 15.3 | 6.93E+07 |
| 12.08124 | 20.25165 | 12.40432 | 5 | 5 | 5 | 12.7 | 6.91E+07 |
| 10.89488 | 13.67795 | 11.80585 | 3 | 3 | 3 | 12.9 | 6.90E+07 |
| 10.01738 | 19.28084 | 11.38765 | 2 | 2 | 2 | 25 | 6.89E+07 |
| 11.68813 | 17.21048 | 18.0828 | 10 | 10 | 10 | 11.6 | 6.85E+07 |
| 13.37367 | 14.1554 | 13.00462 | 9 | 9 | 9 | 17 | 6.85E+07 |
| 11.73761 | 19.19005 | 12.09878 | 7 | 7 | 7 | 16.3 | 6.82E+07 |
| 18.50502 | 17.46698 | 10.45926 | 5 | 5 | 5 | 17.9 | 6.81E+07 |
| 11.80144 | 11.8436 | 10.69071 | 6 | 6 | 6 | 20 | 6.78E+07 |
| 12.4393 | 12.16936 | 12.36704 | 5 | 5 | 5 | 12.2 | 6.77E+07 |
| 11.7279 | 15.91551 | 11.33165 | 7 | 7 | 7 | 13.3 | 6.77E+07 |
| 11.20896 | 12.82756 | 21.91681 | 3 | 3 | 3 | 12.6 | 6.76E+07 |
| 12.94604 | 12.4931 | 12.06196 | 2 | 2 | 2 | 16.2 | 6.75E+07 |
| 16.42016 | 17.33044 | 15.45751 | 6 | 6 | 6 | 16 | 6.71E+07 |
| 19.59705 | 20.75898 | 20.33919 | 5 | 5 | 5 | 15.2 | 6.70E+07 |
| 12.08667 | 14.8451 | 12.22093 | 6 | 6 | 6 | 13.7 | 6.70E+07 |
| 11.84442 | 11.40992 | 11.40699 | 6 | 6 | 6 | 17.3 | 6.68E+07 |
| 12.45594 | 20.53146 | 11.75866 | 4 | 4 | 4 | 18.7 | 6.67E+07 |
| 11.25056 | 17.95543 | 14.22273 | 6 | 6 | 6 | 9.8 | 6.66E+07 |
| 12.50614 | 13.26863 | 12.82939 | 4 | 4 | 4 | 13.4 | 6.66E+07 |
| 11.96716 | 16.48784 | 16.37336 | 7 | 7 | 7 | 9.7 | 6.66E+07 |
| 11.721 | 12.33416 | 11.91555 | 8 | 8 | 8 | 8.3 | 6.63E+07 |
| 13.38049 | 13.19194 | 11.95532 | 2 | 2 | 2 | 9.2 | 6.63E+07 |
| 10.7716 | 12.03512 | 13.12095 | 8 | 8 | 8 | 30.8 | 6.62E+07 |
| 12.26276 | 13.5896 | 11.81905 | 5 | 5 | 5 | 10 | 6.58E+07 |
| 14.05613 | 19.16325 | 12.01441 | 8 | 8 | 8 | 9.2 | 6.57E+07 |
| 20.3043 | 14.4442 | 11.78839 | 5 | 5 | 5 | 26.2 | 6.55E+07 |
| 12.72227 | 13.65093 | 12.05257 | 6 | 6 | 6 | 23.5 | 6.51E+07 |
| 13.71082 | 14.82621 | 12.31278 | 6 | 6 | 6 | 12.1 | 6.49E+07 |
| 9.870546 | 19.20125 | 12.67136 | 5 | 5 | 5 | 7.8 | 6.49E+07 |
| 17.55142 | 13.34881 | 13.49918 | 7 | 7 | 7 | 7.2 | 6.45E+07 |
| 20.2018 | 20.46832 | 20.05603 | 6 | 6 | 6 | 5 | 6.39E+07 |
| 19.50918 | 19.02818 | 18.03209 | 6 | 6 | 6 | 22.2 | 6.39E+07 |
| 11.84066 | 17.82208 | 11.55441 | 6 | 6 | 6 | 28.4 | 6.38E+07 |
| 19.62087 | 12.73137 | 19.92369 | 5 | 5 | 5 | 11 | 6.37E+07 |
| 11.40845 | 13.76965 | 15.4783 | 5 | 5 | 5 | 25 | 6.35E+07 |

| | | | | | | | |
|---|---|---|---|---|---|---|---|
| 11.37659 | 12.95427 | 10.27758 | 6 | 6 | 6 | 15.5 | 6.34E+07 |
| 11.17788 | 13.21486 | 13.27382 | 3 | 3 | 3 | 10.6 | 6.33E+07 |
| 22.43194 | 22.6011 | 22.56725 | 5 | 5 | 5 | 13.3 | 6.29E+07 |
| 13.17841 | 12.81286 | 12.65639 | 2 | 2 | 2 | 3.5 | 6.25E+07 |
| 11.341 | 18.59729 | 10.16081 | 7 | 7 | 7 | 11 | 6.25E+07 |
| 11.97255 | 12.7508 | 11.74419 | 6 | 6 | 6 | 16.7 | 6.24E+07 |
| 11.81944 | 14.52874 | 20.65692 | 7 | 7 | 7 | 22.1 | 6.24E+07 |
| 18.21 | 17.40782 | 16.77004 | 7 | 6 | 5 | 16.5 | 6.24E+07 |
| 11.78231 | 12.41175 | 12.67307 | 18 | 5 | 5 | 15.3 | 6.23E+07 |
| 19.45511 | 18.05027 | 12.92354 | 2 | 2 | 2 | 26.1 | 6.22E+07 |
| 12.60817 | 14.04816 | 12.48046 | 4 | 4 | 4 | 30.1 | 6.22E+07 |
| 12.47579 | 19.02691 | 12.15037 | 4 | 4 | 4 | 21.7 | 6.20E+07 |
| 18.88429 | 24.30869 | 18.54367 | 4 | 4 | 4 | 27.6 | 6.20E+07 |
| 10.53649 | 17.64829 | 12.26546 | 6 | 6 | 5 | 16.5 | 6.16E+07 |
| 19.29744 | 19.26856 | 19.24402 | 5 | 5 | 5 | 7.6 | 6.16E+07 |
| 11.87438 | 14.7812 | 12.13416 | 4 | 4 | 4 | 14.8 | 6.16E+07 |
| 11.00236 | 15.12588 | 12.19468 | 4 | 4 | 4 | 27.3 | 6.13E+07 |
| 13.35553 | 12.68795 | 12.07118 | 2 | 2 | 2 | 12.9 | 6.12E+07 |
| 11.78675 | 13.62227 | 16.64216 | 3 | 3 | 3 | 48.1 | 6.12E+07 |
| 11.66956 | 18.58809 | 13.02932 | 4 | 4 | 4 | 13.5 | 6.10E+07 |
| 12.6052 | 14.67404 | 11.13868 | 8 | 5 | 5 | 64.1 | 6.10E+07 |
| 10.89603 | 22.59817 | 11.62444 | 6 | 6 | 6 | 18.7 | 6.08E+07 |
| 11.73663 | 13.09015 | 13.50889 | 6 | 6 | 6 | 15.3 | 6.07E+07 |
| 12.78279 | 18.65582 | 12.39828 | 4 | 4 | 4 | 21.2 | 6.06E+07 |
| 11.47503 | 13.48154 | 12.06058 | 7 | 7 | 7 | 5.7 | 6.01E+07 |
| 12.12578 | 13.23542 | 13.67625 | 8 | 8 | 8 | 10.2 | 6.01E+07 |
| 12.56006 | 13.85344 | 12.52583 | 9 | 9 | 9 | 9 | 5.98E+07 |
| 10.84213 | 18.84595 | 19.48782 | 27 | 4 | 4 | 53.9 | 5.98E+07 |
| 11.46358 | 20.02025 | 11.25453 | 10 | 10 | 10 | 4.5 | 5.97E+07 |
| 12.59798 | 18.3232 | 17.17231 | 6 | 6 | 6 | 8.3 | 5.97E+07 |
| 11.18228 | 13.73017 | 11.41005 | 5 | 5 | 5 | 34.8 | 5.96E+07 |
| 12.49334 | 18.45495 | 12.73051 | 4 | 4 | 4 | 7.9 | 5.95E+07 |
| 12.71584 | 12.72011 | 11.29447 | 3 | 3 | 3 | 6.5 | 5.95E+07 |
| 13.32917 | 13.32532 | 12.65784 | 4 | 4 | 4 | 20.4 | 5.95E+07 |
| 18.45768 | 18.68332 | 11.40801 | 9 | 4 | 4 | 24.5 | 5.94E+07 |
| 11.50922 | 12.09703 | 21.07549 | 6 | 6 | 6 | 15.1 | 5.93E+07 |
| 13.56641 | 11.52284 | 12.05954 | 2 | 2 | 2 | 14.8 | 5.89E+07 |
| 13.91537 | 12.22398 | 11.90271 | 5 | 5 | 5 | 9.9 | 5.86E+07 |
| 9.548836 | 12.64959 | 11.34865 | 2 | 2 | 2 | 44.8 | 5.86E+07 |
| 11.71746 | 13.30619 | 11.94522 | 3 | 3 | 3 | 22.8 | 5.85E+07 |
| 12.85935 | 16.56145 | 16.59282 | 6 | 6 | 6 | 12.4 | 5.83E+07 |
| 12.05547 | 14.62506 | 11.88021 | 4 | 4 | 4 | 11.4 | 5.81E+07 |
| 13.01906 | 17.74195 | 12.4339 | 4 | 4 | 4 | 16.8 | 5.79E+07 |
| 12.0677 | 13.97522 | 12.33702 | 4 | 4 | 4 | 62.5 | 5.78E+07 |
| 11.00192 | 14.47635 | 11.77162 | 5 | 5 | 5 | 10.4 | 5.75E+07 |
| 11.20996 | 20.61351 | 13.68329 | 6 | 6 | 6 | 10.1 | 5.74E+07 |
| 11.49056 | 12.34458 | 13.37321 | 6 | 2 | 2 | 14.7 | 5.74E+07 |
| 12.40221 | 16.24228 | 16.69493 | 7 | 5 | 5 | 41.3 | 5.74E+07 |

| | | | | | | | |
|---|---|---|---|---|---|---|---|
| 18.6392 | 19.57631 | 19.70597 | 3 | 3 | 3 | 14.5 | 5.74E+07 |
| 12.14089 | 12.36251 | 11.39919 | 3 | 3 | 3 | 8.1 | 5.74E+07 |
| 12.55392 | 18.41043 | 12.34457 | 4 | 4 | 4 | 22.2 | 5.70E+07 |
| 18.13195 | 18.70695 | 11.22613 | 4 | 4 | 4 | 13.9 | 5.68E+07 |
| 11.88133 | 19.17906 | 11.07909 | 4 | 4 | 4 | 36.9 | 5.65E+07 |
| 11.94992 | 12.78004 | 12.893 | 4 | 4 | 4 | 13.4 | 5.65E+07 |
| 12.7404 | 14.10356 | 10.44196 | 7 | 7 | 7 | 9.1 | 5.61E+07 |
| 12.42733 | 17.96764 | 17.24869 | 6 | 6 | 6 | 13.4 | 5.60E+07 |
| 12.82704 | 11.99516 | 11.8499 | 2 | 2 | 2 | 29.9 | 5.59E+07 |
| 11.94947 | 14.86833 | 11.20586 | 7 | 7 | 7 | 14.9 | 5.58E+07 |
| 12.18284 | 17.89587 | 12.81954 | 6 | 6 | 6 | 17.7 | 5.53E+07 |
| 11.9748 | 13.5217 | 11.82819 | 2 | 2 | 2 | 5.3 | 5.52E+07 |
| 12.5109 | 13.81926 | 10.87418 | 4 | 4 | 4 | 13.9 | 5.49E+07 |
| 12.94016 | 13.51722 | 13.40163 | 7 | 7 | 7 | 8.1 | 5.48E+07 |
| 11.04263 | 12.73854 | 12.85603 | 5 | 5 | 5 | 21.3 | 5.47E+07 |
| 12.19113 | 12.42385 | 11.45472 | 7 | 4 | 4 | 28 | 5.43E+07 |
| 12.30221 | 12.63411 | 12.33081 | 8 | 8 | 8 | 13.3 | 5.42E+07 |
| 12.77234 | 14.08073 | 11.58998 | 6 | 6 | 6 | 12.1 | 5.38E+07 |
| 11.89363 | 18.2091 | 11.95166 | 8 | 8 | 8 | 18.1 | 5.37E+07 |
| 12.10529 | 12.46848 | 10.99734 | 7 | 7 | 7 | 7.4 | 5.36E+07 |
| 19.02996 | 23.16869 | 12.7441 | 6 | 6 | 6 | 9.4 | 5.35E+07 |
| 12.03973 | 11.17363 | 12.5305 | 4 | 4 | 4 | 18 | 5.35E+07 |
| 13.2307 | 20.54915 | 12.78851 | 4 | 4 | 4 | 14.8 | 5.33E+07 |
| 11.93907 | 17.83701 | 13.07765 | 3 | 3 | 3 | 7.5 | 5.31E+07 |
| 12.00307 | 13.93112 | 14.50075 | 4 | 4 | 1 | 9.4 | 5.31E+07 |
| 22.27387 | 22.71387 | 22.36856 | 2 | 2 | 2 | 5.2 | 5.30E+07 |
| 11.07814 | 12.87139 | 10.53615 | 5 | 5 | 5 | 26.3 | 5.28E+07 |
| 16.24159 | 13.24506 | 12.36738 | 5 | 5 | 5 | 5.8 | 5.28E+07 |
| 11.56047 | 20.76142 | 13.77616 | 7 | 2 | 2 | 27.6 | 5.26E+07 |
| 12.69247 | 19.33767 | 13.04031 | 5 | 5 | 5 | 5.1 | 5.25E+07 |
| 10.88682 | 12.28346 | 10.91927 | 4 | 4 | 4 | 18.9 | 5.25E+07 |
| 12.73521 | 15.20178 | 12.48851 | 5 | 5 | 5 | 12.1 | 5.24E+07 |
| 21.06171 | 21.08291 | 21.13138 | 4 | 4 | 4 | 22.8 | 5.22E+07 |
| 12.9923 | 20.51586 | 13.25891 | 3 | 3 | 3 | 17.1 | 5.22E+07 |
| 12.57952 | 19.05756 | 11.36547 | 5 | 5 | 5 | 18.4 | 5.22E+07 |
| 18.71018 | 19.79178 | 11.18965 | 4 | 4 | 4 | 18.4 | 5.22E+07 |
| 19.12959 | 20.29749 | 19.80401 | 5 | 5 | 5 | 7.4 | 5.21E+07 |
| 11.85271 | 13.16244 | 12.58356 | 6 | 6 | 6 | 10.5 | 5.20E+07 |
| 21.9113 | 21.17005 | 21.52444 | 4 | 4 | 4 | 33.5 | 5.20E+07 |
| 12.67361 | 16.61569 | 12.17499 | 7 | 7 | 7 | 10.9 | 5.19E+07 |
| 10.87201 | 13.45946 | 19.23196 | 7 | 7 | 7 | 16.7 | 5.16E+07 |
| 18.20027 | 18.07833 | 18.20815 | 5 | 5 | 5 | 17.5 | 5.15E+07 |
| 20.10072 | 13.50914 | 20.3721 | 4 | 4 | 4 | 13.5 | 5.13E+07 |
| 13.33459 | 13.05705 | 12.07219 | 3 | 3 | 3 | 13.2 | 5.13E+07 |
| 16.5826 | 16.34774 | 15.30692 | 4 | 4 | 4 | 17.1 | 5.13E+07 |
| 11.31489 | 13.58458 | 14.08209 | 6 | 6 | 6 | 11.8 | 5.12E+07 |
| 12.17328 | 22.2623 | 10.65388 | 21 | 2 | 2 | 47.3 | 5.12E+07 |
| 12.75179 | 13.90342 | 11.50095 | 6 | 6 | 6 | 30.7 | 5.12E+07 |

| | | | | | | | |
|---|---|---|---|---|---|---|---|
| 12.65819 | 13.85426 | 11.185 | 8 | 8 | 8 | 5.9 | 5.11E+07 |
| 10.95731 | 17.18892 | 18.41741 | 5 | 5 | 5 | 19.4 | 5.09E+07 |
| 10.93972 | 13.36517 | 11.97129 | 2 | 2 | 2 | 5 | 5.06E+07 |
| 19.78344 | 20.12654 | 19.56283 | 5 | 5 | 5 | 10 | 5.06E+07 |
| 17.66719 | 12.86287 | 11.67672 | 5 | 5 | 5 | 22.4 | 5.04E+07 |
| 15.87462 | 13.20555 | 13.65529 | 6 | 6 | 6 | 9.8 | 5.00E+07 |
| 11.59009 | 13.91463 | 12.78494 | 3 | 3 | 3 | 7.3 | 4.99E+07 |
| 9.80567 | 11.86952 | 11.93053 | 2 | 2 | 2 | 1.1 | 4.99E+07 |
| 12.07806 | 13.01483 | 12.61857 | 4 | 4 | 4 | 7.8 | 4.97E+07 |
| 12.0378 | 12.35352 | 11.67067 | 5 | 5 | 5 | 17.6 | 4.96E+07 |
| 10.70305 | 19.5372 | 12.82244 | 5 | 5 | 5 | 17.1 | 4.94E+07 |
| 10.43523 | 12.21951 | 9.67052 | 5 | 5 | 5 | 12.4 | 4.93E+07 |
| 8.47246 | 15.26924 | 11.53926 | 6 | 6 | 5 | 14.6 | 4.92E+07 |
| 12.01641 | 13.3149 | 12.08577 | 6 | 6 | 5 | 13.5 | 4.92E+07 |
| 16.16726 | 18.24261 | 12.31836 | 4 | 4 | 4 | 23.5 | 4.91E+07 |
| 18.10313 | 16.56778 | 18.13606 | 5 | 5 | 5 | 8.8 | 4.89E+07 |
| 11.09309 | 12.81965 | 12.63365 | 4 | 4 | 4 | 15.4 | 4.88E+07 |
| 12.33639 | 19.97155 | 13.3257 | 3 | 3 | 3 | 9.3 | 4.85E+07 |
| 11.50685 | 20.16312 | 12.35648 | 3 | 3 | 3 | 8.9 | 4.85E+07 |
| 11.93458 | 11.36604 | 13.62096 | 4 | 4 | 4 | 16.6 | 4.85E+07 |
| 13.39178 | 13.52457 | 13.69878 | 9 | 9 | 9 | 2.6 | 4.84E+07 |
| 11.57838 | 17.54148 | 11.70184 | 7 | 7 | 7 | 13.5 | 4.83E+07 |
| 12.18719 | 13.1181 | 17.61798 | 3 | 3 | 3 | 12.9 | 4.83E+07 |
| 11.11609 | 19.9812 | 13.54342 | 5 | 5 | 5 | 5.9 | 4.82E+07 |
| 11.04512 | 14.59461 | 12.30236 | 5 | 5 | 5 | 5.2 | 4.81E+07 |
| 12.66405 | 14.03218 | 11.89371 | 4 | 4 | 4 | 9.3 | 4.81E+07 |
| 12.89073 | 18.30466 | 12.60033 | 3 | 3 | 3 | 32 | 4.79E+07 |
| 10.34604 | 14.55952 | 15.15846 | 5 | 5 | 5 | 27.2 | 4.78E+07 |
| 12.5018 | 19.33351 | 12.34251 | 4 | 4 | 4 | 18.7 | 4.75E+07 |
| 12.4592 | 12.85254 | 12.6051 | 5 | 5 | 5 | 19.3 | 4.74E+07 |
| 16.23369 | 16.88298 | 15.84757 | 4 | 4 | 4 | 12.8 | 4.72E+07 |
| 11.60489 | 14.38205 | 13.70048 | 6 | 6 | 6 | 10.1 | 4.70E+07 |
| 12.03972 | 14.57618 | 13.09613 | 4 | 4 | 4 | 5.2 | 4.70E+07 |
| 11.29222 | 14.75426 | 11.94822 | 5 | 5 | 5 | 5.7 | 4.69E+07 |
| 13.85304 | 19.46334 | 19.24379 | 4 | 4 | 4 | 7.3 | 4.65E+07 |
| 12.51436 | 20.86484 | 10.90483 | 3 | 3 | 3 | 14 | 4.65E+07 |
| 18.84064 | 18.61296 | 17.92295 | 3 | 3 | 3 | 20.5 | 4.64E+07 |
| 11.50687 | 13.49057 | 13.47937 | 5 | 5 | 5 | 11.8 | 4.61E+07 |
| 10.91074 | 17.95413 | 12.8688 | 5 | 5 | 5 | 12.5 | 4.61E+07 |
| 12.00328 | 14.23425 | 11.07918 | 4 | 4 | 4 | 2.5 | 4.60E+07 |
| 13.94424 | 18.43522 | 11.59507 | 5 | 5 | 5 | 7.2 | 4.60E+07 |
| 12.65711 | 12.86049 | 12.27862 | 3 | 3 | 3 | 7.3 | 4.59E+07 |
| 17.31416 | 18.70188 | 13.19475 | 3 | 3 | 3 | 11.3 | 4.57E+07 |
| 12.62325 | 13.14989 | 11.79998 | 4 | 4 | 4 | 15.4 | 4.55E+07 |
| 12.37604 | 12.27526 | 12.61748 | 3 | 3 | 3 | 10.6 | 4.55E+07 |
| 10.6989 | 12.70963 | 12.30607 | 2 | 2 | 2 | 4.6 | 4.53E+07 |
| 12.63076 | 16.67742 | 16.5478 | 3 | 3 | 3 | 10.1 | 4.53E+07 |
| 17.33289 | 14.76074 | 13.52792 | 6 | 6 | 6 | 14.9 | 4.52E+07 |

| | | | | | | | |
|---|---|---|---|---|---|---|---|
| 12.66364 | 12.61161 | 12.5559 | 6 | 6 | 6 | 7.3 | 4.52E+07 |
| 12.20496 | 22.51569 | 19.71168 | 3 | 3 | 3 | 14.3 | 4.52E+07 |
| 20.28568 | 21.80467 | 20.03349 | 3 | 3 | 3 | 6.9 | 4.51E+07 |
| 10.79527 | 13.19391 | 18.68808 | 9 | 9 | 9 | 10 | 4.51E+07 |
| 17.96933 | 18.64061 | 17.76344 | 5 | 5 | 5 | 4.7 | 4.50E+07 |
| 10.66514 | 14.04845 | 12.29806 | 3 | 3 | 3 | 35.4 | 4.49E+07 |
| 17.79912 | 12.77162 | 13.15694 | 7 | 7 | 7 | 11 | 4.48E+07 |
| 11.62202 | 13.65013 | 11.78564 | 5 | 5 | 5 | 20.2 | 4.48E+07 |
| 17.96308 | 12.97337 | 13.2911 | 6 | 3 | 3 | 7 | 4.48E+07 |
| 13.81982 | 13.96463 | 12.01455 | 2 | 2 | 2 | 5 | 4.47E+07 |
| 12.37273 | 13.55513 | 11.08628 | 5 | 5 | 5 | 16.6 | 4.46E+07 |
| 17.36905 | 17.80448 | 17.28048 | 5 | 5 | 5 | 6.6 | 4.46E+07 |
| 11.32724 | 19.59596 | 11.52648 | 10 | 10 | 10 | 7.6 | 4.42E+07 |
| 11.82726 | 14.46357 | 12.12958 | 3 | 3 | 2 | 8.2 | 4.41E+07 |
| 11.70068 | 13.8003 | 10.66287 | 3 | 3 | 3 | 27.1 | 4.40E+07 |
| 12.8887 | 19.07347 | 11.96218 | 3 | 3 | 3 | 11.4 | 4.40E+07 |
| 11.54359 | 19.01955 | 13.67876 | 3 | 3 | 3 | 5.9 | 4.40E+07 |
| 11.9349 | 13.27623 | 12.02091 | 2 | 2 | 2 | 6.2 | 4.39E+07 |
| 13.20289 | 18.85836 | 13.41279 | 7 | 7 | 7 | 5.4 | 4.39E+07 |
| 17.61734 | 12.53734 | 12.60851 | 2 | 2 | 2 | 10.6 | 4.39E+07 |
| 10.77503 | 13.47491 | 11.61217 | 4 | 4 | 4 | 8.7 | 4.37E+07 |
| 13.44637 | 14.62389 | 12.25767 | 4 | 4 | 4 | 16.1 | 4.33E+07 |
| 12.33909 | 17.68243 | 12.01778 | 2 | 2 | 2 | 6.2 | 4.33E+07 |
| 13.1486 | 17.41897 | 12.42817 | 4 | 4 | 4 | 10.8 | 4.31E+07 |
| 12.10741 | 14.03364 | 11.54745 | 3 | 3 | 3 | 8.9 | 4.31E+07 |
| 12.20036 | 14.72163 | 12.11366 | 25 | 2 | 2 | 51.9 | 4.30E+07 |
| 12.52955 | 14.85013 | 11.24714 | 3 | 3 | 3 | 13.6 | 4.28E+07 |
| 17.5075 | 20.66014 | 18.30955 | 3 | 3 | 3 | 25.7 | 4.27E+07 |
| 11.66601 | 13.38511 | 13.0538 | 2 | 2 | 2 | 6.8 | 4.27E+07 |
| 12.53063 | 12.64881 | 13.33664 | 4 | 4 | 4 | 8.3 | 4.27E+07 |
| 12.83068 | 12.11181 | 12.85491 | 3 | 3 | 3 | 11.7 | 4.27E+07 |
| 12.95867 | 13.50572 | 12.74033 | 4 | 4 | 4 | 5.8 | 4.26E+07 |
| 11.77474 | 13.83042 | 12.09378 | 4 | 4 | 4 | 14.3 | 4.24E+07 |
| 10.7322 | 24.29144 | 18.32536 | 7 | 7 | 7 | 26 | 4.23E+07 |
| 12.16543 | 13.13183 | 17.86629 | 3 | 3 | 3 | 19.4 | 4.23E+07 |
| 11.98134 | 12.84534 | 11.83331 | 6 | 6 | 6 | 19.6 | 4.22E+07 |
| 11.69306 | 22.76061 | 13.07904 | 3 | 3 | 3 | 22.4 | 4.22E+07 |
| 10.71895 | 16.7129 | 11.87847 | 4 | 4 | 4 | 27.9 | 4.21E+07 |
| 18.70022 | 11.84023 | 12.83063 | 4 | 4 | 4 | 19.8 | 4.20E+07 |
| 17.54026 | 17.89912 | 17.72061 | 7 | 7 | 7 | 9.6 | 4.20E+07 |
| 16.65396 | 19.12563 | 17.06329 | 4 | 4 | 4 | 26.1 | 4.17E+07 |
| 12.35406 | 13.70995 | 11.11684 | 5 | 5 | 5 | 20.4 | 4.17E+07 |
| 12.8491 | 15.01484 | 12.86336 | 3 | 3 | 3 | 9.3 | 4.16E+07 |
| 9.564939 | 15.54943 | 11.67599 | 2 | 2 | 2 | 13.7 | 4.16E+07 |
| 10.10171 | 12.88334 | 13.22103 | 5 | 4 | 3 | 18.9 | 4.15E+07 |
| 13.67023 | 19.64808 | 17.09042 | 5 | 5 | 5 | 10.9 | 4.14E+07 |
| 11.83754 | 14.46711 | 13.23931 | 4 | 3 | 3 | 21.7 | 4.14E+07 |
| 11.97807 | 13.57159 | 12.56784 | 3 | 3 | 3 | 21 | 4.14E+07 |

| | | | | | | | |
|---|---|---|---|---|---|---|---|
| 11.61127 | 14.24286 | 13.05948 | 5 | 5 | 5 | 6.7 | 4.07E+07 |
| 11.03885 | 14.61143 | 14.32232 | 3 | 3 | 3 | 18.1 | 4.06E+07 |
| 11.78476 | 14.27207 | 12.52233 | 3 | 3 | 3 | 18.8 | 4.06E+07 |
| 12.89693 | 12.62426 | 12.64453 | 5 | 5 | 5 | 44.7 | 4.05E+07 |
| 21.36041 | 14.48115 | 12.52576 | 5 | 5 | 5 | 13.2 | 4.05E+07 |
| 18.47235 | 19.17653 | 18.71099 | 5 | 5 | 5 | 25 | 4.05E+07 |
| 12.60204 | 12.51525 | 13.94753 | 3 | 3 | 3 | 12.7 | 4.03E+07 |
| 12.98944 | 19.77509 | 11.98048 | 5 | 5 | 5 | 28.8 | 4.02E+07 |
| 12.11146 | 12.09557 | 11.30759 | 3 | 3 | 3 | 9.7 | 4.01E+07 |
| 11.89298 | 20.3721 | 13.90101 | 3 | 3 | 3 | 30.5 | 4.00E+07 |
| 11.20578 | 14.74997 | 11.51525 | 4 | 4 | 4 | 16.2 | 4.00E+07 |
| 13.69588 | 19.28929 | 12.679 | 5 | 5 | 5 | 15.1 | 3.96E+07 |
| 19.36769 | 18.70816 | 12.18568 | 5 | 5 | 5 | 8.9 | 3.96E+07 |
| 11.02234 | 24.21542 | 17.36384 | 13 | 6 | 6 | 40.5 | 3.96E+07 |
| 12.67587 | 20.74172 | 12.65922 | 3 | 3 | 3 | 10.1 | 3.95E+07 |
| 13.17343 | 12.96576 | 13.6678 | 6 | 6 | 6 | 6.9 | 3.94E+07 |
| 13.47707 | 20.62614 | 12.42776 | 3 | 3 | 3 | 5.3 | 3.91E+07 |
| 11.97667 | 18.85053 | 12.00207 | 5 | 5 | 5 | 16.5 | 3.91E+07 |
| 19.75171 | 22.32215 | 18.68041 | 7 | 7 | 7 | 13 | 3.90E+07 |
| 12.16126 | 20.16349 | 12.78736 | 2 | 2 | 2 | 19.6 | 3.88E+07 |
| 12.1095 | 13.5432 | 13.48724 | 6 | 6 | 6 | 15.2 | 3.85E+07 |
| 11.74762 | 13.9191 | 10.27227 | 4 | 4 | 4 | 11.4 | 3.85E+07 |
| 12.28981 | 12.89883 | 12.76272 | 3 | 3 | 3 | 17.1 | 3.85E+07 |
| 12.00096 | 13.70021 | 10.69133 | 3 | 3 | 3 | 4.1 | 3.85E+07 |
| 11.83603 | 12.96613 | 18.74396 | 2 | 2 | 2 | 6.6 | 3.84E+07 |
| 12.07623 | 14.28759 | 12.53398 | 12 | 6 | 6 | 19 | 3.84E+07 |
| 19.09473 | 12.58728 | 11.31481 | 3 | 3 | 3 | 8.5 | 3.80E+07 |
| 13.29155 | 13.41334 | 11.2823 | 5 | 5 | 5 | 6.5 | 3.80E+07 |
| 12.48535 | 13.41955 | 12.31211 | 4 | 4 | 4 | 6.6 | 3.80E+07 |
| 22.50327 | 20.64685 | 22.33677 | 3 | 3 | 3 | 4.8 | 3.79E+07 |
| 12.89775 | 13.94296 | 13.02595 | 3 | 3 | 3 | 13.1 | 3.79E+07 |
| 12.39979 | 14.23236 | 23.89171 | 2 | 2 | 2 | 1.5 | 3.78E+07 |
| 12.49839 | 19.87325 | 12.74689 | 3 | 3 | 3 | 14.6 | 3.78E+07 |
| 12.50049 | 19.75037 | 10.59254 | 5 | 5 | 5 | 10.4 | 3.77E+07 |
| 10.11273 | 16.52851 | 15.60678 | 5 | 5 | 5 | 18.8 | 3.76E+07 |
| 12.28099 | 13.35434 | 12.68857 | 3 | 3 | 3 | 14.9 | 3.75E+07 |
| 13.35054 | 14.02507 | 12.89884 | 12 | 3 | 3 | 33.9 | 3.75E+07 |
| 12.34296 | 14.80326 | 12.4338 | 3 | 3 | 3 | 14.1 | 3.74E+07 |
| 10.65963 | 19.93948 | 14.05998 | 3 | 3 | 2 | 26.3 | 3.73E+07 |
| 11.16871 | 12.17261 | 11.43966 | 3 | 3 | 3 | 10.4 | 3.73E+07 |
| 12.49775 | 12.61288 | 12.47288 | 3 | 3 | 2 | 10.7 | 3.72E+07 |
| 11.72737 | 20.13496 | 11.57721 | 4 | 4 | 2 | 17.8 | 3.71E+07 |
| 12.38747 | 13.68362 | 11.2645 | 4 | 4 | 4 | 12.8 | 3.70E+07 |
| 11.77082 | 14.3394 | 14.17994 | 6 | 6 | 6 | 10.8 | 3.69E+07 |
| 11.84809 | 13.16183 | 12.3738 | 6 | 6 | 6 | 6.6 | 3.68E+07 |
| 20.51874 | 20.65535 | 20.58206 | 4 | 4 | 4 | 7.6 | 3.67E+07 |
| 12.64368 | 15.03698 | 12.75948 | 3 | 3 | 3 | 12.5 | 3.66E+07 |
| 11.89284 | 13.68266 | 12.67871 | 3 | 3 | 3 | 35.9 | 3.66E+07 |

| | | | | | | | |
|---|---|---|---|---|---|---|---|
| 11.67826 | 12.26621 | 13.23804 | 5 | 5 | 5 | 14.9 | 3.66E+07 |
| 12.18935 | 13.92113 | 11.35805 | 4 | 4 | 4 | 14.9 | 3.66E+07 |
| 12.96004 | 12.93819 | 13.38005 | 2 | 2 | 2 | 11.3 | 3.66E+07 |
| 19.23269 | 17.73026 | 11.33637 | 5 | 5 | 5 | 10.5 | 3.64E+07 |
| 20.29804 | 18.11614 | 10.46708 | 2 | 2 | 2 | 8.4 | 3.63E+07 |
| 13.42819 | 16.93572 | 9.970492 | 2 | 2 | 2 | 7.3 | 3.62E+07 |
| 19.38296 | 22.33694 | 19.72197 | 3 | 3 | 3 | 4.4 | 3.61E+07 |
| 11.75393 | 13.34329 | 11.0936 | 4 | 4 | 4 | 5.7 | 3.61E+07 |
| 11.07144 | 20.96825 | 14.66108 | 2 | 2 | 2 | 8 | 3.59E+07 |
| 10.57242 | 12.43399 | 11.47536 | 4 | 4 | 4 | 4.3 | 3.57E+07 |
| 12.43158 | 18.2322 | 17.71539 | 3 | 3 | 3 | 8.3 | 3.57E+07 |
| 12.17848 | 13.43678 | 11.02473 | 3 | 3 | 3 | 6.3 | 3.54E+07 |
| 10.26954 | 13.4857 | 11.59247 | 7 | 7 | 7 | 4.1 | 3.53E+07 |
| 11.51427 | 13.74683 | 12.48966 | 5 | 5 | 5 | 6.1 | 3.53E+07 |
| 11.54575 | 12.15361 | 12.25603 | 2 | 2 | 2 | 9.7 | 3.53E+07 |
| 18.96452 | 19.46695 | 18.9626 | 4 | 4 | 4 | 9.1 | 3.52E+07 |
| 11.12774 | 14.35801 | 11.81042 | 5 | 5 | 5 | 2.3 | 3.51E+07 |
| 12.24835 | 13.42578 | 12.3942 | 4 | 4 | 4 | 5.8 | 3.51E+07 |
| 10.98365 | 18.40183 | 13.99999 | 2 | 2 | 1 | 10.2 | 3.48E+07 |
| 11.29397 | 15.27424 | 11.69393 | 2 | 2 | 2 | 4.6 | 3.48E+07 |
| 10.75571 | 12.7088 | 13.18378 | 5 | 5 | 5 | 15.6 | 3.47E+07 |
| 13.34717 | 14.6059 | 12.67324 | 3 | 3 | 3 | 8.4 | 3.47E+07 |
| 11.93096 | 20.02958 | 11.30042 | 4 | 4 | 4 | 8 | 3.45E+07 |
| 11.83 | 21.09288 | 16.96093 | 2 | 2 | 2 | 17.3 | 3.44E+07 |
| 12.00184 | 13.0993 | 11.82744 | 3 | 3 | 3 | 8.4 | 3.42E+07 |
| 13.41204 | 13.15841 | 11.2656 | 6 | 4 | 4 | 9.4 | 3.42E+07 |
| 17.42939 | 24.07896 | 10.92845 | 9 | 9 | 9 | 23.7 | 3.40E+07 |
| 18.07838 | 18.67142 | 13.21942 | 4 | 4 | 4 | 7.6 | 3.39E+07 |
| 10.8506 | 13.47208 | 12.55715 | 3 | 3 | 3 | 4 | 3.39E+07 |
| 13.51591 | 17.85172 | 12.76023 | 4 | 4 | 4 | 5 | 3.38E+07 |
| 12.10138 | 12.98719 | 13.72632 | 4 | 4 | 4 | 1.8 | 3.38E+07 |
| 12.52353 | 13.79378 | 11.54511 | 5 | 5 | 5 | 15.8 | 3.37E+07 |
| 13.72945 | 14.34963 | 13.59771 | 2 | 2 | 2 | 11 | 3.37E+07 |
| 13.41497 | 14.20459 | 11.91332 | 3 | 3 | 3 | 9.6 | 3.37E+07 |
| 18.02172 | 18.95886 | 19.16194 | 4 | 4 | 4 | 5.4 | 3.37E+07 |
| 11.37616 | 13.51511 | 12.09089 | 4 | 4 | 4 | 26.5 | 3.35E+07 |
| 15.11235 | 14.81199 | 10.65098 | 5 | 5 | 5 | 6.8 | 3.34E+07 |
| 11.58708 | 13.63214 | 10.58932 | 4 | 4 | 4 | 14 | 3.31E+07 |
| 12.68832 | 14.85474 | 13.12257 | 3 | 3 | 3 | 11.5 | 3.28E+07 |
| 10.35958 | 17.19499 | 11.2828 | 4 | 4 | 4 | 5.8 | 3.28E+07 |
| 11.52951 | 13.56899 | 10.52891 | 3 | 3 | 3 | 23.6 | 3.28E+07 |
| 12.17322 | 18.29536 | 12.57033 | 5 | 5 | 5 | 22.3 | 3.27E+07 |
| 19.02426 | 23.88558 | 19.12528 | 3 | 3 | 3 | 33.3 | 3.26E+07 |
| 11.25899 | 12.96859 | 12.30557 | 4 | 4 | 4 | 6.8 | 3.26E+07 |
| 10.66649 | 16.29726 | 12.59419 | 3 | 3 | 3 | 7.9 | 3.26E+07 |
| 11.99694 | 17.45403 | 11.63971 | 5 | 4 | 4 | 5.4 | 3.26E+07 |
| 13.1269 | 13.06976 | 11.20412 | 4 | 4 | 4 | 10.3 | 3.26E+07 |
| 11.34753 | 13.20655 | 11.32854 | 4 | 4 | 4 | 24 | 3.26E+07 |

| | | | | | | | |
|---|---|---|---|---|---|---|---|
| 12.19714 | 18.35234 | 12.77088 | 3 | 3 | 3 | 3.9 | 3.26E+07 |
| 11.82608 | 13.93267 | 12.04494 | 4 | 4 | 4 | 6.2 | 3.24E+07 |
| 18.28148 | 18.2028 | 18.13074 | 4 | 4 | 4 | 4.8 | 3.24E+07 |
| 13.54897 | 19.70215 | 12.90607 | 4 | 4 | 4 | 12 | 3.20E+07 |
| 12.69243 | 13.64461 | 11.23773 | 2 | 2 | 2 | 8.8 | 3.20E+07 |
| 11.71401 | 13.69276 | 13.78339 | 3 | 3 | 3 | 5.9 | 3.19E+07 |
| 11.91041 | 22.49782 | 12.86404 | 4 | 4 | 4 | 16.7 | 3.18E+07 |
| 11.19136 | 13.3676 | 12.11908 | 3 | 3 | 3 | 8 | 3.18E+07 |
| 11.91633 | 12.21948 | 21.14439 | 3 | 2 | 2 | 8.4 | 3.17E+07 |
| 11.50806 | 20.01141 | 11.52467 | 3 | 3 | 3 | 8.8 | 3.16E+07 |
| 11.35168 | 19.15031 | 13.83169 | 6 | 6 | 6 | 2.1 | 3.14E+07 |
| 11.78335 | 12.28384 | 11.17735 | 4 | 4 | 4 | 26.2 | 3.13E+07 |
| 19.90064 | 23.16704 | 20.37994 | 2 | 2 | 2 | 32.2 | 3.13E+07 |
| 19.52043 | 13.11151 | 20.24725 | 2 | 2 | 2 | 12.7 | 3.12E+07 |
| 12.75189 | 13.08166 | 10.45762 | 3 | 3 | 3 | 8.6 | 3.11E+07 |
| 11.80093 | 11.33077 | 20.77965 | 3 | 3 | 3 | 19.6 | 3.11E+07 |
| 11.6734 | 12.22835 | 12.06263 | 2 | 2 | 2 | 40 | 3.11E+07 |
| 19.95234 | 19.07752 | 12.25213 | 4 | 4 | 4 | 41.8 | 3.11E+07 |
| 13.42518 | 18.07426 | 11.35671 | 5 | 5 | 5 | 8.3 | 3.08E+07 |
| 10.74779 | 13.86893 | 19.07028 | 5 | 5 | 5 | 7.2 | 3.07E+07 |
| 11.24326 | 12.66477 | 11.56325 | 4 | 4 | 4 | 11.8 | 3.04E+07 |
| 14.19675 | 12.41881 | 11.35291 | 4 | 4 | 4 | 6.8 | 3.04E+07 |
| 13.17637 | 12.99085 | 10.96915 | 3 | 3 | 3 | 11.9 | 3.03E+07 |
| 10.26458 | 22.76586 | 13.07454 | 5 | 5 | 5 | 26.4 | 3.03E+07 |
| 11.84847 | 13.4506 | 11.52679 | 2 | 2 | 2 | 13.2 | 3.03E+07 |
| 18.05674 | 17.54894 | 17.36828 | 4 | 4 | 4 | 10.5 | 3.03E+07 |
| 19.59661 | 21.01849 | 18.96635 | 3 | 3 | 3 | 14 | 3.02E+07 |
| 12.51462 | 13.3083 | 13.01268 | 2 | 2 | 2 | 3.6 | 3.01E+07 |
| 13.09046 | 12.83286 | 12.00618 | 5 | 5 | 5 | 3.4 | 3.00E+07 |
| 12.73357 | 14.43747 | 10.54227 | 6 | 6 | 6 | 11.7 | 3.00E+07 |
| 12.7945 | 12.18089 | 12.12827 | 2 | 2 | 2 | 10.9 | 2.99E+07 |
| 11.82051 | 13.32979 | 12.27218 | 2 | 2 | 2 | 6.8 | 2.99E+07 |
| 12.3527 | 13.63988 | 11.70393 | 3 | 3 | 3 | 8.9 | 2.99E+07 |
| 18.90319 | 23.15409 | 19.76875 | 4 | 4 | 4 | 37.5 | 2.98E+07 |
| 12.22483 | 22.77768 | 12.68761 | 11 | 11 | 11 | 16.2 | 2.97E+07 |
| 12.33154 | 16.87316 | 12.95272 | 5 | 5 | 5 | 2.1 | 2.97E+07 |
| 12.29974 | 12.51275 | 12.44503 | 3 | 3 | 3 | 9.4 | 2.97E+07 |
| 12.10106 | 14.15766 | 12.53268 | 2 | 2 | 2 | 9.1 | 2.95E+07 |
| 11.81299 | 20.97134 | 14.10565 | 2 | 2 | 2 | 15.1 | 2.92E+07 |
| 12.56778 | 13.12252 | 12.94407 | 7 | 5 | 5 | 5.4 | 2.91E+07 |
| 13.93433 | 15.08107 | 12.16877 | 2 | 2 | 2 | 3.6 | 2.90E+07 |
| 21.63321 | 21.08213 | 20.74887 | 2 | 2 | 2 | 8.2 | 2.89E+07 |
| 13.63501 | 13.45908 | 10.54369 | 2 | 2 | 2 | 3.7 | 2.89E+07 |
| 13.46528 | 13.58911 | 10.47274 | 2 | 2 | 2 | 4.9 | 2.88E+07 |
| 11.51805 | 12.67157 | 12.67181 | 4 | 4 | 4 | 19 | 2.87E+07 |
| 9.829165 | 18.93064 | 12.86475 | 4 | 4 | 4 | 5.7 | 2.87E+07 |
| 18.59914 | 18.43726 | 12.65333 | 4 | 4 | 4 | 4.6 | 2.87E+07 |
| 10.59133 | 12.44824 | 11.20135 | 2 | 2 | 2 | 13.7 | 2.81E+07 |

| | | | | | | | |
|---|---|---|---|---|---|---|---|
| 10.8828 | 19.42714 | 12.9534 | 4 | 4 | 4 | 4.8 | 2.81E+07 |
| 11.63579 | 17.62257 | 11.53936 | 5 | 5 | 5 | 19.9 | 2.80E+07 |
| 21.12969 | 23.3098 | 12.79992 | 3 | 2 | 2 | 6.2 | 2.80E+07 |
| 12.65534 | 20.22529 | 13.11106 | 3 | 3 | 3 | 7.3 | 2.78E+07 |
| 12.74254 | 12.86298 | 13.55394 | 2 | 2 | 2 | 4.5 | 2.77E+07 |
| 13.32442 | 12.6811 | 12.06191 | 3 | 3 | 3 | 14.8 | 2.77E+07 |
| 12.5713 | 13.00113 | 12.03027 | 3 | 3 | 3 | 14.6 | 2.76E+07 |
| 12.20154 | 19.56856 | 12.65357 | 3 | 3 | 3 | 13.8 | 2.74E+07 |
| 12.33088 | 12.37875 | 11.88328 | 2 | 2 | 2 | 24.2 | 2.73E+07 |
| 12.45793 | 15.80935 | 10.57802 | 4 | 4 | 4 | 4.3 | 2.73E+07 |
| 14.48646 | 12.42959 | 11.03315 | 4 | 4 | 4 | 7.6 | 2.73E+07 |
| 12.91795 | 13.49608 | 11.19083 | 3 | 3 | 3 | 9.3 | 2.72E+07 |
| 12.93273 | 12.38211 | 12.87548 | 3 | 3 | 3 | 3.3 | 2.72E+07 |
| 11.1566 | 13.94493 | 11.57317 | 3 | 3 | 3 | 8.9 | 2.72E+07 |
| 13.66159 | 12.50731 | 13.43069 | 4 | 4 | 4 | 7.5 | 2.71E+07 |
| 11.12689 | 13.30249 | 11.80433 | 3 | 3 | 3 | 6.9 | 2.70E+07 |
| 11.10979 | 13.53142 | 12.16564 | 2 | 2 | 2 | 7 | 2.69E+07 |
| 20.26568 | 20.40655 | 20.31152 | 3 | 3 | 3 | 2 | 2.68E+07 |
| 10.59056 | 11.93178 | 13.27332 | 3 | 2 | 2 | 2.2 | 2.68E+07 |
| 12.43881 | 12.42395 | 10.02987 | 3 | 3 | 3 | 18.2 | 2.68E+07 |
| 10.76896 | 14.61928 | 13.16149 | 2 | 2 | 2 | 2.6 | 2.67E+07 |
| 10.87239 | 13.71129 | 12.22036 | 2 | 2 | 2 | 12.8 | 2.67E+07 |
| 11.5694 | 12.20532 | 13.53538 | 7 | 3 | 3 | 11 | 2.66E+07 |
| 14.36917 | 14.2595 | 12.41989 | 3 | 3 | 3 | 15.5 | 2.65E+07 |
| 13.44688 | 14.73707 | 11.27775 | 2 | 2 | 2 | 5.6 | 2.65E+07 |
| 11.43204 | 11.95685 | 11.9391 | 4 | 4 | 4 | 4.2 | 2.65E+07 |
| 14.14244 | 14.38535 | 12.1875 | 4 | 3 | 3 | 9.7 | 2.64E+07 |
| 11.54921 | 18.48463 | 12.09624 | 2 | 2 | 2 | 12.6 | 2.63E+07 |
| 13.18944 | 19.79028 | 12.9173 | 4 | 4 | 4 | 9.8 | 2.62E+07 |
| 13.9749 | 13.43681 | 10.63332 | 4 | 4 | 4 | 6.1 | 2.62E+07 |
| 12.04315 | 12.38677 | 12.16959 | 20 | 3 | 3 | 57 | 2.62E+07 |
| 12.2261 | 18.04457 | 11.17704 | 2 | 2 | 2 | 23.8 | 2.61E+07 |
| 11.67554 | 13.65654 | 11.04594 | 2 | 2 | 2 | 3.3 | 2.61E+07 |
| 11.6835 | 14.27824 | 10.56624 | 3 | 3 | 3 | 5.3 | 2.61E+07 |
| 11.7607 | 22.47459 | 12.08701 | 4 | 4 | 4 | 6.6 | 2.60E+07 |
| 12.72856 | 17.64836 | 12.75905 | 2 | 2 | 2 | 5.5 | 2.60E+07 |
| 19.60484 | 14.26782 | 12.49029 | 2 | 2 | 2 | 26.7 | 2.59E+07 |
| 12.87973 | 13.58197 | 11.61256 | 3 | 3 | 3 | 9.6 | 2.59E+07 |
| 18.6648 | 13.85283 | 18.32685 | 4 | 4 | 4 | 8.3 | 2.56E+07 |
| 15.98666 | 16.79728 | 16.44854 | 3 | 3 | 3 | 11.7 | 2.55E+07 |
| 13.08963 | 11.75924 | 13.40589 | 2 | 2 | 2 | 3.2 | 2.55E+07 |
| 12.10919 | 12.41893 | 12.71556 | 3 | 3 | 3 | 16.2 | 2.55E+07 |
| 13.48164 | 13.80243 | 11.55413 | 3 | 3 | 3 | 5.9 | 2.54E+07 |
| 12.11246 | 14.36191 | 13.35288 | 3 | 3 | 3 | 18.7 | 2.53E+07 |
| 12.77991 | 13.58364 | 12.80159 | 5 | 5 | 2 | 3 | 2.52E+07 |
| 11.81026 | 14.68154 | 12.48294 | 4 | 4 | 4 | 10.2 | 2.52E+07 |
| 19.32279 | 20.4708 | 19.3117 | 2 | 2 | 2 | 9.3 | 2.51E+07 |
| 12.90869 | 13.47106 | 12.3808 | 2 | 2 | 2 | 11.3 | 2.49E+07 |

| | | | | | | | |
|---|---|---|---|---|---|---|---|
| 11.85905 | 14.26528 | 13.37072 | 3 | 3 | 3 | 9.6 | 2.49E+07 |
| 13.39771 | 13.12744 | 11.86872 | 2 | 2 | 2 | 10.2 | 2.49E+07 |
| 10.99213 | 16.05922 | 17.92851 | 2 | 2 | 2 | 8.6 | 2.47E+07 |
| 10.99128 | 12.70632 | 13.16216 | 6 | 3 | 3 | 11.4 | 2.47E+07 |
| 12.09225 | 13.81132 | 11.72948 | 4 | 4 | 4 | 5.7 | 2.45E+07 |
| 11.51765 | 14.14709 | 13.20384 | 2 | 2 | 2 | 6.3 | 2.45E+07 |
| 12.07015 | 21.07725 | 20.85018 | 4 | 4 | 4 | 20.6 | 2.44E+07 |
| 12.13403 | 12.42343 | 11.53964 | 5 | 2 | 2 | 12.8 | 2.41E+07 |
| 11.17179 | 14.94633 | 11.79546 | 3 | 3 | 3 | 3.8 | 2.39E+07 |
| 10.5263 | 21.30875 | 14.09102 | 3 | 3 | 3 | 42.3 | 2.39E+07 |
| 12.98431 | 12.44567 | 13.04833 | 2 | 2 | 2 | 11.3 | 2.39E+07 |
| 12.45895 | 14.68151 | 11.94161 | 3 | 3 | 3 | 4.2 | 2.39E+07 |
| 14.0559 | 19.37267 | 11.51763 | 3 | 3 | 3 | 2.8 | 2.38E+07 |
| 12.83367 | 12.38535 | 11.66988 | 2 | 2 | 2 | 6.3 | 2.38E+07 |
| 18.24975 | 13.3434 | 11.89273 | 2 | 2 | 2 | 4 | 2.38E+07 |
| 11.97747 | 13.72119 | 11.89287 | 3 | 3 | 3 | 3.9 | 2.37E+07 |
| 11.92092 | 21.09397 | 18.01438 | 6 | 4 | 4 | 8.5 | 2.37E+07 |
| 10.39927 | 13.95342 | 12.67094 | 2 | 2 | 2 | 19.9 | 2.37E+07 |
| 11.36713 | 15.80479 | 15.63691 | 4 | 4 | 4 | 6.9 | 2.36E+07 |
| 11.74231 | 20.44849 | 13.92439 | 4 | 3 | 3 | 8.1 | 2.36E+07 |
| 12.88608 | 13.67913 | 11.47604 | 2 | 2 | 2 | 6.9 | 2.36E+07 |
| 12.00346 | 12.8466 | 11.0919 | 4 | 4 | 4 | 5.9 | 2.35E+07 |
| 11.69481 | 12.53931 | 12.73921 | 4 | 4 | 4 | 5.5 | 2.35E+07 |
| 13.14959 | 14.20393 | 13.13959 | 4 | 4 | 4 | 5.5 | 2.34E+07 |
| 12.23472 | 12.03349 | 12.52252 | 5 | 5 | 5 | 7.3 | 2.33E+07 |
| 11.4478 | 21.73269 | 11.30805 | 4 | 4 | 4 | 5.6 | 2.33E+07 |
| 12.92932 | 22.3633 | 13.91756 | 6 | 6 | 6 | 24 | 2.32E+07 |
| 13.38622 | 19.75274 | 12.20018 | 3 | 3 | 3 | 11.5 | 2.32E+07 |
| 12.5076 | 19.14554 | 12.64273 | 3 | 3 | 3 | 19.4 | 2.32E+07 |
| 12.70345 | 12.44756 | 12.89075 | 4 | 4 | 4 | 9 | 2.31E+07 |
| 11.11835 | 13.97702 | 11.72031 | 2 | 2 | 2 | 2.5 | 2.31E+07 |
| 12.64257 | 20.18326 | 11.18758 | 2 | 2 | 2 | 12.4 | 2.30E+07 |
| 11.07016 | 13.65713 | 11.99626 | 4 | 4 | 4 | 12.9 | 2.30E+07 |
| 13.00078 | 12.37126 | 11.70304 | 3 | 3 | 3 | 9.2 | 2.28E+07 |
| 12.8551 | 13.23357 | 12.4934 | 2 | 2 | 2 | 12.6 | 2.28E+07 |
| 12.9591 | 12.52088 | 19.43123 | 3 | 3 | 3 | 4.4 | 2.28E+07 |
| 9.76375 | 19.33595 | 12.10188 | 2 | 2 | 2 | 16.9 | 2.26E+07 |
| 16.10042 | 11.54049 | 11.59132 | 6 | 4 | 4 | 16.4 | 2.26E+07 |
| 12.08577 | 13.50914 | 13.42538 | 2 | 2 | 2 | 11.3 | 2.26E+07 |
| 12.96076 | 19.90528 | 13.25593 | 5 | 5 | 5 | 3.8 | 2.26E+07 |
| 11.45591 | 14.56036 | 12.28633 | 3 | 3 | 3 | 34.6 | 2.25E+07 |
| 11.75606 | 13.59713 | 12.28389 | 2 | 2 | 2 | 12 | 2.24E+07 |
| 19.72219 | 12.2403 | 13.16868 | 2 | 2 | 2 | 7 | 2.23E+07 |
| 19.4477 | 19.12825 | 18.60816 | 2 | 2 | 2 | 9.7 | 2.23E+07 |
| 11.65143 | 17.21162 | 11.53219 | 2 | 2 | 2 | 32.8 | 2.23E+07 |
| 11.78355 | 14.77662 | 12.70355 | 2 | 2 | 2 | 9.5 | 2.22E+07 |
| 19.11929 | 20.43074 | 19.92104 | 2 | 2 | 2 | 12.1 | 2.21E+07 |
| 12.15544 | 13.1582 | 12.42877 | 2 | 2 | 2 | 12.3 | 2.21E+07 |

| | | | | | | | |
|---|---|---|---|---|---|---|---|
| 12.68143 | 12.64456 | 11.85382 | 2 | 2 | 2 | 10 | 2.21E+07 |
| 17.41221 | 18.16889 | 12.19823 | 2 | 2 | 2 | 6.4 | 2.21E+07 |
| 10.83428 | 12.82574 | 12.20636 | 3 | 3 | 3 | 7.4 | 2.20E+07 |
| 11.25099 | 12.52318 | 12.57024 | 3 | 3 | 3 | 12 | 2.19E+07 |
| 11.01181 | 22.00285 | 11.80915 | 5 | 5 | 5 | 25.7 | 2.19E+07 |
| 11.73484 | 17.70718 | 16.93698 | 2 | 2 | 2 | 9.3 | 2.19E+07 |
| 12.88856 | 21.15542 | 17.12767 | 2 | 2 | 2 | 12 | 2.19E+07 |
| 17.76914 | 18.25962 | 17.7613 | 2 | 2 | 2 | 8.8 | 2.18E+07 |
| 13.01501 | 18.02799 | 13.10967 | 3 | 3 | 3 | 10.8 | 2.17E+07 |
| 11.62166 | 12.25632 | 11.83016 | 2 | 2 | 2 | 8.4 | 2.15E+07 |
| 13.25817 | 14.60215 | 11.96376 | 3 | 3 | 3 | 29.7 | 2.15E+07 |
| 11.846 | 12.97233 | 13.13785 | 2 | 2 | 2 | 6.4 | 2.14E+07 |
| 13.76797 | 14.94861 | 12.19147 | 3 | 3 | 3 | 5.2 | 2.14E+07 |
| 12.04747 | 13.25941 | 11.41085 | 2 | 2 | 2 | 5.2 | 2.14E+07 |
| 11.64063 | 13.03917 | 12.8474 | 3 | 3 | 3 | 19.2 | 2.14E+07 |
| 12.56783 | 13.63143 | 10.38309 | 3 | 3 | 3 | 5.6 | 2.13E+07 |
| 11.91648 | 19.30993 | 12.44927 | 2 | 2 | 2 | 7.5 | 2.12E+07 |
| 12.62054 | 12.49812 | 12.67083 | 5 | 3 | 3 | 11.2 | 2.11E+07 |
| 12.23673 | 13.20774 | 12.26752 | 3 | 3 | 3 | 6 | 2.11E+07 |
| 11.52692 | 17.66331 | 12.11775 | 2 | 2 | 2 | 2.3 | 2.10E+07 |
| 13.26774 | 13.778 | 11.97357 | 2 | 2 | 2 | 4.6 | 2.10E+07 |
| 10.76614 | 14.17339 | 10.14679 | 2 | 2 | 2 | 7.7 | 2.09E+07 |
| 11.12925 | 10.69453 | 12.99474 | 3 | 3 | 3 | 6.1 | 2.07E+07 |
| 13.46042 | 13.60334 | 10.84787 | 4 | 4 | 4 | 4 | 2.05E+07 |
| 13.08146 | 11.35876 | 9.622259 | 2 | 2 | 2 | 7.4 | 2.05E+07 |
| 11.0472 | 17.18389 | 11.54811 | 3 | 3 | 3 | 7.9 | 2.05E+07 |
| 22.44038 | 21.30296 | 13.20533 | 4 | 2 | 2 | 16.4 | 2.03E+07 |
| 11.77594 | 12.01116 | 11.07877 | 4 | 4 | 4 | 11 | 2.02E+07 |
| 11.46955 | 13.43547 | 11.22773 | 2 | 2 | 2 | 7.1 | 2.00E+07 |
| 13.28723 | 18.83399 | 11.75817 | 2 | 2 | 2 | 5.1 | 1.99E+07 |
| 12.79905 | 19.35997 | 11.91155 | 2 | 2 | 2 | 8.5 | 1.99E+07 |
| 20.82788 | 13.60944 | 21.19214 | 3 | 3 | 3 | 8.5 | 1.99E+07 |
| 11.68905 | 13.07907 | 12.50961 | 2 | 2 | 2 | 3.2 | 1.99E+07 |
| 12.34365 | 15.17937 | 12.59919 | 4 | 4 | 4 | 13 | 1.98E+07 |
| 12.37321 | 12.47059 | 11.72571 | 5 | 5 | 5 | 9.5 | 1.98E+07 |
| 10.8705 | 15.29991 | 13.08093 | 2 | 2 | 2 | 8.4 | 1.97E+07 |
| 11.43819 | 19.14617 | 13.54541 | 3 | 3 | 3 | 28.3 | 1.97E+07 |
| 11.91106 | 21.39488 | 11.26051 | 3 | 3 | 3 | 10.9 | 1.96E+07 |
| 9.915457 | 19.46963 | 11.42578 | 2 | 2 | 2 | 7.9 | 1.96E+07 |
| 12.10323 | 15.11272 | 12.58547 | 2 | 2 | 2 | 8 | 1.94E+07 |
| 14.20492 | 12.35235 | 11.08239 | 4 | 4 | 4 | 7 | 1.94E+07 |
| 19.88323 | 22.48417 | 20.25292 | 2 | 2 | 1 | 11.2 | 1.93E+07 |
| 12.58247 | 14.44219 | 12.92829 | 3 | 3 | 3 | 6.7 | 1.92E+07 |
| 12.41439 | 13.91763 | 10.28472 | 2 | 2 | 2 | 3.4 | 1.92E+07 |
| 11.4338 | 13.78538 | 12.54832 | 5 | 5 | 5 | 5.4 | 1.91E+07 |
| 11.86731 | 13.34645 | 21.38501 | 2 | 2 | 2 | 12.3 | 1.91E+07 |
| 11.19552 | 13.13528 | 11.4982 | 2 | 2 | 2 | 5.9 | 1.91E+07 |
| 12.67833 | 13.45526 | 11.70995 | 2 | 2 | 2 | 8.4 | 1.91E+07 |

| | | | | | | | |
|---|---|---|---|---|---|---|---|
| 12.03025 | 13.04521 | 13.17802 | 3 | 3 | 3 | 17.9 | 1.90E+07 |
| 11.94808 | 13.40054 | 12.9893 | 2 | 2 | 2 | 5.1 | 1.90E+07 |
| 10.73582 | 18.34208 | 11.27836 | 3 | 2 | 2 | 2.6 | 1.88E+07 |
| 12.07502 | 19.32974 | 11.97909 | 2 | 2 | 2 | 12.5 | 1.88E+07 |
| 12.21043 | 12.84564 | 11.51131 | 4 | 4 | 4 | 5.4 | 1.88E+07 |
| 12.05583 | 12.05408 | 12.31484 | 4 | 4 | 4 | 4.4 | 1.87E+07 |
| 12.28664 | 14.30584 | 11.92833 | 2 | 2 | 2 | 4.9 | 1.87E+07 |
| 12.39405 | 14.74728 | 12.78115 | 3 | 2 | 2 | 8.5 | 1.86E+07 |
| 11.33822 | 14.17291 | 12.06315 | 4 | 4 | 4 | 5.1 | 1.86E+07 |
| 11.74339 | 13.14577 | 11.44308 | 4 | 3 | 3 | 7.4 | 1.86E+07 |
| 12.33249 | 14.04792 | 11.69785 | 4 | 4 | 4 | 3.3 | 1.85E+07 |
| 16.56212 | 17.53039 | 16.19963 | 2 | 2 | 2 | 6.6 | 1.85E+07 |
| 11.88807 | 20.08589 | 11.37013 | 3 | 2 | 2 | 17 | 1.82E+07 |
| 12.83826 | 14.36784 | 14.72838 | 3 | 3 | 3 | 6.6 | 1.81E+07 |
| 19.87518 | 12.61601 | 11.67257 | 6 | 3 | 3 | 33.5 | 1.80E+07 |
| 11.54047 | 13.88055 | 11.77746 | 2 | 2 | 2 | 3.6 | 1.79E+07 |
| 12.40806 | 12.74488 | 10.56146 | 3 | 3 | 3 | 16.7 | 1.78E+07 |
| 11.15287 | 12.23766 | 10.81986 | 3 | 3 | 3 | 3.2 | 1.78E+07 |
| 11.17142 | 14.99178 | 13.1 | 3 | 2 | 2 | 6.5 | 1.77E+07 |
| 12.55103 | 12.2192 | 13.52841 | 2 | 2 | 2 | 14.4 | 1.77E+07 |
| 11.95958 | 12.1394 | 11.68265 | 3 | 3 | 3 | 8.1 | 1.77E+07 |
| 12.90101 | 12.87121 | 13.73556 | 4 | 4 | 4 | 3.3 | 1.77E+07 |
| 10.79889 | 20.97897 | 13.37792 | 2 | 2 | 2 | 14.1 | 1.76E+07 |
| 13.06849 | 13.85133 | 12.13807 | 2 | 2 | 2 | 1.5 | 1.76E+07 |
| 11.96747 | 13.49283 | 13.07633 | 3 | 3 | 3 | 8.1 | 1.76E+07 |
| 12.70262 | 18.99857 | 12.28832 | 3 | 3 | 3 | 8.2 | 1.75E+07 |
| 12.18687 | 13.03218 | 12.93645 | 2 | 2 | 2 | 12 | 1.74E+07 |
| 11.96163 | 12.29698 | 13.57978 | 2 | 2 | 2 | 2.9 | 1.72E+07 |
| 11.839 | 13.33452 | 10.70976 | 3 | 3 | 3 | 5 | 1.72E+07 |
| 12.65308 | 13.28623 | 14.24489 | 2 | 2 | 2 | 4 | 1.72E+07 |
| 12.33886 | 12.95884 | 10.97321 | 3 | 3 | 3 | 19.8 | 1.72E+07 |
| 17.49997 | 13.7361 | 17.6235 | 2 | 2 | 2 | 11.6 | 1.71E+07 |
| 20.34873 | 20.43288 | 20.3849 | 4 | 4 | 4 | 1.6 | 1.71E+07 |
| 13.03996 | 14.20069 | 11.01913 | 2 | 2 | 2 | 3.7 | 1.70E+07 |
| 14.40615 | 19.37515 | 12.08784 | 3 | 3 | 3 | 5.2 | 1.70E+07 |
| 13.24869 | 13.88072 | 12.38762 | 2 | 2 | 2 | 3 | 1.69E+07 |
| 18.42333 | 20.51335 | 17.61712 | 2 | 2 | 2 | 9.7 | 1.69E+07 |
| 14.02347 | 13.60119 | 10.2143 | 4 | 4 | 4 | 8.6 | 1.69E+07 |
| 10.20912 | 13.08833 | 12.1354 | 2 | 2 | 2 | 4.7 | 1.69E+07 |
| 12.47696 | 19.48526 | 12.45649 | 2 | 2 | 2 | 3.5 | 1.68E+07 |
| 11.62239 | 12.71353 | 11.87567 | 3 | 3 | 3 | 3.5 | 1.68E+07 |
| 11.1396 | 20.30251 | 17.95146 | 3 | 3 | 3 | 6.4 | 1.67E+07 |
| 13.63677 | 13.24683 | 11.97644 | 4 | 4 | 4 | 1.2 | 1.66E+07 |
| 12.12956 | 20.59441 | 19.64959 | 2 | 2 | 2 | 25.5 | 1.66E+07 |
| 11.67074 | 13.20056 | 11.00783 | 4 | 4 | 4 | 19.1 | 1.66E+07 |
| 12.96805 | 12.86153 | 17.81547 | 2 | 2 | 2 | 8.2 | 1.64E+07 |
| 11.10882 | 12.7475 | 12.37985 | 2 | 2 | 2 | 7.9 | 1.64E+07 |
| 11.1444 | 13.49567 | 10.76119 | 3 | 3 | 3 | 5.9 | 1.63E+07 |

| | | | | | | | |
|---|---|---|---|---|---|---|---|
| 11.20322 | 14.27894 | 12.04944 | 13 | 3 | 3 | 11.4 | 1.61E+07 |
| 12.04655 | 13.48205 | 10.91236 | 2 | 2 | 2 | 12.1 | 1.61E+07 |
| 13.55953 | 13.82223 | 12.19101 | 2 | 2 | 2 | 6.6 | 1.61E+07 |
| 12.40844 | 13.95817 | 12.17019 | 2 | 2 | 2 | 10.8 | 1.61E+07 |
| 11.26325 | 13.96722 | 11.94854 | 3 | 2 | 2 | 21.2 | 1.60E+07 |
| 11.7823 | 19.97729 | 11.7041 | 2 | 2 | 2 | 7.9 | 1.60E+07 |
| 12.31841 | 13.20621 | 11.50714 | 2 | 2 | 2 | 5.9 | 1.60E+07 |
| 12.84494 | 13.9278 | 11.54213 | 2 | 2 | 2 | 5.1 | 1.60E+07 |
| 11.88289 | 20.23257 | 11.73912 | 2 | 2 | 2 | 5.3 | 1.59E+07 |
| 11.16435 | 14.25874 | 13.55266 | 2 | 2 | 2 | 3.1 | 1.58E+07 |
| 10.25044 | 14.98467 | 13.87838 | 2 | 2 | 2 | 1.8 | 1.58E+07 |
| 12.89334 | 13.7384 | 10.82534 | 2 | 2 | 2 | 8.9 | 1.58E+07 |
| 11.85212 | 12.43594 | 10.91838 | 2 | 2 | 2 | 5.3 | 1.58E+07 |
| 10.71811 | 13.50931 | 10.33558 | 2 | 2 | 2 | 4.4 | 1.57E+07 |
| 12.91486 | 14.06197 | 11.89858 | 2 | 2 | 2 | 12.7 | 1.56E+07 |
| 12.3946 | 11.17645 | 11.34689 | 2 | 2 | 2 | 4.6 | 1.56E+07 |
| 12.61871 | 13.18753 | 13.60122 | 2 | 2 | 2 | 8.1 | 1.56E+07 |
| 12.00797 | 15.31822 | 12.05632 | 2 | 2 | 2 | 5 | 1.55E+07 |
| 12.36562 | 14.58594 | 11.47621 | 2 | 2 | 2 | 3 | 1.55E+07 |
| 16.96804 | 14.21003 | 11.48469 | 2 | 2 | 2 | 3.8 | 1.55E+07 |
| 13.15175 | 13.20036 | 12.69724 | 2 | 2 | 2 | 3.6 | 1.55E+07 |
| 10.35336 | 12.06549 | 13.00692 | 3 | 3 | 3 | 5.6 | 1.54E+07 |
| 12.03041 | 19.72492 | 12.37382 | 2 | 2 | 2 | 6.2 | 1.54E+07 |
| 10.56535 | 13.48668 | 13.83929 | 2 | 2 | 2 | 7.6 | 1.54E+07 |
| 13.02557 | 13.07334 | 11.70362 | 2 | 2 | 2 | 3 | 1.53E+07 |
| 13.76399 | 15.06366 | 12.96754 | 2 | 2 | 2 | 10.2 | 1.52E+07 |
| 11.14379 | 13.70441 | 9.892752 | 2 | 2 | 2 | 2.2 | 1.51E+07 |
| 12.02079 | 13.97037 | 11.92871 | 2 | 2 | 2 | 12.6 | 1.51E+07 |
| 11.92318 | 19.33353 | 12.57063 | 2 | 2 | 2 | 5.3 | 1.50E+07 |
| 12.03257 | 12.86155 | 11.04364 | 3 | 3 | 3 | 3.8 | 1.50E+07 |
| 11.74449 | 13.40357 | 12.55563 | 2 | 2 | 2 | 8.6 | 1.49E+07 |
| 11.53069 | 21.91652 | 14.63503 | 7 | 7 | 7 | 5.2 | 1.48E+07 |
| 18.63357 | 18.75245 | 13.90471 | 2 | 2 | 2 | 4.9 | 1.48E+07 |
| 12.38579 | 13.91069 | 12.5563 | 2 | 2 | 2 | 28.2 | 1.48E+07 |
| 16.84046 | 12.80582 | 19.27314 | 2 | 2 | 2 | 14.2 | 1.47E+07 |
| 13.00122 | 13.55874 | 10.87159 | 15 | 2 | 2 | 11.8 | 1.47E+07 |
| 10.39525 | 14.7265 | 11.73052 | 3 | 3 | 3 | 2.2 | 1.46E+07 |
| 11.81922 | 12.71521 | 11.1749 | 4 | 3 | 3 | 15.3 | 1.46E+07 |
| 10.78133 | 15.29143 | 11.83805 | 2 | 2 | 2 | 2.2 | 1.45E+07 |
| 17.67418 | 21.38585 | 11.21066 | 2 | 2 | 2 | 9.9 | 1.44E+07 |
| 12.59034 | 14.28378 | 19.04081 | 3 | 3 | 3 | 6.1 | 1.43E+07 |
| 11.91331 | 13.52954 | 12.55119 | 2 | 2 | 2 | 3.4 | 1.43E+07 |
| 12.06442 | 12.91925 | 11.38732 | 2 | 2 | 2 | 3 | 1.43E+07 |
| 11.38112 | 19.25022 | 11.9979 | 2 | 2 | 2 | 1.1 | 1.42E+07 |
| 13.53787 | 13.52814 | 13.22866 | 2 | 2 | 2 | 4 | 1.41E+07 |
| 13.002 | 19.62587 | 10.93152 | 2 | 2 | 2 | 7.9 | 1.41E+07 |
| 10.03504 | 13.01185 | 14.18882 | 2 | 2 | 2 | 5.3 | 1.41E+07 |
| 12.09922 | 14.07734 | 11.6051 | 3 | 3 | 3 | 3.5 | 1.39E+07 |

| | | | | | | | |
|---|---|---|---|---|---|---|---|
| 12.50396 | 13.40686 | 13.71972 | 2 | 2 | 2 | 9.8 | 1.39E+07 |
| 13.00709 | 14.22588 | 11.60733 | 2 | 2 | 2 | 4.2 | 1.38E+07 |
| 13.2202 | 13.20061 | 10.52575 | 2 | 2 | 2 | 8 | 1.38E+07 |
| 11.39293 | 15.16888 | 12.36742 | 2 | 2 | 2 | 13.2 | 1.38E+07 |
| 12.89829 | 19.05635 | 12.70297 | 2 | 2 | 2 | 17.2 | 1.37E+07 |
| 12.29506 | 12.07405 | 11.36696 | 2 | 2 | 2 | 2.1 | 1.37E+07 |
| 11.549 | 12.71503 | 12.02458 | 3 | 3 | 3 | 11.9 | 1.36E+07 |
| 12.31596 | 14.15797 | 12.70585 | 2 | 2 | 2 | 9.7 | 1.36E+07 |
| 11.55533 | 12.4923 | 12.25097 | 2 | 2 | 2 | 4.3 | 1.35E+07 |
| 12.44295 | 12.00489 | 12.83019 | 2 | 2 | 2 | 7.8 | 1.35E+07 |
| 10.70665 | 13.97404 | 12.23477 | 2 | 2 | 2 | 10.3 | 1.34E+07 |
| 12.92284 | 13.69709 | 12.45573 | 2 | 2 | 2 | 4.4 | 1.33E+07 |
| 11.67301 | 13.07354 | 13.54366 | 2 | 2 | 2 | 4.6 | 1.32E+07 |
| 11.9498 | 14.88623 | 11.58795 | 2 | 2 | 2 | 3.5 | 1.32E+07 |
| 11.8762 | 12.10038 | 11.78943 | 2 | 2 | 2 | 6 | 1.31E+07 |
| 13.05976 | 13.47114 | 11.26646 | 2 | 2 | 2 | 12.8 | 1.31E+07 |
| 11.80508 | 12.88787 | 12.30482 | 3 | 3 | 3 | 6.7 | 1.31E+07 |
| 13.77047 | 13.17361 | 12.99085 | 2 | 2 | 2 | 2.9 | 1.30E+07 |
| 12.19845 | 13.24661 | 13.52196 | 2 | 2 | 2 | 6 | 1.29E+07 |
| 11.80069 | 18.46587 | 12.52718 | 2 | 2 | 2 | 12.2 | 1.28E+07 |
| 13.23148 | 22.19874 | 12.10661 | 2 | 2 | 2 | 4.8 | 1.28E+07 |
| 10.92776 | 14.58847 | 12.65186 | 3 | 3 | 3 | 17.1 | 1.27E+07 |
| 14.39629 | 20.61 | 12.5192 | 2 | 2 | 2 | 17.6 | 1.26E+07 |
| 12.84591 | 19.03625 | 12.31167 | 2 | 2 | 2 | 15 | 1.26E+07 |
| 11.6126 | 14.95554 | 11.47055 | 3 | 2 | 2 | 5.5 | 1.26E+07 |
| 12.45553 | 14.12412 | 11.95418 | 3 | 3 | 3 | 0.7 | 1.26E+07 |
| 11.75201 | 21.79891 | 12.87733 | 5 | 5 | 5 | 22.8 | 1.26E+07 |
| 11.346 | 12.18887 | 11.98442 | 2 | 2 | 2 | 11.7 | 1.25E+07 |
| 12.38187 | 12.54308 | 12.10559 | 2 | 2 | 2 | 15 | 1.24E+07 |
| 12.16372 | 13.51683 | 10.90792 | 2 | 2 | 2 | 1.1 | 1.24E+07 |
| 12.84597 | 14.1001 | 11.18903 | 2 | 2 | 2 | 7.3 | 1.23E+07 |
| 11.37197 | 12.5095 | 11.13793 | 3 | 3 | 3 | 13.1 | 1.22E+07 |
| 12.25347 | 14.17779 | 10.51807 | 2 | 2 | 2 | 6.1 | 1.21E+07 |
| 13.27321 | 13.01769 | 10.21989 | 2 | 2 | 2 | 3.6 | 1.20E+07 |
| 12.59341 | 13.14484 | 10.70153 | 3 | 3 | 3 | 3.1 | 1.20E+07 |
| 12.84074 | 12.18617 | 11.90362 | 4 | 4 | 4 | 3.8 | 1.19E+07 |
| 12.72861 | 14.11386 | 12.07409 | 2 | 2 | 2 | 2.9 | 1.19E+07 |
| 11.88794 | 11.92077 | 12.23501 | 2 | 2 | 2 | 5.8 | 1.19E+07 |
| 11.54212 | 13.09998 | 11.4309 | 2 | 2 | 2 | 3.5 | 1.18E+07 |
| 12.08022 | 20.98635 | 11.23423 | 2 | 2 | 2 | 2.1 | 1.18E+07 |
| 12.15733 | 21.58495 | 13.68132 | 2 | 2 | 2 | 22.9 | 1.18E+07 |
| 11.12207 | 18.76937 | 13.03011 | 3 | 3 | 3 | 4.1 | 1.18E+07 |
| 20.01672 | 13.71555 | 12.36786 | 4 | 4 | 4 | 15.1 | 1.15E+07 |
| 12.64334 | 13.79855 | 11.48996 | 2 | 2 | 2 | 6.1 | 1.14E+07 |
| 12.00316 | 13.25904 | 10.45349 | 2 | 2 | 2 | 2.9 | 1.14E+07 |
| 13.00807 | 12.684 | 11.15262 | 2 | 2 | 2 | 3.3 | 1.13E+07 |
| 10.50119 | 12.31642 | 17.7602 | 2 | 2 | 2 | 9 | 1.12E+07 |
| 12.77081 | 12.26181 | 13.06809 | 2 | 2 | 2 | 3.4 | 1.12E+07 |

| | | | | | | | |
|---|---|---|---|---|---|---|---|
| 11.98313 | 14.58681 | 11.50463 | 3 | 3 | 3 | 12.8 | 1.09E+07 |
| 12.3153 | 13.69629 | 11.48184 | 2 | 2 | 2 | 6.3 | 1.08E+07 |
| 13.55727 | 14.04779 | 11.57314 | 3 | 3 | 3 | 3.2 | 1.08E+07 |
| 11.9981 | 14.76544 | 12.21915 | 2 | 2 | 2 | 0.9 | 1.07E+07 |
| 11.49844 | 11.1935 | 12.1516 | 2 | 2 | 2 | 4.7 | 1.06E+07 |
| 12.7091 | 13.98599 | 11.3129 | 2 | 2 | 2 | 24.1 | 1.05E+07 |
| 12.73526 | 14.57149 | 12.76409 | 2 | 2 | 2 | 20.2 | 1.03E+07 |
| 13.048 | 12.99035 | 13.10029 | 2 | 2 | 2 | 7.9 | 1.03E+07 |
| 12.52499 | 15.34592 | 13.59877 | 2 | 2 | 2 | 6.9 | 1.02E+07 |
| 11.04852 | 13.18491 | 11.84112 | 2 | 2 | 2 | 2.3 | 1.01E+07 |
| 10.81338 | 14.97001 | 11.81117 | 2 | 2 | 2 | 2.7 | 1.00E+07 |
| 11.44132 | 13.67647 | 13.83605 | 2 | 2 | 2 | 2.9 | 9.83E+06 |
| 11.27361 | 19.99341 | 12.39585 | 2 | 2 | 2 | 16.2 | 9.75E+06 |
| 10.77545 | 13.75181 | 12.99965 | 2 | 2 | 2 | 2.1 | 9.64E+06 |
| 12.22091 | 14.62873 | 11.5199 | 2 | 2 | 2 | 6.3 | 9.43E+06 |
| 10.64124 | 11.89751 | 11.81377 | 2 | 2 | 2 | 2.2 | 9.31E+06 |
| 13.40929 | 20.78197 | 12.04229 | 3 | 3 | 3 | 7.9 | 9.23E+06 |
| 11.41038 | 12.93292 | 11.57821 | 2 | 2 | 2 | 5.1 | 9.15E+06 |
| 13.27117 | 14.2277 | 13.67353 | 2 | 2 | 2 | 5.8 | 9.13E+06 |
| 12.73234 | 13.95236 | 11.63177 | 2 | 2 | 2 | 4.3 | 8.99E+06 |
| 12.39062 | 13.66758 | 12.24581 | 2 | 2 | 2 | 1.6 | 8.72E+06 |
| 11.06492 | 13.72661 | 10.8663 | 2 | 2 | 2 | 9 | 8.71E+06 |
| 18.18916 | 18.64699 | 18.6768 | 3 | 3 | 3 | 12 | 8.65E+06 |
| 12.17848 | 12.50874 | 12.11139 | 2 | 2 | 2 | 2.7 | 8.58E+06 |
| 11.97171 | 19.3869 | 13.14214 | 3 | 3 | 3 | 5.8 | 8.43E+06 |
| 11.03202 | 15.28664 | 11.78343 | 3 | 3 | 3 | 3.3 | 8.40E+06 |
| 12.3677 | 14.23083 | 11.69245 | 3 | 3 | 3 | 4.4 | 8.36E+06 |
| 12.17298 | 12.34246 | 10.82535 | 3 | 3 | 3 | 12.5 | 8.36E+06 |
| 18.72214 | 13.7717 | 13.18805 | 2 | 2 | 2 | 4.5 | 8.33E+06 |
| 12.11431 | 13.92123 | 18.17767 | 3 | 3 | 3 | 10.3 | 8.21E+06 |
| 12.14709 | 13.12407 | 12.61487 | 3 | 3 | 3 | 5.3 | 8.19E+06 |
| 9.932918 | 19.27182 | 9.878994 | 2 | 2 | 2 | 8.1 | 8.16E+06 |
| 10.96 | 19.52978 | 10.29817 | 2 | 2 | 2 | 8.7 | 8.01E+06 |
| 9.874801 | 13.25861 | 11.08915 | 3 | 3 | 3 | 4.3 | 7.86E+06 |
| 12.93152 | 20.57828 | 13.4872 | 3 | 3 | 3 | 8.5 | 7.83E+06 |
| 12.79698 | 18.60567 | 12.77317 | 2 | 2 | 2 | 1.1 | 7.82E+06 |
| 12.23494 | 12.13302 | 11.27149 | 3 | 3 | 3 | 2.5 | 7.81E+06 |
| 12.32497 | 14.73172 | 12.02394 | 2 | 2 | 2 | 2.4 | 7.77E+06 |
| 13.05909 | 14.93342 | 11.27388 | 2 | 2 | 2 | 7.5 | 7.65E+06 |
| 18.78918 | 21.07215 | 13.29035 | 2 | 2 | 2 | 3.5 | 7.60E+06 |
| 18.55921 | 13.50893 | 17.69513 | 2 | 2 | 2 | 4.4 | 7.58E+06 |
| 12.67194 | 21.18979 | 17.96961 | 2 | 2 | 2 | 7.7 | 7.53E+06 |
| 13.09926 | 13.68916 | 13.37846 | 2 | 2 | 2 | 3.5 | 7.53E+06 |
| 13.0648 | 21.19809 | 12.12719 | 2 | 2 | 2 | 3 | 7.44E+06 |
| 13.68267 | 14.7636 | 13.11791 | 3 | 3 | 3 | 10.7 | 7.31E+06 |
| 12.94066 | 14.77871 | 12.02826 | 2 | 2 | 2 | 4.1 | 7.25E+06 |
| 12.6038 | 13.02823 | 12.33062 | 2 | 2 | 2 | 5.3 | 7.12E+06 |
| 12.01948 | 14.04759 | 11.83824 | 2 | 2 | 2 | 2.2 | 6.97E+06 |

| | | | | | | | |
|---|---|---|---|---|---|---|---|
| 12.18008 | 20.572 | 12.35773 | 2 | 2 | 2 | 11.3 | 6.94E+06 |
| 10.92984 | 19.44845 | 12.49589 | 3 | 3 | 3 | 5.8 | 6.86E+06 |
| 12.60764 | 13.44531 | 12.319 | 4 | 2 | 2 | 9.7 | 6.85E+06 |
| 12.95248 | 14.69826 | 13.46047 | 2 | 2 | 2 | 1.9 | 6.68E+06 |
| 13.17431 | 19.94464 | 11.1803 | 4 | 2 | 2 | 5 | 6.49E+06 |
| 14.34289 | 13.19834 | 13.57135 | 3 | 3 | 3 | 4.9 | 6.45E+06 |
| 10.45535 | 13.38498 | 13.10365 | 2 | 2 | 2 | 3.3 | 6.29E+06 |
| 12.21824 | 13.40353 | 12.70915 | 2 | 2 | 2 | 2.4 | 6.22E+06 |
| 13.49341 | 13.45258 | 11.41917 | 2 | 2 | 2 | 6.1 | 6.17E+06 |
| 12.35752 | 19.61929 | 11.59936 | 2 | 2 | 2 | 4.2 | 6.14E+06 |
| 11.12632 | 18.93822 | 11.57894 | 2 | 2 | 2 | 3.9 | 6.12E+06 |
| 13.48965 | 12.9507 | 12.84997 | 2 | 2 | 1 | 6.7 | 5.83E+06 |
| 11.43466 | 11.78021 | 12.73394 | 2 | 2 | 2 | 3.9 | 5.66E+06 |
| 12.01416 | 19.77444 | 11.35361 | 7 | 2 | 2 | 8.9 | 5.52E+06 |
| 12.28133 | 14.72365 | 12.72633 | 2 | 2 | 2 | 6.6 | 5.49E+06 |
| 12.61226 | 13.93771 | 11.76401 | 2 | 2 | 2 | 4.1 | 5.22E+06 |
| 13.08851 | 12.87496 | 12.09336 | 2 | 2 | 2 | 1.8 | 5.20E+06 |
| 12.45901 | 12.95153 | 11.19258 | 2 | 2 | 2 | 5.7 | 5.12E+06 |
| 11.26971 | 11.52799 | 12.82122 | 2 | 2 | 2 | 3.2 | 5.02E+06 |
| 11.75348 | 13.68629 | 11.97836 | 2 | 2 | 2 | 4.2 | 4.99E+06 |
| 12.06985 | 13.83918 | 11.58715 | 2 | 2 | 2 | 6.1 | 4.87E+06 |
| 13.48549 | 14.75088 | 12.50384 | 2 | 2 | 2 | 11.4 | 4.86E+06 |
| 11.76063 | 19.3449 | 12.62457 | 2 | 2 | 2 | 4.3 | 4.82E+06 |
| 19.71272 | 13.78442 | 10.56139 | 2 | 2 | 2 | 3.1 | 4.78E+06 |
| 11.92612 | 11.93203 | 12.62941 | 2 | 2 | 2 | 12.1 | 4.78E+06 |
| 12.62642 | 19.0827 | 13.37739 | 2 | 2 | 2 | 12.7 | 4.59E+06 |
| 11.98449 | 14.33071 | 12.86306 | 2 | 2 | 2 | 1.3 | 4.58E+06 |
| 13.91945 | 20.1493 | 11.64721 | 2 | 2 | 2 | 12.2 | 4.15E+06 |
| 13.1944 | 18.48133 | 13.02418 | 2 | 2 | 2 | 14.8 | 4.04E+06 |
| 12.99673 | 14.49548 | 11.08531 | 2 | 2 | 2 | 6.6 | 3.94E+06 |
| 11.721 | 14.07581 | 12.22148 | 2 | 2 | 2 | 12.6 | 3.87E+06 |
| 12.51828 | 15.42899 | 11.84592 | 3 | 3 | 3 | 7.3 | 2.43E+06 |

| MS/MS counts VR R1 | MS/MS counts VR R2 | MS/MS counts VR R3 | Log 2 LFQ intensity IgG R1 | MS/MS counts IgG R2 | MS/MS counts IgG R3 | MS/MS count 6 |
|---|---|---|---|---|---|---|
| 1210 | 276 | 303 | 328 | 51 | 141 | 111 |
| 256 | 56 | 56 | 62 | 14 | 37 | 31 |
| 269 | 88 | 92 | 85 | 0 | 4 | 0 |
| 56 | 21 | 20 | 14 | 0 | 1 | 0 |
| 104 | 28 | 30 | 27 | 2 | 14 | 3 |
| 135 | 41 | 46 | 42 | 2 | 4 | 0 |
| 221 | 75 | 74 | 72 | 0 | 0 | 0 |
| 115 | 26 | 31 | 33 | 5 | 14 | 6 |
| 557 | 174 | 197 | 186 | 0 | 0 | 0 |
| 198 | 64 | 66 | 68 | 0 | 0 | 0 |
| 191 | 63 | 57 | 54 | 3 | 12 | 2 |
| 219 | 65 | 70 | 80 | 0 | 4 | 0 |
| 534 | 174 | 176 | 184 | 0 | 0 | 0 |
| 126 | 35 | 41 | 38 | 0 | 10 | 2 |
| 97 | 32 | 32 | 33 | 0 | 0 | 0 |
| 159 | 44 | 48 | 48 | 3 | 13 | 3 |
| 216 | 73 | 72 | 71 | 0 | 0 | 0 |
| 127 | 35 | 33 | 40 | 1 | 16 | 2 |
| 126 | 41 | 37 | 39 | 1 | 5 | 3 |
| 142 | 50 | 44 | 48 | 0 | 0 | 0 |
| 60 | 16 | 21 | 15 | 1 | 6 | 1 |
| 207 | 68 | 66 | 64 | 1 | 6 | 2 |
| 415 | 136 | 140 | 139 | 0 | 0 | 0 |
| 64 | 18 | 20 | 22 | 0 | 4 | 0 |
| 86 | 26 | 27 | 24 | 1 | 7 | 1 |
| 108 | 36 | 34 | 34 | 0 | 3 | 1 |
| 68 | 21 | 23 | 21 | 0 | 3 | 0 |
| 237 | 81 | 77 | 78 | 0 | 1 | 0 |
| 113 | 34 | 34 | 39 | 0 | 6 | 0 |
| 88 | 27 | 32 | 25 | 1 | 2 | 1 |
| 71 | 22 | 21 | 24 | 0 | 3 | 1 |
| 48 | 13 | 16 | 16 | 0 | 2 | 1 |
| 55 | 15 | 18 | 14 | 0 | 5 | 3 |
| 47 | 13 | 15 | 13 | 1 | 5 | 0 |
| 132 | 38 | 43 | 43 | 0 | 8 | 0 |
| 161 | 40 | 41 | 38 | 9 | 23 | 10 |
| 55 | 13 | 12 | 17 | 3 | 5 | 5 |
| 94 | 33 | 28 | 30 | 0 | 2 | 1 |
| 42 | 12 | 13 | 14 | 0 | 2 | 1 |
| 75 | 22 | 25 | 25 | 0 | 2 | 1 |
| 205 | 67 | 69 | 68 | 0 | 1 | 0 |
| 81 | 18 | 20 | 23 | 5 | 11 | 4 |
| 92 | 26 | 31 | 31 | 0 | 3 | 1 |
| 160 | 52 | 53 | 55 | 0 | 0 | 0 |

| | | | | | | |
|---|---|---|---|---|---|---|
| 149 | 46 | 45 | 51 | 0 | 7 | 0 |
| 65 | 21 | 21 | 18 | 0 | 4 | 1 |
| 70 | 16 | 15 | 14 | 5 | 14 | 6 |
| 61 | 19 | 19 | 15 | 1 | 5 | 2 |
| 62 | 21 | 18 | 21 | 0 | 1 | 1 |
| 182 | 51 | 35 | 60 | 0 | 35 | 1 |
| 141 | 48 | 46 | 45 | 0 | 2 | 0 |
| 137 | 44 | 46 | 38 | 0 | 9 | 0 |
| 68 | 21 | 21 | 17 | 1 | 6 | 2 |
| 165 | 51 | 58 | 56 | 0 | 0 | 0 |
| 20 | 6 | 6 | 6 | 0 | 2 | 0 |
| 114 | 39 | 32 | 35 | 0 | 8 | 0 |
| 67 | 16 | 21 | 17 | 2 | 8 | 3 |
| 189 | 62 | 61 | 62 | 0 | 4 | 0 |
| 72 | 26 | 23 | 23 | 0 | 0 | 0 |
| 71 | 28 | 18 | 24 | 0 | 1 | 0 |
| 91 | 26 | 29 | 31 | 0 | 5 | 0 |
| 36 | 9 | 12 | 10 | 1 | 2 | 2 |
| 83 | 24 | 22 | 22 | 2 | 7 | 6 |
| 156 | 49 | 45 | 51 | 1 | 9 | 1 |
| 289 | 90 | 100 | 99 | 0 | 0 | 0 |
| 127 | 36 | 47 | 44 | 0 | 0 | 0 |
| 46 | 14 | 15 | 15 | 0 | 2 | 0 |
| 51 | 16 | 16 | 16 | 0 | 2 | 1 |
| 71 | 21 | 24 | 21 | 1 | 3 | 1 |
| 180 | 55 | 61 | 63 | 0 | 1 | 0 |
| 82 | 29 | 24 | 23 | 1 | 4 | 1 |
| 59 | 18 | 17 | 16 | 2 | 5 | 1 |
| 30 | 5 | 7 | 7 | 2 | 7 | 2 |
| 39 | 15 | 11 | 12 | 0 | 1 | 0 |
| 85 | 28 | 30 | 27 | 0 | 0 | 0 |
| 96 | 28 | 26 | 30 | 2 | 10 | 0 |
| 76 | 25 | 27 | 24 | 0 | 0 | 0 |
| 140 | 37 | 41 | 37 | 6 | 15 | 4 |
| 66 | 21 | 24 | 21 | 0 | 0 | 0 |
| 41 | 14 | 12 | 14 | 0 | 1 | 0 |
| 159 | 55 | 54 | 50 | 0 | 0 | 0 |
| 344 | 120 | 110 | 114 | 0 | 0 | 0 |
| 38 | 15 | 12 | 11 | 0 | 0 | 0 |
| 268 | 93 | 85 | 90 | 0 | 0 | 0 |
| 46 | 13 | 13 | 12 | 1 | 5 | 2 |
| 31 | 9 | 10 | 9 | 1 | 2 | 0 |
| 60 | 20 | 21 | 19 | 0 | 0 | 0 |
| 77 | 23 | 21 | 22 | 1 | 7 | 3 |
| 153 | 57 | 48 | 48 | 0 | 0 | 0 |
| 101 | 35 | 35 | 31 | 0 | 0 | 0 |
| 131 | 45 | 43 | 43 | 0 | 0 | 0 |
| 46 | 15 | 15 | 15 | 0 | 1 | 0 |

| | | | | | | |
|---|---|---|---|---|---|---|
| 45 | 11 | 18 | 13 | 1 | 1 | 1 |
| 103 | 34 | 33 | 36 | 0 | 0 | 0 |
| 125 | 39 | 34 | 34 | 0 | 15 | 3 |
| 40 | 13 | 13 | 14 | 0 | 0 | 0 |
| 101 | 35 | 33 | 33 | 0 | 0 | 0 |
| 38 | 12 | 10 | 13 | 0 | 3 | 0 |
| 74 | 26 | 23 | 25 | 0 | 0 | 0 |
| 111 | 40 | 36 | 34 | 0 | 1 | 0 |
| 319 | 111 | 100 | 108 | 0 | 0 | 0 |
| 55 | 19 | 14 | 17 | 0 | 4 | 1 |
| 33 | 11 | 8 | 11 | 0 | 2 | 1 |
| 21 | 8 | 6 | 5 | 0 | 2 | 0 |
| 140 | 45 | 42 | 48 | 0 | 4 | 1 |
| 32 | 9 | 10 | 10 | 0 | 2 | 1 |
| 28 | 7 | 8 | 10 | 0 | 2 | 1 |
| 109 | 39 | 36 | 34 | 0 | 0 | 0 |
| 229 | 77 | 73 | 75 | 0 | 3 | 1 |
| 61 | 20 | 19 | 22 | 0 | 0 | 0 |
| 259 | 71 | 80 | 87 | 0 | 11 | 10 |
| 68 | 22 | 21 | 25 | 0 | 0 | 0 |
| 24 | 6 | 5 | 9 | 0 | 2 | 2 |
| 137 | 31 | 31 | 38 | 5 | 26 | 6 |
| 66 | 15 | 13 | 13 | 5 | 17 | 3 |
| 100 | 30 | 31 | 34 | 1 | 2 | 2 |
| 57 | 19 | 18 | 20 | 0 | 0 | 0 |
| 148 | 44 | 36 | 38 | 7 | 15 | 8 |
| 36 | 11 | 13 | 11 | 0 | 1 | 0 |
| 96 | 27 | 26 | 30 | 0 | 11 | 2 |
| 52 | 13 | 21 | 15 | 0 | 3 | 0 |
| 65 | 26 | 18 | 20 | 0 | 1 | 0 |
| 90 | 30 | 28 | 32 | 0 | 0 | 0 |
| 92 | 31 | 29 | 32 | 0 | 0 | 0 |
| 92 | 32 | 33 | 27 | 0 | 0 | 0 |
| 29 | 8 | 11 | 10 | 0 | 0 | 0 |
| 52 | 15 | 17 | 16 | 0 | 3 | 1 |
| 115 | 37 | 27 | 31 | 2 | 12 | 6 |
| 251 | 83 | 85 | 83 | 0 | 0 | 0 |
| 53 | 17 | 16 | 19 | 0 | 1 | 0 |
| 19 | 6 | 5 | 6 | 0 | 2 | 0 |
| 140 | 44 | 49 | 47 | 0 | 0 | 0 |
| 92 | 29 | 30 | 33 | 0 | 0 | 0 |
| 131 | 40 | 46 | 42 | 0 | 2 | 1 |
| 190 | 51 | 52 | 64 | 1 | 17 | 5 |
| 39 | 9 | 15 | 12 | 0 | 3 | 0 |
| 139 | 44 | 48 | 47 | 0 | 0 | 0 |
| 101 | 32 | 35 | 34 | 0 | 0 | 0 |
| 74 | 25 | 23 | 26 | 0 | 0 | 0 |
| 151 | 53 | 51 | 47 | 0 | 0 | 0 |

| | | | | | | |
|---|---|---|---|---|---|---|
| 84 | 30 | 24 | 23 | 0 | 7 | 0 |
| 99 | 32 | 29 | 37 | 0 | 1 | 0 |
| 128 | 45 | 42 | 41 | 0 | 0 | 0 |
| 77 | 23 | 27 | 27 | 0 | 0 | 0 |
| 56 | 19 | 18 | 17 | 0 | 2 | 0 |
| 24 | 9 | 7 | 5 | 1 | 2 | 0 |
| 10 | 3 | 4 | 3 | 0 | 0 | 0 |
| 47 | 16 | 15 | 16 | 0 | 0 | 0 |
| 80 | 28 | 25 | 27 | 0 | 0 | 0 |
| 20 | 6 | 7 | 6 | 0 | 1 | 0 |
| 20 | 7 | 6 | 6 | 0 | 1 | 0 |
| 107 | 29 | 22 | 31 | 1 | 23 | 1 |
| 60 | 21 | 20 | 19 | 0 | 0 | 0 |
| 27 | 7 | 9 | 10 | 0 | 1 | 0 |
| 111 | 42 | 32 | 36 | 0 | 1 | 0 |
| 52 | 16 | 20 | 16 | 0 | 0 | 0 |
| 95 | 33 | 31 | 31 | 0 | 0 | 0 |
| 81 | 24 | 28 | 26 | 0 | 3 | 0 |
| 49 | 16 | 17 | 16 | 0 | 0 | 0 |
| 115 | 40 | 36 | 39 | 0 | 0 | 0 |
| 162 | 54 | 50 | 58 | 0 | 0 | 0 |
| 108 | 35 | 38 | 35 | 0 | 0 | 0 |
| 97 | 29 | 34 | 34 | 0 | 0 | 0 |
| 38 | 15 | 12 | 10 | 0 | 1 | 0 |
| 108 | 26 | 28 | 34 | 2 | 11 | 7 |
| 27 | 9 | 8 | 9 | 0 | 1 | 0 |
| 24 | 8 | 9 | 7 | 0 | 0 | 0 |
| 79 | 25 | 27 | 27 | 0 | 0 | 0 |
| 65 | 22 | 15 | 20 | 0 | 5 | 3 |
| 29 | 10 | 8 | 8 | 1 | 2 | 0 |
| 50 | 14 | 16 | 20 | 0 | 0 | 0 |
| 92 | 29 | 30 | 33 | 0 | 0 | 0 |
| 56 | 17 | 18 | 21 | 0 | 0 | 0 |
| 116 | 40 | 39 | 37 | 0 | 0 | 0 |
| 21 | 6 | 7 | 7 | 0 | 1 | 0 |
| 105 | 36 | 36 | 32 | 0 | 1 | 0 |
| 136 | 45 | 43 | 48 | 0 | 0 | 0 |
| 66 | 23 | 21 | 20 | 0 | 2 | 0 |
| 52 | 15 | 15 | 14 | 2 | 6 | 0 |
| 107 | 35 | 34 | 38 | 0 | 0 | 0 |
| 147 | 47 | 49 | 51 | 0 | 0 | 0 |
| 47 | 15 | 13 | 13 | 1 | 4 | 1 |
| 132 | 49 | 43 | 40 | 0 | 0 | 0 |
| 53 | 14 | 16 | 17 | 0 | 5 | 1 |
| 103 | 33 | 32 | 38 | 0 | 0 | 0 |
| 79 | 27 | 26 | 26 | 0 | 0 | 0 |
| 71 | 19 | 27 | 23 | 0 | 2 | 0 |
| 58 | 19 | 19 | 20 | 0 | 0 | 0 |

| | | | | | | |
|---|---|---|---|---|---|---|
| 46 | 12 | 11 | 12 | 3 | 8 | 0 |
| 87 | 29 | 29 | 29 | 0 | 0 | 0 |
| 57 | 20 | 17 | 17 | 0 | 2 | 1 |
| 115 | 39 | 41 | 35 | 0 | 0 | 0 |
| 10 | 4 | 3 | 3 | 0 | 0 | 0 |
| 48 | 15 | 17 | 16 | 0 | 0 | 0 |
| 51 | 18 | 17 | 16 | 0 | 0 | 0 |
| 56 | 17 | 14 | 13 | 1 | 11 | 0 |
| 139 | 46 | 42 | 46 | 1 | 3 | 1 |
| 47 | 17 | 17 | 13 | 0 | 0 | 0 |
| 80 | 25 | 26 | 29 | 0 | 0 | 0 |
| 179 | 56 | 63 | 60 | 0 | 0 | 0 |
| 76 | 28 | 22 | 21 | 2 | 3 | 0 |
| 78 | 24 | 30 | 24 | 0 | 0 | 0 |
| 69 | 21 | 26 | 20 | 0 | 2 | 0 |
| 82 | 26 | 29 | 27 | 0 | 0 | 0 |
| 28 | 8 | 8 | 8 | 0 | 3 | 1 |
| 141 | 44 | 49 | 48 | 0 | 0 | 0 |
| 15 | 4 | 4 | 6 | 0 | 1 | 0 |
| 102 | 35 | 31 | 32 | 1 | 3 | 0 |
| 4 | 1 | 1 | 2 | 0 | 0 | 0 |
| 75 | 27 | 24 | 23 | 0 | 1 | 0 |
| 103 | 35 | 36 | 32 | 0 | 0 | 0 |
| 131 | 43 | 38 | 44 | 0 | 6 | 0 |
| 174 | 58 | 57 | 59 | 0 | 0 | 0 |
| 65 | 22 | 21 | 20 | 0 | 2 | 0 |
| 73 | 20 | 23 | 30 | 0 | 0 | 0 |
| 57 | 20 | 17 | 20 | 0 | 0 | 0 |
| 54 | 16 | 17 | 20 | 0 | 1 | 0 |
| 55 | 17 | 19 | 19 | 0 | 0 | 0 |
| 40 | 14 | 9 | 12 | 0 | 5 | 0 |
| 30 | 10 | 9 | 11 | 0 | 0 | 0 |
| 60 | 17 | 20 | 23 | 0 | 0 | 0 |
| 57 | 19 | 21 | 16 | 0 | 1 | 0 |
| 47 | 17 | 16 | 14 | 0 | 0 | 0 |
| 96 | 27 | 32 | 35 | 0 | 1 | 1 |
| 77 | 20 | 22 | 28 | 2 | 3 | 2 |
| 152 | 51 | 50 | 51 | 0 | 0 | 0 |
| 99 | 20 | 21 | 24 | 8 | 16 | 10 |
| 99 | 32 | 34 | 33 | 0 | 0 | 0 |
| 21 | 7 | 7 | 7 | 0 | 0 | 0 |
| 39 | 13 | 14 | 12 | 0 | 0 | 0 |
| 124 | 39 | 47 | 38 | 0 | 0 | 0 |
| 72 | 21 | 21 | 24 | 0 | 3 | 3 |
| 139 | 45 | 51 | 43 | 0 | 0 | 0 |
| 137 | 45 | 45 | 47 | 0 | 0 | 0 |
| 40 | 7 | 8 | 7 | 6 | 7 | 5 |
| 105 | 33 | 37 | 35 | 0 | 0 | 0 |

| | | | | | | |
|---|---|---|---|---|---|---|
| 23 | 7 | 9 | 7 | 0 | 0 | 0 |
| 69 | 21 | 23 | 25 | 0 | 0 | 0 |
| 140 | 24 | 17 | 34 | 12 | 36 | 17 |
| 34 | 9 | 10 | 9 | 1 | 4 | 1 |
| 33 | 9 | 11 | 13 | 0 | 0 | 0 |
| 89 | 21 | 19 | 24 | 3 | 12 | 10 |
| 67 | 21 | 23 | 23 | 0 | 0 | 0 |
| 64 | 19 | 10 | 12 | 0 | 21 | 2 |
| 21 | 4 | 6 | 7 | 1 | 3 | 0 |
| 59 | 15 | 16 | 19 | 3 | 6 | 0 |
| 57 | 18 | 20 | 19 | 0 | 0 | 0 |
| 67 | 14 | 14 | 16 | 6 | 9 | 8 |
| 126 | 41 | 41 | 40 | 0 | 4 | 0 |
| 91 | 29 | 34 | 28 | 0 | 0 | 0 |
| 90 | 26 | 28 | 26 | 0 | 10 | 0 |
| 63 | 21 | 18 | 24 | 0 | 0 | 0 |
| 30 | 11 | 8 | 10 | 0 | 1 | 0 |
| 102 | 34 | 31 | 37 | 0 | 0 | 0 |
| 35 | 10 | 11 | 8 | 0 | 4 | 2 |
| 53 | 17 | 21 | 15 | 0 | 0 | 0 |
| 45 | 15 | 12 | 16 | 0 | 2 | 0 |
| 112 | 37 | 39 | 36 | 0 | 0 | 0 |
| 109 | 42 | 30 | 37 | 0 | 0 | 0 |
| 45 | 11 | 9 | 9 | 5 | 6 | 5 |
| 43 | 14 | 17 | 12 | 0 | 0 | 0 |
| 106 | 28 | 36 | 37 | 0 | 5 | 0 |
| 65 | 19 | 21 | 20 | 0 | 5 | 0 |
| 20 | 5 | 7 | 7 | 0 | 1 | 0 |
| 105 | 38 | 30 | 37 | 0 | 0 | 0 |
| 52 | 12 | 10 | 15 | 3 | 11 | 1 |
| 107 | 26 | 29 | 43 | 0 | 7 | 2 |
| 112 | 37 | 34 | 41 | 0 | 0 | 0 |
| 112 | 36 | 34 | 42 | 0 | 0 | 0 |
| 96 | 33 | 31 | 32 | 0 | 0 | 0 |
| 60 | 17 | 21 | 20 | 0 | 2 | 0 |
| 23 | 6 | 7 | 7 | 1 | 2 | 0 |
| 67 | 20 | 25 | 22 | 0 | 0 | 0 |
| 80 | 24 | 26 | 22 | 1 | 7 | 0 |
| 76 | 25 | 24 | 26 | 0 | 1 | 0 |
| 21 | 6 | 8 | 6 | 0 | 1 | 0 |
| 62 | 20 | 23 | 19 | 0 | 0 | 0 |
| 51 | 16 | 18 | 17 | 0 | 0 | 0 |
| 36 | 12 | 13 | 11 | 0 | 0 | 0 |
| 89 | 30 | 28 | 31 | 0 | 0 | 0 |
| 14 | 3 | 5 | 4 | 1 | 1 | 0 |
| 91 | 34 | 26 | 31 | 0 | 0 | 0 |
| 129 | 46 | 11 | 13 | 0 | 59 | 0 |
| 37 | 14 | 11 | 12 | 0 | 0 | 0 |

| | | | | | | |
|---|---|---|---|---|---|---|
| 30 | 10 | 9 | 11 | 0 | 0 | 0 |
| 96 | 31 | 35 | 30 | 0 | 0 | 0 |
| 43 | 13 | 16 | 14 | 0 | 0 | 0 |
| 11 | 3 | 3 | 3 | 0 | 1 | 1 |
| 33 | 10 | 9 | 8 | 0 | 5 | 1 |
| 63 | 20 | 21 | 22 | 0 | 0 | 0 |
| 33 | 11 | 11 | 11 | 0 | 0 | 0 |
| 36 | 10 | 11 | 15 | 0 | 0 | 0 |
| 44 | 13 | 14 | 16 | 0 | 1 | 0 |
| 95 | 27 | 30 | 38 | 0 | 0 | 0 |
| 26 | 7 | 9 | 9 | 0 | 1 | 0 |
| 32 | 10 | 11 | 11 | 0 | 0 | 0 |
| 74 | 26 | 23 | 25 | 0 | 0 | 0 |
| 66 | 20 | 24 | 21 | 0 | 1 | 0 |
| 38 | 13 | 12 | 13 | 0 | 0 | 0 |
| 64 | 22 | 21 | 21 | 0 | 0 | 0 |
| 83 | 28 | 26 | 29 | 0 | 0 | 0 |
| 69 | 22 | 17 | 19 | 0 | 11 | 0 |
| 25 | 9 | 7 | 7 | 0 | 2 | 0 |
| 106 | 37 | 32 | 37 | 0 | 0 | 0 |
| 35 | 11 | 14 | 10 | 0 | 0 | 0 |
| 60 | 19 | 17 | 24 | 0 | 0 | 0 |
| 74 | 22 | 27 | 25 | 0 | 0 | 0 |
| 75 | 28 | 23 | 24 | 0 | 0 | 0 |
| 86 | 27 | 30 | 29 | 0 | 0 | 0 |
| 12 | 3 | 3 | 3 | 1 | 1 | 1 |
| 51 | 19 | 18 | 14 | 0 | 0 | 0 |
| 51 | 13 | 18 | 14 | 2 | 4 | 0 |
| 77 | 21 | 27 | 29 | 0 | 0 | 0 |
| 41 | 14 | 13 | 12 | 0 | 2 | 0 |
| 17 | 7 | 5 | 5 | 0 | 0 | 0 |
| 41 | 14 | 12 | 15 | 0 | 0 | 0 |
| 43 | 13 | 13 | 12 | 0 | 5 | 0 |
| 49 | 14 | 18 | 17 | 0 | 0 | 0 |
| 37 | 11 | 13 | 13 | 0 | 0 | 0 |
| 70 | 23 | 26 | 21 | 0 | 0 | 0 |
| 51 | 15 | 18 | 18 | 0 | 0 | 0 |
| 43 | 12 | 14 | 9 | 1 | 6 | 1 |
| 14 | 5 | 4 | 5 | 0 | 0 | 0 |
| 58 | 20 | 18 | 20 | 0 | 0 | 0 |
| 51 | 14 | 12 | 13 | 1 | 8 | 3 |
| 72 | 19 | 15 | 17 | 0 | 21 | 0 |
| 51 | 17 | 16 | 14 | 0 | 4 | 0 |
| 26 | 8 | 9 | 9 | 0 | 0 | 0 |
| 30 | 10 | 9 | 11 | 0 | 0 | 0 |
| 59 | 21 | 21 | 17 | 0 | 0 | 0 |
| 29 | 7 | 10 | 7 | 1 | 4 | 0 |
| 38 | 14 | 13 | 11 | 0 | 0 | 0 |

| | | | | | | |
|---|---|---|---|---|---|---|
| 33 | 10 | 9 | 11 | 0 | 3 | 0 |
| 40 | 10 | 10 | 13 | 0 | 7 | 0 |
| 67 | 23 | 22 | 22 | 0 | 0 | 0 |
| 80 | 26 | 27 | 27 | 0 | 0 | 0 |
| 60 | 19 | 20 | 21 | 0 | 0 | 0 |
| 84 | 24 | 23 | 31 | 0 | 3 | 3 |
| 33 | 10 | 12 | 11 | 0 | 0 | 0 |
| 11 | 3 | 3 | 2 | 1 | 2 | 0 |
| 29 | 9 | 10 | 10 | 0 | 0 | 0 |
| 32 | 10 | 10 | 10 | 0 | 2 | 0 |
| 20 | 7 | 5 | 8 | 0 | 0 | 0 |
| 65 | 23 | 24 | 18 | 0 | 0 | 0 |
| 82 | 26 | 26 | 30 | 0 | 0 | 0 |
| 70 | 26 | 23 | 20 | 0 | 1 | 0 |
| 40 | 12 | 15 | 13 | 0 | 0 | 0 |
| 39 | 14 | 9 | 10 | 0 | 6 | 0 |
| 71 | 25 | 23 | 23 | 0 | 0 | 0 |
| 38 | 14 | 11 | 13 | 0 | 0 | 0 |
| 43 | 12 | 14 | 17 | 0 | 0 | 0 |
| 64 | 17 | 21 | 21 | 0 | 3 | 2 |
| 34 | 10 | 12 | 12 | 0 | 0 | 0 |
| 34 | 11 | 12 | 11 | 0 | 0 | 0 |
| 42 | 15 | 12 | 15 | 0 | 0 | 0 |
| 32 | 10 | 10 | 11 | 0 | 1 | 0 |
| 34 | 10 | 13 | 11 | 0 | 0 | 0 |
| 61 | 19 | 24 | 18 | 0 | 0 | 0 |
| 41 | 9 | 11 | 13 | 2 | 3 | 3 |
| 74 | 27 | 21 | 26 | 0 | 0 | 0 |
| 33 | 11 | 12 | 10 | 0 | 0 | 0 |
| 80 | 29 | 24 | 27 | 0 | 0 | 0 |
| 44 | 13 | 16 | 15 | 0 | 0 | 0 |
| 70 | 25 | 25 | 20 | 0 | 0 | 0 |
| 43 | 15 | 15 | 13 | 0 | 0 | 0 |
| 44 | 13 | 14 | 17 | 0 | 0 | 0 |
| 36 | 12 | 12 | 12 | 0 | 0 | 0 |
| 44 | 11 | 11 | 12 | 1 | 9 | 0 |
| 52 | 16 | 18 | 18 | 0 | 0 | 0 |
| 44 | 14 | 15 | 15 | 0 | 0 | 0 |
| 49 | 19 | 17 | 13 | 0 | 0 | 0 |
| 47 | 15 | 14 | 18 | 0 | 0 | 0 |
| 55 | 19 | 19 | 17 | 0 | 0 | 0 |
| 41 | 13 | 13 | 15 | 0 | 0 | 0 |
| 29 | 10 | 9 | 10 | 0 | 0 | 0 |
| 26 | 8 | 9 | 9 | 0 | 0 | 0 |
| 50 | 17 | 17 | 16 | 0 | 0 | 0 |
| 73 | 22 | 27 | 24 | 0 | 0 | 0 |
| 54 | 20 | 16 | 18 | 0 | 0 | 0 |
| 7 | 4 | 2 | 1 | 0 | 0 | 0 |

| | | | | | | |
|---|---|---|---|---|---|---|
| 16 | 5 | 5 | 6 | 0 | 0 | 0 |
| 14 | 5 | 1 | 2 | 0 | 5 | 1 |
| 59 | 20 | 20 | 19 | 0 | 0 | 0 |
| 61 | 22 | 20 | 19 | 0 | 0 | 0 |
| 35 | 13 | 12 | 10 | 0 | 0 | 0 |
| 53 | 17 | 14 | 15 | 0 | 6 | 1 |
| 29 | 4 | 4 | 5 | 3 | 8 | 5 |
| 43 | 13 | 17 | 13 | 0 | 0 | 0 |
| 50 | 18 | 16 | 16 | 0 | 0 | 0 |
| 65 | 20 | 21 | 24 | 0 | 0 | 0 |
| 61 | 18 | 21 | 22 | 0 | 0 | 0 |
| 59 | 19 | 17 | 23 | 0 | 0 | 0 |
| 53 | 17 | 16 | 19 | 0 | 1 | 0 |
| 45 | 15 | 13 | 16 | 0 | 1 | 0 |
| 54 | 16 | 19 | 19 | 0 | 0 | 0 |
| 65 | 20 | 23 | 22 | 0 | 0 | 0 |
| 33 | 11 | 11 | 10 | 0 | 1 | 0 |
| 36 | 10 | 14 | 12 | 0 | 0 | 0 |
| 35 | 14 | 10 | 11 | 0 | 0 | 0 |
| 13 | 4 | 4 | 5 | 0 | 0 | 0 |
| 45 | 14 | 14 | 17 | 0 | 0 | 0 |
| 50 | 17 | 16 | 17 | 0 | 0 | 0 |
| 33 | 11 | 10 | 12 | 0 | 0 | 0 |
| 40 | 14 | 13 | 13 | 0 | 0 | 0 |
| 81 | 27 | 28 | 26 | 0 | 0 | 0 |
| 47 | 16 | 17 | 14 | 0 | 0 | 0 |
| 45 | 15 | 11 | 8 | 0 | 11 | 0 |
| 30 | 11 | 11 | 8 | 0 | 0 | 0 |
| 66 | 22 | 22 | 22 | 0 | 0 | 0 |
| 47 | 17 | 13 | 17 | 0 | 0 | 0 |
| 66 | 22 | 23 | 21 | 0 | 0 | 0 |
| 39 | 11 | 12 | 12 | 2 | 2 | 0 |
| 31 | 11 | 10 | 10 | 0 | 0 | 0 |
| 36 | 12 | 13 | 11 | 0 | 0 | 0 |
| 17 | 6 | 0 | 0 | 0 | 10 | 1 |
| 18 | 6 | 5 | 7 | 0 | 0 | 0 |
| 73 | 26 | 23 | 24 | 0 | 0 | 0 |
| 48 | 16 | 15 | 17 | 0 | 0 | 0 |
| 53 | 17 | 17 | 19 | 0 | 0 | 0 |
| 53 | 16 | 18 | 19 | 0 | 0 | 0 |
| 38 | 13 | 15 | 10 | 0 | 0 | 0 |
| 54 | 21 | 16 | 17 | 0 | 0 | 0 |
| 66 | 24 | 21 | 21 | 0 | 0 | 0 |
| 62 | 21 | 21 | 20 | 0 | 0 | 0 |
| 61 | 24 | 18 | 19 | 0 | 0 | 0 |
| 33 | 11 | 11 | 11 | 0 | 0 | 0 |
| 37 | 13 | 11 | 13 | 0 | 0 | 0 |
| 37 | 13 | 13 | 11 | 0 | 0 | 0 |

| | | | | | | |
|---|---|---|---|---|---|---|
| 28 | 9 | 8 | 11 | 0 | 0 | 0 |
| 55 | 18 | 15 | 22 | 0 | 0 | 0 |
| 25 | 9 | 8 | 8 | 0 | 0 | 0 |
| 51 | 10 | 11 | 10 | 6 | 8 | 6 |
| 62 | 22 | 22 | 18 | 0 | 0 | 0 |
| 32 | 10 | 13 | 9 | 0 | 0 | 0 |
| 22 | 8 | 7 | 7 | 0 | 0 | 0 |
| 73 | 22 | 24 | 27 | 0 | 0 | 0 |
| 53 | 18 | 16 | 19 | 0 | 0 | 0 |
| 48 | 17 | 17 | 14 | 0 | 0 | 0 |
| 36 | 12 | 13 | 11 | 0 | 0 | 0 |
| 53 | 19 | 17 | 16 | 0 | 1 | 0 |
| 47 | 15 | 17 | 15 | 0 | 0 | 0 |
| 38 | 11 | 14 | 13 | 0 | 0 | 0 |
| 74 | 22 | 24 | 28 | 0 | 0 | 0 |
| 47 | 16 | 16 | 15 | 0 | 0 | 0 |
| 31 | 10 | 12 | 9 | 0 | 0 | 0 |
| 49 | 15 | 14 | 20 | 0 | 0 | 0 |
| 34 | 9 | 12 | 13 | 0 | 0 | 0 |
| 64 | 21 | 22 | 21 | 0 | 0 | 0 |
| 37 | 15 | 9 | 13 | 0 | 0 | 0 |
| 52 | 18 | 16 | 18 | 0 | 0 | 0 |
| 16 | 5 | 5 | 6 | 0 | 0 | 0 |
| 16 | 4 | 4 | 4 | 1 | 1 | 2 |
| 40 | 11 | 14 | 15 | 0 | 0 | 0 |
| 16 | 6 | 5 | 4 | 1 | 0 | 0 |
| 18 | 8 | 0 | 0 | 0 | 10 | 0 |
| 34 | 11 | 11 | 12 | 0 | 0 | 0 |
| 72 | 23 | 25 | 24 | 0 | 0 | 0 |
| 25 | 8 | 9 | 8 | 0 | 0 | 0 |
| 16 | 5 | 6 | 5 | 0 | 0 | 0 |
| 29 | 10 | 10 | 9 | 0 | 0 | 0 |
| 15 | 5 | 6 | 4 | 0 | 0 | 0 |
| 43 | 5 | 7 | 7 | 8 | 8 | 8 |
| 53 | 19 | 15 | 18 | 0 | 0 | 1 |
| 39 | 12 | 12 | 15 | 0 | 0 | 0 |
| 50 | 16 | 17 | 17 | 0 | 0 | 0 |
| 42 | 15 | 12 | 15 | 0 | 0 | 0 |
| 34 | 12 | 13 | 9 | 0 | 0 | 0 |
| 44 | 14 | 15 | 14 | 0 | 1 | 0 |
| 27 | 9 | 11 | 7 | 0 | 0 | 0 |
| 19 | 6 | 6 | 6 | 0 | 1 | 0 |
| 51 | 20 | 17 | 14 | 0 | 0 | 0 |
| 10 | 3 | 2 | 3 | 1 | 1 | 0 |
| 27 | 10 | 6 | 6 | 0 | 4 | 1 |
| 37 | 14 | 12 | 11 | 0 | 0 | 0 |
| 39 | 15 | 13 | 11 | 0 | 0 | 0 |
| 51 | 18 | 17 | 16 | 0 | 0 | 0 |

| | | | | | | |
|---|---|---|---|---|---|---|
| 58 | 20 | 18 | 20 | 0 | 0 | 0 |
| 35 | 13 | 11 | 11 | 0 | 0 | 0 |
| 59 | 19 | 18 | 22 | 0 | 0 | 0 |
| 55 | 17 | 19 | 19 | 0 | 0 | 0 |
| 49 | 16 | 14 | 19 | 0 | 0 | 0 |
| 20 | 4 | 6 | 5 | 1 | 1 | 3 |
| 24 | 10 | 9 | 5 | 0 | 0 | 0 |
| 40 | 15 | 12 | 13 | 0 | 0 | 0 |
| 24 | 8 | 8 | 8 | 0 | 0 | 0 |
| 27 | 7 | 10 | 10 | 0 | 0 | 0 |
| 40 | 14 | 12 | 14 | 0 | 0 | 0 |
| 38 | 12 | 9 | 17 | 0 | 0 | 0 |
| 44 | 11 | 13 | 17 | 0 | 3 | 0 |
| 17 | 5 | 5 | 5 | 0 | 2 | 0 |
| 31 | 9 | 11 | 11 | 0 | 0 | 0 |
| 25 | 7 | 10 | 8 | 0 | 0 | 0 |
| 30 | 12 | 11 | 7 | 0 | 0 | 0 |
| 42 | 14 | 13 | 15 | 0 | 0 | 0 |
| 55 | 22 | 18 | 15 | 0 | 0 | 0 |
| 57 | 13 | 15 | 13 | 2 | 10 | 4 |
| 29 | 7 | 12 | 7 | 0 | 3 | 0 |
| 33 | 10 | 10 | 11 | 0 | 2 | 0 |
| 29 | 10 | 8 | 11 | 0 | 0 | 0 |
| 27 | 11 | 8 | 8 | 0 | 0 | 0 |
| 25 | 9 | 8 | 8 | 0 | 0 | 0 |
| 19 | 6 | 7 | 6 | 0 | 0 | 0 |
| 45 | 16 | 14 | 15 | 0 | 0 | 0 |
| 22 | 7 | 5 | 10 | 0 | 0 | 0 |
| 49 | 16 | 16 | 17 | 0 | 0 | 0 |
| 16 | 4 | 4 | 4 | 1 | 2 | 1 |
| 34 | 12 | 12 | 10 | 0 | 0 | 0 |
| 48 | 17 | 16 | 15 | 0 | 0 | 0 |
| 12 | 4 | 4 | 4 | 0 | 0 | 0 |
| 20 | 7 | 8 | 5 | 0 | 0 | 0 |
| 30 | 9 | 10 | 11 | 0 | 0 | 0 |
| 42 | 13 | 15 | 11 | 1 | 2 | 0 |
| 46 | 15 | 16 | 15 | 0 | 0 | 0 |
| 38 | 16 | 10 | 12 | 0 | 0 | 0 |
| 23 | 7 | 8 | 8 | 0 | 0 | 0 |
| 26 | 8 | 9 | 9 | 0 | 0 | 0 |
| 63 | 21 | 18 | 24 | 0 | 0 | 0 |
| 22 | 8 | 6 | 8 | 0 | 0 | 0 |
| 40 | 13 | 13 | 14 | 0 | 0 | 0 |
| 62 | 19 | 17 | 26 | 0 | 0 | 0 |
| 56 | 22 | 16 | 18 | 0 | 0 | 0 |
| 46 | 14 | 17 | 15 | 0 | 0 | 0 |
| 42 | 16 | 13 | 13 | 0 | 0 | 0 |
| 49 | 19 | 14 | 16 | 0 | 0 | 0 |

| | | | | | | |
|---|---|---|---|---|---|---|
| 40 | 13 | 13 | 14 | 0 | 0 | 0 |
| 28 | 7 | 8 | 9 | 0 | 4 | 0 |
| 48 | 16 | 16 | 16 | 0 | 0 | 0 |
| 54 | 15 | 16 | 23 | 0 | 0 | 0 |
| 41 | 12 | 15 | 14 | 0 | 0 | 0 |
| 22 | 8 | 7 | 7 | 0 | 0 | 0 |
| 29 | 9 | 10 | 10 | 0 | 0 | 0 |
| 30 | 9 | 12 | 9 | 0 | 0 | 0 |
| 40 | 12 | 13 | 15 | 0 | 0 | 0 |
| 35 | 10 | 13 | 12 | 0 | 0 | 0 |
| 26 | 11 | 6 | 8 | 0 | 1 | 0 |
| 52 | 18 | 17 | 17 | 0 | 0 | 0 |
| 25 | 8 | 8 | 9 | 0 | 0 | 0 |
| 26 | 10 | 8 | 8 | 0 | 0 | 0 |
| 37 | 6 | 8 | 6 | 4 | 9 | 4 |
| 51 | 15 | 17 | 19 | 0 | 0 | 0 |
| 52 | 15 | 18 | 19 | 0 | 0 | 0 |
| 33 | 11 | 10 | 12 | 0 | 0 | 0 |
| 25 | 6 | 6 | 6 | 1 | 5 | 1 |
| 57 | 21 | 19 | 17 | 0 | 0 | 0 |
| 14 | 5 | 4 | 4 | 0 | 1 | 0 |
| 34 | 12 | 11 | 11 | 0 | 0 | 0 |
| 27 | 10 | 9 | 8 | 0 | 0 | 0 |
| 25 | 9 | 9 | 7 | 0 | 0 | 0 |
| 53 | 20 | 16 | 17 | 0 | 0 | 0 |
| 25 | 7 | 7 | 8 | 0 | 2 | 1 |
| 7 | 3 | 2 | 2 | 0 | 0 | 0 |
| 41 | 13 | 15 | 13 | 0 | 0 | 0 |
| 23 | 6 | 9 | 8 | 0 | 0 | 0 |
| 23 | 7 | 7 | 9 | 0 | 0 | 0 |
| 28 | 11 | 9 | 8 | 0 | 0 | 0 |
| 34 | 11 | 13 | 10 | 0 | 0 | 0 |
| 32 | 9 | 12 | 10 | 0 | 1 | 0 |
| 18 | 3 | 4 | 2 | 1 | 5 | 3 |
| 39 | 13 | 13 | 13 | 0 | 0 | 0 |
| 35 | 12 | 13 | 10 | 0 | 0 | 0 |
| 9 | 4 | 0 | 0 | 0 | 5 | 0 |
| 34 | 8 | 13 | 13 | 0 | 0 | 0 |
| 31 | 12 | 11 | 8 | 0 | 0 | 0 |
| 28 | 9 | 10 | 9 | 0 | 0 | 0 |
| 28 | 9 | 11 | 8 | 0 | 0 | 0 |
| 50 | 19 | 16 | 15 | 0 | 0 | 0 |
| 25 | 10 | 6 | 8 | 0 | 1 | 0 |
| 40 | 9 | 16 | 15 | 0 | 0 | 0 |
| 23 | 8 | 7 | 8 | 0 | 0 | 0 |
| 38 | 11 | 13 | 14 | 0 | 0 | 0 |
| 24 | 8 | 8 | 8 | 0 | 0 | 0 |
| 37 | 11 | 11 | 15 | 0 | 0 | 0 |

| | | | | | | |
|---|---|---|---|---|---|---|
| 38 | 12 | 11 | 15 | 0 | 0 | 0 |
| 33 | 8 | 13 | 12 | 0 | 0 | 0 |
| 38 | 13 | 12 | 13 | 0 | 0 | 0 |
| 28 | 9 | 11 | 7 | 0 | 1 | 0 |
| 30 | 10 | 11 | 9 | 0 | 0 | 0 |
| 36 | 15 | 12 | 9 | 0 | 0 | 0 |
| 42 | 14 | 15 | 13 | 0 | 0 | 0 |
| 23 | 8 | 7 | 8 | 0 | 0 | 0 |
| 21 | 7 | 7 | 7 | 0 | 0 | 0 |
| 42 | 15 | 15 | 12 | 0 | 0 | 0 |
| 33 | 10 | 8 | 14 | 0 | 1 | 0 |
| 43 | 15 | 13 | 15 | 0 | 0 | 0 |
| 41 | 13 | 11 | 17 | 0 | 0 | 0 |
| 32 | 11 | 11 | 10 | 0 | 0 | 0 |
| 13 | 2 | 4 | 4 | 1 | 0 | 2 |
| 18 | 2 | 5 | 4 | 1 | 3 | 3 |
| 38 | 13 | 12 | 13 | 0 | 0 | 0 |
| 24 | 6 | 8 | 7 | 1 | 1 | 1 |
| 24 | 8 | 8 | 7 | 0 | 1 | 0 |
| 38 | 14 | 13 | 11 | 0 | 0 | 0 |
| 18 | 5 | 7 | 6 | 0 | 0 | 0 |
| 44 | 17 | 13 | 14 | 0 | 0 | 0 |
| 9 | 2 | 3 | 3 | 0 | 1 | 0 |
| 26 | 8 | 8 | 10 | 0 | 0 | 0 |
| 18 | 5 | 5 | 8 | 0 | 0 | 0 |
| 40 | 13 | 16 | 11 | 0 | 0 | 0 |
| 13 | 5 | 3 | 5 | 0 | 0 | 0 |
| 41 | 15 | 10 | 16 | 0 | 0 | 0 |
| 29 | 8 | 10 | 10 | 0 | 1 | 0 |
| 52 | 15 | 16 | 21 | 0 | 0 | 0 |
| 49 | 16 | 14 | 19 | 0 | 0 | 0 |
| 20 | 5 | 5 | 5 | 1 | 4 | 0 |
| 34 | 10 | 10 | 14 | 0 | 0 | 0 |
| 9 | 2 | 2 | 3 | 1 | 1 | 0 |
| 39 | 11 | 13 | 14 | 0 | 1 | 0 |
| 27 | 8 | 9 | 10 | 0 | 0 | 0 |
| 34 | 13 | 10 | 11 | 0 | 0 | 0 |
| 41 | 11 | 18 | 12 | 0 | 0 | 0 |
| 45 | 13 | 14 | 18 | 0 | 0 | 0 |
| 29 | 10 | 10 | 8 | 0 | 1 | 0 |
| 27 | 9 | 8 | 10 | 0 | 0 | 0 |
| 25 | 9 | 9 | 7 | 0 | 0 | 0 |
| 29 | 9 | 9 | 11 | 0 | 0 | 0 |
| 18 | 5 | 6 | 7 | 0 | 0 | 0 |
| 4 | 1 | 1 | 2 | 0 | 0 | 0 |
| 26 | 8 | 8 | 10 | 0 | 0 | 0 |
| 18 | 6 | 6 | 5 | 0 | 1 | 0 |
| 21 | 7 | 7 | 7 | 0 | 0 | 0 |

| | | | | | | |
|---|---|---|---|---|---|---|
| 38 | 13 | 10 | 15 | 0 | 0 | 0 |
| 42 | 14 | 15 | 13 | 0 | 0 | 0 |
| 12 | 4 | 4 | 4 | 0 | 0 | 0 |
| 30 | 4 | 3 | 7 | 2 | 8 | 6 |
| 35 | 11 | 13 | 11 | 0 | 0 | 0 |
| 40 | 13 | 14 | 13 | 0 | 0 | 0 |
| 18 | 5 | 7 | 6 | 0 | 0 | 0 |
| 37 | 12 | 15 | 10 | 0 | 0 | 0 |
| 34 | 10 | 14 | 10 | 0 | 0 | 0 |
| 25 | 9 | 8 | 8 | 0 | 0 | 0 |
| 30 | 9 | 7 | 8 | 1 | 3 | 2 |
| 50 | 14 | 19 | 17 | 0 | 0 | 0 |
| 39 | 10 | 17 | 12 | 0 | 0 | 0 |
| 25 | 7 | 9 | 9 | 0 | 0 | 0 |
| 21 | 8 | 6 | 7 | 0 | 0 | 0 |
| 26 | 7 | 10 | 9 | 0 | 0 | 0 |
| 25 | 11 | 6 | 8 | 0 | 0 | 0 |
| 26 | 8 | 10 | 8 | 0 | 0 | 0 |
| 10 | 4 | 4 | 2 | 0 | 0 | 0 |
| 29 | 9 | 9 | 11 | 0 | 0 | 0 |
| 8 | 3 | 3 | 2 | 0 | 0 | 0 |
| 15 | 5 | 5 | 5 | 0 | 0 | 0 |
| 33 | 11 | 11 | 11 | 0 | 0 | 0 |
| 27 | 10 | 9 | 8 | 0 | 0 | 0 |
| 15 | 4 | 6 | 4 | 0 | 1 | 0 |
| 43 | 16 | 15 | 10 | 0 | 2 | 0 |
| 21 | 8 | 7 | 6 | 0 | 0 | 0 |
| 32 | 9 | 11 | 12 | 0 | 0 | 0 |
| 29 | 12 | 8 | 9 | 0 | 0 | 0 |
| 21 | 6 | 6 | 9 | 0 | 0 | 0 |
| 36 | 13 | 12 | 11 | 0 | 0 | 0 |
| 12 | 5 | 3 | 4 | 0 | 0 | 0 |
| 26 | 8 | 9 | 9 | 0 | 0 | 0 |
| 28 | 5 | 11 | 12 | 0 | 0 | 0 |
| 15 | 4 | 3 | 4 | 0 | 4 | 0 |
| 15 | 6 | 4 | 5 | 0 | 0 | 0 |
| 31 | 10 | 11 | 10 | 0 | 0 | 0 |
| 16 | 6 | 5 | 5 | 0 | 0 | 0 |
| 15 | 4 | 5 | 5 | 0 | 1 | 0 |
| 38 | 16 | 9 | 13 | 0 | 0 | 0 |
| 16 | 7 | 3 | 6 | 0 | 0 | 0 |
| 16 | 7 | 4 | 5 | 0 | 0 | 0 |
| 37 | 11 | 12 | 14 | 0 | 0 | 0 |
| 35 | 11 | 11 | 13 | 0 | 0 | 0 |
| 34 | 12 | 12 | 10 | 0 | 0 | 0 |
| 21 | 7 | 7 | 7 | 0 | 0 | 0 |
| 22 | 7 | 8 | 7 | 0 | 0 | 0 |
| 36 | 12 | 12 | 12 | 0 | 0 | 0 |

| | | | | | | |
|---|---|---|---|---|---|---|
| 21 | 7 | 7 | 5 | 0 | 2 | 0 |
| 26 | 9 | 8 | 9 | 0 | 0 | 0 |
| 31 | 11 | 10 | 10 | 0 | 0 | 0 |
| 14 | 4 | 4 | 6 | 0 | 0 | 0 |
| 25 | 8 | 8 | 9 | 0 | 0 | 0 |
| 18 | 6 | 5 | 7 | 0 | 0 | 0 |
| 22 | 8 | 6 | 8 | 0 | 0 | 0 |
| 10 | 3 | 3 | 4 | 0 | 0 | 0 |
| 31 | 8 | 11 | 10 | 0 | 2 | 0 |
| 19 | 7 | 6 | 6 | 0 | 0 | 0 |
| 24 | 9 | 9 | 6 | 0 | 0 | 0 |
| 17 | 5 | 6 | 6 | 0 | 0 | 0 |
| 29 | 10 | 9 | 10 | 0 | 0 | 0 |
| 21 | 8 | 7 | 6 | 0 | 0 | 0 |
| 20 | 8 | 6 | 6 | 0 | 0 | 0 |
| 32 | 10 | 11 | 11 | 0 | 0 | 0 |
| 16 | 5 | 5 | 6 | 0 | 0 | 0 |
| 36 | 11 | 11 | 14 | 0 | 0 | 0 |
| 30 | 12 | 8 | 10 | 0 | 0 | 0 |
| 20 | 7 | 5 | 8 | 0 | 0 | 0 |
| 16 | 4 | 6 | 6 | 0 | 0 | 0 |
| 30 | 10 | 11 | 9 | 0 | 0 | 0 |
| 37 | 13 | 11 | 13 | 0 | 0 | 0 |
| 22 | 7 | 7 | 8 | 0 | 0 | 0 |
| 47 | 17 | 14 | 16 | 0 | 0 | 0 |
| 21 | 5 | 5 | 7 | 0 | 4 | 0 |
| 26 | 7 | 9 | 9 | 0 | 1 | 0 |
| 23 | 8 | 7 | 8 | 0 | 0 | 0 |
| 21 | 7 | 5 | 9 | 0 | 0 | 0 |
| 19 | 5 | 6 | 4 | 0 | 3 | 1 |
| 31 | 10 | 11 | 10 | 0 | 0 | 0 |
| 8 | 2 | 4 | 2 | 0 | 0 | 0 |
| 21 | 7 | 7 | 7 | 0 | 0 | 0 |
| 33 | 13 | 9 | 11 | 0 | 0 | 0 |
| 33 | 11 | 11 | 11 | 0 | 0 | 0 |
| 20 | 8 | 6 | 6 | 0 | 0 | 0 |
| 31 | 12 | 11 | 8 | 0 | 0 | 0 |
| 28 | 9 | 7 | 12 | 0 | 0 | 0 |
| 14 | 4 | 5 | 5 | 0 | 0 | 0 |
| 13 | 5 | 5 | 3 | 0 | 0 | 0 |
| 47 | 13 | 19 | 15 | 0 | 0 | 0 |
| 17 | 8 | 5 | 4 | 0 | 0 | 0 |
| 29 | 8 | 12 | 9 | 0 | 0 | 0 |
| 31 | 10 | 11 | 10 | 0 | 0 | 0 |
| 17 | 6 | 5 | 6 | 0 | 0 | 0 |
| 15 | 5 | 4 | 6 | 0 | 0 | 0 |
| 32 | 12 | 9 | 10 | 0 | 1 | 0 |
| 31 | 14 | 9 | 8 | 0 | 0 | 0 |

| | | | | | | |
|---|---|---|---|---|---|---|
| 28 | 11 | 7 | 10 | 0 | 0 | 0 |
| 25 | 9 | 7 | 9 | 0 | 0 | 0 |
| 6 | 2 | 2 | 2 | 0 | 0 | 0 |
| 6 | 2 | 2 | 2 | 0 | 0 | 0 |
| 33 | 11 | 10 | 12 | 0 | 0 | 0 |
| 27 | 10 | 8 | 9 | 0 | 0 | 0 |
| 24 | 7 | 8 | 9 | 0 | 0 | 0 |
| 24 | 8 | 9 | 7 | 0 | 0 | 0 |
| 29 | 8 | 12 | 9 | 0 | 0 | 0 |
| 20 | 5 | 7 | 8 | 0 | 0 | 0 |
| 19 | 6 | 5 | 8 | 0 | 0 | 0 |
| 39 | 12 | 11 | 14 | 0 | 2 | 0 |
| 26 | 6 | 10 | 9 | 0 | 1 | 0 |
| 26 | 9 | 8 | 9 | 0 | 0 | 0 |
| 26 | 7 | 9 | 10 | 0 | 0 | 0 |
| 17 | 5 | 6 | 6 | 0 | 0 | 0 |
| 13 | 3 | 5 | 5 | 0 | 0 | 0 |
| 21 | 5 | 9 | 7 | 0 | 0 | 0 |
| 10 | 3 | 3 | 3 | 0 | 1 | 0 |
| 35 | 15 | 10 | 10 | 0 | 0 | 0 |
| 34 | 6 | 14 | 14 | 0 | 0 | 0 |
| 22 | 6 | 8 | 8 | 0 | 0 | 0 |
| 18 | 7 | 5 | 6 | 0 | 0 | 0 |
| 9 | 1 | 2 | 2 | 0 | 4 | 0 |
| 24 | 8 | 8 | 8 | 0 | 0 | 0 |
| 17 | 9 | 3 | 1 | 0 | 4 | 0 |
| 9 | 4 | 3 | 2 | 0 | 0 | 0 |
| 25 | 9 | 9 | 7 | 0 | 0 | 0 |
| 22 | 10 | 4 | 8 | 0 | 0 | 0 |
| 20 | 8 | 6 | 6 | 0 | 0 | 0 |
| 26 | 8 | 8 | 10 | 0 | 0 | 0 |
| 19 | 9 | 6 | 4 | 0 | 0 | 0 |
| 27 | 12 | 7 | 8 | 0 | 0 | 0 |
| 29 | 10 | 9 | 10 | 0 | 0 | 0 |
| 21 | 5 | 9 | 7 | 0 | 0 | 0 |
| 15 | 4 | 6 | 5 | 0 | 0 | 0 |
| 33 | 10 | 10 | 13 | 0 | 0 | 0 |
| 26 | 10 | 6 | 10 | 0 | 0 | 0 |
| 17 | 5 | 8 | 4 | 0 | 0 | 0 |
| 19 | 6 | 6 | 7 | 0 | 0 | 0 |
| 25 | 8 | 9 | 8 | 0 | 0 | 0 |
| 13 | 1 | 0 | 0 | 0 | 12 | 0 |
| 2 | 1 | 0 | 0 | 0 | 1 | 0 |
| 24 | 7 | 9 | 8 | 0 | 0 | 0 |
| 15 | 6 | 5 | 4 | 0 | 0 | 0 |
| 28 | 8 | 8 | 12 | 0 | 0 | 0 |
| 20 | 8 | 6 | 6 | 0 | 0 | 0 |
| 25 | 9 | 8 | 8 | 0 | 0 | 0 |

| | | | | | | |
|---|---|---|---|---|---|---|
| 16 | 5 | 5 | 6 | 0 | 0 | 0 |
| 26 | 8 | 9 | 9 | 0 | 0 | 0 |
| 16 | 6 | 5 | 5 | 0 | 0 | 0 |
| 28 | 9 | 8 | 11 | 0 | 0 | 0 |
| 21 | 9 | 5 | 7 | 0 | 0 | 0 |
| 17 | 8 | 5 | 4 | 0 | 0 | 0 |
| 22 | 6 | 7 | 9 | 0 | 0 | 0 |
| 36 | 10 | 16 | 10 | 0 | 0 | 0 |
| 33 | 11 | 10 | 12 | 0 | 0 | 0 |
| 26 | 6 | 4 | 5 | 1 | 6 | 4 |
| 3 | 0 | 1 | 2 | 0 | 0 | 0 |
| 8 | 3 | 2 | 3 | 0 | 0 | 0 |
| 25 | 9 | 7 | 9 | 0 | 0 | 0 |
| 27 | 9 | 10 | 8 | 0 | 0 | 0 |
| 9 | 3 | 3 | 3 | 0 | 0 | 0 |
| 26 | 9 | 8 | 9 | 0 | 0 | 0 |
| 14 | 2 | 0 | 0 | 0 | 12 | 0 |
| 39 | 15 | 13 | 11 | 0 | 0 | 0 |
| 16 | 5 | 4 | 7 | 0 | 0 | 0 |
| 16 | 7 | 4 | 5 | 0 | 0 | 0 |
| 30 | 9 | 10 | 11 | 0 | 0 | 0 |
| 33 | 11 | 11 | 11 | 0 | 0 | 0 |
| 19 | 5 | 7 | 7 | 0 | 0 | 0 |
| 11 | 4 | 3 | 4 | 0 | 0 | 0 |
| 29 | 13 | 7 | 8 | 0 | 1 | 0 |
| 23 | 9 | 5 | 9 | 0 | 0 | 0 |
| 35 | 11 | 11 | 13 | 0 | 0 | 0 |
| 23 | 9 | 7 | 7 | 0 | 0 | 0 |
| 34 | 9 | 12 | 13 | 0 | 0 | 0 |
| 23 | 6 | 9 | 8 | 0 | 0 | 0 |
| 33 | 8 | 11 | 14 | 0 | 0 | 0 |
| 20 | 7 | 7 | 6 | 0 | 0 | 0 |
| 14 | 5 | 3 | 6 | 0 | 0 | 0 |
| 10 | 4 | 2 | 4 | 0 | 0 | 0 |
| 17 | 6 | 5 | 6 | 0 | 0 | 0 |
| 25 | 8 | 7 | 10 | 0 | 0 | 0 |
| 19 | 6 | 8 | 5 | 0 | 0 | 0 |
| 30 | 9 | 9 | 12 | 0 | 0 | 0 |
| 25 | 6 | 8 | 11 | 0 | 0 | 0 |
| 17 | 6 | 5 | 6 | 0 | 0 | 0 |
| 22 | 7 | 8 | 7 | 0 | 0 | 0 |
| 14 | 6 | 4 | 4 | 0 | 0 | 0 |
| 35 | 14 | 8 | 13 | 0 | 0 | 0 |
| 33 | 9 | 13 | 11 | 0 | 0 | 0 |
| 17 | 6 | 4 | 6 | 0 | 1 | 0 |
| 23 | 6 | 9 | 8 | 0 | 0 | 0 |
| 18 | 6 | 3 | 9 | 0 | 0 | 0 |
| 22 | 6 | 9 | 5 | 0 | 2 | 0 |

| | | | | | | |
|---|---|---|---|---|---|---|
| 30 | 8 | 8 | 14 | 0 | 0 | 0 |
| 13 | 5 | 2 | 5 | 0 | 1 | 0 |
| 11 | 4 | 3 | 4 | 0 | 0 | 0 |
| 19 | 5 | 8 | 6 | 0 | 0 | 0 |
| 21 | 8 | 2 | 3 | 0 | 8 | 0 |
| 14 | 5 | 3 | 6 | 0 | 0 | 0 |
| 29 | 8 | 13 | 8 | 0 | 0 | 0 |
| 21 | 7 | 8 | 6 | 0 | 0 | 0 |
| 26 | 8 | 10 | 8 | 0 | 0 | 0 |
| 22 | 8 | 5 | 9 | 0 | 0 | 0 |
| 15 | 5 | 5 | 5 | 0 | 0 | 0 |
| 17 | 6 | 3 | 8 | 0 | 0 | 0 |
| 27 | 7 | 11 | 9 | 0 | 0 | 0 |
| 27 | 9 | 10 | 8 | 0 | 0 | 0 |
| 17 | 6 | 6 | 5 | 0 | 0 | 0 |
| 23 | 8 | 8 | 7 | 0 | 0 | 0 |
| 18 | 7 | 5 | 6 | 0 | 0 | 0 |
| 23 | 9 | 8 | 6 | 0 | 0 | 0 |
| 14 | 5 | 5 | 4 | 0 | 0 | 0 |
| 25 | 11 | 9 | 5 | 0 | 0 | 0 |
| 12 | 5 | 4 | 3 | 0 | 0 | 0 |
| 11 | 4 | 3 | 4 | 0 | 0 | 0 |
| 12 | 5 | 4 | 3 | 0 | 0 | 0 |
| 39 | 11 | 15 | 13 | 0 | 0 | 0 |
| 12 | 5 | 3 | 4 | 0 | 0 | 0 |
| 17 | 7 | 4 | 6 | 0 | 0 | 0 |
| 21 | 6 | 7 | 8 | 0 | 0 | 0 |
| 24 | 6 | 8 | 10 | 0 | 0 | 0 |
| 18 | 8 | 5 | 5 | 0 | 0 | 0 |
| 20 | 7 | 6 | 7 | 0 | 0 | 0 |
| 10 | 3 | 3 | 4 | 0 | 0 | 0 |
| 19 | 6 | 6 | 7 | 0 | 0 | 0 |
| 19 | 6 | 6 | 5 | 0 | 2 | 0 |
| 4 | 1 | 2 | 1 | 0 | 0 | 0 |
| 9 | 3 | 4 | 2 | 0 | 0 | 0 |
| 21 | 6 | 9 | 6 | 0 | 0 | 0 |
| 19 | 9 | 5 | 5 | 0 | 0 | 0 |
| 19 | 6 | 7 | 6 | 0 | 0 | 0 |
| 20 | 6 | 6 | 8 | 0 | 0 | 0 |
| 12 | 5 | 3 | 4 | 0 | 0 | 0 |
| 20 | 7 | 6 | 7 | 0 | 0 | 0 |
| 23 | 6 | 9 | 8 | 0 | 0 | 0 |
| 11 | 4 | 4 | 3 | 0 | 0 | 0 |
| 8 | 3 | 2 | 3 | 0 | 0 | 0 |
| 27 | 8 | 10 | 9 | 0 | 0 | 0 |
| 21 | 8 | 6 | 7 | 0 | 0 | 0 |
| 10 | 3 | 3 | 4 | 0 | 0 | 0 |
| 32 | 11 | 11 | 10 | 0 | 0 | 0 |

| | | | | | | |
|---|---|---|---|---|---|---|
| 23 | 8 | 8 | 7 | 0 | 0 | 0 |
| 22 | 7 | 7 | 8 | 0 | 0 | 0 |
| 7 | 2 | 3 | 2 | 0 | 0 | 0 |
| 14 | 3 | 6 | 4 | 0 | 1 | 0 |
| 24 | 9 | 6 | 9 | 0 | 0 | 0 |
| 19 | 4 | 8 | 7 | 0 | 0 | 0 |
| 14 | 4 | 6 | 4 | 0 | 0 | 0 |
| 14 | 5 | 4 | 5 | 0 | 0 | 0 |
| 14 | 5 | 5 | 4 | 0 | 0 | 0 |
| 7 | 2 | 0 | 0 | 0 | 5 | 0 |
| 18 | 7 | 5 | 6 | 0 | 0 | 0 |
| 10 | 3 | 4 | 3 | 0 | 0 | 0 |
| 15 | 4 | 4 | 7 | 0 | 0 | 0 |
| 26 | 8 | 8 | 10 | 0 | 0 | 0 |
| 17 | 6 | 5 | 6 | 0 | 0 | 0 |
| 18 | 7 | 5 | 4 | 0 | 2 | 0 |
| 13 | 4 | 4 | 5 | 0 | 0 | 0 |
| 24 | 8 | 9 | 7 | 0 | 0 | 0 |
| 13 | 5 | 4 | 4 | 0 | 0 | 0 |
| 19 | 6 | 7 | 6 | 0 | 0 | 0 |
| 16 | 3 | 6 | 7 | 0 | 0 | 0 |
| 13 | 3 | 3 | 7 | 0 | 0 | 0 |
| 20 | 10 | 5 | 5 | 0 | 0 | 0 |
| 16 | 5 | 7 | 4 | 0 | 0 | 0 |
| 31 | 12 | 9 | 10 | 0 | 0 | 0 |
| 19 | 7 | 5 | 7 | 0 | 0 | 0 |
| 14 | 6 | 4 | 4 | 0 | 0 | 0 |
| 21 | 8 | 5 | 8 | 0 | 0 | 0 |
| 28 | 8 | 9 | 11 | 0 | 0 | 0 |
| 23 | 8 | 8 | 7 | 0 | 0 | 0 |
| 11 | 3 | 4 | 4 | 0 | 0 | 0 |
| 10 | 3 | 4 | 3 | 0 | 0 | 0 |
| 16 | 4 | 6 | 5 | 0 | 1 | 0 |
| 23 | 9 | 5 | 6 | 0 | 3 | 0 |
| 13 | 5 | 4 | 4 | 0 | 0 | 0 |
| 25 | 9 | 9 | 7 | 0 | 0 | 0 |
| 10 | 3 | 2 | 4 | 0 | 1 | 0 |
| 28 | 10 | 8 | 10 | 0 | 0 | 0 |
| 26 | 9 | 10 | 7 | 0 | 0 | 0 |
| 19 | 5 | 7 | 7 | 0 | 0 | 0 |
| 19 | 6 | 6 | 7 | 0 | 0 | 0 |
| 25 | 9 | 3 | 6 | 0 | 7 | 0 |
| 19 | 5 | 7 | 7 | 0 | 0 | 0 |
| 10 | 3 | 4 | 3 | 0 | 0 | 0 |
| 16 | 5 | 6 | 5 | 0 | 0 | 0 |
| 24 | 7 | 8 | 9 | 0 | 0 | 0 |
| 15 | 7 | 5 | 3 | 0 | 0 | 0 |
| 20 | 5 | 7 | 8 | 0 | 0 | 0 |

| | | | | | | |
|---|---|---|---|---|---|---|
| 10 | 4 | 3 | 3 | 0 | 0 | 0 |
| 23 | 7 | 7 | 9 | 0 | 0 | 0 |
| 15 | 4 | 5 | 6 | 0 | 0 | 0 |
| 13 | 6 | 3 | 4 | 0 | 0 | 0 |
| 20 | 4 | 8 | 8 | 0 | 0 | 0 |
| 16 | 6 | 4 | 6 | 0 | 0 | 0 |
| 14 | 5 | 4 | 5 | 0 | 0 | 0 |
| 13 | 2 | 5 | 4 | 1 | 1 | 0 |
| 24 | 8 | 8 | 8 | 0 | 0 | 0 |
| 6 | 1 | 3 | 2 | 0 | 0 | 0 |
| 27 | 12 | 7 | 8 | 0 | 0 | 0 |
| 17 | 6 | 6 | 5 | 0 | 0 | 0 |
| 7 | 3 | 2 | 2 | 0 | 0 | 0 |
| 15 | 5 | 4 | 6 | 0 | 0 | 0 |
| 24 | 8 | 7 | 9 | 0 | 0 | 0 |
| 30 | 10 | 10 | 10 | 0 | 0 | 0 |
| 13 | 4 | 5 | 4 | 0 | 0 | 0 |
| 20 | 7 | 6 | 7 | 0 | 0 | 0 |
| 11 | 4 | 4 | 3 | 0 | 0 | 0 |
| 14 | 5 | 4 | 5 | 0 | 0 | 0 |
| 10 | 2 | 5 | 3 | 0 | 0 | 0 |
| 17 | 6 | 5 | 5 | 0 | 1 | 0 |
| 19 | 4 | 6 | 9 | 0 | 0 | 0 |
| 33 | 12 | 11 | 10 | 0 | 0 | 0 |
| 11 | 3 | 3 | 3 | 0 | 1 | 1 |
| 13 | 4 | 2 | 3 | 0 | 4 | 0 |
| 11 | 3 | 3 | 5 | 0 | 0 | 0 |
| 16 | 5 | 5 | 6 | 0 | 0 | 0 |
| 9 | 4 | 1 | 4 | 0 | 0 | 0 |
| 21 | 7 | 6 | 8 | 0 | 0 | 0 |
| 15 | 5 | 5 | 5 | 0 | 0 | 0 |
| 25 | 13 | 2 | 5 | 0 | 5 | 0 |
| 13 | 2 | 6 | 5 | 0 | 0 | 0 |
| 19 | 6 | 7 | 6 | 0 | 0 | 0 |
| 12 | 5 | 4 | 3 | 0 | 0 | 0 |
| 20 | 6 | 6 | 8 | 0 | 0 | 0 |
| 16 | 3 | 7 | 6 | 0 | 0 | 0 |
| 10 | 3 | 2 | 2 | 1 | 1 | 1 |
| 16 | 6 | 4 | 6 | 0 | 0 | 0 |
| 9 | 2 | 4 | 3 | 0 | 0 | 0 |
| 15 | 5 | 5 | 5 | 0 | 0 | 0 |
| 13 | 5 | 3 | 5 | 0 | 0 | 0 |
| 16 | 6 | 5 | 5 | 0 | 0 | 0 |
| 3 | 1 | 1 | 1 | 0 | 0 | 0 |
| 18 | 4 | 7 | 7 | 0 | 0 | 0 |
| 13 | 4 | 5 | 4 | 0 | 0 | 0 |
| 18 | 6 | 6 | 5 | 0 | 1 | 0 |
| 13 | 4 | 4 | 5 | 0 | 0 | 0 |

| | | | | | | |
|---|---|---|---|---|---|---|
| 13 | 7 | 4 | 2 | 0 | 0 | 0 |
| 15 | 2 | 6 | 7 | 0 | 0 | 0 |
| 18 | 6 | 6 | 6 | 0 | 0 | 0 |
| 22 | 7 | 8 | 7 | 0 | 0 | 0 |
| 24 | 9 | 6 | 9 | 0 | 0 | 0 |
| 17 | 4 | 2 | 2 | 0 | 9 | 0 |
| 11 | 4 | 3 | 4 | 0 | 0 | 0 |
| 20 | 8 | 5 | 7 | 0 | 0 | 0 |
| 18 | 6 | 4 | 8 | 0 | 0 | 0 |
| 18 | 5 | 5 | 8 | 0 | 0 | 0 |
| 13 | 3 | 4 | 6 | 0 | 0 | 0 |
| 15 | 5 | 5 | 5 | 0 | 0 | 0 |
| 18 | 6 | 5 | 7 | 0 | 0 | 0 |
| 36 | 13 | 10 | 13 | 0 | 0 | 0 |
| 13 | 4 | 5 | 4 | 0 | 0 | 0 |
| 7 | 3 | 1 | 3 | 0 | 0 | 0 |
| 8 | 3 | 3 | 2 | 0 | 0 | 0 |
| 7 | 2 | 3 | 2 | 0 | 0 | 0 |
| 17 | 5 | 6 | 6 | 0 | 0 | 0 |
| 22 | 8 | 6 | 8 | 0 | 0 | 0 |
| 21 | 6 | 8 | 7 | 0 | 0 | 0 |
| 19 | 8 | 5 | 6 | 0 | 0 | 0 |
| 20 | 7 | 6 | 7 | 0 | 0 | 0 |
| 9 | 3 | 3 | 3 | 0 | 0 | 0 |
| 17 | 8 | 3 | 6 | 0 | 0 | 0 |
| 14 | 4 | 6 | 4 | 0 | 0 | 0 |
| 10 | 3 | 0 | 0 | 0 | 7 | 0 |
| 17 | 5 | 5 | 7 | 0 | 0 | 0 |
| 17 | 5 | 6 | 6 | 0 | 0 | 0 |
| 19 | 7 | 6 | 6 | 0 | 0 | 0 |
| 13 | 4 | 5 | 4 | 0 | 0 | 0 |
| 15 | 5 | 5 | 5 | 0 | 0 | 0 |
| 15 | 5 | 5 | 5 | 0 | 0 | 0 |
| 5 | 2 | 1 | 2 | 0 | 0 | 0 |
| 17 | 6 | 6 | 5 | 0 | 0 | 0 |
| 9 | 3 | 3 | 3 | 0 | 0 | 0 |
| 25 | 6 | 9 | 10 | 0 | 0 | 0 |
| 20 | 7 | 6 | 7 | 0 | 0 | 0 |
| 15 | 6 | 4 | 5 | 0 | 0 | 0 |
| 9 | 2 | 3 | 4 | 0 | 0 | 0 |
| 18 | 4 | 8 | 6 | 0 | 0 | 0 |
| 21 | 7 | 10 | 4 | 0 | 0 | 0 |
| 20 | 8 | 6 | 6 | 0 | 0 | 0 |
| 22 | 10 | 4 | 8 | 0 | 0 | 0 |
| 11 | 4 | 3 | 4 | 0 | 0 | 0 |
| 10 | 2 | 4 | 4 | 0 | 0 | 0 |
| 15 | 5 | 5 | 5 | 0 | 0 | 0 |
| 13 | 5 | 4 | 4 | 0 | 0 | 0 |

| | | | | | | |
|---|---|---|---|---|---|---|
| 15 | 4 | 5 | 6 | 0 | 0 | 0 |
| 20 | 6 | 5 | 9 | 0 | 0 | 0 |
| 18 | 6 | 7 | 5 | 0 | 0 | 0 |
| 14 | 5 | 4 | 5 | 0 | 0 | 0 |
| 18 | 6 | 5 | 7 | 0 | 0 | 0 |
| 19 | 6 | 6 | 7 | 0 | 0 | 0 |
| 23 | 9 | 8 | 6 | 0 | 0 | 0 |
| 14 | 5 | 3 | 6 | 0 | 0 | 0 |
| 24 | 7 | 10 | 7 | 0 | 0 | 0 |
| 5 | 1 | 1 | 1 | 0 | 2 | 0 |
| 19 | 7 | 5 | 7 | 0 | 0 | 0 |
| 23 | 7 | 9 | 7 | 0 | 0 | 0 |
| 14 | 6 | 5 | 3 | 0 | 0 | 0 |
| 5 | 1 | 2 | 2 | 0 | 0 | 0 |
| 14 | 4 | 4 | 6 | 0 | 0 | 0 |
| 17 | 3 | 5 | 5 | 0 | 4 | 0 |
| 18 | 6 | 5 | 7 | 0 | 0 | 0 |
| 11 | 3 | 3 | 5 | 0 | 0 | 0 |
| 19 | 4 | 8 | 7 | 0 | 0 | 0 |
| 12 | 4 | 3 | 5 | 0 | 0 | 0 |
| 6 | 2 | 2 | 2 | 0 | 0 | 0 |
| 16 | 5 | 5 | 6 | 0 | 0 | 0 |
| 15 | 5 | 4 | 6 | 0 | 0 | 0 |
| 6 | 3 | 1 | 2 | 0 | 0 | 0 |
| 13 | 2 | 5 | 6 | 0 | 0 | 0 |
| 23 | 8 | 8 | 7 | 0 | 0 | 0 |
| 21 | 6 | 7 | 8 | 0 | 0 | 0 |
| 23 | 7 | 7 | 9 | 0 | 0 | 0 |
| 8 | 3 | 2 | 3 | 0 | 0 | 0 |
| 13 | 5 | 3 | 5 | 0 | 0 | 0 |
| 6 | 2 | 2 | 2 | 0 | 0 | 0 |
| 11 | 4 | 4 | 3 | 0 | 0 | 0 |
| 10 | 3 | 4 | 3 | 0 | 0 | 0 |
| 17 | 7 | 3 | 1 | 0 | 6 | 0 |
| 11 | 4 | 3 | 4 | 0 | 0 | 0 |
| 18 | 5 | 7 | 6 | 0 | 0 | 0 |
| 15 | 6 | 5 | 4 | 0 | 0 | 0 |
| 12 | 4 | 3 | 4 | 0 | 1 | 0 |
| 22 | 8 | 7 | 7 | 0 | 0 | 0 |
| 17 | 6 | 5 | 6 | 0 | 0 | 0 |
| 17 | 6 | 5 | 6 | 0 | 0 | 0 |
| 11 | 2 | 5 | 4 | 0 | 0 | 0 |
| 13 | 4 | 0 | 0 | 0 | 9 | 0 |
| 8 | 3 | 2 | 3 | 0 | 0 | 0 |
| 17 | 5 | 5 | 7 | 0 | 0 | 0 |
| 12 | 5 | 4 | 3 | 0 | 0 | 0 |
| 10 | 3 | 1 | 1 | 0 | 5 | 0 |
| 13 | 5 | 3 | 5 | 0 | 0 | 0 |

| | | | | | | |
|---|---|---|---|---|---|---|
| 17 | 7 | 7 | 3 | 0 | 0 | 0 |
| 12 | 3 | 4 | 5 | 0 | 0 | 0 |
| 18 | 6 | 6 | 6 | 0 | 0 | 0 |
| 20 | 9 | 5 | 6 | 0 | 0 | 0 |
| 14 | 5 | 5 | 4 | 0 | 0 | 0 |
| 3 | 1 | 2 | 0 | 0 | 0 | 0 |
| 9 | 4 | 2 | 3 | 0 | 0 | 0 |
| 10 | 3 | 4 | 3 | 0 | 0 | 0 |
| 10 | 2 | 3 | 5 | 0 | 0 | 0 |
| 8 | 2 | 2 | 4 | 0 | 0 | 0 |
| 13 | 6 | 3 | 4 | 0 | 0 | 0 |
| 16 | 6 | 4 | 6 | 0 | 0 | 0 |
| 22 | 6 | 8 | 8 | 0 | 0 | 0 |
| 16 | 5 | 3 | 8 | 0 | 0 | 0 |
| 21 | 8 | 8 | 5 | 0 | 0 | 0 |
| 16 | 4 | 6 | 6 | 0 | 0 | 0 |
| 7 | 2 | 3 | 2 | 0 | 0 | 0 |
| 4 | 1 | 2 | 1 | 0 | 0 | 0 |
| 16 | 2 | 2 | 1 | 3 | 6 | 2 |
| 21 | 7 | 7 | 7 | 0 | 0 | 0 |
| 13 | 4 | 4 | 5 | 0 | 0 | 0 |
| 10 | 3 | 4 | 3 | 0 | 0 | 0 |
| 21 | 7 | 8 | 6 | 0 | 0 | 0 |
| 10 | 5 | 2 | 3 | 0 | 0 | 0 |
| 15 | 7 | 3 | 5 | 0 | 0 | 0 |
| 18 | 6 | 5 | 7 | 0 | 0 | 0 |
| 8 | 2 | 3 | 3 | 0 | 0 | 0 |
| 13 | 3 | 5 | 5 | 0 | 0 | 0 |
| 14 | 3 | 4 | 7 | 0 | 0 | 0 |
| 12 | 5 | 3 | 4 | 0 | 0 | 0 |
| 15 | 4 | 5 | 6 | 0 | 0 | 0 |
| 19 | 7 | 6 | 6 | 0 | 0 | 0 |
| 12 | 5 | 3 | 4 | 0 | 0 | 0 |
| 10 | 3 | 3 | 4 | 0 | 0 | 0 |
| 21 | 6 | 7 | 8 | 0 | 0 | 0 |
| 16 | 5 | 5 | 6 | 0 | 0 | 0 |
| 19 | 5 | 8 | 6 | 0 | 0 | 0 |
| 19 | 6 | 6 | 7 | 0 | 0 | 0 |
| 7 | 2 | 2 | 3 | 0 | 0 | 0 |
| 14 | 4 | 5 | 5 | 0 | 0 | 0 |
| 18 | 6 | 6 | 6 | 0 | 0 | 0 |
| 13 | 4 | 4 | 5 | 0 | 0 | 0 |
| 13 | 5 | 5 | 3 | 0 | 0 | 0 |
| 10 | 6 | 1 | 3 | 0 | 0 | 0 |
| 12 | 6 | 4 | 2 | 0 | 0 | 0 |
| 12 | 4 | 5 | 3 | 0 | 0 | 0 |
| 13 | 4 | 4 | 5 | 0 | 0 | 0 |
| 13 | 4 | 5 | 4 | 0 | 0 | 0 |

| | | | | | | |
|---|---|---|---|---|---|---|
| 13 | 5 | 3 | 5 | 0 | 0 | 0 |
| 11 | 5 | 3 | 3 | 0 | 0 | 0 |
| 6 | 2 | 2 | 2 | 0 | 0 | 0 |
| 12 | 3 | 4 | 5 | 0 | 0 | 0 |
| 11 | 4 | 3 | 4 | 0 | 0 | 0 |
| 4 | 1 | 2 | 1 | 0 | 0 | 0 |
| 12 | 4 | 6 | 2 | 0 | 0 | 0 |
| 14 | 5 | 4 | 5 | 0 | 0 | 0 |
| 18 | 6 | 4 | 8 | 0 | 0 | 0 |
| 15 | 5 | 5 | 5 | 0 | 0 | 0 |
| 16 | 5 | 5 | 6 | 0 | 0 | 0 |
| 14 | 7 | 3 | 4 | 0 | 0 | 0 |
| 17 | 5 | 5 | 7 | 0 | 0 | 0 |
| 11 | 3 | 2 | 6 | 0 | 0 | 0 |
| 16 | 5 | 5 | 6 | 0 | 0 | 0 |
| 8 | 2 | 3 | 3 | 0 | 0 | 0 |
| 19 | 7 | 6 | 6 | 0 | 0 | 0 |
| 15 | 4 | 6 | 5 | 0 | 0 | 0 |
| 12 | 4 | 4 | 4 | 0 | 0 | 0 |
| 10 | 4 | 3 | 3 | 0 | 0 | 0 |
| 13 | 5 | 5 | 3 | 0 | 0 | 0 |
| 7 | 3 | 2 | 2 | 0 | 0 | 0 |
| 21 | 7 | 6 | 8 | 0 | 0 | 0 |
| 9 | 4 | 3 | 2 | 0 | 0 | 0 |
| 5 | 2 | 2 | 1 | 0 | 0 | 0 |
| 13 | 4 | 4 | 5 | 0 | 0 | 0 |
| 14 | 3 | 4 | 3 | 0 | 2 | 2 |
| 11 | 3 | 3 | 5 | 0 | 0 | 0 |
| 11 | 5 | 3 | 3 | 0 | 0 | 0 |
| 9 | 3 | 3 | 3 | 0 | 0 | 0 |
| 13 | 5 | 5 | 3 | 0 | 0 | 0 |
| 8 | 3 | 2 | 3 | 0 | 0 | 0 |
| 9 | 2 | 2 | 5 | 0 | 0 | 0 |
| 11 | 3 | 3 | 5 | 0 | 0 | 0 |
| 7 | 1 | 4 | 2 | 0 | 0 | 0 |
| 15 | 6 | 4 | 5 | 0 | 0 | 0 |
| 10 | 4 | 3 | 3 | 0 | 0 | 0 |
| 14 | 3 | 4 | 7 | 0 | 0 | 0 |
| 8 | 3 | 2 | 3 | 0 | 0 | 0 |
| 8 | 4 | 2 | 2 | 0 | 0 | 0 |
| 13 | 4 | 3 | 6 | 0 | 0 | 0 |
| 8 | 3 | 2 | 3 | 0 | 0 | 0 |
| 14 | 3 | 4 | 7 | 0 | 0 | 0 |
| 13 | 3 | 4 | 6 | 0 | 0 | 0 |
| 14 | 6 | 3 | 5 | 0 | 0 | 0 |
| 14 | 6 | 4 | 4 | 0 | 0 | 0 |
| 8 | 4 | 3 | 1 | 0 | 0 | 0 |
| 11 | 3 | 4 | 4 | 0 | 0 | 0 |

| | | | | | | |
|---|---|---|---|---|---|---|
| 10 | 4 | 4 | 2 | 0 | 0 | 0 |
| 8 | 2 | 3 | 3 | 0 | 0 | 0 |
| 10 | 1 | 4 | 5 | 0 | 0 | 0 |
| 5 | 2 | 2 | 1 | 0 | 0 | 0 |
| 8 | 3 | 3 | 2 | 0 | 0 | 0 |
| 15 | 5 | 5 | 5 | 0 | 0 | 0 |
| 11 | 3 | 4 | 4 | 0 | 0 | 0 |
| 10 | 4 | 3 | 3 | 0 | 0 | 0 |
| 11 | 3 | 4 | 4 | 0 | 0 | 0 |
| 5 | 2 | 1 | 2 | 0 | 0 | 0 |
| 10 | 3 | 4 | 3 | 0 | 0 | 0 |
| 10 | 3 | 4 | 3 | 0 | 0 | 0 |
| 5 | 1 | 1 | 0 | 0 | 3 | 0 |
| 15 | 5 | 6 | 4 | 0 | 0 | 0 |
| 11 | 3 | 3 | 5 | 0 | 0 | 0 |
| 8 | 3 | 3 | 2 | 0 | 0 | 0 |
| 7 | 1 | 3 | 3 | 0 | 0 | 0 |
| 7 | 2 | 1 | 4 | 0 | 0 | 0 |
| 6 | 2 | 3 | 1 | 0 | 0 | 0 |
| 8 | 3 | 4 | 1 | 0 | 0 | 0 |
| 9 | 2 | 3 | 4 | 0 | 0 | 0 |
| 10 | 5 | 0 | 0 | 0 | 5 | 0 |
| 11 | 4 | 2 | 5 | 0 | 0 | 0 |
| 7 | 2 | 3 | 2 | 0 | 0 | 0 |
| 15 | 4 | 3 | 8 | 0 | 0 | 0 |
| 15 | 7 | 4 | 4 | 0 | 0 | 0 |
| 17 | 5 | 6 | 6 | 0 | 0 | 0 |
| 5 | 1 | 1 | 3 | 0 | 0 | 0 |
| 9 | 3 | 2 | 4 | 0 | 0 | 0 |
| 11 | 2 | 6 | 3 | 0 | 0 | 0 |
| 10 | 4 | 3 | 3 | 0 | 0 | 0 |
| 12 | 4 | 4 | 4 | 0 | 0 | 0 |
| 9 | 4 | 3 | 2 | 0 | 0 | 0 |
| 9 | 3 | 3 | 3 | 0 | 0 | 0 |
| 12 | 4 | 4 | 4 | 0 | 0 | 0 |
| 6 | 3 | 1 | 2 | 0 | 0 | 0 |
| 4 | 1 | 1 | 2 | 0 | 0 | 0 |
| 11 | 5 | 3 | 3 | 0 | 0 | 0 |
| 5 | 1 | 2 | 2 | 0 | 0 | 0 |
| 8 | 3 | 2 | 3 | 0 | 0 | 0 |
| 8 | 1 | 2 | 5 | 0 | 0 | 0 |
| 7 | 2 | 1 | 4 | 0 | 0 | 0 |
| 7 | 4 | 1 | 2 | 0 | 0 | 0 |
| 8 | 3 | 2 | 3 | 0 | 0 | 0 |
| 12 | 5 | 2 | 5 | 0 | 0 | 0 |
| 9 | 3 | 2 | 4 | 0 | 0 | 0 |
| 8 | 1 | 3 | 4 | 0 | 0 | 0 |
| 11 | 2 | 3 | 6 | 0 | 0 | 0 |

| | | | | | | |
|---|---|---|---|---|---|---|
| 11 | 3 | 3 | 3 | 0 | 1 | 1 |
| 7 | 2 | 2 | 3 | 0 | 0 | 0 |
| 7 | 2 | 3 | 2 | 0 | 0 | 0 |
| 11 | 5 | 3 | 3 | 0 | 0 | 0 |
| 8 | 4 | 2 | 2 | 0 | 0 | 0 |
| 11 | 5 | 4 | 2 | 0 | 0 | 0 |
| 17 | 6 | 4 | 7 | 0 | 0 | 0 |
| 14 | 4 | 3 | 7 | 0 | 0 | 0 |
| 6 | 2 | 2 | 2 | 0 | 0 | 0 |
| 11 | 3 | 4 | 4 | 0 | 0 | 0 |
| 12 | 4 | 3 | 5 | 0 | 0 | 0 |
| 5 | 2 | 2 | 1 | 0 | 0 | 0 |
| 8 | 4 | 2 | 2 | 0 | 0 | 0 |
| 8 | 5 | 1 | 2 | 0 | 0 | 0 |
| 14 | 5 | 5 | 4 | 0 | 0 | 0 |
| 4 | 1 | 2 | 1 | 0 | 0 | 0 |
| 13 | 5 | 3 | 5 | 0 | 0 | 0 |
| 11 | 4 | 4 | 3 | 0 | 0 | 0 |
| 14 | 4 | 5 | 5 | 0 | 0 | 0 |
| 14 | 6 | 3 | 5 | 0 | 0 | 0 |
| 11 | 2 | 0 | 1 | 0 | 8 | 0 |
| 7 | 2 | 2 | 3 | 0 | 0 | 0 |
| 6 | 2 | 2 | 2 | 0 | 0 | 0 |
| 7 | 3 | 2 | 2 | 0 | 0 | 0 |
| 10 | 3 | 3 | 4 | 0 | 0 | 0 |
| 5 | 2 | 0 | 0 | 0 | 2 | 1 |
| 10 | 2 | 4 | 4 | 0 | 0 | 0 |
| 19 | 5 | 7 | 7 | 0 | 0 | 0 |
| 8 | 2 | 2 | 2 | 0 | 2 | 0 |
| 9 | 3 | 3 | 3 | 0 | 0 | 0 |
| 9 | 2 | 4 | 3 | 0 | 0 | 0 |
| 7 | 2 | 4 | 1 | 0 | 0 | 0 |
| 6 | 1 | 1 | 4 | 0 | 0 | 0 |
| 6 | 2 | 1 | 2 | 0 | 1 | 0 |
| 12 | 3 | 5 | 4 | 0 | 0 | 0 |
| 5 | 2 | 1 | 2 | 0 | 0 | 0 |
| 10 | 2 | 1 | 4 | 0 | 2 | 1 |
| 11 | 5 | 2 | 4 | 0 | 0 | 0 |
| 7 | 1 | 2 | 1 | 1 | 0 | 2 |
| 13 | 3 | 5 | 5 | 0 | 0 | 0 |
| 9 | 2 | 4 | 3 | 0 | 0 | 0 |
| 10 | 3 | 3 | 4 | 0 | 0 | 0 |
| 5 | 1 | 3 | 1 | 0 | 0 | 0 |
| 8 | 3 | 1 | 4 | 0 | 0 | 0 |
| 9 | 3 | 2 | 4 | 0 | 0 | 0 |
| 14 | 4 | 5 | 5 | 0 | 0 | 0 |
| 9 | 3 | 2 | 2 | 0 | 2 | 0 |
| 16 | 4 | 6 | 6 | 0 | 0 | 0 |

| | | | | | | |
|---|---|---|---|---|---|---|
| 15 | 5 | 5 | 5 | 0 | 0 | 0 |
| 13 | 4 | 5 | 4 | 0 | 0 | 0 |
| 5 | 2 | 1 | 2 | 0 | 0 | 0 |
| 6 | 1 | 3 | 2 | 0 | 0 | 0 |
| 11 | 3 | 4 | 4 | 0 | 0 | 0 |
| 14 | 4 | 5 | 5 | 0 | 0 | 0 |
| 8 | 2 | 3 | 3 | 0 | 0 | 0 |
| 1 | 0 | 0 | 1 | 0 | 0 | 0 |
| 11 | 3 | 4 | 4 | 0 | 0 | 0 |
| 14 | 4 | 5 | 5 | 0 | 0 | 0 |
| 9 | 4 | 3 | 2 | 0 | 0 | 0 |
| 12 | 3 | 4 | 5 | 0 | 0 | 0 |
| 9 | 2 | 4 | 3 | 0 | 0 | 0 |
| 11 | 4 | 4 | 3 | 0 | 0 | 0 |
| 10 | 2 | 3 | 5 | 0 | 0 | 0 |
| 12 | 3 | 4 | 5 | 0 | 0 | 0 |
| 9 | 3 | 3 | 3 | 0 | 0 | 0 |
| 3 | 0 | 2 | 1 | 0 | 0 | 0 |
| 3 | 1 | 0 | 1 | 0 | 1 | 0 |
| 6 | 1 | 2 | 3 | 0 | 0 | 0 |
| 12 | 3 | 7 | 2 | 0 | 0 | 0 |
| 13 | 4 | 3 | 6 | 0 | 0 | 0 |
| 6 | 2 | 3 | 1 | 0 | 0 | 0 |
| 10 | 3 | 4 | 3 | 0 | 0 | 0 |
| 13 | 5 | 3 | 5 | 0 | 0 | 0 |
| 7 | 1 | 4 | 2 | 0 | 0 | 0 |
| 11 | 3 | 4 | 4 | 0 | 0 | 0 |
| 9 | 5 | 2 | 2 | 0 | 0 | 0 |
| 7 | 3 | 2 | 2 | 0 | 0 | 0 |
| 12 | 4 | 4 | 4 | 0 | 0 | 0 |
| 7 | 4 | 1 | 2 | 0 | 0 | 0 |
| 11 | 3 | 4 | 4 | 0 | 0 | 0 |
| 4 | 1 | 2 | 1 | 0 | 0 | 0 |
| 7 | 2 | 3 | 2 | 0 | 0 | 0 |
| 9 | 4 | 3 | 2 | 0 | 0 | 0 |
| 5 | 2 | 2 | 1 | 0 | 0 | 0 |
| 3 | 1 | 2 | 0 | 0 | 0 | 0 |
| 7 | 2 | 3 | 2 | 0 | 0 | 0 |
| 11 | 3 | 3 | 5 | 0 | 0 | 0 |
| 14 | 5 | 4 | 5 | 0 | 0 | 0 |
| 9 | 4 | 3 | 2 | 0 | 0 | 0 |
| 7 | 3 | 2 | 2 | 0 | 0 | 0 |
| 6 | 1 | 2 | 3 | 0 | 0 | 0 |
| 10 | 4 | 3 | 3 | 0 | 0 | 0 |
| 6 | 2 | 2 | 2 | 0 | 0 | 0 |
| 2 | 1 | 0 | 1 | 0 | 0 | 0 |
| 6 | 2 | 2 | 2 | 0 | 0 | 0 |
| 13 | 4 | 5 | 4 | 0 | 0 | 0 |

| | | | | | | |
|---|---|---|---|---|---|---|
| 11 | 2 | 5 | 4 | 0 | 0 | 0 |
| 5 | 1 | 1 | 1 | 0 | 2 | 0 |
| 5 | 1 | 1 | 3 | 0 | 0 | 0 |
| 10 | 2 | 3 | 5 | 0 | 0 | 0 |
| 10 | 3 | 2 | 5 | 0 | 0 | 0 |
| 9 | 4 | 2 | 3 | 0 | 0 | 0 |
| 10 | 4 | 3 | 3 | 0 | 0 | 0 |
| 13 | 4 | 4 | 5 | 0 | 0 | 0 |
| 8 | 3 | 2 | 3 | 0 | 0 | 0 |
| 6 | 2 | 2 | 2 | 0 | 0 | 0 |
| 7 | 2 | 2 | 3 | 0 | 0 | 0 |
| 9 | 4 | 2 | 3 | 0 | 0 | 0 |
| 20 | 6 | 8 | 6 | 0 | 0 | 0 |
| 4 | 1 | 1 | 2 | 0 | 0 | 0 |
| 5 | 1 | 2 | 2 | 0 | 0 | 0 |
| 4 | 2 | 1 | 1 | 0 | 0 | 0 |
| 7 | 2 | 2 | 3 | 0 | 0 | 0 |
| 4 | 1 | 2 | 1 | 0 | 0 | 0 |
| 9 | 4 | 3 | 2 | 0 | 0 | 0 |
| 6 | 2 | 2 | 2 | 0 | 0 | 0 |
| 7 | 4 | 1 | 2 | 0 | 0 | 0 |
| 8 | 2 | 2 | 4 | 0 | 0 | 0 |
| 6 | 2 | 2 | 2 | 0 | 0 | 0 |
| 8 | 3 | 2 | 3 | 0 | 0 | 0 |
| 7 | 2 | 2 | 3 | 0 | 0 | 0 |
| 5 | 2 | 1 | 2 | 0 | 0 | 0 |
| 4 | 2 | 2 | 0 | 0 | 0 | 0 |
| 7 | 2 | 2 | 1 | 0 | 2 | 0 |
| 2 | 1 | 0 | 1 | 0 | 0 | 0 |
| 5 | 2 | 1 | 2 | 0 | 0 | 0 |
| 8 | 3 | 2 | 3 | 0 | 0 | 0 |
| 8 | 2 | 3 | 3 | 0 | 0 | 0 |
| 12 | 4 | 4 | 4 | 0 | 0 | 0 |
| 8 | 0 | 0 | 0 | 0 | 8 | 0 |
| 8 | 3 | 2 | 3 | 0 | 0 | 0 |
| 8 | 3 | 4 | 1 | 0 | 0 | 0 |
| 5 | 3 | 0 | 0 | 0 | 2 | 0 |
| 13 | 4 | 4 | 5 | 0 | 0 | 0 |
| 5 | 2 | 1 | 2 | 0 | 0 | 0 |
| 13 | 3 | 3 | 7 | 0 | 0 | 0 |
| 9 | 4 | 4 | 1 | 0 | 0 | 0 |
| 12 | 3 | 5 | 4 | 0 | 0 | 0 |
| 6 | 2 | 2 | 2 | 0 | 0 | 0 |
| 4 | 1 | 2 | 1 | 0 | 0 | 0 |
| 8 | 4 | 2 | 2 | 0 | 0 | 0 |
| 14 | 4 | 5 | 5 | 0 | 0 | 0 |
| 5 | 1 | 2 | 2 | 0 | 0 | 0 |
| 9 | 3 | 4 | 2 | 0 | 0 | 0 |

| | | | | | | |
|---|---|---|---|---|---|---|
| 11 | 5 | 3 | 3 | 0 | 0 | 0 |
| 7 | 2 | 2 | 3 | 0 | 0 | 0 |
| 4 | 4 | 0 | 0 | 0 | 0 | 0 |
| 7 | 2 | 3 | 2 | 0 | 0 | 0 |
| 9 | 5 | 3 | 1 | 0 | 0 | 0 |
| 6 | 3 | 1 | 1 | 0 | 1 | 0 |
| 7 | 2 | 3 | 2 | 0 | 0 | 0 |
| 7 | 2 | 3 | 2 | 0 | 0 | 0 |
| 7 | 3 | 1 | 3 | 0 | 0 | 0 |
| 3 | 1 | 2 | 0 | 0 | 0 | 0 |
| 11 | 3 | 4 | 4 | 0 | 0 | 0 |
| 6 | 2 | 3 | 1 | 0 | 0 | 0 |
| 9 | 4 | 2 | 3 | 0 | 0 | 0 |
| 6 | 0 | 0 | 0 | 0 | 6 | 0 |
| 5 | 2 | 1 | 2 | 0 | 0 | 0 |
| 13 | 4 | 4 | 5 | 0 | 0 | 0 |
| 4 | 0 | 2 | 2 | 0 | 0 | 0 |
| 5 | 0 | 3 | 2 | 0 | 0 | 0 |
| 7 | 0 | 0 | 0 | 0 | 7 | 0 |
| 6 | 1 | 2 | 3 | 0 | 0 | 0 |
| 9 | 4 | 2 | 3 | 0 | 0 | 0 |
| 4 | 1 | 2 | 1 | 0 | 0 | 0 |
| 9 | 2 | 4 | 3 | 0 | 0 | 0 |
| 8 | 2 | 3 | 3 | 0 | 0 | 0 |
| 5 | 1 | 2 | 2 | 0 | 0 | 0 |
| 10 | 4 | 1 | 5 | 0 | 0 | 0 |
| 8 | 3 | 2 | 3 | 0 | 0 | 0 |
| 11 | 5 | 3 | 3 | 0 | 0 | 0 |
| 7 | 2 | 3 | 2 | 0 | 0 | 0 |
| 7 | 2 | 3 | 1 | 0 | 1 | 0 |
| 7 | 3 | 2 | 2 | 0 | 0 | 0 |
| 4 | 2 | 1 | 1 | 0 | 0 | 0 |
| 6 | 2 | 3 | 1 | 0 | 0 | 0 |
| 9 | 2 | 3 | 4 | 0 | 0 | 0 |
| 10 | 4 | 3 | 3 | 0 | 0 | 0 |
| 8 | 2 | 4 | 2 | 0 | 0 | 0 |
| 11 | 5 | 3 | 3 | 0 | 0 | 0 |
| 3 | 1 | 0 | 2 | 0 | 0 | 0 |
| 7 | 2 | 2 | 3 | 0 | 0 | 0 |
| 6 | 4 | 1 | 1 | 0 | 0 | 0 |
| 6 | 2 | 1 | 3 | 0 | 0 | 0 |
| 8 | 3 | 3 | 2 | 0 | 0 | 0 |
| 9 | 2 | 3 | 4 | 0 | 0 | 0 |
| 11 | 3 | 5 | 3 | 0 | 0 | 0 |
| 14 | 4 | 4 | 6 | 0 | 0 | 0 |
| 9 | 4 | 2 | 3 | 0 | 0 | 0 |
| 8 | 2 | 4 | 2 | 0 | 0 | 0 |
| 9 | 3 | 3 | 3 | 0 | 0 | 0 |

| | | | | | | |
|---|---|---|---|---|---|---|
| 9 | 2 | 2 | 5 | 0 | 0 | 0 |
| 6 | 3 | 1 | 2 | 0 | 0 | 0 |
| 4 | 1 | 1 | 2 | 0 | 0 | 0 |
| 10 | 3 | 3 | 4 | 0 | 0 | 0 |
| 5 | 2 | 2 | 1 | 0 | 0 | 0 |
| 5 | 2 | 2 | 1 | 0 | 0 | 0 |
| 6 | 1 | 1 | 1 | 0 | 3 | 0 |
| 6 | 3 | 1 | 2 | 0 | 0 | 0 |
| 8 | 2 | 3 | 3 | 0 | 0 | 0 |
| 11 | 3 | 4 | 4 | 0 | 0 | 0 |
| 6 | 1 | 2 | 3 | 0 | 0 | 0 |
| 5 | 3 | 1 | 1 | 0 | 0 | 0 |
| 7 | 3 | 2 | 2 | 0 | 0 | 0 |
| 8 | 3 | 2 | 3 | 0 | 0 | 0 |
| 5 | 1 | 1 | 3 | 0 | 0 | 0 |
| 5 | 1 | 2 | 2 | 0 | 0 | 0 |
| 8 | 3 | 2 | 3 | 0 | 0 | 0 |
| 8 | 1 | 3 | 4 | 0 | 0 | 0 |
| 6 | 2 | 2 | 2 | 0 | 0 | 0 |
| 4 | 1 | 2 | 1 | 0 | 0 | 0 |
| 14 | 4 | 6 | 4 | 0 | 0 | 0 |
| 7 | 3 | 2 | 2 | 0 | 0 | 0 |
| 3 | 1 | 1 | 1 | 0 | 0 | 0 |
| 6 | 2 | 1 | 0 | 0 | 3 | 0 |
| 6 | 2 | 3 | 1 | 0 | 0 | 0 |
| 3 | 0 | 2 | 1 | 0 | 0 | 0 |
| 12 | 1 | 0 | 0 | 0 | 11 | 0 |
| 8 | 2 | 3 | 3 | 0 | 0 | 0 |
| 4 | 1 | 2 | 1 | 0 | 0 | 0 |
| 7 | 3 | 2 | 2 | 0 | 0 | 0 |
| 9 | 3 | 3 | 3 | 0 | 0 | 0 |
| 12 | 4 | 4 | 4 | 0 | 0 | 0 |
| 3 | 1 | 1 | 1 | 0 | 0 | 0 |
| 3 | 1 | 1 | 1 | 0 | 0 | 0 |
| 6 | 1 | 2 | 3 | 0 | 0 | 0 |
| 8 | 3 | 2 | 3 | 0 | 0 | 0 |
| 9 | 4 | 1 | 4 | 0 | 0 | 0 |
| 8 | 2 | 2 | 4 | 0 | 0 | 0 |
| 5 | 1 | 2 | 2 | 0 | 0 | 0 |
| 11 | 5 | 3 | 3 | 0 | 0 | 0 |
| 5 | 2 | 1 | 2 | 0 | 0 | 0 |
| 10 | 6 | 2 | 2 | 0 | 0 | 0 |
| 3 | 1 | 0 | 0 | 0 | 2 | 0 |
| 4 | 1 | 1 | 2 | 0 | 0 | 0 |
| 7 | 2 | 2 | 3 | 0 | 0 | 0 |
| 5 | 2 | 2 | 1 | 0 | 0 | 0 |
| 7 | 3 | 3 | 1 | 0 | 0 | 0 |
| 8 | 2 | 3 | 3 | 0 | 0 | 0 |

| | | | | | | |
|---|---|---|---|---|---|---|
| 4 | 1 | 1 | 2 | 0 | 0 | 0 |
| 6 | 2 | 3 | 1 | 0 | 0 | 0 |
| 5 | 0 | 3 | 2 | 0 | 0 | 0 |
| 10 | 4 | 4 | 2 | 0 | 0 | 0 |
| 4 | 2 | 1 | 1 | 0 | 0 | 0 |
| 4 | 1 | 2 | 1 | 0 | 0 | 0 |
| 5 | 2 | 2 | 1 | 0 | 0 | 0 |
| 6 | 2 | 2 | 2 | 0 | 0 | 0 |
| 4 | 2 | 1 | 1 | 0 | 0 | 0 |
| 5 | 2 | 2 | 1 | 0 | 0 | 0 |
| 8 | 3 | 4 | 1 | 0 | 0 | 0 |
| 4 | 3 | 1 | 0 | 0 | 0 | 0 |
| 3 | 0 | 1 | 0 | 0 | 2 | 0 |
| 4 | 3 | 1 | 0 | 0 | 0 | 0 |
| 6 | 2 | 2 | 2 | 0 | 0 | 0 |
| 6 | 2 | 3 | 1 | 0 | 0 | 0 |
| 7 | 2 | 3 | 2 | 0 | 0 | 0 |
| 4 | 2 | 2 | 0 | 0 | 0 | 0 |
| 11 | 4 | 2 | 5 | 0 | 0 | 0 |
| 8 | 2 | 4 | 2 | 0 | 0 | 0 |
| 4 | 1 | 1 | 2 | 0 | 0 | 0 |
| 4 | 1 | 1 | 2 | 0 | 0 | 0 |
| 6 | 3 | 1 | 2 | 0 | 0 | 0 |
| 9 | 4 | 0 | 0 | 0 | 5 | 0 |
| 3 | 1 | 2 | 0 | 0 | 0 | 0 |
| 9 | 3 | 3 | 3 | 0 | 0 | 0 |
| 3 | 0 | 1 | 1 | 0 | 1 | 0 |
| 4 | 1 | 2 | 1 | 0 | 0 | 0 |
| 8 | 2 | 3 | 3 | 0 | 0 | 0 |
| 8 | 4 | 2 | 2 | 0 | 0 | 0 |
| 4 | 2 | 1 | 1 | 0 | 0 | 0 |
| 3 | 1 | 1 | 1 | 0 | 0 | 0 |
| 6 | 2 | 2 | 2 | 0 | 0 | 0 |
| 4 | 0 | 0 | 0 | 0 | 4 | 0 |
| 13 | 0 | 0 | 0 | 0 | 13 | 0 |
| 10 | 4 | 3 | 3 | 0 | 0 | 0 |
| 4 | 3 | 1 | 0 | 0 | 0 | 0 |
| 7 | 2 | 3 | 2 | 0 | 0 | 0 |
| 3 | 1 | 1 | 1 | 0 | 0 | 0 |
| 8 | 4 | 1 | 3 | 0 | 0 | 0 |
| 2 | 1 | 0 | 1 | 0 | 0 | 0 |
| 5 | 1 | 0 | 1 | 0 | 2 | 1 |
| 3 | 1 | 1 | 1 | 0 | 0 | 0 |
| 5 | 2 | 2 | 1 | 0 | 0 | 0 |
| 5 | 2 | 1 | 2 | 0 | 0 | 0 |
| 11 | 4 | 4 | 3 | 0 | 0 | 0 |
| 6 | 2 | 2 | 2 | 0 | 0 | 0 |
| 5 | 1 | 3 | 1 | 0 | 0 | 0 |

| | | | | | | |
|---|---|---|---|---|---|---|
| 5 | 3 | 1 | 1 | 0 | 0 | 0 |
| 8 | 2 | 2 | 4 | 0 | 0 | 0 |
| 2 | 1 | 1 | 0 | 0 | 0 | 0 |
| 4 | 2 | 1 | 1 | 0 | 0 | 0 |
| 5 | 2 | 2 | 1 | 0 | 0 | 0 |
| 7 | 3 | 2 | 2 | 0 | 0 | 0 |
| 6 | 3 | 2 | 1 | 0 | 0 | 0 |
| 6 | 3 | 2 | 1 | 0 | 0 | 0 |
| 6 | 2 | 2 | 2 | 0 | 0 | 0 |
| 7 | 1 | 3 | 3 | 0 | 0 | 0 |
| 6 | 3 | 1 | 2 | 0 | 0 | 0 |
| 3 | 1 | 2 | 0 | 0 | 0 | 0 |
| 4 | 0 | 2 | 2 | 0 | 0 | 0 |
| 7 | 2 | 2 | 3 | 0 | 0 | 0 |
| 6 | 1 | 2 | 3 | 0 | 0 | 0 |
| 6 | 3 | 2 | 1 | 0 | 0 | 0 |
| 6 | 2 | 2 | 2 | 0 | 0 | 0 |
| 7 | 3 | 2 | 1 | 0 | 1 | 0 |
| 4 | 1 | 1 | 2 | 0 | 0 | 0 |
| 7 | 2 | 2 | 3 | 0 | 0 | 0 |
| 4 | 1 | 1 | 2 | 0 | 0 | 0 |
| 7 | 3 | 2 | 2 | 0 | 0 | 0 |
| 4 | 1 | 2 | 1 | 0 | 0 | 0 |
| 9 | 3 | 3 | 3 | 0 | 0 | 0 |
| 4 | 1 | 1 | 2 | 0 | 0 | 0 |
| 9 | 3 | 4 | 2 | 0 | 0 | 0 |
| 7 | 2 | 2 | 3 | 0 | 0 | 0 |
| 4 | 1 | 2 | 1 | 0 | 0 | 0 |
| 5 | 2 | 3 | 0 | 0 | 0 | 0 |
| 4 | 2 | 0 | 2 | 0 | 0 | 0 |
| 7 | 3 | 3 | 1 | 0 | 0 | 0 |
| 4 | 1 | 2 | 1 | 0 | 0 | 0 |
| 1 | 0 | 0 | 1 | 0 | 0 | 0 |
| 1 | 1 | 0 | 0 | 0 | 0 | 0 |
| 7 | 2 | 3 | 2 | 0 | 0 | 0 |
| 7 | 3 | 2 | 2 | 0 | 0 | 0 |
| 4 | 2 | 1 | 1 | 0 | 0 | 0 |
| 5 | 3 | 1 | 1 | 0 | 0 | 0 |
| 7 | 1 | 2 | 4 | 0 | 0 | 0 |
| 7 | 3 | 1 | 3 | 0 | 0 | 0 |
| 6 | 2 | 2 | 2 | 0 | 0 | 0 |
| 7 | 2 | 3 | 2 | 0 | 0 | 0 |
| 5 | 3 | 1 | 1 | 0 | 0 | 0 |
| 8 | 3 | 2 | 3 | 0 | 0 | 0 |
| 8 | 2 | 4 | 2 | 0 | 0 | 0 |
| 9 | 4 | 3 | 2 | 0 | 0 | 0 |
| 5 | 1 | 2 | 1 | 0 | 1 | 0 |
| 4 | 2 | 1 | 1 | 0 | 0 | 0 |

| | | | | | | |
|---|---|---|---|---|---|---|
| 6 | 2 | 3 | 1 | 0 | 0 | 0 |
| 3 | 0 | 1 | 2 | 0 | 0 | 0 |
| 5 | 2 | 1 | 2 | 0 | 0 | 0 |
| 4 | 1 | 1 | 2 | 0 | 0 | 0 |
| 7 | 1 | 2 | 4 | 0 | 0 | 0 |
| 3 | 1 | 1 | 1 | 0 | 0 | 0 |
| 5 | 1 | 1 | 0 | 0 | 3 | 0 |
| 5 | 1 | 2 | 2 | 0 | 0 | 0 |
| 3 | 1 | 1 | 1 | 0 | 0 | 0 |
| 5 | 2 | 1 | 2 | 0 | 0 | 0 |
| 5 | 2 | 2 | 1 | 0 | 0 | 0 |
| 6 | 2 | 2 | 2 | 0 | 0 | 0 |
| 3 | 1 | 1 | 1 | 0 | 0 | 0 |
| 6 | 2 | 2 | 2 | 0 | 0 | 0 |
| 3 | 0 | 1 | 2 | 0 | 0 | 0 |
| 4 | 1 | 1 | 2 | 0 | 0 | 0 |
| 6 | 1 | 0 | 1 | 0 | 4 | 0 |
| 4 | 2 | 0 | 2 | 0 | 0 | 0 |
| 9 | 2 | 3 | 4 | 0 | 0 | 0 |
| 5 | 2 | 1 | 2 | 0 | 0 | 0 |
| 4 | 1 | 1 | 2 | 0 | 0 | 0 |
| 8 | 1 | 3 | 4 | 0 | 0 | 0 |
| 4 | 1 | 1 | 2 | 0 | 0 | 0 |
| 6 | 4 | 1 | 1 | 0 | 0 | 0 |
| 8 | 3 | 3 | 2 | 0 | 0 | 0 |
| 6 | 2 | 1 | 3 | 0 | 0 | 0 |
| 7 | 0 | 0 | 0 | 0 | 7 | 0 |
| 4 | 1 | 2 | 1 | 0 | 0 | 0 |
| 4 | 2 | 2 | 0 | 0 | 0 | 0 |
| 7 | 3 | 1 | 3 | 0 | 0 | 0 |
| 5 | 2 | 1 | 2 | 0 | 0 | 0 |
| 5 | 1 | 2 | 2 | 0 | 0 | 0 |
| 9 | 3 | 3 | 3 | 0 | 0 | 0 |
| 5 | 1 | 1 | 3 | 0 | 0 | 0 |
| 4 | 2 | 1 | 1 | 0 | 0 | 0 |
| 6 | 2 | 2 | 2 | 0 | 0 | 0 |
| 4 | 1 | 1 | 2 | 0 | 0 | 0 |
| 5 | 3 | 2 | 0 | 0 | 0 | 0 |
| 5 | 1 | 1 | 3 | 0 | 0 | 0 |
| 9 | 3 | 3 | 3 | 0 | 0 | 0 |
| 9 | 2 | 3 | 4 | 0 | 0 | 0 |
| 6 | 2 | 2 | 2 | 0 | 0 | 0 |
| 5 | 2 | 1 | 2 | 0 | 0 | 0 |
| 2 | 1 | 0 | 1 | 0 | 0 | 0 |
| 7 | 2 | 3 | 2 | 0 | 0 | 0 |
| 5 | 1 | 2 | 2 | 0 | 0 | 0 |
| 3 | 1 | 1 | 0 | 0 | 1 | 0 |
| 4 | 1 | 1 | 2 | 0 | 0 | 0 |

| | | | | | | |
|---|---|---|---|---|---|---|
| 6 | 2 | 2 | 2 | 0 | 0 | 0 |
| 5 | 1 | 2 | 2 | 0 | 0 | 0 |
| 5 | 1 | 1 | 3 | 0 | 0 | 0 |
| 5 | 0 | 2 | 3 | 0 | 0 | 0 |
| 7 | 2 | 0 | 0 | 0 | 5 | 0 |
| 5 | 1 | 2 | 2 | 0 | 0 | 0 |
| 6 | 2 | 1 | 1 | 0 | 2 | 0 |
| 4 | 1 | 1 | 2 | 0 | 0 | 0 |
| 4 | 1 | 1 | 2 | 0 | 0 | 0 |
| 6 | 2 | 2 | 2 | 0 | 0 | 0 |
| 6 | 1 | 3 | 2 | 0 | 0 | 0 |
| 2 | 0 | 0 | 2 | 0 | 0 | 0 |
| 3 | 1 | 1 | 1 | 0 | 0 | 0 |
| 4 | 1 | 1 | 2 | 0 | 0 | 0 |
| 5 | 2 | 2 | 1 | 0 | 0 | 0 |
| 6 | 2 | 3 | 1 | 0 | 0 | 0 |
| 7 | 1 | 3 | 3 | 0 | 0 | 0 |
| 6 | 1 | 2 | 3 | 0 | 0 | 0 |
| 6 | 2 | 2 | 2 | 0 | 0 | 0 |
| 4 | 2 | 1 | 1 | 0 | 0 | 0 |
| 2 | 1 | 1 | 0 | 0 | 0 | 0 |
| 1 | 1 | 0 | 0 | 0 | 0 | 0 |
| 6 | 2 | 2 | 2 | 0 | 0 | 0 |
| 6 | 3 | 1 | 2 | 0 | 0 | 0 |
| 2 | 1 | 0 | 1 | 0 | 0 | 0 |
| 8 | 3 | 3 | 2 | 0 | 0 | 0 |
| 4 | 2 | 1 | 1 | 0 | 0 | 0 |
| 8 | 3 | 3 | 2 | 0 | 0 | 0 |
| 5 | 2 | 2 | 1 | 0 | 0 | 0 |
| 3 | 1 | 1 | 1 | 0 | 0 | 0 |
| 3 | 0 | 2 | 1 | 0 | 0 | 0 |
| 5 | 1 | 1 | 3 | 0 | 0 | 0 |
| 5 | 2 | 2 | 1 | 0 | 0 | 0 |
| 9 | 3 | 2 | 4 | 0 | 0 | 0 |
| 5 | 2 | 3 | 0 | 0 | 0 | 0 |
| 3 | 1 | 0 | 2 | 0 | 0 | 0 |
| 7 | 3 | 2 | 2 | 0 | 0 | 0 |
| 4 | 1 | 0 | 0 | 0 | 3 | 0 |
| 3 | 1 | 0 | 2 | 0 | 0 | 0 |
| 4 | 2 | 2 | 0 | 0 | 0 | 0 |
| 4 | 2 | 0 | 2 | 0 | 0 | 0 |
| 2 | 0 | 0 | 1 | 0 | 1 | 0 |
| 4 | 2 | 1 | 1 | 0 | 0 | 0 |
| 5 | 2 | 1 | 2 | 0 | 0 | 0 |
| 12 | 4 | 4 | 4 | 0 | 0 | 0 |
| 1 | 1 | 0 | 0 | 0 | 0 | 0 |
| 4 | 2 | 1 | 1 | 0 | 0 | 0 |
| 4 | 1 | 2 | 1 | 0 | 0 | 0 |

| | | | | | | |
|---|---|---|---|---|---|---|
| 6 | 1 | 2 | 3 | 0 | 0 | 0 |
| 5 | 2 | 2 | 1 | 0 | 0 | 0 |
| 4 | 2 | 0 | 2 | 0 | 0 | 0 |
| 2 | 1 | 0 | 1 | 0 | 0 | 0 |
| 5 | 2 | 1 | 2 | 0 | 0 | 0 |
| 8 | 2 | 2 | 4 | 0 | 0 | 0 |
| 5 | 2 | 1 | 2 | 0 | 0 | 0 |
| 4 | 1 | 1 | 2 | 0 | 0 | 0 |
| 6 | 2 | 2 | 2 | 0 | 0 | 0 |
| 2 | 1 | 1 | 0 | 0 | 0 | 0 |
| 6 | 2 | 2 | 2 | 0 | 0 | 0 |
| 3 | 0 | 2 | 1 | 0 | 0 | 0 |
| 4 | 1 | 1 | 1 | 0 | 1 | 0 |
| 4 | 1 | 2 | 1 | 0 | 0 | 0 |
| 6 | 2 | 3 | 1 | 0 | 0 | 0 |
| 2 | 2 | 0 | 0 | 0 | 0 | 0 |
| 4 | 1 | 1 | 2 | 0 | 0 | 0 |
| 3 | 1 | 1 | 1 | 0 | 0 | 0 |
| 3 | 1 | 0 | 2 | 0 | 0 | 0 |
| 5 | 2 | 1 | 2 | 0 | 0 | 0 |
| 3 | 2 | 1 | 0 | 0 | 0 | 0 |
| 4 | 0 | 2 | 2 | 0 | 0 | 0 |
| 4 | 2 | 0 | 0 | 0 | 2 | 0 |
| 3 | 1 | 1 | 1 | 0 | 0 | 0 |
| 5 | 1 | 2 | 2 | 0 | 0 | 0 |
| 5 | 2 | 1 | 2 | 0 | 0 | 0 |
| 2 | 0 | 1 | 1 | 0 | 0 | 0 |
| 1 | 1 | 0 | 0 | 0 | 0 | 0 |
| 7 | 3 | 1 | 3 | 0 | 0 | 0 |
| 6 | 2 | 2 | 2 | 0 | 0 | 0 |
| 3 | 1 | 1 | 1 | 0 | 0 | 0 |
| 3 | 1 | 1 | 1 | 0 | 0 | 0 |
| 4 | 2 | 1 | 1 | 0 | 0 | 0 |
| 5 | 1 | 2 | 2 | 0 | 0 | 0 |
| 2 | 0 | 1 | 1 | 0 | 0 | 0 |
| 3 | 1 | 1 | 1 | 0 | 0 | 0 |
| 4 | 1 | 0 | 2 | 0 | 1 | 0 |
| 7 | 3 | 2 | 2 | 0 | 0 | 0 |
| 3 | 1 | 1 | 1 | 0 | 0 | 0 |
| 4 | 1 | 1 | 2 | 0 | 0 | 0 |
| 3 | 0 | 1 | 2 | 0 | 0 | 0 |
| 6 | 2 | 1 | 3 | 0 | 0 | 0 |
| 4 | 1 | 2 | 1 | 0 | 0 | 0 |
| 2 | 0 | 0 | 0 | 0 | 2 | 0 |
| 7 | 2 | 4 | 1 | 0 | 0 | 0 |
| 5 | 2 | 1 | 2 | 0 | 0 | 0 |
| 3 | 0 | 2 | 1 | 0 | 0 | 0 |
| 2 | 0 | 0 | 2 | 0 | 0 | 0 |

| | | | | | | |
|---|---|---|---|---|---|---|
| 6 | 2 | 2 | 2 | 0 | 0 | 0 |
| 6 | 2 | 2 | 2 | 0 | 0 | 0 |
| 3 | 1 | 2 | 0 | 0 | 0 | 0 |
| 6 | 2 | 2 | 2 | 0 | 0 | 0 |
| 4 | 2 | 1 | 1 | 0 | 0 | 0 |
| 1 | 0 | 0 | 0 | 0 | 1 | 0 |
| 3 | 0 | 1 | 2 | 0 | 0 | 0 |
| 5 | 1 | 2 | 2 | 0 | 0 | 0 |
| 1 | 0 | 0 | 1 | 0 | 0 | 0 |
| 4 | 1 | 1 | 2 | 0 | 0 | 0 |
| 1 | 1 | 0 | 0 | 0 | 0 | 0 |
| 5 | 2 | 2 | 1 | 0 | 0 | 0 |
| 4 | 1 | 1 | 2 | 0 | 0 | 0 |
| 3 | 0 | 1 | 2 | 0 | 0 | 0 |
| 3 | 1 | 1 | 1 | 0 | 0 | 0 |
| 1 | 1 | 0 | 0 | 0 | 0 | 0 |
| 3 | 1 | 1 | 1 | 0 | 0 | 0 |
| 4 | 0 | 2 | 2 | 0 | 0 | 0 |
| 5 | 1 | 1 | 3 | 0 | 0 | 0 |
| 3 | 1 | 1 | 1 | 0 | 0 | 0 |
| 4 | 1 | 2 | 1 | 0 | 0 | 0 |
| 2 | 1 | 0 | 1 | 0 | 0 | 0 |
| 2 | 1 | 0 | 0 | 0 | 1 | 0 |
| 2 | 0 | 1 | 1 | 0 | 0 | 0 |
| 4 | 2 | 1 | 1 | 0 | 0 | 0 |
| 3 | 2 | 0 | 1 | 0 | 0 | 0 |
| 4 | 1 | 1 | 2 | 0 | 0 | 0 |
| 5 | 2 | 1 | 2 | 0 | 0 | 0 |
| 4 | 2 | 0 | 1 | 0 | 1 | 0 |
| 4 | 1 | 2 | 1 | 0 | 0 | 0 |
| 2 | 0 | 2 | 0 | 0 | 0 | 0 |
| 7 | 0 | 0 | 0 | 0 | 7 | 0 |
| 2 | 2 | 0 | 0 | 0 | 0 | 0 |
| 3 | 1 | 0 | 2 | 0 | 0 | 0 |
| 4 | 2 | 1 | 1 | 0 | 0 | 0 |
| 2 | 1 | 0 | 1 | 0 | 0 | 0 |
| 5 | 2 | 2 | 1 | 0 | 0 | 0 |
| 4 | 1 | 2 | 1 | 0 | 0 | 0 |
| 3 | 1 | 1 | 1 | 0 | 0 | 0 |
| 4 | 1 | 0 | 0 | 0 | 3 | 0 |
| 5 | 2 | 1 | 2 | 0 | 0 | 0 |
| 3 | 1 | 2 | 0 | 0 | 0 | 0 |
| 3 | 1 | 1 | 1 | 0 | 0 | 0 |
| 4 | 1 | 2 | 1 | 0 | 0 | 0 |
| 3 | 1 | 1 | 1 | 0 | 0 | 0 |
| 3 | 0 | 2 | 1 | 0 | 0 | 0 |
| 3 | 1 | 1 | 1 | 0 | 0 | 0 |
| 4 | 1 | 1 | 2 | 0 | 0 | 0 |

| | | | | | | |
|---|---|---|---|---|---|---|
| 3 | 1 | 0 | 2 | 0 | 0 | 0 |
| 5 | 2 | 1 | 2 | 0 | 0 | 0 |
| 5 | 1 | 2 | 2 | 0 | 0 | 0 |
| 4 | 2 | 1 | 1 | 0 | 0 | 0 |
| 4 | 1 | 2 | 1 | 0 | 0 | 0 |
| 3 | 1 | 1 | 1 | 0 | 0 | 0 |
| 11 | 5 | 4 | 2 | 0 | 0 | 0 |
| 5 | 2 | 2 | 1 | 0 | 0 | 0 |
| 1 | 0 | 0 | 1 | 0 | 0 | 0 |
| 3 | 2 | 1 | 0 | 0 | 0 | 0 |
| 3 | 0 | 2 | 1 | 0 | 0 | 0 |
| 2 | 0 | 2 | 0 | 0 | 0 | 0 |
| 3 | 1 | 1 | 1 | 0 | 0 | 0 |
| 3 | 1 | 1 | 1 | 0 | 0 | 0 |
| 1 | 0 | 0 | 1 | 0 | 0 | 0 |
| 4 | 1 | 2 | 1 | 0 | 0 | 0 |
| 4 | 3 | 0 | 0 | 0 | 1 | 0 |
| 1 | 1 | 0 | 0 | 0 | 0 | 0 |
| 5 | 2 | 1 | 2 | 0 | 0 | 0 |
| 2 | 1 | 0 | 1 | 0 | 0 | 0 |
| 4 | 0 | 0 | 0 | 1 | 2 | 1 |
| 6 | 2 | 3 | 1 | 0 | 0 | 0 |
| 2 | 0 | 1 | 1 | 0 | 0 | 0 |
| 3 | 1 | 1 | 1 | 0 | 0 | 0 |
| 6 | 2 | 2 | 2 | 0 | 0 | 0 |
| 4 | 1 | 2 | 1 | 0 | 0 | 0 |
| 5 | 0 | 0 | 0 | 0 | 5 | 0 |
| 4 | 1 | 1 | 2 | 0 | 0 | 0 |
| 4 | 1 | 1 | 2 | 0 | 0 | 0 |
| 2 | 0 | 0 | 2 | 0 | 0 | 0 |
| 6 | 2 | 2 | 2 | 0 | 0 | 0 |
| 5 | 1 | 1 | 3 | 0 | 0 | 0 |
| 4 | 1 | 1 | 2 | 0 | 0 | 0 |
| 2 | 0 | 1 | 1 | 0 | 0 | 0 |
| 4 | 1 | 2 | 1 | 0 | 0 | 0 |
| 4 | 1 | 1 | 2 | 0 | 0 | 0 |
| 4 | 2 | 0 | 2 | 0 | 0 | 0 |
| 1 | 0 | 1 | 0 | 0 | 0 | 0 |
| 4 | 1 | 1 | 2 | 0 | 0 | 0 |
| 3 | 1 | 1 | 1 | 0 | 0 | 0 |
| 2 | 1 | 0 | 0 | 0 | 1 | 0 |
| 4 | 0 | 0 | 0 | 0 | 4 | 0 |
| 7 | 2 | 1 | 4 | 0 | 0 | 0 |
| 3 | 1 | 1 | 1 | 0 | 0 | 0 |
| 3 | 1 | 0 | 2 | 0 | 0 | 0 |
| 3 | 1 | 1 | 1 | 0 | 0 | 0 |
| 2 | 0 | 0 | 2 | 0 | 0 | 0 |
| 4 | 1 | 1 | 2 | 0 | 0 | 0 |

| | | | | | | |
|---|---|---|---|---|---|---|
| 6 | 2 | 1 | 3 | 0 | 0 | 0 |
| 2 | 1 | 0 | 1 | 0 | 0 | 0 |
| 5 | 2 | 1 | 2 | 0 | 0 | 0 |
| 2 | 1 | 1 | 0 | 0 | 0 | 0 |
| 3 | 0 | 0 | 3 | 0 | 0 | 0 |
| 2 | 2 | 0 | 0 | 0 | 0 | 0 |
| 7 | 2 | 3 | 2 | 0 | 0 | 0 |
| 2 | 1 | 1 | 0 | 0 | 0 | 0 |
| 1 | 0 | 1 | 0 | 0 | 0 | 0 |
| 6 | 2 | 2 | 2 | 0 | 0 | 0 |
| 3 | 1 | 1 | 1 | 0 | 0 | 0 |
| 2 | 1 | 1 | 0 | 0 | 0 | 0 |
| 2 | 2 | 0 | 0 | 0 | 0 | 0 |
| 2 | 0 | 0 | 2 | 0 | 0 | 0 |
| 4 | 2 | 1 | 1 | 0 | 0 | 0 |
| 4 | 2 | 1 | 1 | 0 | 0 | 0 |
| 3 | 0 | 0 | 0 | 0 | 3 | 0 |
| 4 | 1 | 2 | 1 | 0 | 0 | 0 |
| 3 | 1 | 1 | 1 | 0 | 0 | 0 |
| 2 | 1 | 1 | 0 | 0 | 0 | 0 |
| 2 | 0 | 2 | 0 | 0 | 0 | 0 |
| 2 | 1 | 0 | 1 | 0 | 0 | 0 |
| 3 | 1 | 2 | 0 | 0 | 0 | 0 |
| 2 | 2 | 0 | 0 | 0 | 0 | 0 |
| 4 | 1 | 0 | 1 | 0 | 2 | 0 |
| 3 | 2 | 0 | 1 | 0 | 0 | 0 |
| 2 | 0 | 0 | 2 | 0 | 0 | 0 |
| 5 | 2 | 1 | 1 | 0 | 1 | 0 |
| 4 | 2 | 1 | 1 | 0 | 0 | 0 |
| 3 | 0 | 1 | 2 | 0 | 0 | 0 |
| 2 | 1 | 1 | 0 | 0 | 0 | 0 |
| 4 | 1 | 1 | 1 | 0 | 1 | 0 |
| 3 | 1 | 0 | 2 | 0 | 0 | 0 |
| 5 | 1 | 2 | 2 | 0 | 0 | 0 |
| 3 | 1 | 0 | 0 | 0 | 2 | 0 |
| 2 | 2 | 0 | 0 | 0 | 0 | 0 |
| 3 | 0 | 1 | 2 | 0 | 0 | 0 |
| 3 | 1 | 1 | 1 | 0 | 0 | 0 |
| 2 | 1 | 0 | 1 | 0 | 0 | 0 |
| 2 | 0 | 0 | 1 | 0 | 1 | 0 |
| 2 | 2 | 0 | 0 | 0 | 0 | 0 |
| 2 | 0 | 0 | 0 | 0 | 2 | 0 |
| 3 | 1 | 1 | 1 | 0 | 0 | 0 |
| 2 | 0 | 0 | 0 | 0 | 2 | 0 |
| 3 | 2 | 1 | 0 | 0 | 0 | 0 |
| 4 | 0 | 2 | 2 | 0 | 0 | 0 |
| 3 | 1 | 1 | 1 | 0 | 0 | 0 |
| 2 | 1 | 1 | 0 | 0 | 0 | 0 |

| | | | | | | |
|---|---|---|---|---|---|---|
| 2 | 1 | 0 | 0 | 0 | 1 | 0 |
| 4 | 3 | 1 | 0 | 0 | 0 | 0 |
| 2 | 1 | 0 | 1 | 0 | 0 | 0 |
| 3 | 0 | 2 | 1 | 0 | 0 | 0 |
| 2 | 1 | 1 | 0 | 0 | 0 | 0 |
| 3 | 2 | 0 | 1 | 0 | 0 | 0 |
| 6 | 1 | 2 | 3 | 0 | 0 | 0 |
| 4 | 2 | 1 | 1 | 0 | 0 | 0 |
| 5 | 2 | 1 | 2 | 0 | 0 | 0 |
| 2 | 0 | 0 | 0 | 0 | 2 | 0 |
| 3 | 2 | 0 | 1 | 0 | 0 | 0 |
| 3 | 1 | 0 | 2 | 0 | 0 | 0 |
| 2 | 0 | 1 | 1 | 0 | 0 | 0 |
| 3 | 1 | 0 | 0 | 0 | 2 | 0 |
| 2 | 0 | 1 | 1 | 0 | 0 | 0 |
| 3 | 1 | 0 | 2 | 0 | 0 | 0 |
| 4 | 2 | 1 | 1 | 0 | 0 | 0 |
| 2 | 1 | 0 | 1 | 0 | 0 | 0 |
| 6 | 1 | 2 | 3 | 0 | 0 | 0 |
| 3 | 1 | 1 | 1 | 0 | 0 | 0 |
| 2 | 1 | 1 | 0 | 0 | 0 | 0 |
| 1 | 0 | 0 | 1 | 0 | 0 | 0 |
| 3 | 0 | 1 | 1 | 0 | 1 | 0 |
| 3 | 0 | 0 | 3 | 0 | 0 | 0 |
| 6 | 2 | 2 | 2 | 0 | 0 | 0 |
| 5 | 1 | 2 | 1 | 0 | 1 | 0 |
| 2 | 1 | 1 | 0 | 0 | 0 | 0 |
| 2 | 0 | 0 | 0 | 0 | 2 | 0 |
| 2 | 0 | 0 | 2 | 0 | 0 | 0 |
| 2 | 1 | 0 | 1 | 0 | 0 | 0 |
| 4 | 1 | 1 | 2 | 0 | 0 | 0 |
| 4 | 2 | 2 | 0 | 0 | 0 | 0 |

**Supplemental Table 2.** GO Terms. DAVID results of gene ontology analysis of proteins identified as enriched in the antiVR condition compared to IgG control.

| DAVID Database | GO Term | Figure Label |
|---|---|---|
| GOTERM_CC_DIRECT | GO:0015935 | small ribosomal subunit |
| GOTERM_CC_DIRECT | GO:0005763 | mitochondrial small ribosomal subunit |
| GOTERM_CC_DIRECT | GO:0022626 | cytosolic ribosome |
| GOTERM_CC_DIRECT | GO:0022625 | cytosolic large ribosomal subunit |
| GOTERM_CC_DIRECT | GO:0022627 | cytosolic small ribosomal subunit |
| GOTERM_CC_DIRECT | GO:0005840 | ribosome |
| GOTERM_MF_DIRECT | GO:0003735 | structural constituent of ribosome |
| GOTERM_CC_DIRECT | GO:0015934 | large ribosomal subunit |
| GOTERM_BP_DIRECT | GO:0042254 | ribosome biogenesis |
| GOTERM_CC_DIRECT | GO:0005762 | mitochondrial large ribosomal subunit |
| GOTERM_BP_DIRECT | GO:0032543 | mitochondrial translation |
| GOTERM_BP_DIRECT | GO:0006412 | translation |
| GOTERM_BP_DIRECT | GO:0006413 | translational initiation |
| GOTERM_CC_DIRECT | GO:0072686 | mitotic spindle |
| GOTERM_BP_DIRECT | GO:0000278 | mitotic cell cycle |
| GOTERM_MF_DIRECT | GO:0036402 | proteasome-activating ATPase activity |
| GOTERM_CC_DIRECT | GO:0005838 | proteasome regulatory particle |
| GOTERM_CC_DIRECT | GO:0000502 | proteasome complex |
